# A whole-body micro-CT scan library that captures the skeletal diversity of Lake Malawi cichlid fishes

**DOI:** 10.1101/2023.11.01.565162

**Authors:** Callum V. Bucklow, Martin J. Genner, George F. Turner, James Maclaine, Roger Benson, Berta Verd

## Abstract

Here we describe a dataset of freely available, readily processed, whole-body *μ*CT-scans of 56 species (116 specimens) of Lake Malawi cichlid fishes that captures a considerable majority of the morphological variation present in this remarkable adaptive radiation. We contextualise the scanned specimens within a discussion of their respective ecomorphological groupings and suggest possible macroevolutionary studies that could be conducted with these data. We also describe a methodology to efficiently *μ*CT-scan (on average) 23 specimens per hour, limiting scanning time and alleviating the financial cost whilst maintaining high resolution. We demonstrate the utility of this method by reconstructing 3D models of multiple bones from multiple specimens within the dataset. We hope this dataset will enable further morphological study of this fascinating system and permit wider-scale comparisons with other cichlid adaptive radiations.

## Background & Summary

Cichlids are one of the most speciose families of vertebrates, with over 1000 species in the African Rift Valley alone^1,2^. Multiple, independent, adaptive radiations of these fishes have evolved in the Great Lakes of East Africa, their associated satellite water bodies, as well as their connecting riverine systems. The radiations of these fishes (Subfamily: Pseudocrenilabrinae^3^), particularly those associated with Lakes Malawi, Victoria and Tanganyika, have become powerful models for the study of macroevolutionary processes^4–10^, behaviour and physiology^11–15^, and have emerged more recently as models in evolutionary developmental biology^16–19^.

Lake Malawi haplochromine cichlids represent a particularly speciose, phenotypically diverse but genetically homogeneous adaptive radiation of lacustrine fishes. This diversity, comprising approximately 850 species of maternal mouthbrooders, is the most extensive adaptive radiation of vertebrates so far identified^1,9^. Molecular clock analyses estimate the radiation to be approximately 800 thousand years old^4^, a relatively young radiation when compared to the older system of Lake Tanganyika (∼10myr) which contains just 250 species^7,20^. Despite their high phenotypic diversity, genetic variation in Lake Malawi cichlids is extremely low. Whole genomic comparisons of representatives from all seven distinct ecomorphological groups within Lake Malawi, estimated an average DNA sequence divergence of just 0.19-0.27%^4^ – a range comparable to that within human populations^6^. In addition, a relatively low DNA mutation rate; that alone cannot account for the estimated divergence time of Lake Malawi cichlids^4,6^, overlapping distributions of interspecific pairwise sequence differences and heterozygosity (intraspecific genetic variation)^4^ only further complicates this enigmatic adaptive radiation.

East African cichlids, including those belonging to the Lake Malawi radiation, have recently emerged as powerful models in evolutionary developmental biology^17^. Evolutionary modification of embryological mechanisms drives the evolution of novel adaptations and requires genetic variation^21^. Thus, comparing the embryological development of cichlids, which have limited genetic variation, can enable us to identify specific cases where evolution has modified developmental mechanisms. The diversity of feeding habits of Lake Malawi cichlids, and the ability to causally link morphological differences in craniofacial morphology to these ecological niches, has enabled integrative genetic and morphological studies examining the evolution of these traits^22–25^. More recent studies have expanded the scope beyond craniofacial phenotypes, including pigmentation patterning^26–28^; body and fin shape^19,29^ and axial elongation^18^. In parallel, aided by developments in whole-genome sequencing technologies^6^, it has been possible to considerably improve our understanding of the phylogenetic relationships among Lake Malawi cichlids^4,9,30,31^. Previously intractable macroevolutionary studies, such as the convergent evolution of hypertrophied lips^32^ can now take advantage of the relatively robust phylogenies based on whole-genome sequences. Moreover, there are now opportunities to use this new phylogenetic information to focus on the evolution of other traits, such as the axial and appendicular skeleton, that is of key importance in teleost diversification^33,34^. However, a whole-body *μ*CT-scan dataset of Lake Malawi cichlid fishes has not yet been described.

Here we present a new database of high-resolution X-ray micro-computed tomography (*μ*CT) scans of Lake Malawi cichlids, providing 3D data on skeletal morphology for the whole body of 56 species across 26 genera. In total these data comprise 116 individuals from seven recognized ecomorphological groupings^4^ (Table 1), contrasting in multiple aspects of morphology, size, behaviour, and habitat preference. We demonstrate the resolution and utility of our dataset by illustrating 3D whole-body renderings of several species, and of several skeletal regions of interest. The data will be useful resource for researchers interested in the emergence of morphological variation, including the macroevolutionary patterns common to adaptive radiations^35,36^. We argue that where digitisation efforts are being taken to characterise adult cichlid morphology, whole-body scans should be standard to ensure that other sources of morphological variation can be investigated. We propose methods that minimise scanning time, thus alleviating financial and time constraints of scanning large numbers of specimens, whilst maintaining sufficiently high resolution for macroevolutionary studies and geometric morphometric analyses of morphological variation.

**Table 1.** Genera and species represented within the dataset. The number of genera and species for each ecomorphological group are the same as those used in Malinsky et al., 2018^4^. \**Astatotilapia calliptera, Astatotilapia* sp. ‘Ruaha blue’ and *Astatotilapia gigliolli*. ^†^If considering the addition of *Lethrinops albus* and *Lethrinops auritus* that cluster within the ‘Shallow Benthics’. ^‡^Includes *Pallidochromis*. ^§^Includes *Copadichromis trimaculatus* which clusters within the shallow benthics in the phylogeny depicted in Figure 1.

| Ecomorphology | Genera Sampled | Genera Sampled (%) | Species Sampled | Species Sampled (%) |
| --- | --- | --- | --- | --- |
| *Astatotilapia (calliptera)* | 1 of 1 | 100% | 3* | N/A |
| Deep Benthic | 2 of 5 | 40% | 3 (5<sup>†</sup>) of 150 | 2.00% (3.33%) |
| *Diplotaxodon* | 2 of 2<sup>‡</sup> | 100% | 9 of 19 | 47.37% |
| Mbuna | 7 of 12 | 58.33% | 7 of 328 | 2.13% |
| *Rhamphochromis* | 1 of 1 | 100% | 10 of 14 | 71.43% |
| Shallow Benthic | 12 of 32 | 37.50% | 20 of 287 | 6.97% |
| Utaka | 1 of 1 | 100% | 4<sup>§</sup> of 55 | 7.27% |
| **Total** | **26 of 54** | **48.15%** | **56 of 856** | **6.54%** |

## Methods

### Sample Selection

There are an estimated 850 species of Lake Malawi cichlid fishes^4^. Many of which have not been described, preserved in museum collections or are available on phylogenies based on whole genome evidence. Therefore, to maximise the utility and morphological variation captured by our dataset, we focused on species present in published phylogenies^4,9,24^, and sought to include as many genera as possible in dataset (see Table 1). We prioritised scanning the type species for genera, and avoided inclusion of species which already had whole body scans freely available online. Our specimens were sourced from the collections at the Natural History Museum in London (NHMUK), from the School of Biological Sciences of the University of Bristol (Martin Genner) and from the School of Natural Sciences of Bangor University (George Turner). In total we scanned 116 specimens from 56 species (Supplementary Table 1). Of these 99 were wild-caught, and 17 were laboratory-reared. These laboratory-reared fish included *Astatotilapia calliptera* (Mbaka River, n=10), *Maylandia zebra* (Boadzulu island, n=5) and *Rhamphochromis* sp. ‘Chilingali’ (n=2), all of which died naturally or were euthanised by anaesthetic overdose [Schedule 1; Animals (Scientific Procedures) Act 1986].

### *μ*CT-Scanning

Since there was already a large collection of specimens present at the Natural History Museum in London (NHMUK) and in the extensive research collections of Martin J Genner (School of Biological Sciences, University of Bristol) and George F Turner (School of Natural Sciences, Bangor University) we decided to take advantage of the scanners present in the CT facility of NHMUK and at the XTM Facility based in the Paleobiology Research Group at the University of Bristol. Of the total 116 individuals scanned (56 species), 56 specimens (28 species) were scanned at the NHMUK Imaging and Analysis Centre and 60 specimens (28 species) were scanned at the XTM Facility at the University of Bristol. A flowchart, describing all the necessary decisions and required processing steps is provided as Figure 2.

**Figure 1.**
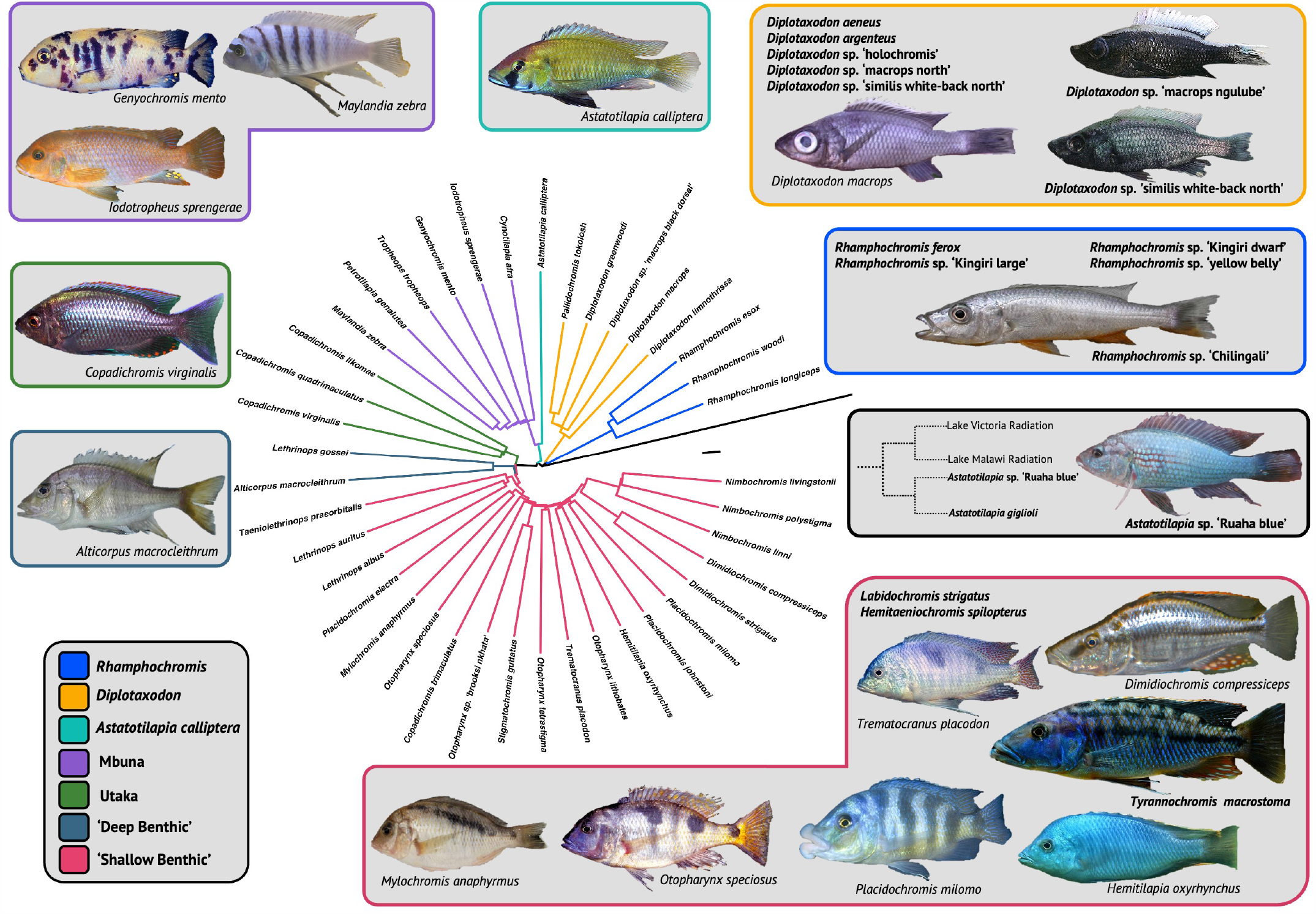
A summary of the *μ*CT-scan dataset. We were able to sample species from all seven ecomorphological groups in the Lake Malawi haplochromine radiation. The phylogenetic relationships between the majority of the species scanned is indicated and coloured according to the respective ecomorphology. The tree is a pruned version of the full (no intermediates) neighbour-joining tree published by Malinsky et al.^4^, which is rooted to *Neolamprologous brichardi*, a non-haplochromine cichlid endemic to Lake Tanganyika^20^. A cladogram depicting the relationship between the Lake Victoria, Lake Malawi and the *Astatotilapia* species native to the Great Ruaha River is indicated in the black box. Longer terminal branches reflect a higher ratio of within-species to between-species variation. We also scanned 18 species of cichlid whose phylogenetic relationships are not resolved in the phylogeny shown. The names of these species, most of which are undescribed, are indicated in their respective ecomorphological group in bold. Pictures (not to scale) of example species belonging to each ecomorphological group are also shown. Black bar: 2 *×* 10^−4^ substitutions per base pair.

**Figure 2.**
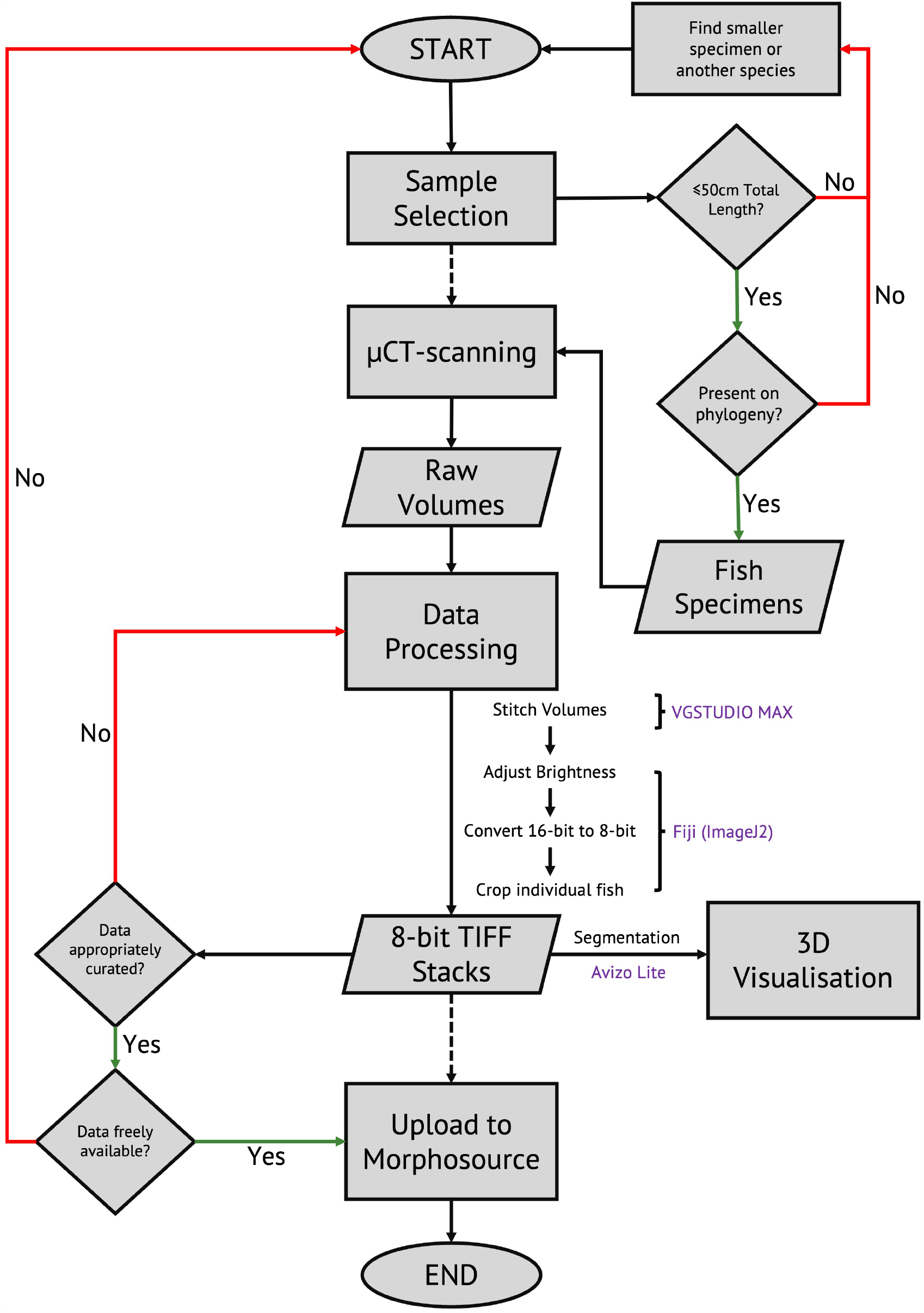
Flowchart of *μ*CT-scanning, image processing and segmentation methodology. The flowchart outlines the necessary decisions that were made during collation of the described *μ*-CT scan dataset. Rectangles represent processes; parallelograms represent inputs or outputs; diamonds represent decisions. It is sufficiently generalised that it can be reused for future data collection. We were focused on generating data for a specific macroevolutionary study, so we restricted the dataset to species with known phylogenetic placements. Software associated with data processing steps are indicated in purple.

#### Scanning Arrangement

To maximise the utility of our time and the number of species scanned, multiple specimens were scanned in each individual scan (3). Each batch of specimens was fit to the width of the scan field of view to maximise resolution, and multiple scans were conducted along a the vertical axis in order to scan the full body length of each specimen. Practically, this meant specimens were arranged into batches of similar total length. Batch sizes varied between two and five specimens, with the number of each batch ultimately dependent upon the overall size of the specimens within the batch. Of the 32 batches scanned: 20 were comprised of four specimens; nine of three specimens, and two and one batch(es) of one and five specimens, respectively. Since multiple individuals of different species were often scanned together (Figure 3A, B) and it was critical that individuals of the same species could be readily identified. Therefore, unique, low density objects such as plastic bricks, pipette tips and rubber bands were placed in physical proximity to each specimen, to act as recognisable markers (Figure 3C) which would readily resolve in the reconstructed image stacks (see Post-Scanning Processing). These objects were attached to each individual specimen-containing bag, and the specimens were bundled together, ideally ensuring that the objects faced outwards (Figure 3D-F). Specimens were tightly packed into plastic containers, sealed with tape, and allowed to rest upright (head-up) for at least ten minutes so the contents could settle to prevent movement during scanning (Figure 3G-K). We were able to scan, on average, 23 specimens per hour at maximal efficiency, an efficiency that was primarily the result of having two people per scanning visit (one scanning and one packing). The scanning rate could be further increased by packaging specimens in advance of the scanning so that each batch can be scanned continuously.

**Figure 3.**
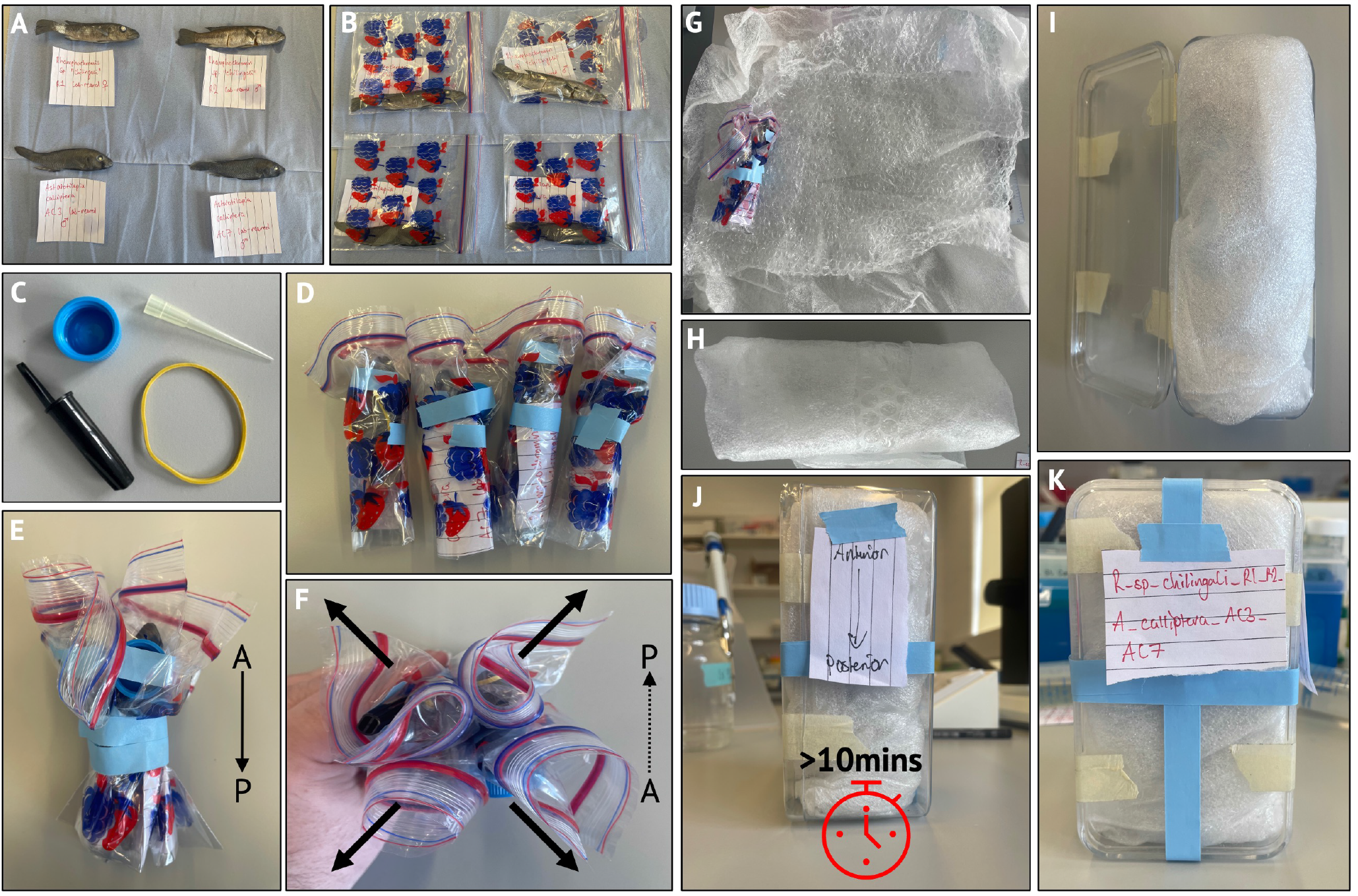
Specimen preparation for *μ*CT-scanning. Multiple fish were scanned at the same time. (A-B) Individual fish were labelled and placed in separate plastic bags (so they could be correctly identified and correctly stored after imaging). Unique objects (C) that would be readily identifiable were attached to the outside of these bags, ideally close to the heads, positioned outwards (F, arrows), and bundled together with tape (D-F) all with the same orientation (head-up). Bundles were then wrapped in bubble wrap and other packaging material and tightly sealed inside a plastic container, again head-up. Containers were left for at least ten minutes to settle to prevent movement during scanning.

#### Scanning Procedure

Prior to scanning, a visual inspection of the specimens was made following X-ray exposure to check for internal damage (such as damage to the vertebral column). If present, and possible, specimens were switched for another specimen of the same batch. To maximise scan resolution, each batch of fish was scanned using the full width of the scanner field of view. This necessitates that multiple, overlapping scans were conducted along the vertical axis of the scanner (Z-axis) in order to capture the full body length of each fish. The overlapping scans were subsequently stitched together in the processing stage using the software, VGStudio Max 3.2.5 64-bit (see below). The maximise efficiency, individuals with similar body lengths were scanned together (see above). The fish scanned ranged in standard length of 3.8cm to 33cm, which meant that scanning parameters varied between batches. The number of projections generated range from 901 to 2001. Exposure times varied between 250 and 500 seconds, with a median of 354 seconds. Similarly, power varied greatly between scans, ranging between 12.48 and 37.995 watts (mode 37.44W). All scanning parameters, for each batch conducted, as well as for each individual scan can be found in the metadata supplied in Supplementary Table 1.

### Post-Scanning Processing

All processing and segmentation was conducted on a machine specifically built for image analysis, with the following specifications: 2 x Intel^®^ Xeon^®^ CPU-ES-2640 @2.60GHz, 2601MHz, 8 Core(s) processors, 128GB of dedicated DDR3 RAM running on Microsoft Windows 10 Pro (Build Number: 10.0.19045). Raw, isometric volumes generated from the CT-scanning were imported into VGStudio Max 3.2.5 64-bit and anterior and posterior halves were subsequently stitched together by defining overlapping regions of interest in both the anterior, middle (if applicable) and posterior volumes. 16-bit tiff stacks were exported from VGStudio Max 3.2.5 64-bit and imported into FIJI^37^, a GUI for ImageJ^38^. In FIJI, individual fish were cropped out of the 16-bit stacks, which were identifiable due to the unique objects associated with each individual (see above). The brightness was adjusted by extending the distribution of pixel values to remove 0 values, and the tiff stacks were converted into 8-bit to decrease file size. Total stack file size was further decreased by removing images at the beginning and end of the stacks that did not contain readily identifiable bone.

### Data Availability

Cropped 8-bit tiff stacks of all 116 specimens have been deposited into a dedicated project on Morphosource, accessible here, where they are freely available and can be downloaded by anyone. We provide a short discussion about segmenting, including methods to optimise this these reconstructed image stacks below. We politely request that a direct reference be made to this paper if any scans from our dataset are used in future analyses. See Supplementary Table 1 for a breakdown of all scanned specimens.

## Data Records

Seven major ecomorphological groupings make up the Lake Malawi haplochromine radiation. The *Diplotaxodon* and *Rhamphochromis* groups are comprised of elongate piscivores and zooplanktivores adapted to the deeper and more open water niches within Lake Malawi^39,40^. The ‘shallow benthic’ group is speciose set of Lake Malawi cichlids^41^, comprised of a morphologically diverse array of species found across a range of habitats, but is largely restricted to benthic habitats less than 50m. The ‘deep benthic’ is a comprised a group of species specialised for deep water benthic habitats, typically more than 50m in depth. The ‘utaka’ are a group of typically shallow open water shoaling species, primarily represented by members of the genus *Copadichromis*^42^. The ‘mbuna’ are a distinct clade of relatively small-bodied rock-associated species typically found to depths of ∼30m, and with a diverse mix of foraging strategies^43^. The final grouping is represented by only one species, *Astatotilapia calliptera*, that is often present in shallow macrophyte-rich marginal habitats of the Lake Malawi. It is often referred to as a generalist species — consuming phytoplankton, zooplankton and littoral arthropods. It the only representative of the haplochromine radiation that is widespread in rivers and smaller lakes of the Lake Malawi catchment^44,45^. Given the number of species sampled in our dataset, the inclusion of several undescribed species as well as limited species-specific data for some of the species in our dataset, we are unable to offer descriptions of every specimen within our dataset. We have instead limited our wider discussion of species within some of the ecomorphological groups to a selection of well-studied species that demonstrate the diversity captured within our dataset.

### *Diplotaxodon* and *Rhamphochromis*

Nine species of the *Diplotaxodon* group (47%; Table 1), including the type species *Diplotaxodon argenteus* (n=1) and *Pallidochromis tokolosh* (n=2) are present within our dataset. We also sampled 10 species of the *Rhamphochromis* group (71%; Table 1), including the type species *Rhamphochromis longiceps* (n=2) and the remarkably large *Rhamphochromis woodi* (n=2, see below), that are endemic to Lake Malawi. We were also able to sample two sympatric species from the crater lake, Lake Kingiri, *Rhamphochromis* sp. “Kingiri Dwarf” (n=2) and *Rhamphochromis* sp. “Kingiri Large” (n=2), as well as *Rhamphochromis* sp. ‘Chilingali’ (n=4) from the satellite, Lake Chilingali (Figure 4), which is now presumed extinct in the wild^46^.

**Figure 4.**
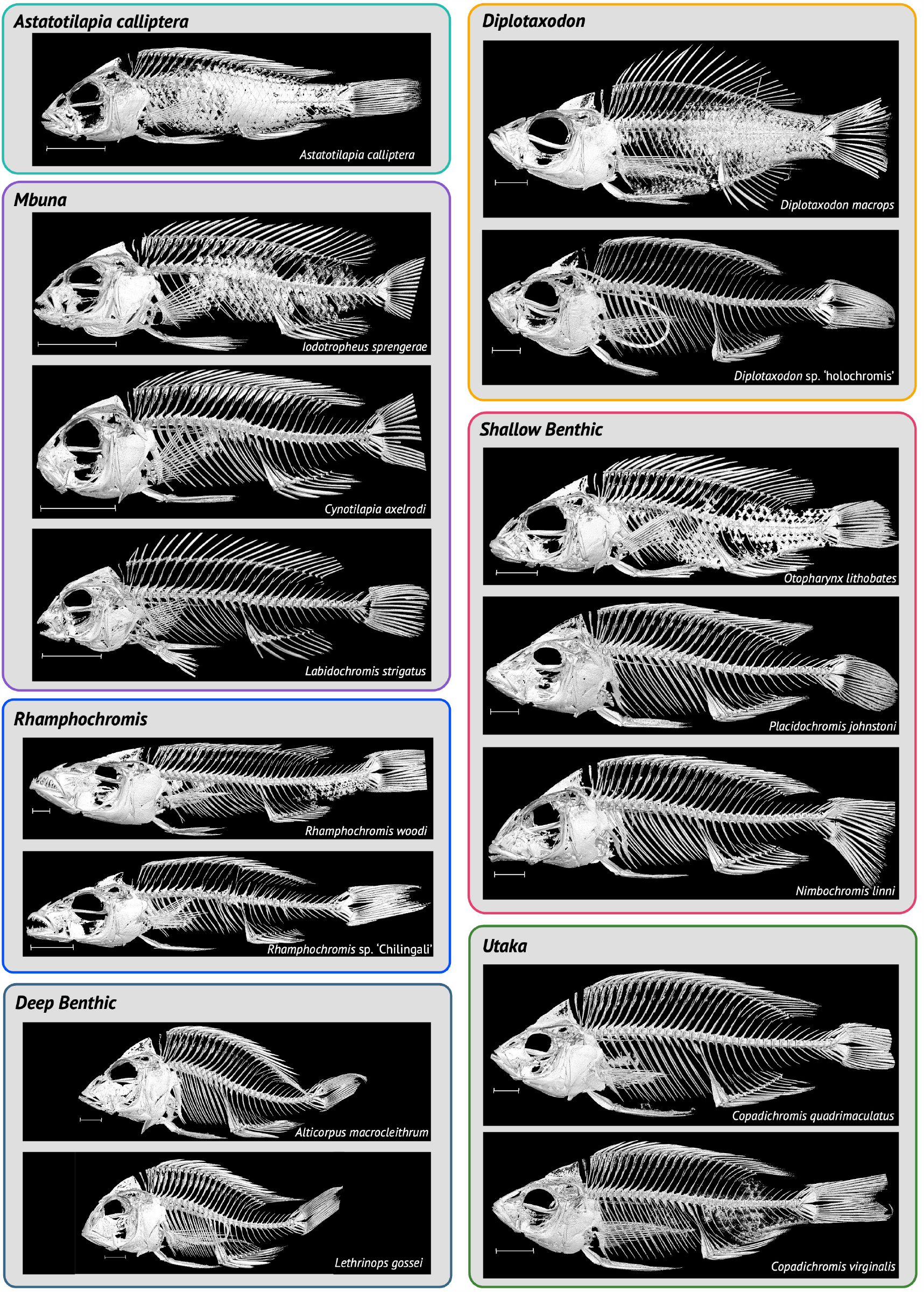
Whole-body 3D models of select specimens from the dataset. Scale is 1cm. Specimens are arranged according to the ecomorphological group they belong to. Species names are indicated. Specimens used are indicated in Supplementary Table 1. Some of the specimens have been rendered with their scales, mainly as we found that several specimens had particularly thick scales making it difficult render whole-body models without including the scales. It is worth noting, however, that the lateral lines have resolved quite well in the models with scales present and with a sufficiently high threshold value scales could resolve well. The ring structure in *Diplotaxodon* sp. ‘holochromis’ is a rubber band used for identification purposes.

The *Diplotaxodon* and *Rhamphochromis* groups are two reciprocally monophyletic diverging lineages of Lake Malawi cichlids^4^ that have adapted to the pelagic zone of Lake Malawi^40^. The majority of species in the groups are piscivorous, although several species, including *Diplotaxodon limnothrissa* (n=2) are predominantly zooplanktivorous^47^. Large-bodied *Rhamphochromis* primarily feed on Lake Malawi sardines (usipa; *Engraulicypris sardella*) and endemic cichlids (e.g. utaka). Members of *Diplotaxodon* and *Rhamphochromis* are among the deepest-living of all Lake Malawi cichlids, with representatives of both being caught at depths exceeding 200 metres – the ‘twilight zone’ where light is almost completely absent^40^. Species within the *Diplotaxodon macrops* complex, which is represented in the scanned samples by *Diplotaxodon macrops* (n=1) (Figure 4), *Diplotaxodon* sp. ‘macrops north’ (n=2), *Diplotaxodon* sp. ‘macrops black dorsal’ (n=2) and *Diplotaxodon* sp. ‘macrops ngulube’ (n=2), have been found between 100 and 220m, a depth similarly reported to be occupied by *Rhamphochromis* during the day^48^.

Morphological comparisons of *Diplotaxodon* and *Rhamphochromis* with Lake Malawi cichlids from other habitats could provide valuable insights into convergent adaptation of traits enabling occupation of pelagic niches. Divergence along depth gradients is associated with the evolution of reproductive isolation in many marine and freshwater species groups, likely a consequence of the strong selective pressures associated with deeper water, such as the absence of sunlight, greater hydrostatic pressure, and reduced levels of dissolved oxygen^49,50^. Morphological comparisons of *Diplotaxodon* and *Rhamphochromis*, against closely related littoral species, could be a powerful model for the study of evolution of convergent phenotypes necessary for adapting to pelagic environments. Body elongation, supported by increased vertebral counts, is a common adaptive trait in teleosts adapted to pelagic (and piscivorous) niches^51^ and is also the case with *Rhamphochromis*^52^. Vertebral morphology has also been related to swimming kinematics, body shape and habitat preference^53^ and the evolutionary modification of internal vertebral traits appears to have taken place during adaptation to pelagic environments^34^. A study of these traits using the Lake Malawi may help to determine the rate at which these phenotypes can become fixed, and provide insight into the role of these morphological adaptations along the benthic-pelagic speciation axis^49,50^.

Remarkable size variation is present within the *Rhamphochromis* genus. *Rhamphochromis woodi* is considered to be one of the largest Lake Malawi cichlids, measuring up to 40 cm standard length^39^. In contrast, wild-caught specimens of *Rhamphochromis sp. ‘Kingiri Dwarf’*, endemic to the crater lake Kingiri, do not exceed 7.5 cm standard length^46^. Similarly, wild caught *Rhamphochromis* sp. ‘Chilingali’ are also small bodied, with maximum observed standard length of 10.6 cm^8^. Notably *Rhamphochromis* sp. ‘Chilingali’ is relatively amenable to laboratory study, and its elongated body has made it a useful model in evolutionary developmental biology^17,18^. Given the exceptional size difference in the genus *Rhamphochromis*, our dataset represents a potentially valuable resource for the study of the evolution of allometric scaling, which has not been well studied in cichlids^54^.

### Shallow Benthic

The shallow benthic species group is extremely speciose, with remarkable amount of morphological diversity^4,41^, and hundreds of species. The majority of shallow benthic species inhabit relatively shallow inshore habitats of Lake Malawi, such as the sand or mud lake floor, or sand-rock transitional zones. Our dataset includes 20 shallow benthic species in 12 genera^4^ (Table 1), including several large, ambush predators, as well as a collection of trophic specialists. For a complete list of shallow benthics see Supplementary Table 1.

Large ambush predators represented in the dataset include *Dimidochromis strigatus* (n=1), *Dimidochromis compressiceps* (n=1), *Tyrannochromis macrostoma* (n=1), *Nimbochromis livingstonii* (n=1) and *Nimbochromis polystigma* (n=2). *Dimi-dochromis compressiceps* has a generalist piscivore lifestyle, and occupies reed-beds of the Lake. *Nimbochromis livingstonii* and *N. polystigma* are both considered to be ‘sleeper’ (Chichewa: “kaligono”), and have been observed burying themselves within sandy substrate and snatching prey attracted by the disturbed sediment^41^. Another member of *Nimbochromis, Nimbochromis linni* (n=1) has a characteristic downward-projecting snout (Figure 4), enabling it to extract prey from rock crevices^41,55^.

We sampled several shallow-benthic predators, including *Otopharynx speciosus* (n=2), one of the few piscivores within the genus *Otopharynx*. Males of this species have been encountered at depths exceeding 25m^41^, suggesting tolerance of relatively deep water, and suggesting the species may have morphological adaptations enabling occupation of deep-water niches similar to *Rhamphochromis* and *Diplotaxodon*. Of the approximately 20 species of *Otopharynx*^56^ we were able to sample an additional three species: *Otopharynx lithobates* (n=3, including the holotype NHMUK 1974.7.5.1); *Otopharynx tetrastigma* (n=2); and the undescribed *Otopharynx* sp. “brooksi nkhata” (n=1). We also sampled several specialised trophic specialists including the molluscivores *Mylochromis anaphyrmus* (n=1) and *Trematocranus placodon* (n=1) and the invertebrate picker *Placidochromis johnstoni* (n=1, Figure 4). The diet of *T. placodon* predominately comprises the gastropods *Bulinus nyassanus* and *Melanoides tuberculata*^57^. Enlarged sensory pores and lateral lines form a sonar-like detection system that allows *T. placodon* to sense the movement of these prey within the sediment. Curiously, this strategy and associated morphological characteristics are also associated with *Aulonocara* and *Lethrinops*, both ‘deep benthics’, suggesting convergent evolution of lateral line phenotypes^9^. The specimens in our dataset may enable some morphological comparisons to further investigate differences in the sensory pore characteristics among species.

### ‘Rock-dwelling’ *Mbuna*

The mbuna group dominate the rocky shores of Lake Malawi, and are used as a model system for the study of rapid speciation and adaptive radiation^25,43,58^. Similar to the shallow-benthics, there are hundreds of species, many of which are undescribed^41,43^. We aimed to maximise our coverage of the phenotypic diversity in the group by sampling multiple genera, which are largely differentiated on the basis of head, jaw and tooth morphology^43^. Our dataset includes 7 species (15 individuals) of mbuna, covering 7 of the 14 described mbuna genera (Table 1).

*Cynotilapia* can be distinguished from other genera by the presence of unicuspid (conical) teeth^41,59,60^ and is represented in our dataset by *Cynotilapia axelrodi* (n=1, Figure 4). This is a genus of typically planktivorous species^41^ and their relatively simple dentition may reflect this lifestyle^61^. By contrast, *Maylandia* (*Metriaclima*^62^), represented by *Maylandia zebra* (n=5), has closely arranged bicuspid teeth, that is uses for pulling and scraping loose Aufwuchs (periphyton) attached to the rocks found in their preferred preferred rocky habitats^59,63^. *Tropheops* and *Iodotropheus*, represented by *Tropheops tropheops* (n=2) and *Iodotropheus sprengerae* (n=2), also have closely packed bicuspid teeth, that they use to feed on epilithic algae which they pluck with sideways, upwards head jerks, a behaviour likely supported by *Tropheops*’ characteristic steeply sloped vomer (71-96°)^41^. Members of *Petrotilapia*, represented by *Petrotilapia genalutea* (n=1) have a mixed combination of tricupsid and unicupsid teeth that they use to comb loose peripyton from rock surfaces^64^. A further represented mbuna genus is the monotypic *Genyochromis*, represented by *Genyochromis mento* (n=2). Like the majority of mbuna, *G. mento* has prominent outer bicupsid teeth, and are supported by smaller inner tricupsid teeth^41,59^. In contrast to most other mbuna, however, *G. mento* is a highly specialised feeder, a lepidophage (scale-eater), that targets the the caudal and anal fins of other cichlids in rocky habitats^41,59,65^. The preferred striking side of *G. mento* significantly correlates with left-right asymmetry of the dentary, with right and left-leaning individuals preferring to strike the corresponding side, respectively, of their prey. Interestingly, however, a comparison of their jaw laterality with *Perissodus microlepis*, a lepidophage endemic to Lake Tanganyika^20^, showed that laterality in *G. mento* is weaker than in *P. microlepis* – likely a result of phylogenetic constraint from their shorter evolutionary history and their herbivorous ancestors^65^.

The craniofacial bones commonly studied in mbuna, such as the dentary, premaxilla, pharyngeal jaws, as well as their associated teeth, can be segmented from specimens in the dataset (see *G. mento*, Figure 5). Future sampling should focus on the seven remaining genera not sampled in our dataset: *Abactochromis, Chindongo, Cyathochromis, Gepyrochromis, Labeotropheus, Melanochromis* and *Psuedotropheus*.

**Figure 5.**
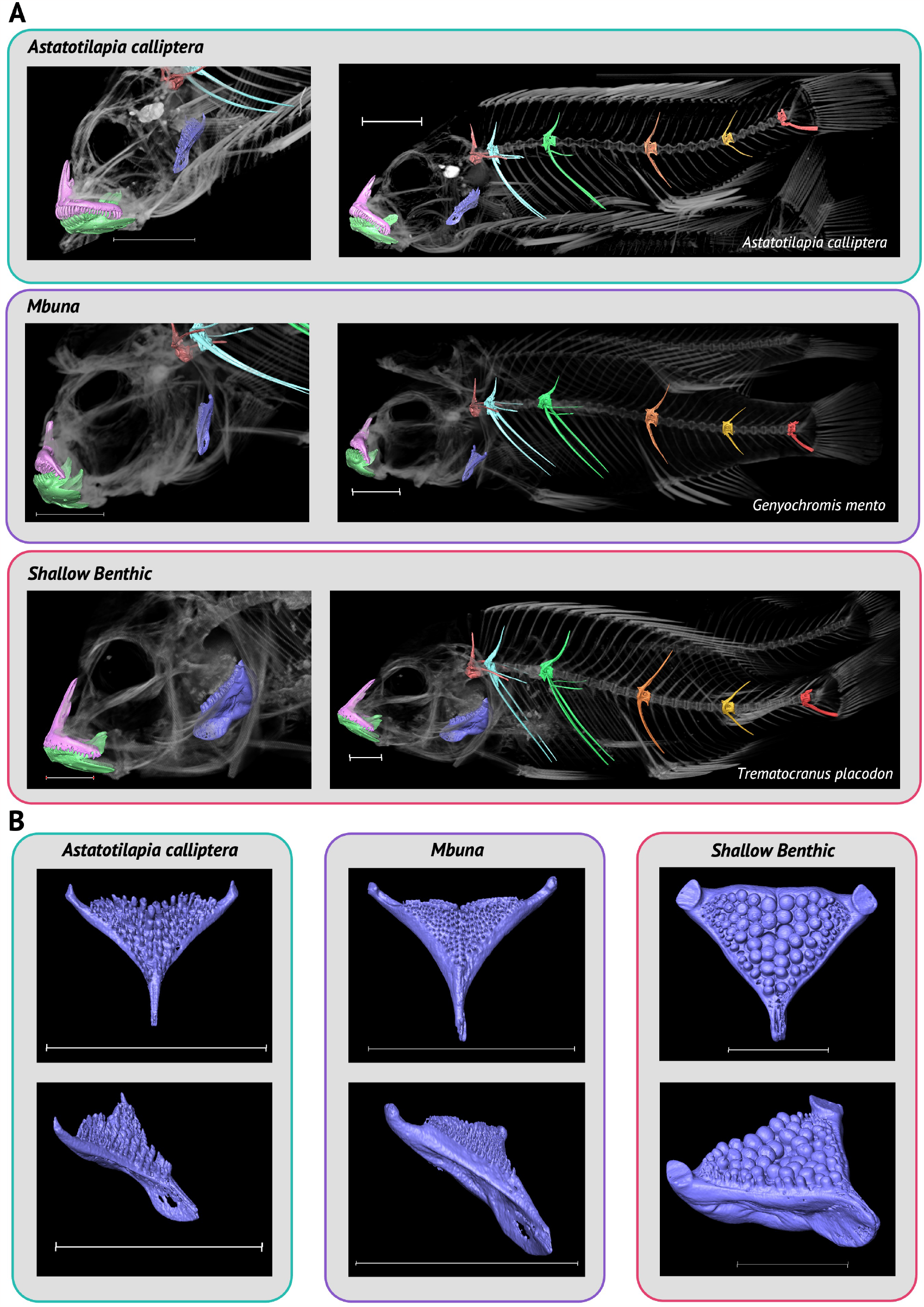
Segmented Bones from *Astatotilapia calliptera, Genyochromis mento* (mbuna) and *Trematocranus placodon* (shallow benthic). (A, left) A close up, lateral view of the head of each species (species name indicated on right), showing the dentary (green), premaxilla (pink) and lower pharyngeal jaw (purple) positioned within the a volume render of the head. (A, right) A whole body lateral view showing the aforementioned jaw bones, as well as the first non-rib-bearing vertebra (orange), the first rib-bearing (precaudal) vertebra (light blue), precaudal vertebra 8 (green), caudal vertebra 3 (orange), caudal vertebra 10 (gold) and the pre-urostyle (final caudal) vertebra (red). (B) Anterior (top) and anterolateral (bottom) view of the lower pharyngeal jaws for each species in (A). Scale for all images is shown as 1cm. See Supplementary Table 1 for details of the specimens used.

#### *Astatotilapia calliptera* and Ruaha Catchment

*Astatotilapia* is polyphyletic and current members of the genus are widespread across East and North Africa^6,66,67^. Only one species of *Astatotilapia* is native to Lake Malawi, *Astatotilapia calliptera*, which is also found in East African rivers flowing eastward to the Indian Ocean, from the Rovuma River in the north, to the Save River in the south. Given the wide distribution of the species, it is perhaps unsurprising that intraspecific genetic variation within the species is comparable to that of the whole Lake Malawi radiation^4,6^. Despite their wide distribution and relatively large intraspecific genetic variation, they phylogenetically cluster within the Lake Malawi radiation (Figure 1), forming a sister clade to the mbuna, with which they share an excess of alleles^4^. This pattern, alongside a perceived riverine ‘generalist’ lifestyle, has led to the hypothesis that either Lake Malawi cichlids radiated from an *A. calliptera*-*like* ancestor or that *A. calliptera* is the sympatric ancestor of all Lake Malawi cichlids^4,6,41,67^.

Given the importance of *A. calliptera* in the Lake Malawi radiation, we sampled multiple individuals from multiple populations. We were able to scan nine laboratory-reared individuals from the Mbaka river population, which flows into the northern end of Lake Malawi^68^. We also scanned individuals from Lake Chilwa (an endorheic lake south-east of Lake Malawi^69^; n=2), Lake ‘Misoko’, presumably Lake Masoko (a crater lake north of Lake Malawi^68^, n=2), and wild-caught individuals from the main body of Lake Malawi (n=2). Populations of *A. calliptera* differ in life history strategies^70^ and are also undergoing sympatric speciation along a depth gradient in at least one location (Lake Masoko)^8^, where littoral and benthic *A. calliptera* ecomorphs have diverged in multiple characteristics, including body shape and trophic specialism, in approximately 1000 years^8^. Therefore, it is possible that morphological evaluations of more populations of *A. calliptera* will reveal further diversity, potentially providing greater insight into the role it has taken in generating the wider Lake Malawi haplochromine radiation.

A key part of macroevolutionary studies is the estimation of ancestral state of traits based on the morphology of their descendants^71^. This relies on a comprehensive understanding of trait diversity across taxa, and such data can also inform models of morphological evolution, and enable estimates of rates of phenotypic evolution. Since the genetic diversity of the Lake Malawi radiation was possibly seeded by multiple riverine species^5^, we sought to add specimens to the dataset that could enable the morphological reconstruction of the common ancestor of the Lake Malawi radiation. Therefore, we also sampled two additional species of *Astatotilapia*. These included *Astatotilapia gigliolli* (n=2) and *Astatotilapia* sp. ‘Ruaha blue’ (n=2), native to the Great Ruaha River^66,67^. Construction of a mtDNA-based phylogeny initially placed *Astatotilapia* sp. ‘Ruaha blue’ as a sister taxa to the Lake Malawi radiation^66^. However, a phylogeny based on variation within whole-genome sequences has shown *A. gigliolli* and *A*. sp. ‘Ruaha blue’, sister taxa, form a sister clade with both the Lake Malawi and Lake Victoria radiations (see Figure 1) – a topology that is likely the result of an ancestral hybridisation event with the ancestors of both lineages prior to their respective adaptive radiations^5^. Therefore, the addition of species from the Ruaha catchment, may therefore enable a more robust estimation of the ancestral phenotype of Lake Malawi cichlids.

### Deep Benthic and ‘*Utaka*’

We sampled deep-water benthic species from two genera; *Alticorpus* and *Lethrinops*^9,45^ (Table 1). *Alticorpus*, like *Aulonocara* (not represented in the dataset) is characterised by the presence of greatly enlarged cranial sensory openings and lateral line canals used to detect prey in the sediment. Deep-water benthic species are found below 50m, a ‘twilight’ zone with very little visible light. *Alticorpus macrocleithrum* (n=3) is found between 75m and 125m, with abundance peaking above 100m^72^, a depth similarly occupied by deep-water Lethrinops^73^, including *Lethrinops gossei* (n=1). Several species of *Lethrinops*, however, inhabit shallower water^4,41^. We sampled two spcies of shallow water *Lethrinops*, including *Lethrinops auritus* (n=2), and *Lethrinops albus* (n=2), both of which phylogenetically cluster within the ‘shallow benthic’ lineage (Figure 1).

Our dataset also contains four species of zooplankton-feeding, shoaling cichlids which are commonly referred to as ‘utaka’. Utaka is primarily made up of species belonging to *Copadichromis*^42^, with a small number of species also belonging to *Mchenga* and the older *Nyassachromis*^74^. However, their placement within the utaka is disputed and they have not been considered in our species/genera counts (see Table 1). We sampled four species of *Copadichromis*: *Copadichromis likomae* (n=2), *Copadichromis quadrimaculatus* (n=2), *Copadichromis trimaculatus* (n=2, see Figure 6) and *Copadichromis virginalis* (n=2). Utaka feed in the water column, and can be commonly found close to the shore^41^. *Copadichromis* are generally characterised by their relatively small, highly protrusible mouths, that they use to suck zooplankton into their mouths, as well as numerous long gill rakers which strain plankton from the water that enters their mouths as a result of their sucking feeding mechanism^41,75^.

**Figure 6.**
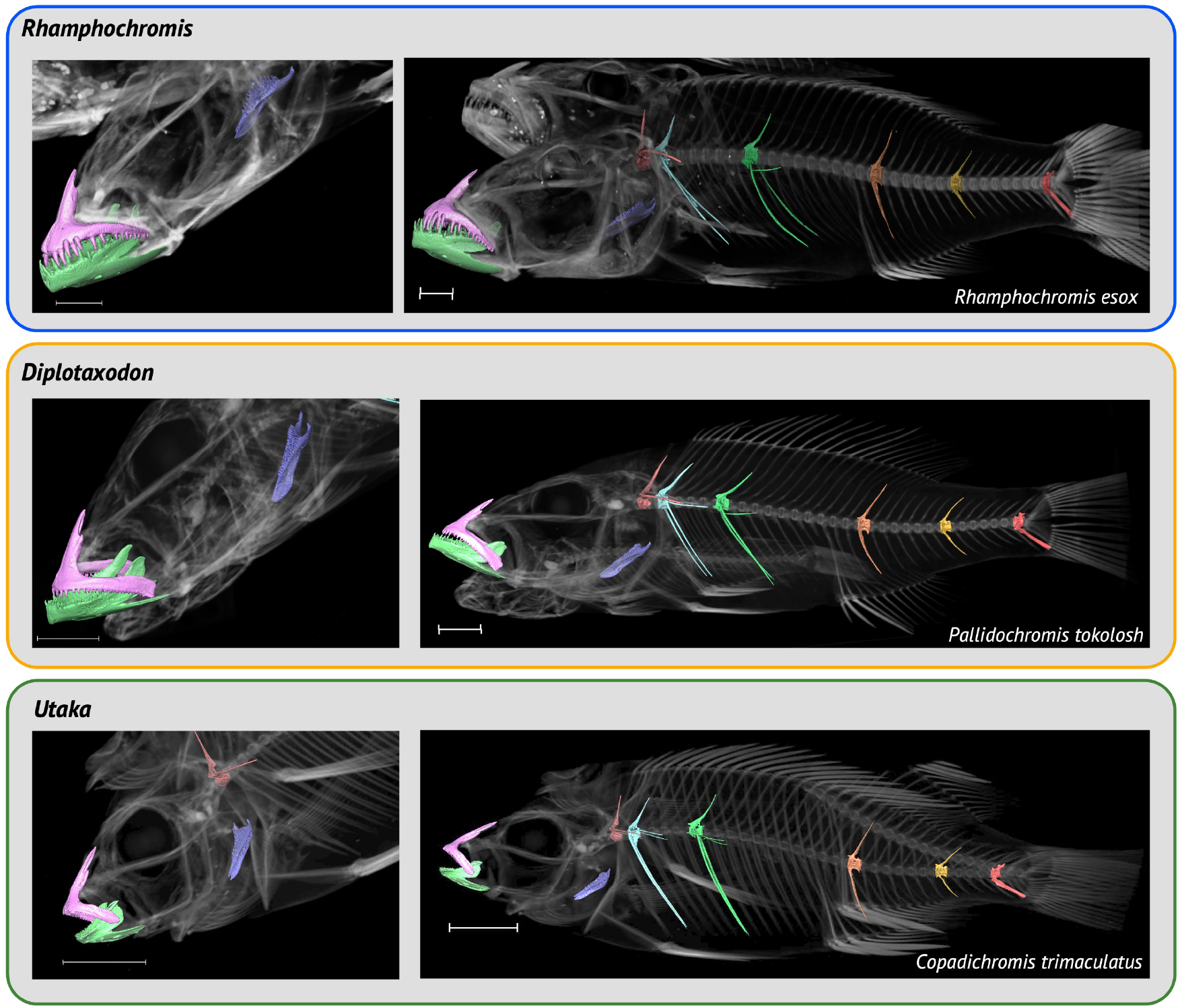
Segmented Bones from *Rhamphochromis esox* (*Rhamphochromis*), *Pallidochromis tokolosh* (*Diplotaxodon*) and *Copadichromis trimaculatus* (Utaka). A close up, lateral view of the head of each species (species name indicated on right), showing the dentary (green), premaxilla (pink) and lower pharyngeal jaw (purple) positioned within a volume render of the head is shown for each specimen on the left. On the right are whole body lateral views showing the aforementioned jaw bones, as well as the first non-rib-bearing vertebrae (orange), the first rib-bearing (precaudal) vertebrae (light blue), precaudal vertebrae 8 (green), caudal vertebrae 3 (orange), caudal vertebrae 10 (gold) and the pre-urostyle (final caudal) vertebrae (red) within a volume rendering of the whole body. Scale for all images is shown as 1cm. See Supplementary Table 1 for details of the specimens used.

Both the deep benthics and utaka are currently underrepresented within our dataset and future sampling should aim to add additional species of *Copadichromis*. In addition, sampling missing ‘deep benthic’ genera, such as *Aulonocara* and *Tramitichromis* should be prioritised for future sampling efforts. *Aulonocara stuartgranti* and *Aulonocara steveni* would be particularly interesting future additions and could offer interesting morphological comparisons with deeper living species. Moreover, given that *Lethrinops* is polyphyletic^73,76^, additional sampling of *Lethrinops* species could provide morphological data to support future systematic studies.

## Technical Validation and Usage Notes

As mentioned above, entries for each specimen can be found in Supplementary Table 1 and cropped, reconstructed 8-bit tiff images stacks of all specimens scanned can be accessed and downloaded from a dedicated project on Morphosource.

To validate our *μ*CT-scan data, we imported the reconstructed image stacks into Avizo Lite (v9.3.0), a proprietary software developed by Thermo Fisher Scientific and either generated a full body volume render or rendered 3D models of the whole body. Surfaces from 3D whole-body renderings were exported from AvizoLite as Polygon File Format (.ply) files and imported into MeshLab for visualisation and manipulation. We note that there are free, open access alternatives to Avizo Lite (v9.3.0) for the segmentation of 3D-image data, such as 3D-Slicer, which has a large and active community of users^77^ and Dragonfly which supports the use of deep learning to automatically segment 3D image data and offers non-commercial licenses for academic use free-of-charge. Example whole-body 3D renderings for representatives from each ecomorphological group can be found in Figure 4 and for each specimen in the dataset in the Supplementary Material.

In addition, we manually segmented multiple bones, including the dentary, premaxilla, lower pharyngeal jaw and multiple vertebral types in several species discussed above. This included: the ‘generalist’ *Astatotilapia calliptera*; the ‘mbuna’ and lepidophage, *Genyochromis mento*; the ‘shallow benthic’ and snail crusher, *Trematocranus placodon* (Figure 5); the pelagic piscivores, *Rhamphochromis esox* and *Pallidochromis tokolosh* and the zooplanktivorous (utaka), *Copadichromis trimaculatus* (Figure 6). Lower pharyngeal jaws that are highly variable among species of Lake Malawi cichlids^25^ were particularly well resolved. For example, newly erupting teeth were visible on the relatively large, and dense, lower pharyngeal jaw of *Trematocranus placodon* (Figure 5B). Similarly, renderings of multiple vertebral types (Figures 5-6) were also good quality. The zygapophyses and fine structure of the vertebral centra, sometimes including the neural canal, were also well resolved. All 3D-renderings of these bones can be found in the Supplementary Materials as downloadable .ply files.

The computer specifications we used for the all analyses (see Methodology) are hard to find on personal, or older machines and some users may find it difficult to work with some of our larger image stacks. To minimise memory usage during segmentation and speed up processing, cropped reconstructed stacks can be loaded in multiple increments (note that the Z-voxel size must be multiplied by said increment). We tested this and found that roughly comparable models could be generated, although it was clear that finer morphological detail was absent (data not shown). Therefore, where possible, the whole stack should be used when segmenting regions of interest. In addition, since these regions were manually segmented, many of the segmentation steps rely on the judgment of the individual segmenting and rendering of the regions of interest. We found that segmenting from median-filtered reconstructed image stacks drastically lowered the quality of the rendered models (data not shown) and would suggest refraining from segmenting from a median-filtered image stack. In addition, we found that relatively low smoothing factors were best for rendering surfaces from segmented regions of interest. In Avizo Lite (v9.3.0), a smoothing factor between 0-10 (including rational intermediates) can be applied when rendering surfaces of segmented regions of interest. We rarely found it necessary to use a value above ‘3’; indeed, all whole-body 3D models were smoothed with a factor of 2.5, and we suggest, regardless of the tool used to smooth and render segmented surfaces, to use smoothing cautiously. Furthermore, we were able to scan multiple species of *Copadichromis* (utaka), see above and were able to generate relatively good quality models (see Figure 6 and the Supplementary Models). However, it is clear that some of the jaw structures did not resolve as well as in other specimens. It is possible that the jaw bones of these fish species are not particularly dense, which made it difficult to image them using the same scanning procedure as that was used for all of the other specimens. Therefore, in future, we would recommend care when segmenting bones from the *Copadichromis* species present in this dataset and also to increase the power and exposure time for future sampling of *Copadichromis*.

Here we have presented the first, comprehensive, freely available, whole-body *μ*CT-scan dataset of Lake Malawi cichlids. We have described several macroevolutionary studies that could be conducted with this dataset to better understand this remarkable cichlid adaptive radiation and include suggestions for future sampling. Our Lake Malawi dataset now joins two other East African adaptive radiation datasets, the recent haplochromine Lake Victoria library^78^ and the extensive *μ*CT-scan dataset of Lake Tanganyika cichlid fishes^7^. Therefore, the addition of our dataset now offers a unique opportunity for much wider, and systematic, morphological comparisons within East African cichlids. We have also described a methodology to efficiently *μ*CT-scan multiple specimens simultaneously; reducing scanning time and financial cost, whilst maintaining scan quality, demonstrating the utility of this method by reconstructing 3D-models of multiple bones from multiple specimens within our dataset. We hope the availability of these data will inspire people to address some of the many questions left still to understand this remarkable adaptive radiation, permit wider-scale comparisons with other cichlid adaptive radiations and set a precedent to make whole-body *μ*CT scans the automatic default for any sampling efforts involving cichlids.

## Supporting information

Whole Body Renderings

3D Bone Models

## Acknowledgements

We thank Vincent Fernandez, CT facility manager at the NHMUK and Liz Martin-Silverstone at the XTM Facility at the University of Bristol for organising access to their respective imaging facilities. Thank you to the fish whose lives were sacrificed for this work. This research was partly funded by a Biotechnology and Biological Sciences Research Council (BBSRC) studentship (Grant Number: 2445747).

## Author contributions statement

C.V.B. and B.V. conceived the study. C.V.B and R.B. designed the data acquisition pipeline and dataset curation methodology. C.V.B. and R.B. conducted the imaging. C.V.B. performed the sample collection, processed all the raw image data, and wrote the manuscript. B.V., R.B. and M.J.G. edited the manuscript. J.M. organised specimens at the NHMUK. G.F.T. and M.J.G. contributed specimens to be imaged. All authors reviewed the manuscript.

## References

1. Turner, G. F., Seehausen, O., Knight, M. E., Allender, C. J. & Robinson, R. L. How many species of cichlid fishes are there in african lakes? Mol. Ecol. 10, 793–806, 10.1046/j.1365-294x.2001.01200.x (2001).

2. Barlow, G. The Cichlid Fishes: Nature’s Grand Experiment In Evolution (Hachette UK, London, 2008).

3. Sparks, J. S. & Smith, W. L. Phylogeny and biogeography of cichlid fishes (teleostei: Perciformes: Cichlidae). Cladistics 20, 501–517, 10.1111/j.1096-0031.2004.00038.x (2004).

4. Malinsky, M. et al. Whole-genome sequences of malawi cichlids reveal multiple radiations interconnected by gene flow. Nat. Ecol. Evol. 2, 1940–1955, 10.1038/s41559-018-0717-x (2018).

5. Svardal, H. et al. Ancestral hybridization facilitated species diversification in the lake malawi cichlid fish adaptive radiation. Mol. biology evolution 37, 1100–1113, 10.1093/molbev/msz294 (2020).

6. Svardal, H., Salzburger, W. & Malinsky, M. Genetic variation and hybridization in evolutionary radiations of cichlid fishes. Annu. Rev. Animal Biosci. 9, 55–79, 10.1146/annurev-animal-061220-023129 (2021).

7. Ronco, F. et al. Drivers and dynamics of a massive adaptive radiation in cichlid fishes. Nature 589, 76–81, 10.1038/s41586-020-2930-4 (2021).

8. Genner, M. J. et al. Evolution of a cichlid fish in a lake malawi satellite lake. Proc. Royal Soc. B: Biol. Sci. 274, 2249–2257, 10.1098/rspb.2007.0619 (2007).

9. Genner, M. J. & Turner, G. F. Ancient hybridization and phenotypic novelty within lake malawi’s cichlid fish radiation. Mol. Biol. Evol. 29, 195–206, 10.1093/molbev/msr183 (2012).

10. Meier, J. I. et al. Ancient hybridization fuels rapid cichlid fish adaptive radiations. Nat. communications 8, 14363, 10.1038/ncomms14363 (2017).

11. Keller-Costa, T., Canário, A. V. & Hubbard, P. C. Chemical communication in cichlids: a mini-review. Gen. comparative endocrinology 221, 64–74, 10.1016/j.ygcen.2015.01.001 (2015).

12. Faber-Hammond, J. J. & Renn, S. C. Transcriptomic changes associated with maternal care in the brain of mouthbrooding cichlid astatotilapia burtoni reflect adaptation to self-induced metabolic stress. J. Exp. Biol. 226, jeb244734, 10.1242/jeb.244734 (2023).

13. Plenderleith, M., Oosterhout, C. v., Robinson, R. L. & Turner, G. F. Female preference for conspecific males based on olfactory cues in a lake malawi cichlid fish. Biol. Lett. 1, 411–414, 10.1098/rsbl.2005.0355 (2005).

14. Morita, M. et al. Bower-building behaviour is associated with increased sperm longevity in tanganyikan cichlids. J. evolutionary biology 27, 2629–2643, 10.1111/jeb.12522 (2014).

15. McKaye, K. R. & Kocher, T. Head ramming behaviour by three paedophagous cichlids in lake malawi, africa. Animal Behav. 31, 206–210, 10.1016/S0003-3472(83)80190-0 (1983).

16. Woltering, J. M., Holzem, M., Schneider, R. F., Nanos, V. & Meyer, A. The skeletal ontogeny of astatotilapia burtoni–a direct-developing model system for the evolution and development of the teleost body plan. BMC developmental biology 18, 1–23, 10.1186/s12861-018-0166-4 (2018).

17. Santos, M. E., Lopes, J. F. & Kratochwil, C. F. East african cichlid fishes. EvoDevo 14, 1, 10.1186/s13227-022-00205-5 (2023).

18. Marconi, A., Yang, C. Z., McKay, S. & Santos, M. E. Morphological and temporal variation in early embryogenesis contributes to species divergence in malawi cichlid fishes. Evol. & Dev. 25, 170–193, 10.1111/ede.12429 (2023).

19. Navon, D., Olearczyk, N. & Albertson, R. C. Genetic and developmental basis for fin shape variation in african cichlid fishes. Mol. Ecol. 26, 291–303, 10.1111/mec.13905 (2017).

20. Ronco, F., Büscher, H. H., Indermaur, A. & Salzburger, W. The taxonomic diversity of the cichlid fish fauna of ancient lake tanganyika, east africa. J. Gt. Lakes Res. 46, 1067–1078, 10.1016/j.jglr.2019.05.009 (2020).

21. Arthur, W. The emerging conceptual framework of evolutionary developmental biology. Nature 415, 757–764, 10.1038/415757a (2002).

22. Albertson, R. C. & Kocher, T. D. Assessing morphological differences in an adaptive trait: a landmark-based morphometric approach. The J. Exp. Zool. 289, 385–403, 10.1002/jez.1020 (2001).

23. Adams, D., Yamaoka, K. & Kassam, D. Functional significance of variation in trophic morphology within feeding microhabitat-differentiated cichlid species in lake malawi. Animal Biol. 54, 77–90, 10.1163/157075604323010060 (2004).

24. Hulsey, C. D., Alfaro, M. E., Zheng, J., Meyer, A. & Holzman, R. Pleiotropic jaw morphology links the evolution of mechanical modularity and functional feeding convergence in lake malawi cichlids. Proc. Royal Soc. B: Biol. Sci. 286, 20182358, 10.1098/rspb.2018.2358 (2019).

25. Conith, A. J. & Albertson, R. C. The cichlid oral and pharyngeal jaws are evolutionarily and genetically coupled. Nat. Commun. 12, 5477, 10.1038/s41467-021-25755-5 (2021).

26. Kratochwil, C. F. et al. Agouti-related peptide 2 facilitates convergent evolution of stripe patterns across cichlid fish radiations. Science 362, 457–460, 10.1126/science.aao6809 (2018).

27. Gerwin, J., Urban, S., Meyer, A. & Kratochwil, C. F. Of bars and stripes: A malawi cichlid hybrid cross provides insights into genetic modularity and evolution of modifier loci underlying colour pattern diversification. Mol. Ecol. 30, 4789–4803, 10.1111/mec.16097 (2021).

28. Clark, B. et al. Oca2 targeting using crispr/cas9 in the malawi cichlid astatotilapia calliptera. Royal Soc. Open Sci. 9, 220077, 10.1098/rsos.220077 (2022).

29. DeLorenzo, L. et al. Genetic basis of ecologically relevant body shape variation among four genera of cichlid fishes. Mol. ecology 32, 3975–3988, 10.1111/mec.16977 (2023).

30. Darrin Hulsey, C., Keck, B. P., Alamillo, H. & O’Meara, B. C. Mitochondrial genome primers for lake malawi cichlids. Mol. Ecol. Resour. 13, 347–353, 10.1111/1755-0998.12066 (2013).

31. McGee, M. D. et al. The ecological and genomic basis of explosive adaptive radiation. Nature 586, 75–79, 10.1038/s41586-020-2652-7 (2020).

32. Masonick, P., Meyer, A. & Hulsey, C. D. Phylogenomic analyses show repeated evolution of hypertrophied lips among lake malawi cichlid fishes. Genome Biol. Evol. 14, evac051, 10.1093/gbe/evac051 (2022).

33. Price, S. A., Friedman, S. T. & Wainwright, P. C. How predation shaped fish: the impact of fin spines on body form evolution across teleosts. Proc. Royal Soc. B: Biol. Sci. 282, 20151428, 10.1098/rspb.2015.1428 (2015).

34. Baxter, D., Cohen, K. E., Donatelli, C. M. & Tytell, E. D. Internal vertebral morphology of bony fishes matches the mechanical demands of different environments. Ecol. Evol. 12, e9499, 10.1002/ece3.9499 (2022).

35. Todd Streelman, J. & Danley, P. D. The stages of vertebrate evolutionary radiation. Trends Ecol. & Evol. 18, 126–131, 10.1016/S0169-5347(02)00036-8 (2003).

36. Gavrilets, S. & Losos, J. B. Adaptive radiation: Contrasting theory with data. Science 323, 732–737, 10.1126/science.1157966 (2009).

37. Schindelin, J. et al. Fiji: an open-source platform for biological-image analysis. Nat. methods 9, 676–682, 10.1038/nmeth.2019 (2012).

38. Schneider, C. A., Rasband, W. S. & Eliceiri, K. W. Nih image to imagej: 25 years of image analysis. Nat. methods 9, 671–675, 10.1038/nmeth.2089 (2012).

39. Turner, G., Robinson, R., Shaw, P., Carvalho, G. & Snoeks, J. Identification and biology of Diplotaxodon, Rhamphochromis and Pallidochromis (Cichlid Press, 2004).

40. Hahn, C., Genner, M. J., Turner, G. F. & Joyce, D. A. The genomic basis of cichlid fish adaptation within the deepwater “twilight zone” of lake malawi. Evol. Lett. 1, 184–198, 10.1002/evl3.20 (2017).

41. Konings, A. Malaŵi Cichlids in their Natural Habitat (Cichlid Press, 2016), 5th edition edn.

42. Anseeuw, D., Nevado, B., Busselen, P., Snoeks, J. & Verheyen, E. Extensive introgression among ancestral mtDNA lineages: Phylogenetic relationships of the utaka within the lake malawi cichlid flock. Int. J. Evol. Biol. 2012, 1–9, 10.1155/2012/865603 (2012).

43. Genner, M. J. & Turner, G. F. The mbuna cichlids of lake malawi: a model for rapid speciation and adaptive radiation. Fish fisheries 6, 1–34, 10.1111/j.1467-2679.2005.00173.x (2005).

44. Nichols, P. et al. Secondary contact seeds phenotypic novelty in cichlid fishes. Proc. Royal Soc. B: Biol. Sci. 282, 20142272, 10.1098/rspb.2014.2272 (2015).

45. Joyce, D. A. et al. Repeated colonization and hybridization in lake malawi cichlids. Curr. Biol. 21, R108–R109, 10.1016/j.cub.2010.11.029 (2011).

46. Turner, G., Ngatunga, B. P. & Genner, M. J. The natural history of the satellite lakes of lake malawi. Preprint at 10.32942/osf.io/sehdq (2019).

47. Turner, G. F. Description of a commercially important pelagic species of the genus diplotaxodon (pisces: Cichlidae) from lake malawi, africa. J. Fish Biol. 44, 799–807, 10.1111/j.1095-8649.1994.tb01256.x (1994).

48. Lowe-McConnell, R. Recent research in the african great lakes: fisheries, biodiversity and cichlid evolution. Accesible here =https://aquadocs.org/handle/1834/22270 (2003).

49. Wilson, G. D. & Hessler, R. R. Speciation in the deep sea. Annu. Rev. Ecol. Syst. 18, 185–207, 10.1146/annurev.es.18.110187.001153 (1987).

50. Jennings, R. M., Etter, R. J. & Ficarra, L. Population differentiation and species formation in the deep sea: the potential role of environmental gradients and depth. PLoS One 8, e77594, 10.1371/journal.pone.0077594 (2013).

51. Neat, F. & Campbell, N. Proliferation of elongate fishes in the deep sea. J. Fish Biol. 83, 1576–1591, 10.1111/jfb.12266 (2013).

52. Stiassny, M. Phylogenetic versus convergent relationships between piscivorous cichlid fishes from lakes malawi and tanganyika. Bull. Br. Mus. (Natural Hist. Zool. 40, 67 –101 (1981).

53. Donatelli, C. M. et al. Foretelling the flex—vertebral shape predicts behavior and ecology of fishes. Integr. Comp. Biol. 61, 414–426, 10.1093/icb/icab110 (2021).

54. Fujimura, K. & Okada, N. Shaping of the lower jaw bone during growth of nile tilapia oreochromis niloticus and a lake victoria cichlid haplochromis chilotes: A geometric morphometric approach. Dev. Growth & Differ. 50, 653–663, 10.1111/j.1440-169X.2008.01063.x (2008).

55. Gatumu, E. M. Redescription of the genera nimbochromis and tyrannochromis (teleostei: Cichlidae) from lake malawi, africa. https://solo.bodleian.ox.ac.uk/permalink/44OXF_INST/ao2p7t/cdi_proquest_journals_305235338 (2003).

56. Oliver, M. K. Six new species of the cichlid genus otopharynx from lake malaŵi (teleostei: Cichlidae). Bull. Peabody Mus. Nat. Hist. 59, 159–197, 10.3374/014.059.0204 (2018).

57. Evers, B. N., Madsen, H., McKaye, K. M. & Stauffer, J. R. The schistosome intermediate host, bulinus nyassanus, is a ‘preferred’ food for the cichlid fish, trematocranus placodon, at cape maclear, lake malawi. Annals Trop. Medicine & Parasitol. 100, 75–85, 10.1179/136485906X78553 (2006).

58. Albertson, R. C. Morphological divergence predicts habitat partitioning in a lake malawi cichlid species complex. Copeia 2008, 689–698, 10.1643/CG-07-217 (2008).

59. Ribbink, A., Marsh, B., Marsh, A., Ribbink, A. & Sharp, B. A preliminary survey of the cichlid fishes of rocky habitats in lake malawi. South Afr. J. Zool. 18, 149–310, 10.1080/02541858.1983.11447831 (1983).

60. Kassam, D., Seki, S., Rusuwa, B., Ambali, A. J. & Yamaoka, K. Genetic diversity within the genus cynotilapia and its phylogenetic position among lake malawi’s mbuna cichlids. Afr. J. Biotechnol. 4, 10.4314/ajb.v4i10.71319 (2005).

61. Genner, M., Turner, G., Barker, S. & Hawkins, S. Niche segregation among lake malawi cichlid fishes? evidence from stable isotope signatures. Ecol. Lett. 2, 185–190, 10.1046/j.1461-0248.1999.00068.x (1999).

62. Stauffer Jr, J. R., Bowers, N. J., Kellogg, K. A. & McKaye, K. R. A revision of the blue-black pseudotropheus zebra (teleostei: Cichlidae) complex from lake malaŵi, africa, with a description of a new genus and ten new species. Proc. Acad. Nat. Sci. Phila. 189–230, http://www.jstor.org/stable/4065053 (1997).

63. Holzberg, S. A field and laboratory study of the behaviour and ecology of pseudotropheus zebra (boulenger), an endemic cichlid of lake malawi (pisces; cichlidae). J. Zool. Syst. Evol. Res. 16, 171–187, 10.1111/j.1439-0469.1978.tb00929.x (1978).

64. Marsh, A. A taxonomic study of the fish genus petrotilapia (pisces: Cichlidae) from lake malawi. Ichthyol. Bull. J.L.B. Smith Inst. Ichthyol. 48, 1–14 (1983).

65. Takeuchi, Y. et al. Specialized movement and laterality of fin-biting behaviour in genyochromis mento in lake malawi. J. Exp. Biol. 222, jeb191676, 10.1242/jeb.191676 (2019).

66. Genner, M. J., Ngatunga, B. P., Mzighani, S., Smith, A. & Turner, G. F. Geographical ancestry of lake malawi’s cichlid fish diversity. Biol. Lett. 11, 20150232, 10.1098/rsbl.2015.0232 (2015).

67. Turner, G., Ngatunga, B. P. & Genner, M. J. Astatotilapia species (teleostei, cichlidae) from malawi, mozambique and tanzania, excluding the basin of lake victoria. Preprint at 10.32942/osf.io/eu6rx (2021).

68. Malinsky, M. et al. Genomic islands of speciation separate cichlid ecomorphs in an east african crater lake. Science 350, 1493–1498, 10.1126/science.aac9927 (2015).

69. Njaya, F. et al. The natural history and fisheries ecology of lake chilwa, southern malawi. J. Gt. Lakes Res. 37, 15–25, 10.1016/j.jglr.2010.09.008 (2011).

70. Parsons, P. J., Bridle, J. R., Rüber, L. & Genner, M. J. Evolutionary divergence in life history traits among populations of the lake malawi cichlid fish astatotilapia calliptera. Ecol. evolution 7, 8488–8506, 10.1002/ece3.3311 (2017).

71. Omland, K. E. The assumptions and challenges of ancestral state reconstructions. Syst. biology 48, 604–611, 10.1080/106351599260175 (1999).

72. Duponchelle, F., Ribbink, A., Msukwa, A., Mafuka, J. & Mandere, D. Depth distribution and breeding patterns of the demersal species most commonly caught by trawling in the south west arm of lake malawi.Available at https://malawicichlids.com/duponchelle_ch2.pdf (2000).

73. Turner, G. F. A new species of deep-water lethrinops (cichlidae) from lake malawi. J. Fish Biol. 101, 1405–1410, 10.1111/jfb.15208 (2022).

74. Stauffer, J. & Konings, A. Review of copadichromis (teleostei: Cichlidae) with the description of a new genus and six new species. Ichthyol. Explor. Freshwaters 17, 9–42 (2006).

75. Turner, G. F., Crampton, D. A., Rusuwa, B., Hooft van Huysduynen, A. & Svardal, H. Taxonomic investigation of the zooplanktivorous lake malawi cichlids copadichromis mloto (iles) and c. virginalis (iles). Hydrobiologia 1–11, 10.1007/s10750-022-05025-1 (2022).

76. Turner, G. F., Crampton, D. A. & Genner, M. J. A new species of lethrinops (cichliformes: Cichlidae) from a lake malawi satellite lake, believed to be extinct in the wild. Preprint at 10.1101/2023.03.17.533142 (2023).

77. Kikinis, R., Pieper, S. D. & Vosburgh, K. G. 3D Slicer: A Platform for Subject-Specific Image Analysis, Visualization, and Clinical Support (Springer New York, New York, NY, 2014).

78. Haberthür, D. et al. Microtomographic investigation of a large corpus of cichlids. Plos one 18, e0291003, 10.1371/journal.pone.0291003 (2023).

