## Supplementary figures and images for "A whole-body micro-CT scan library that captures the skeletal diversity of Lake Malawi cichlid fishes"

### Rhamphochromis_sp_kingiri_large_UniBri_site1_8bit_a.tif

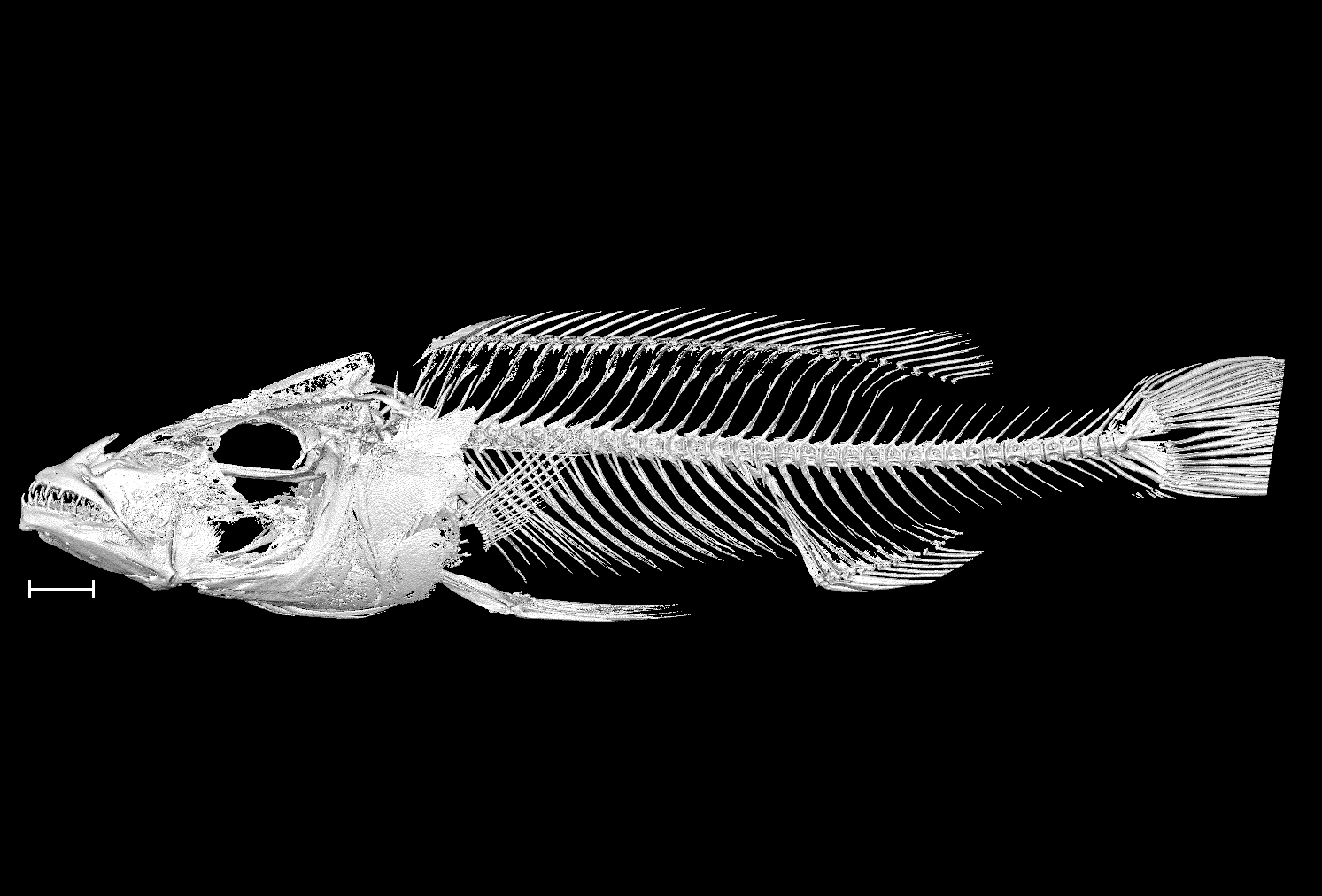

### Rhamphochromis_sp_kingiri_large_UniBri_site1_8bit_b.tif

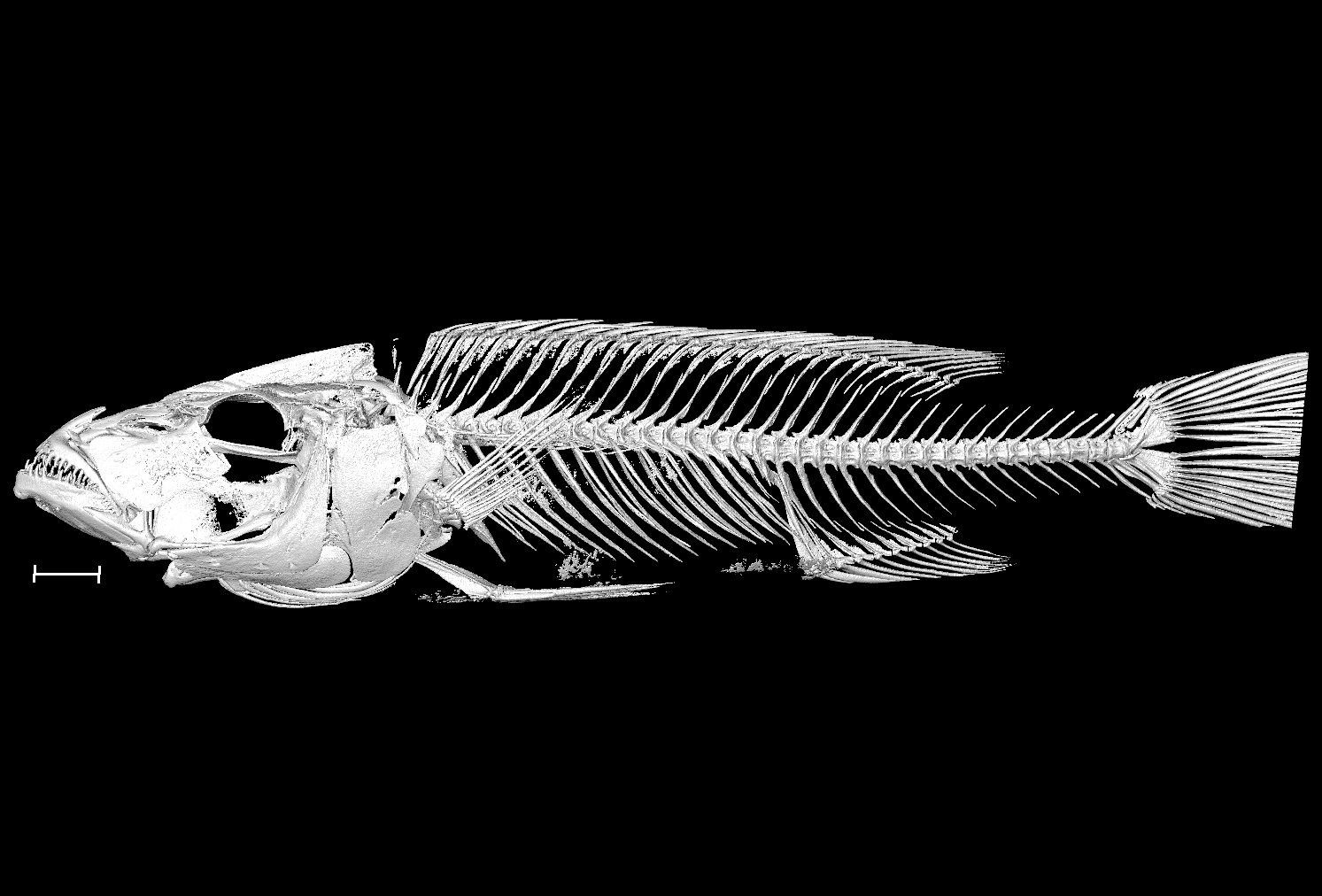

### Rhamphochromis_sp_longiceps_blue_back_UniBri_RRC037_1_8bit.tif

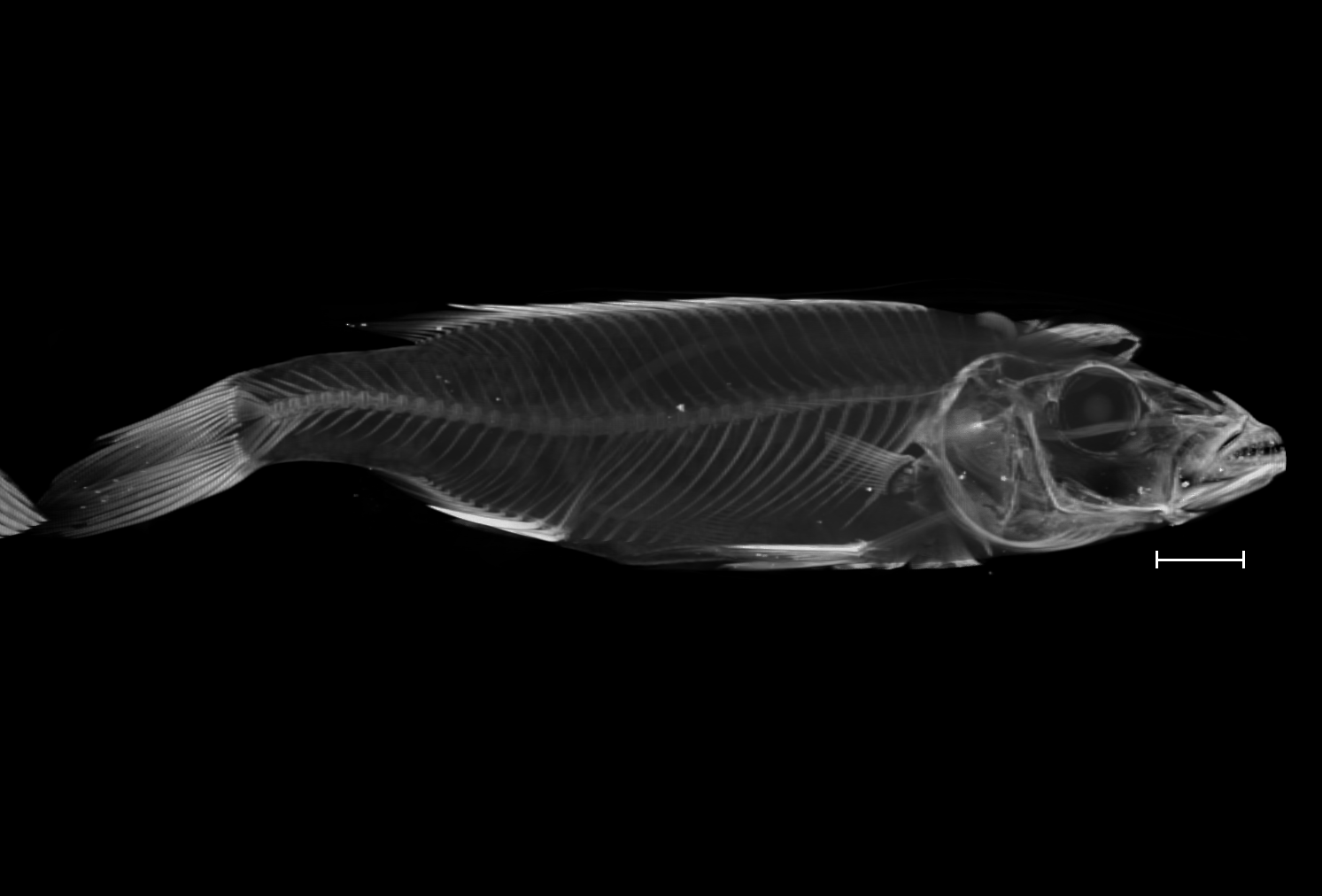

### Rhamphochromis_sp_longiceps_blue_back_UniBri_RRC037_3_8bit.tif

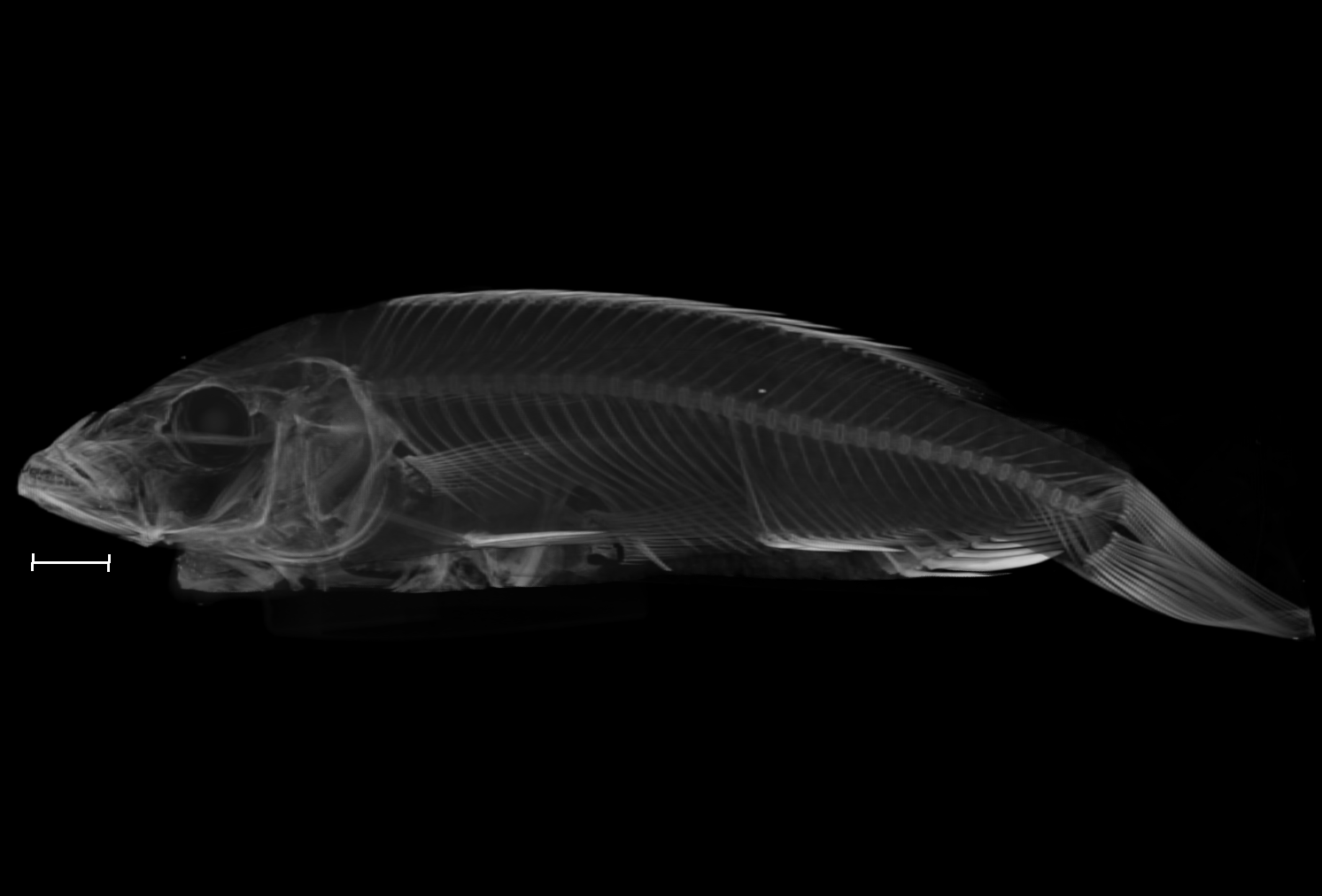

### Rhamphochromis_sp_longiceps_grey_back_UniBri_RRC045_8bit_a.tif

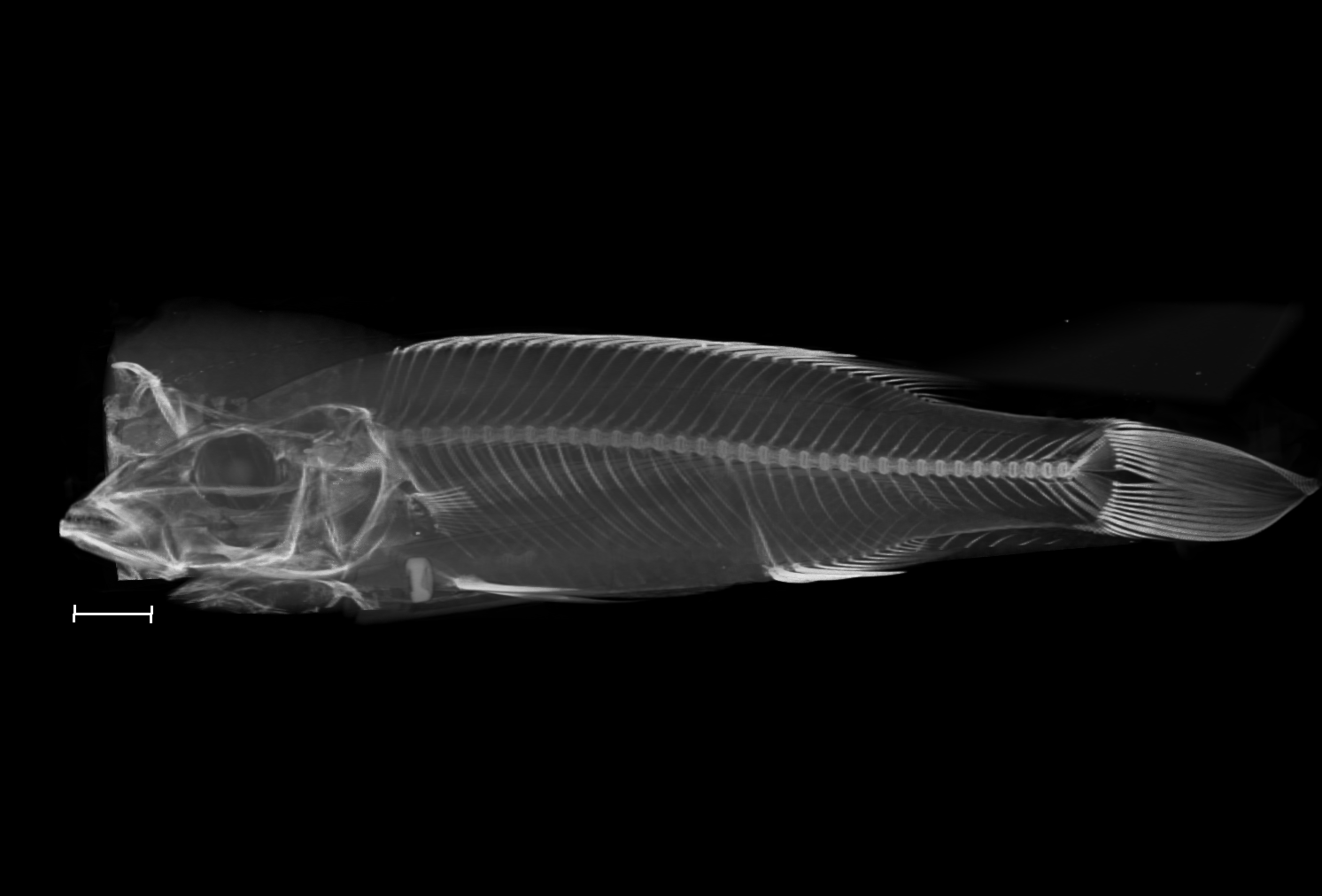

### Rhamphochromis_sp_longiceps_grey_back_UniBri_RRC045_8bit_b.tif

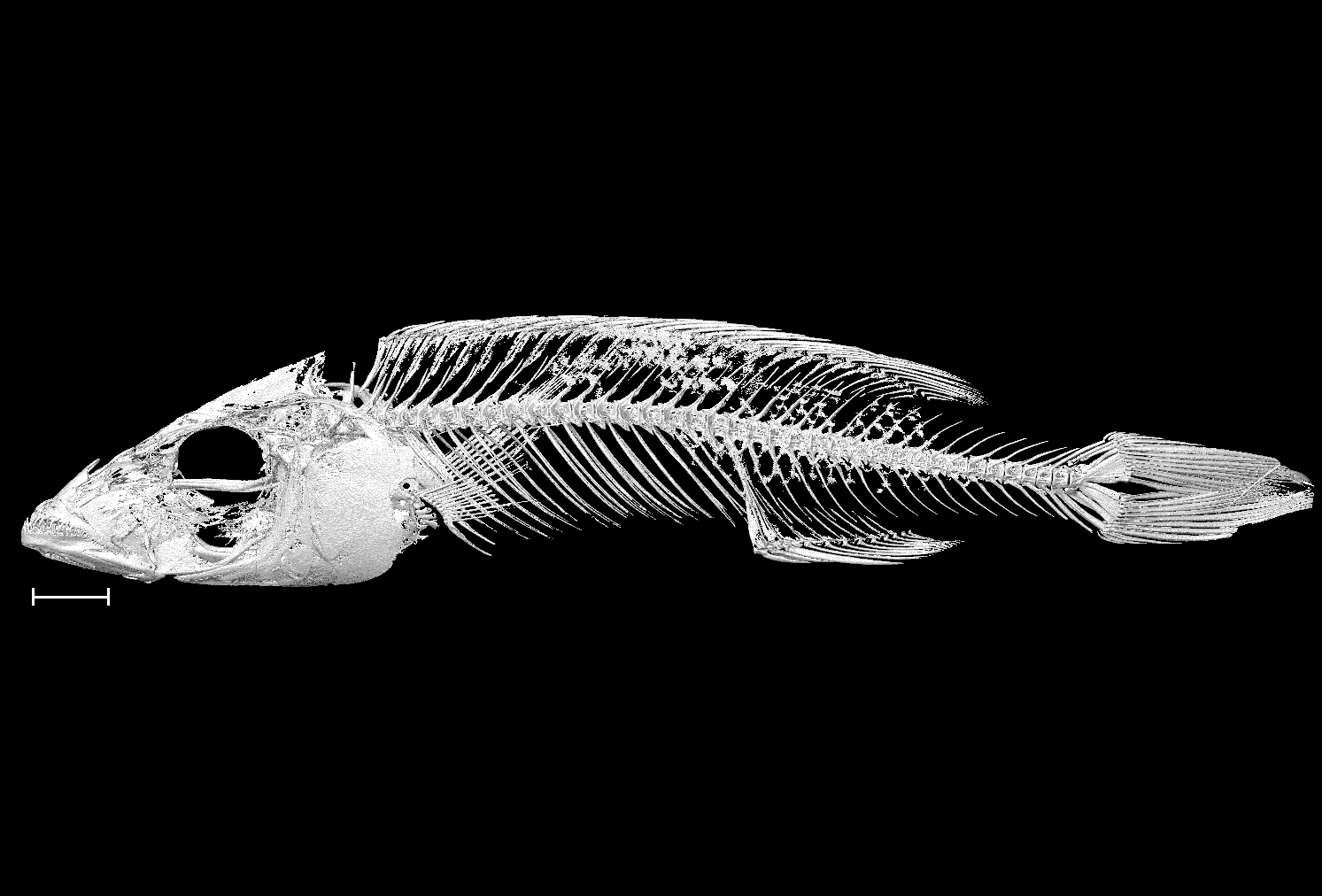

### Rhamphochromis_sp_yellow_belly_UniBri_RRC049_8bit_a.tif

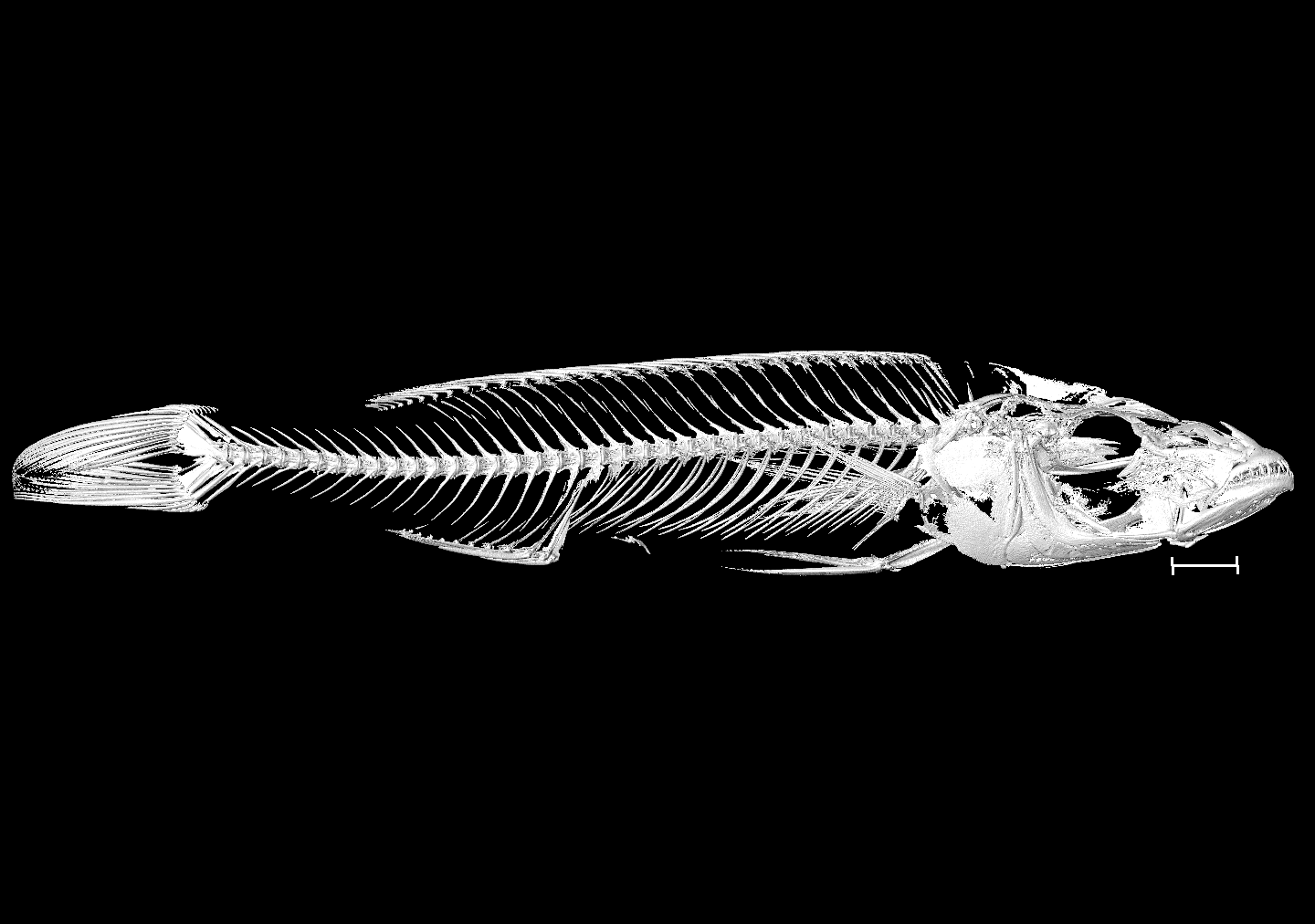

### Rhamphochromis_sp_yellow_belly_UniBri_RRC049_8bit_b.tif

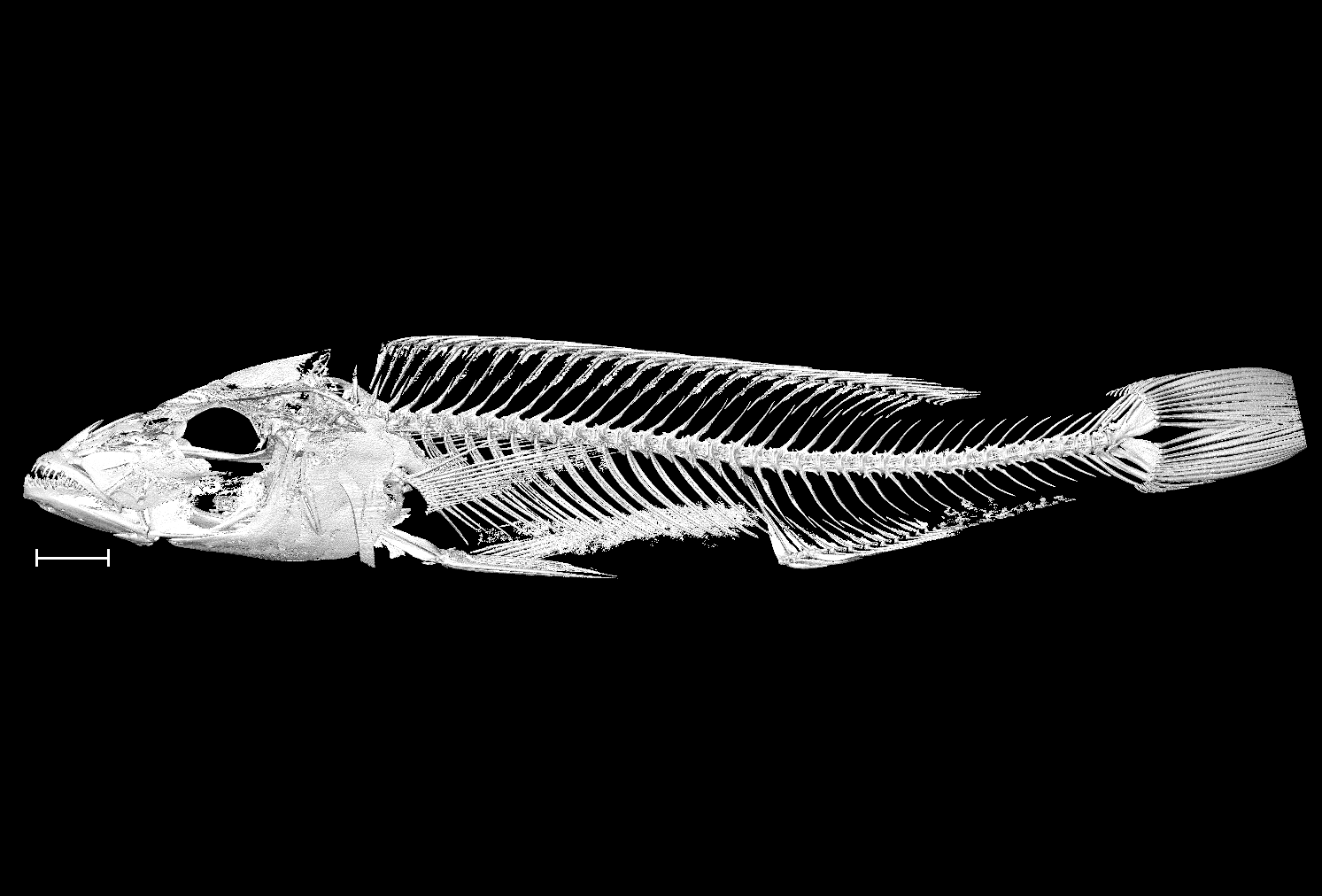

### Rhamphochromis_woodi_NHMUK_1935_6_14_2185_2187_8bit_a.tif

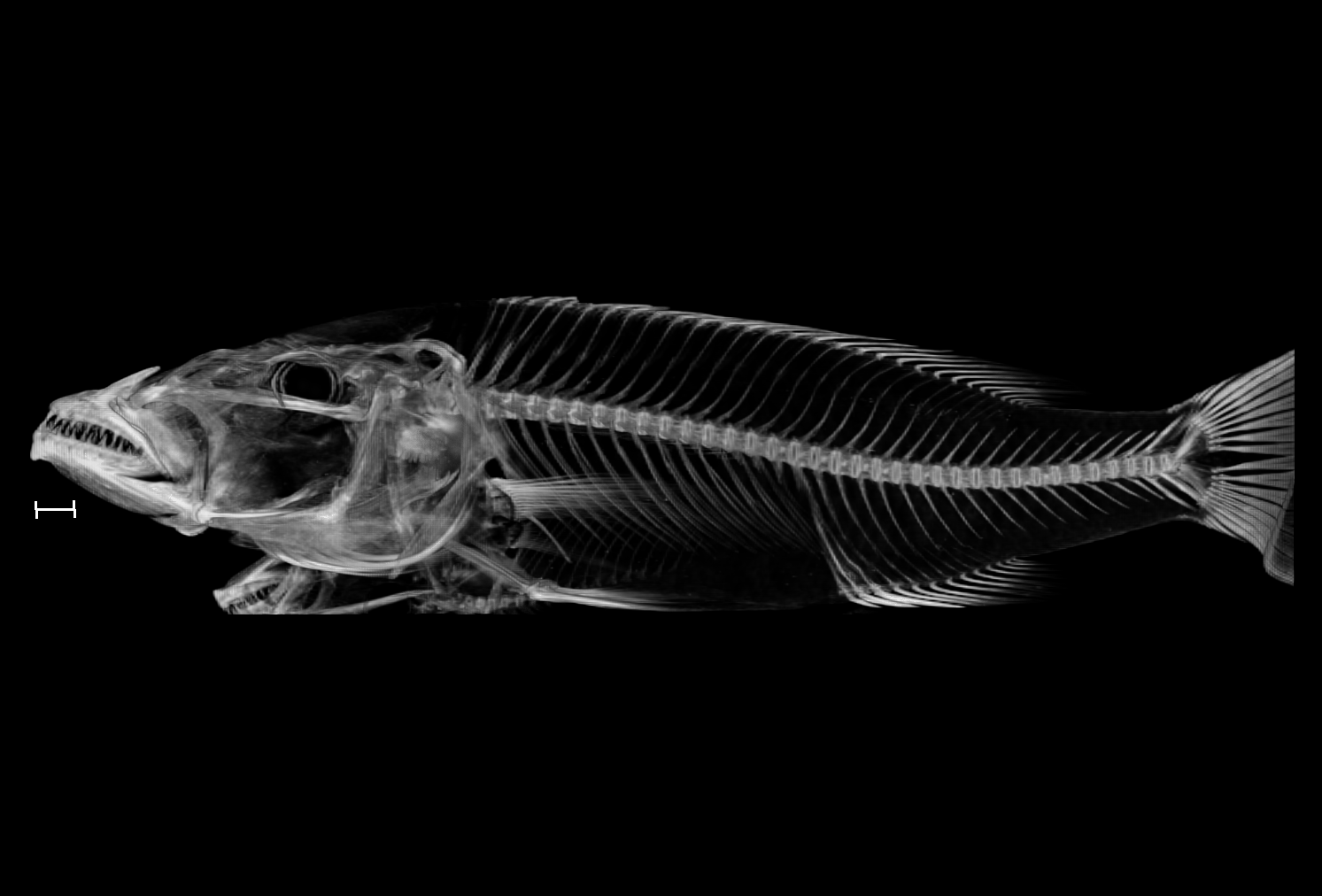

### Rhamphochromis_woodi_NHMUK_1935_6_14_2185_2187_8bit_b.tif

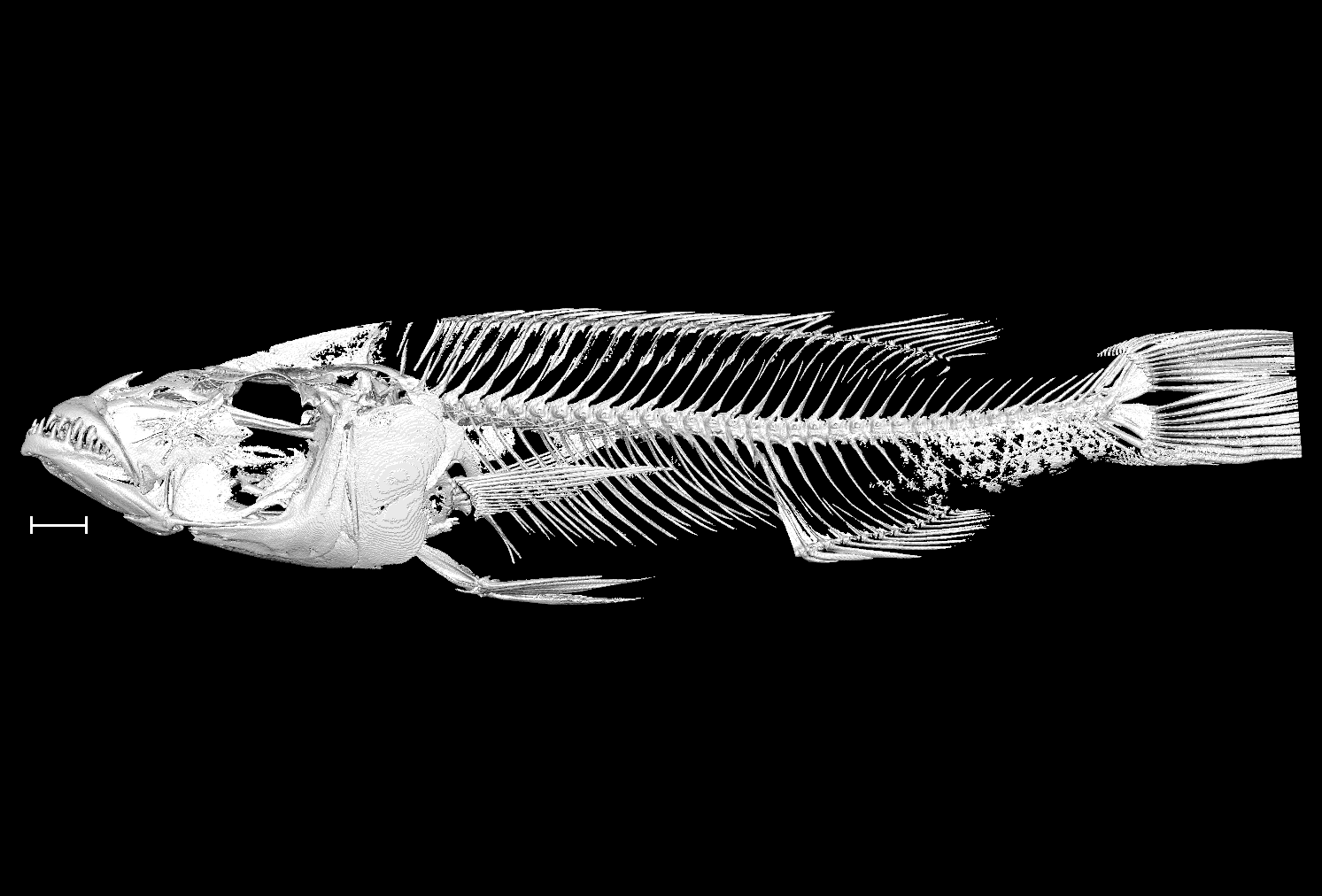

### Stigmatochromis_macrorhynchos_UniBri_SG9_8bit.tif

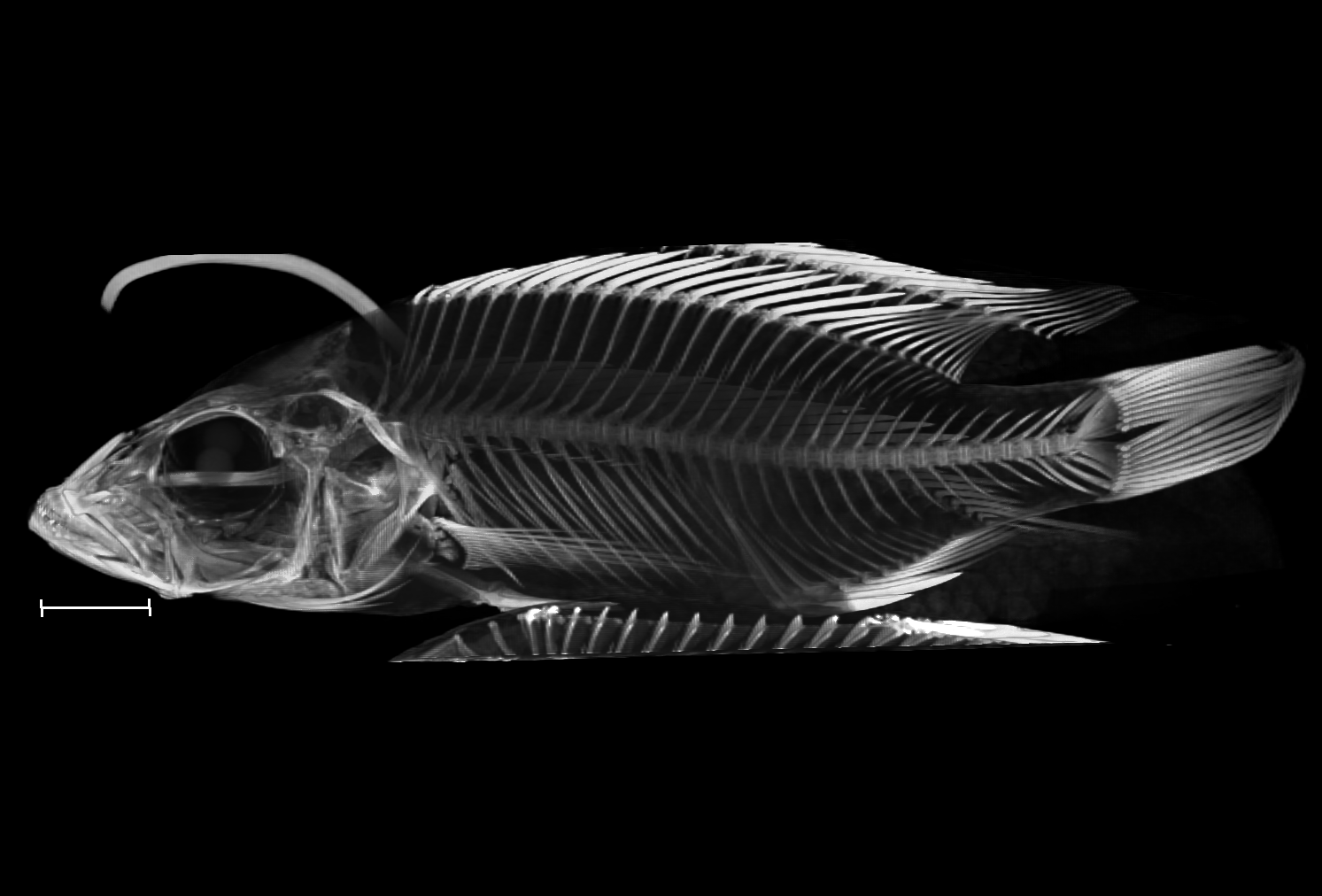

### Taeniolethrinops_praeorbitalis_UniBri_31_9_14_8bit.tif

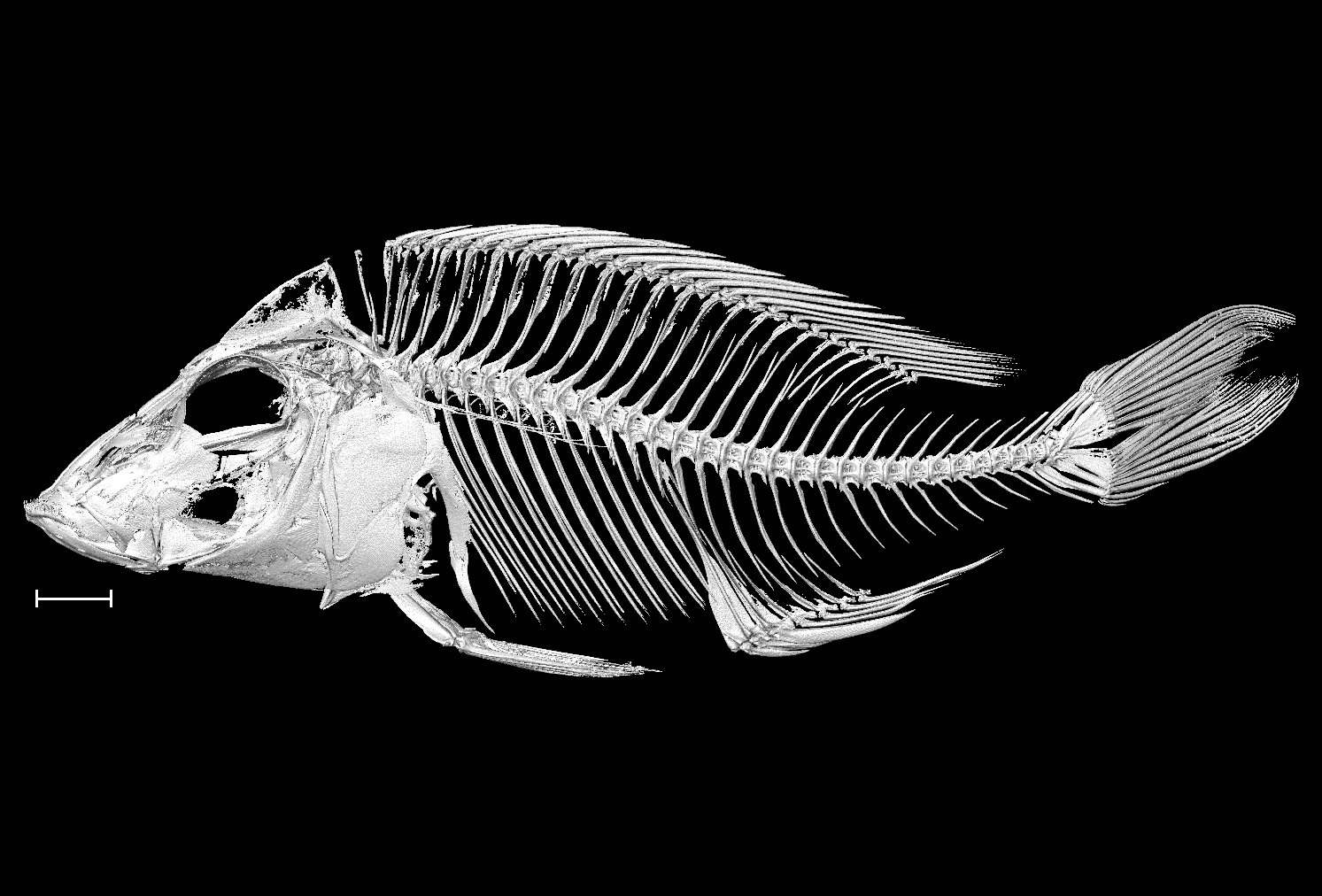

### Trematocranus_placodon_UniBri_220_8bit.tiff

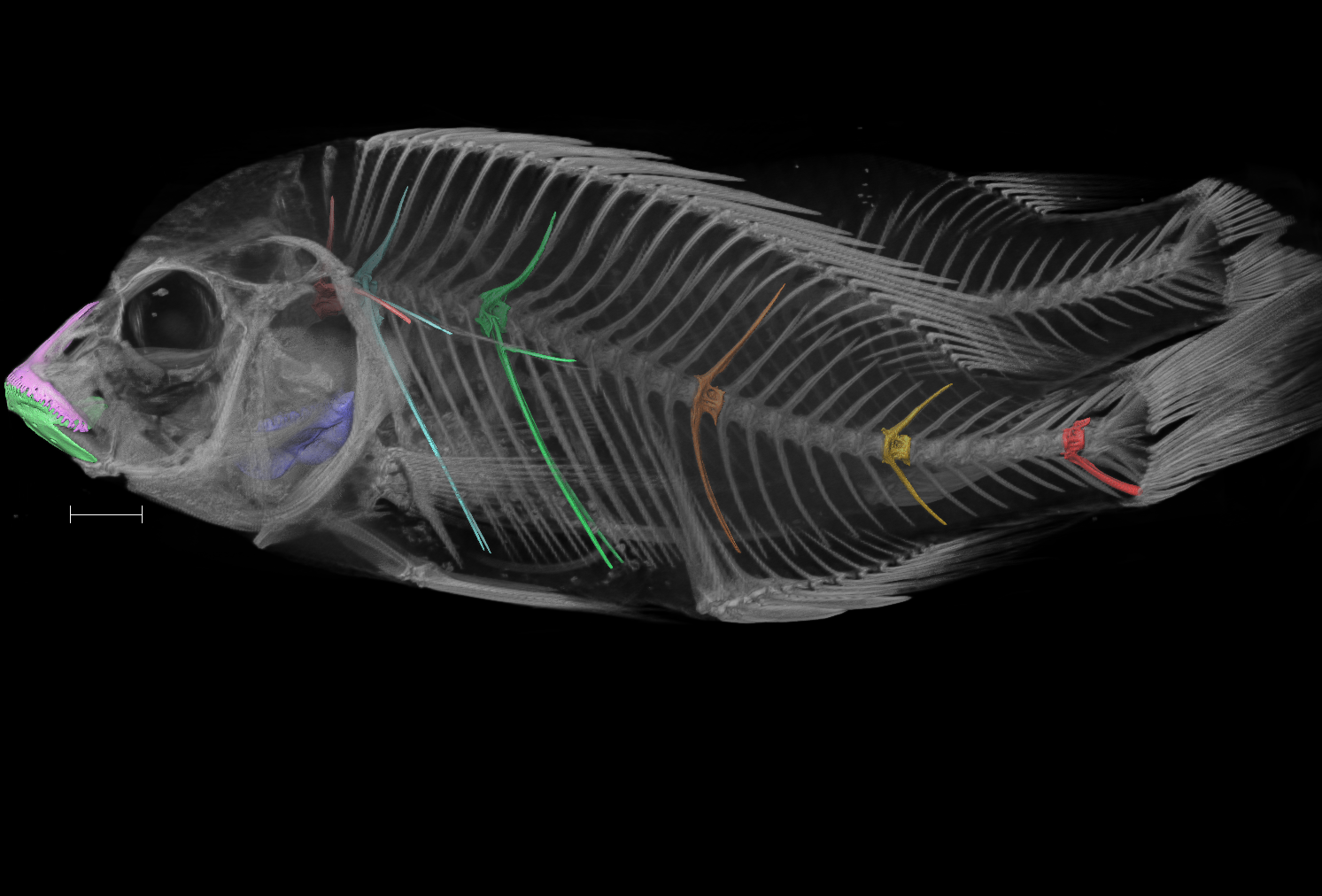

### Tropheops_tropheops_NHMUK_2012_1_17_18_25_8bit_a.tif

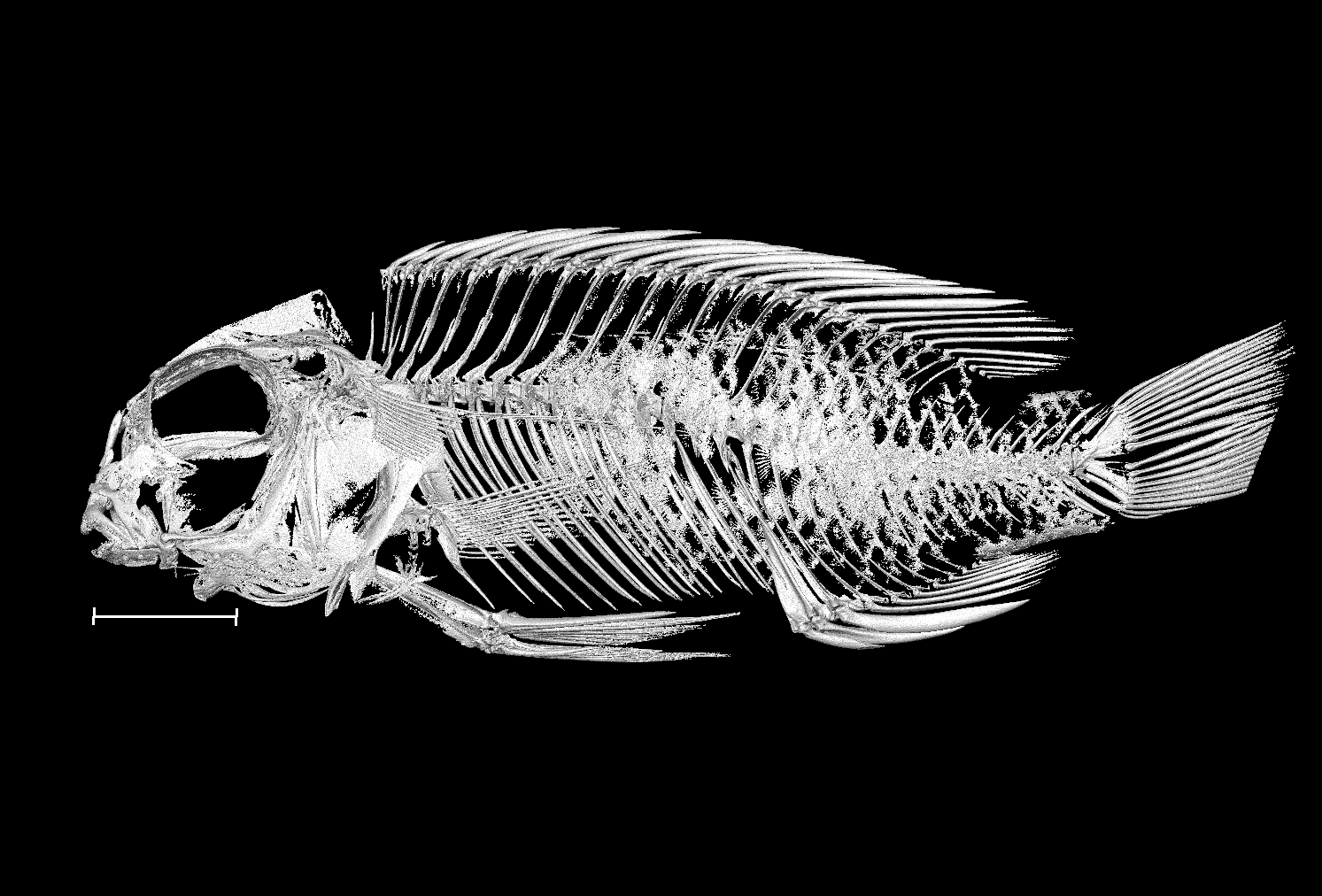

### Tropheops_tropheops_NHMUK_2012_1_17_18_25_8bit_b.tif

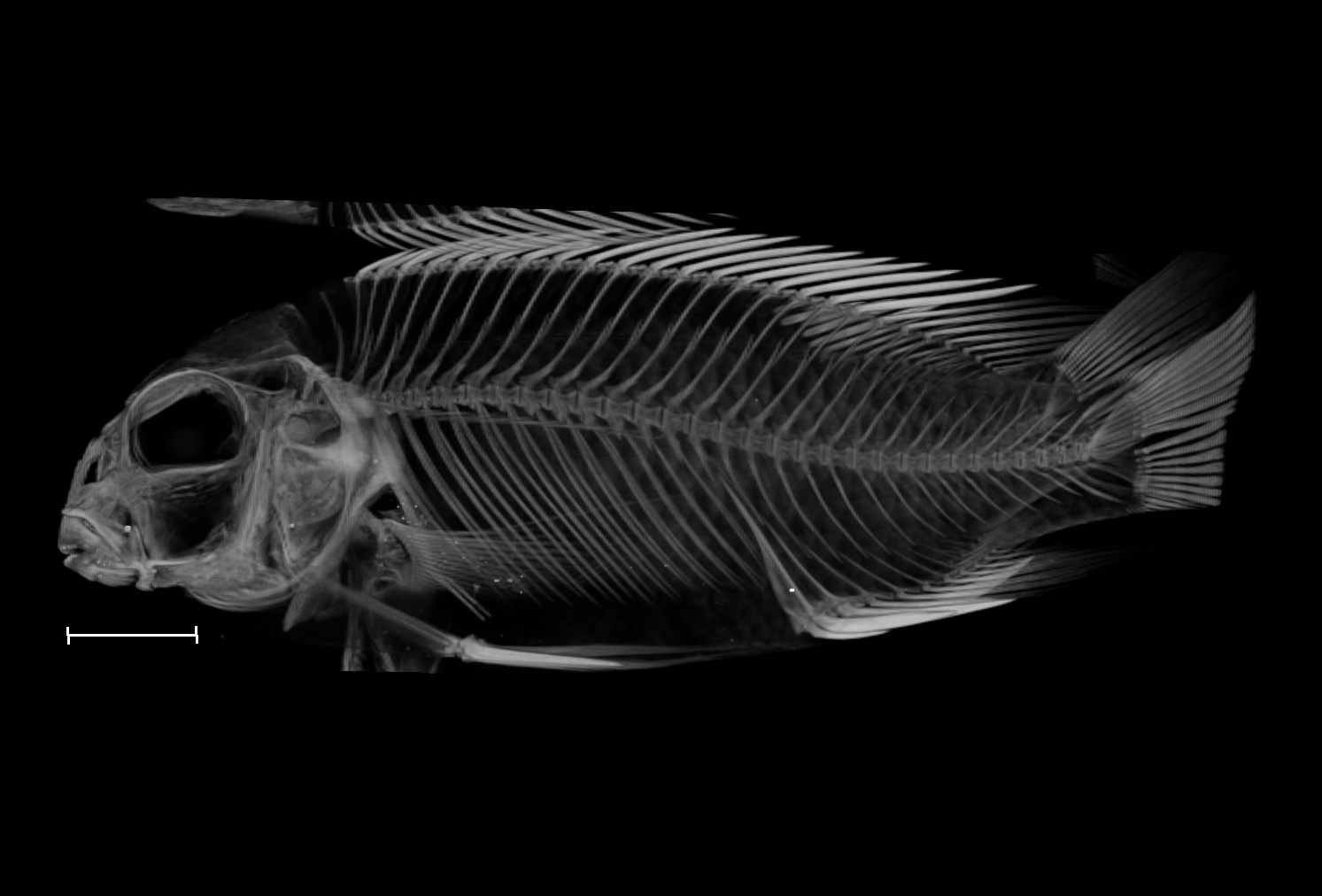

### Tyrannochromis_macrostoma_UniBri_201_8bit.tif

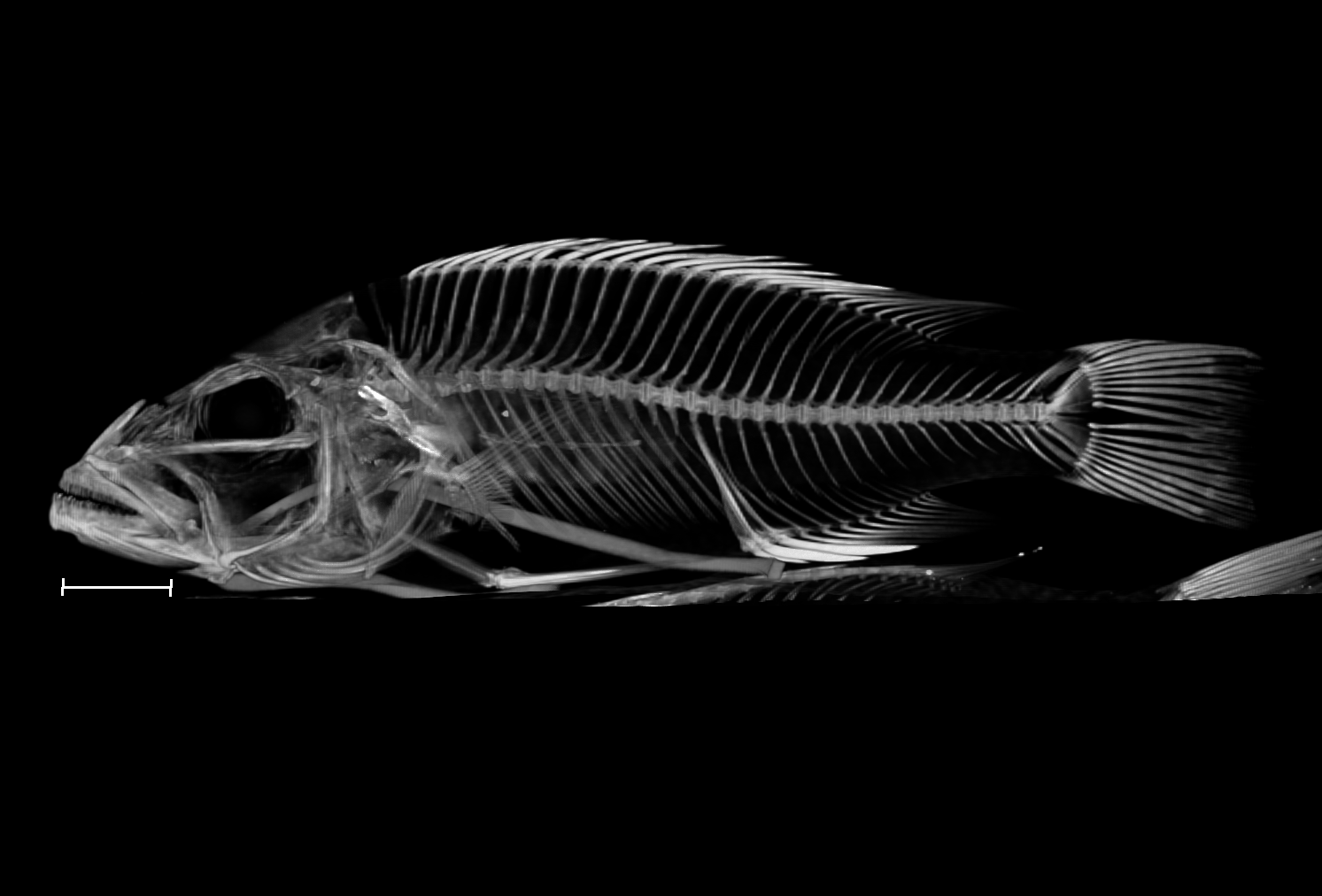
