## Supplementary figures and images for "A whole-body micro-CT scan library that captures the skeletal diversity of Lake Malawi cichlid fishes"

### Alticorpus_macrocleithrum_NHMUK_1984_10_22_2_5_8bit_a.tif

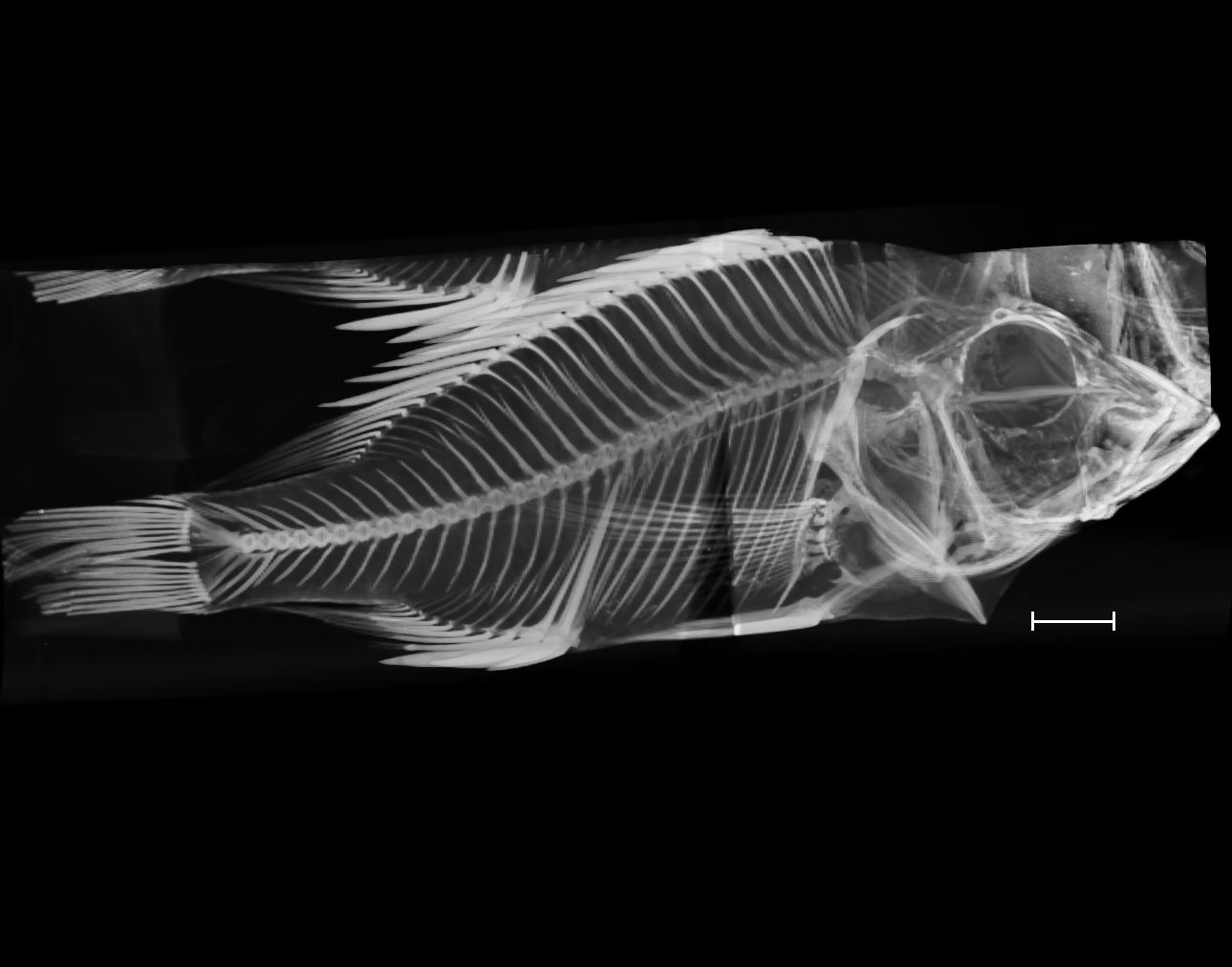

### Alticorpus_macrocleithrum_NHMUK_1984_10_22_2_5_8bit_b.tif

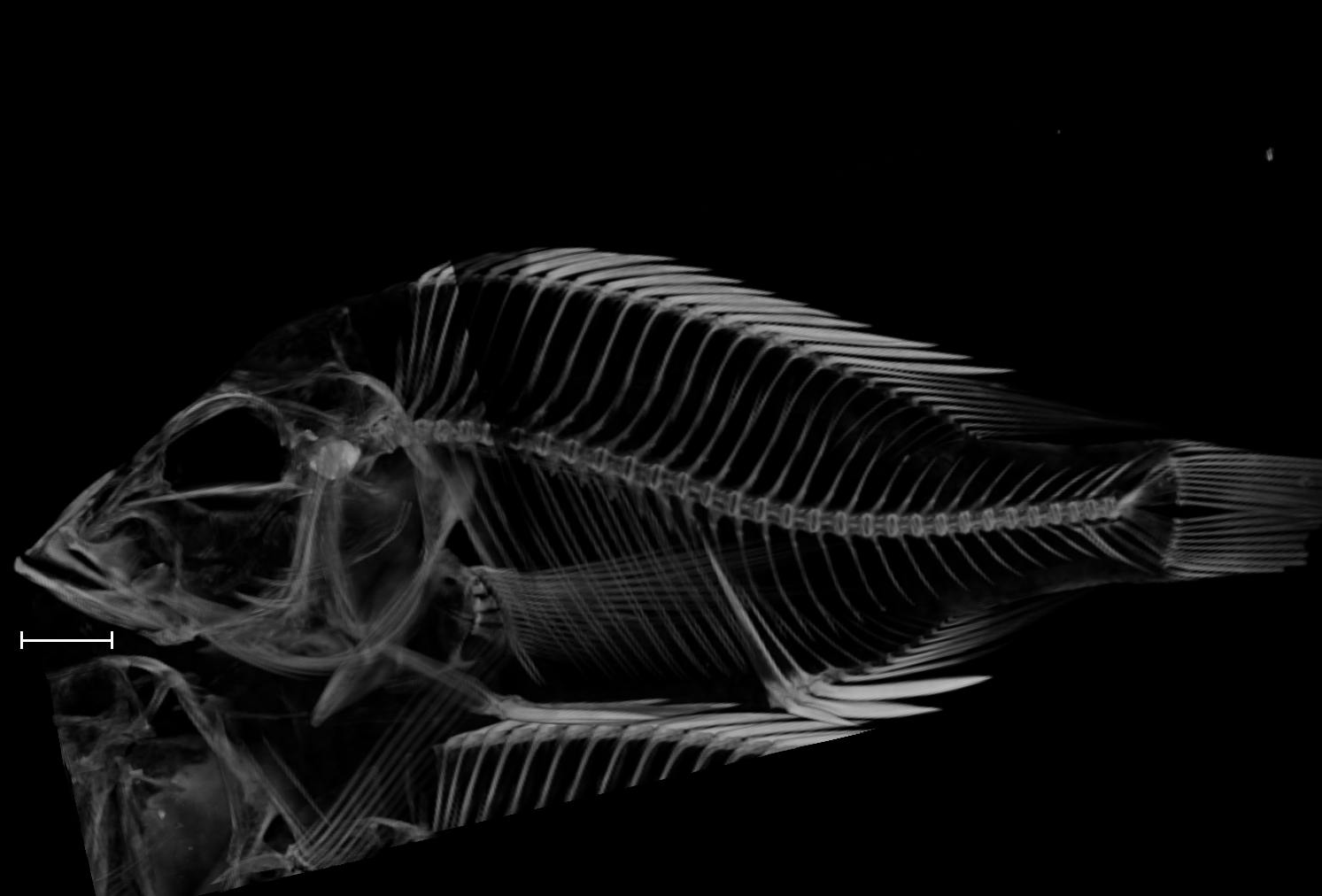

### Alticorpus_macrocleithrum_UniBri_maldeco_19_10_04_8bit.tif

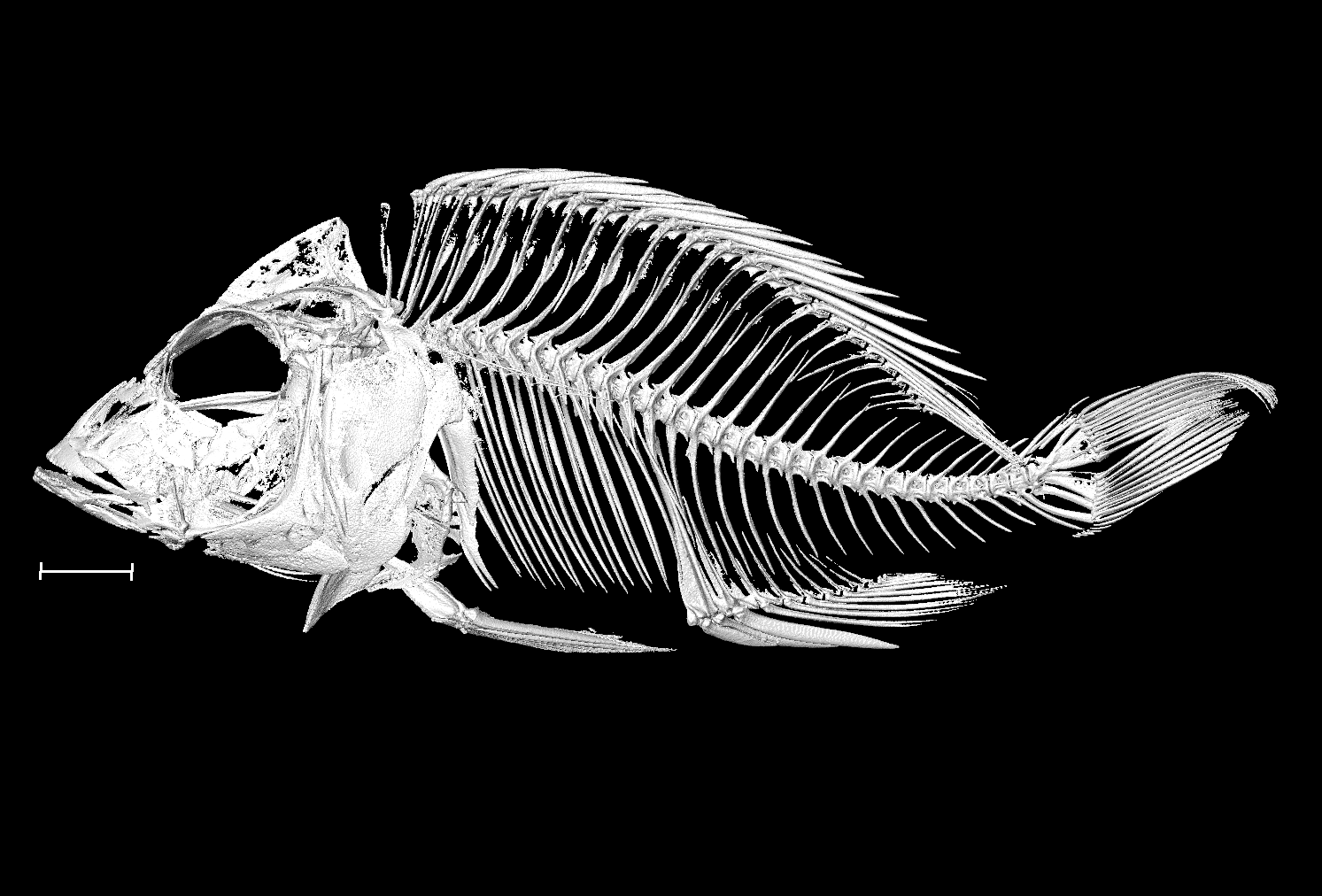

### Astatotilapia_calliptera_NHMUK_1921_9_6_94_95_8bit_a.tif

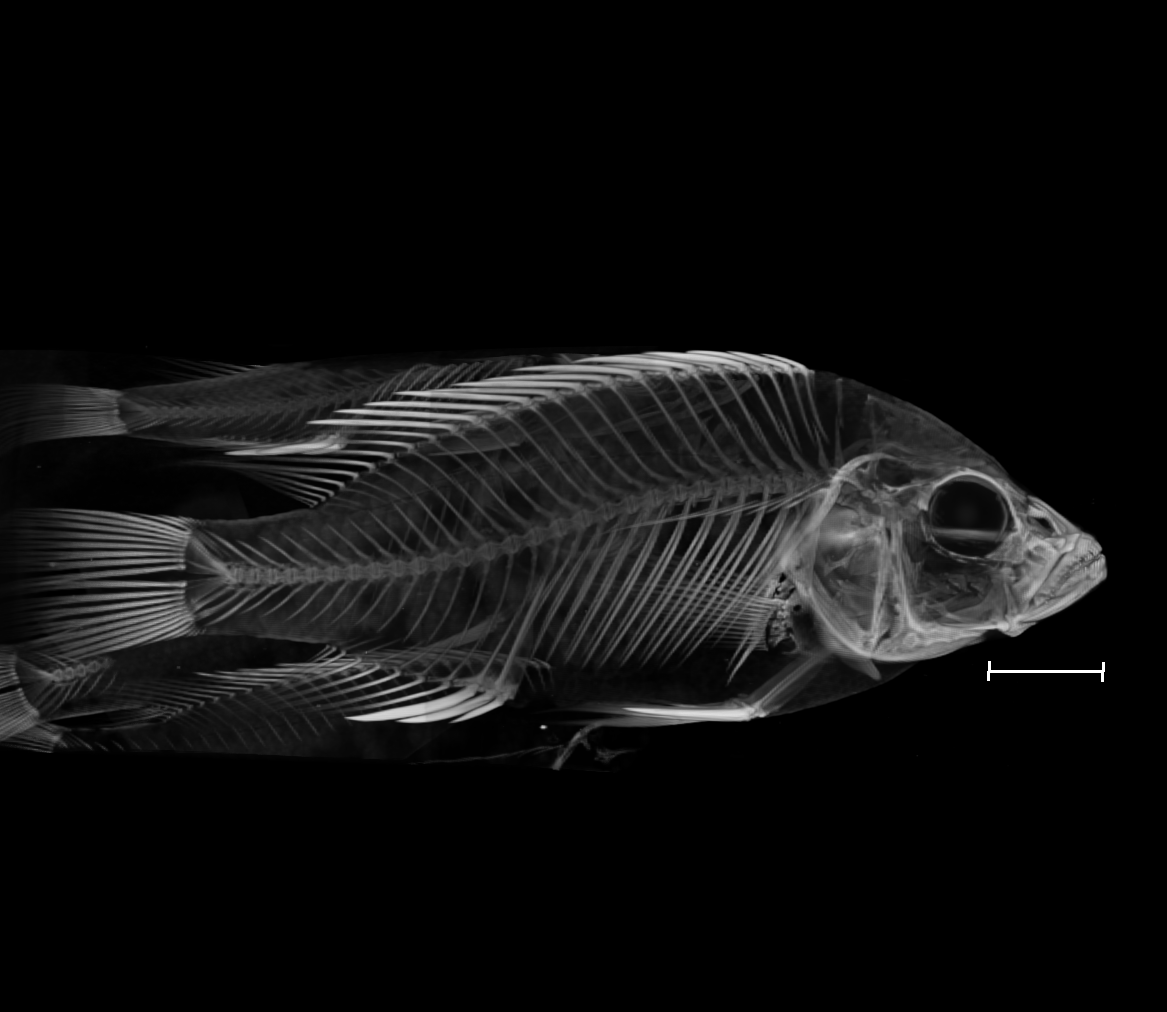

### Astatotilapia_calliptera_NHMUK_1921_9_6_94_95_8bit_b.tif

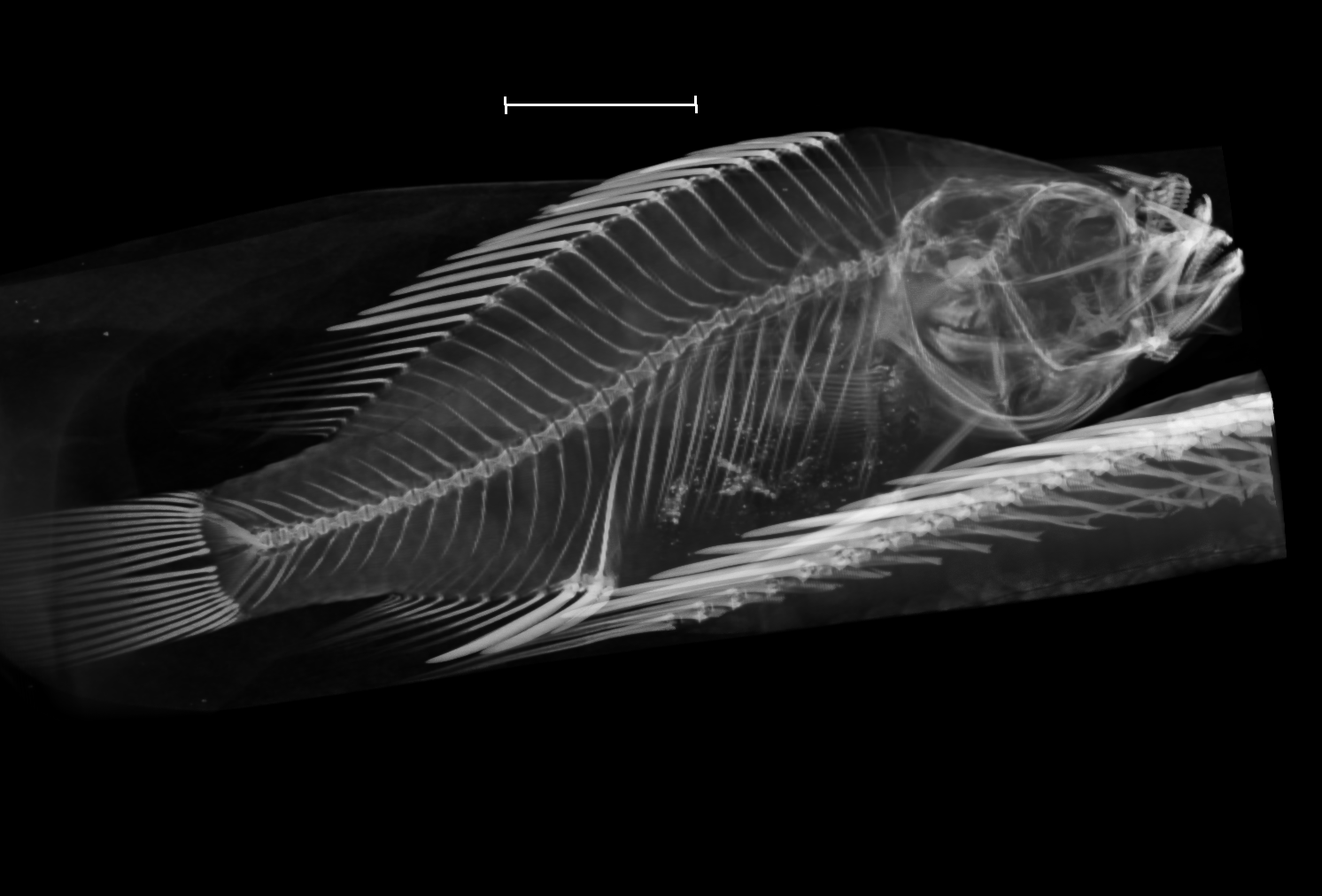

### Astatotilapia_calliptera_NHMUK_1957_11_29_16_26_8bit_a.tif

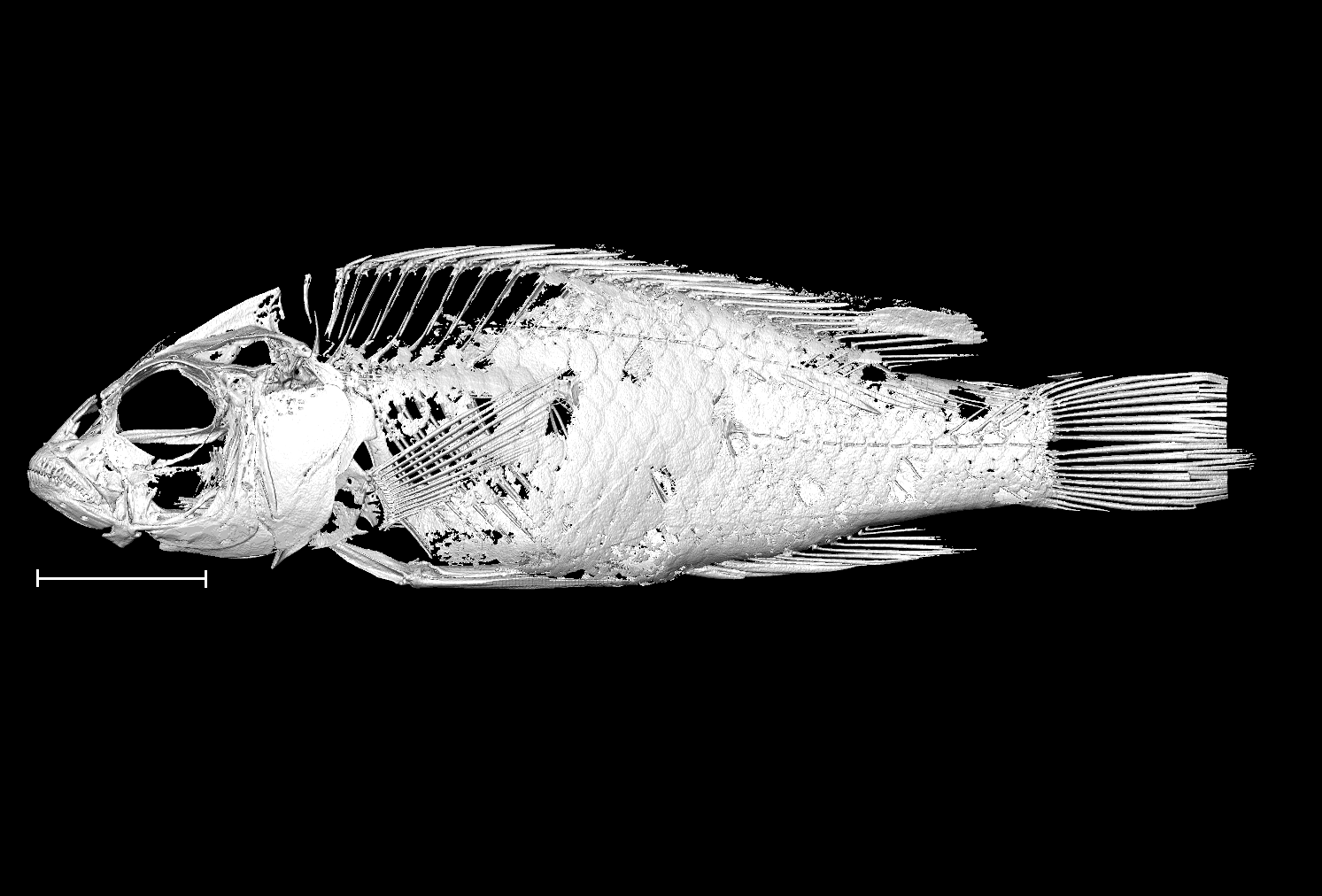

### Astatotilapia_calliptera_NHMUK_1957_11_29_16_26_8bit_b.tif

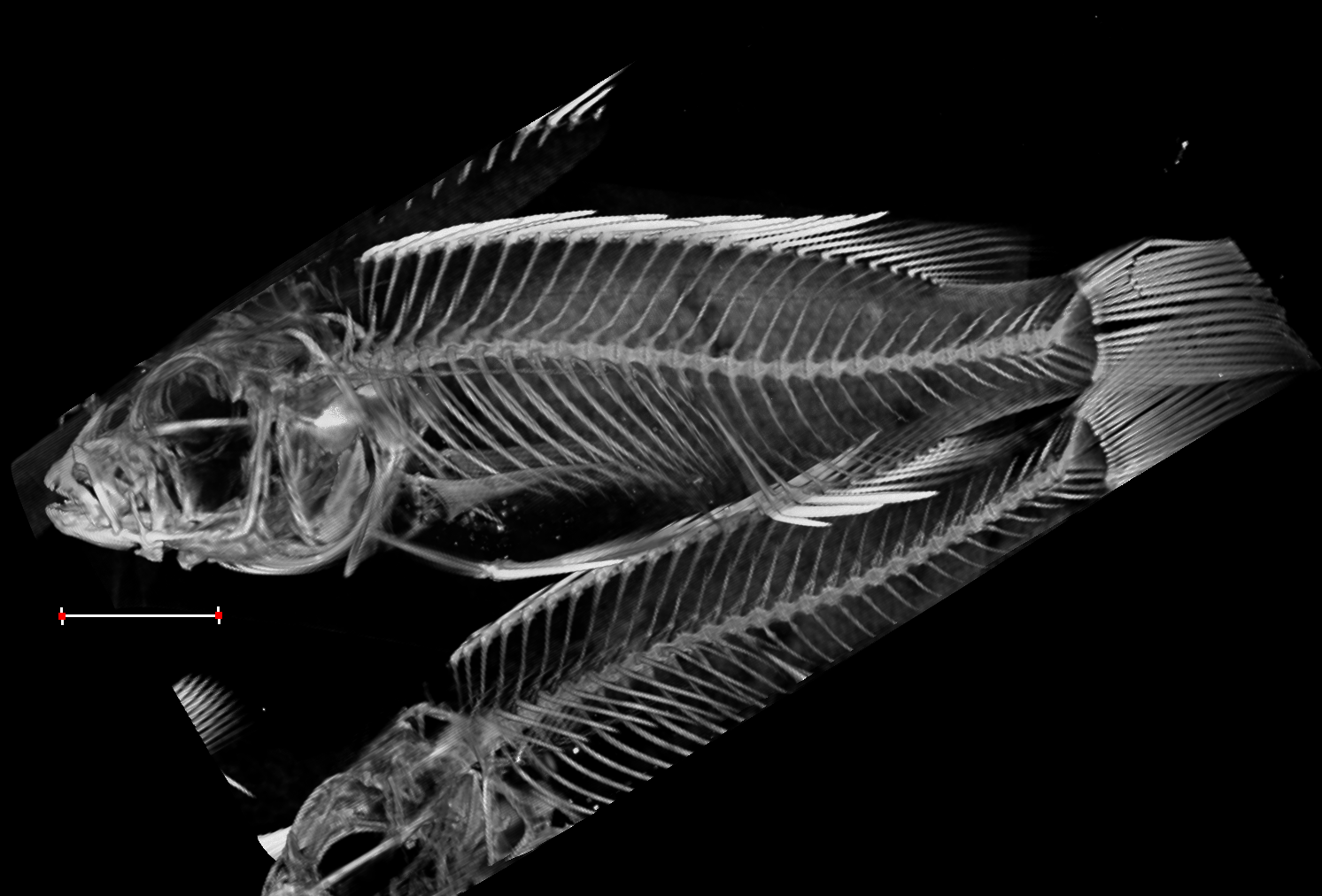

### Astatotilapia_calliptera_NHMUK_1966_7_28_21_24_8bit_a.tif

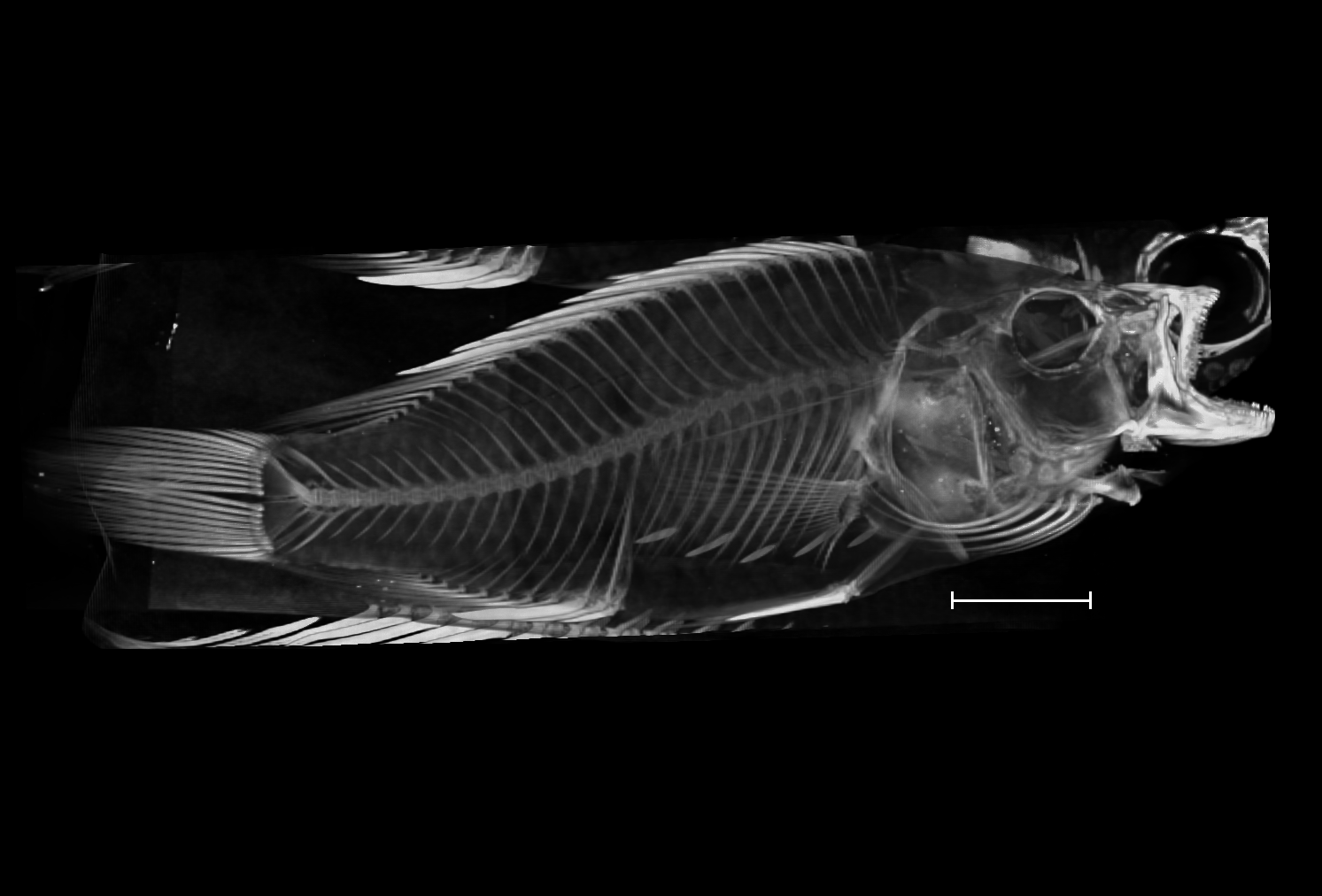

### Astatotilapia_calliptera_NHMUK_1966_7_28_21_24_8bit_b.tif

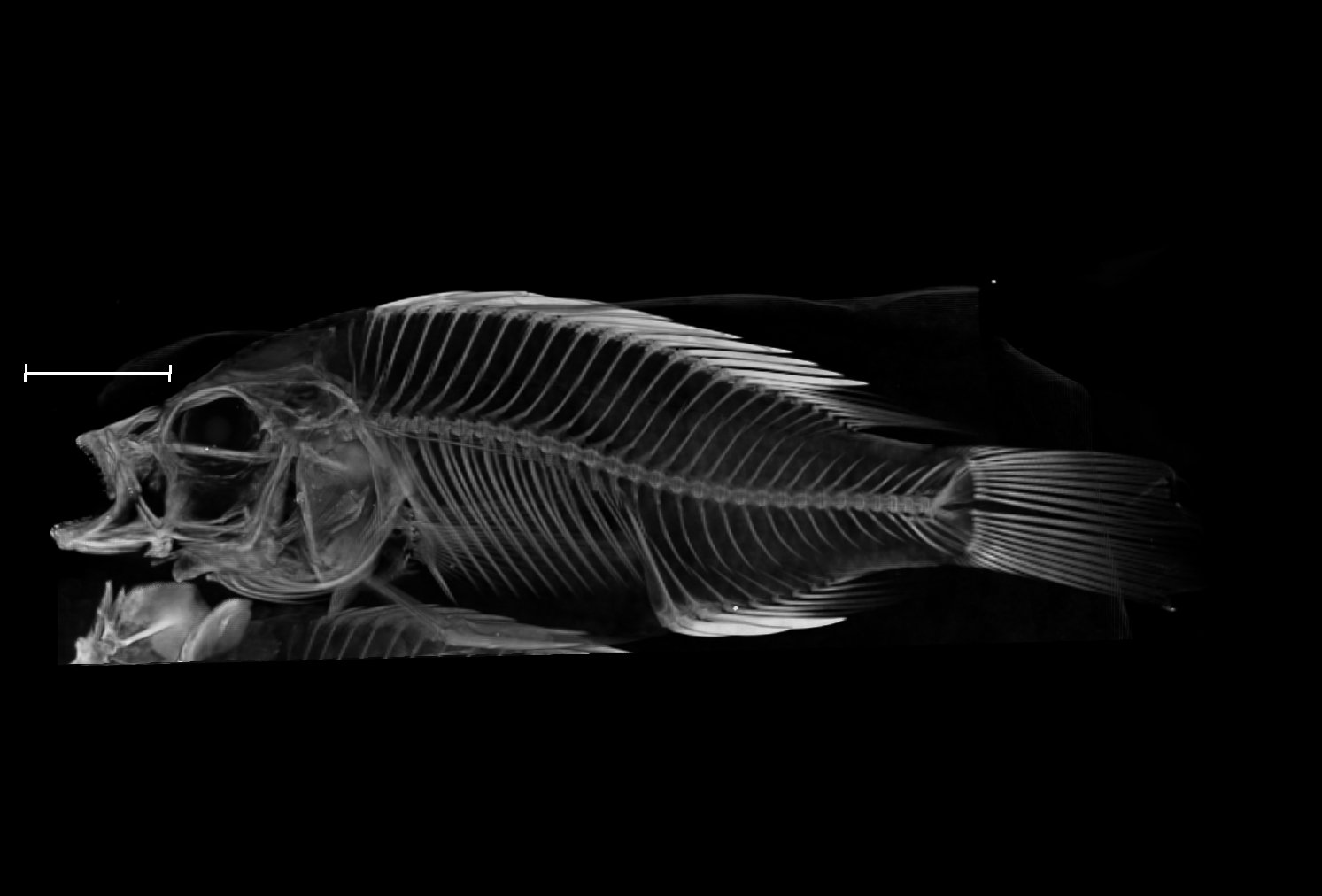

### Astatotilapia_calliptera_UniOxf_AC1_8bit.tif

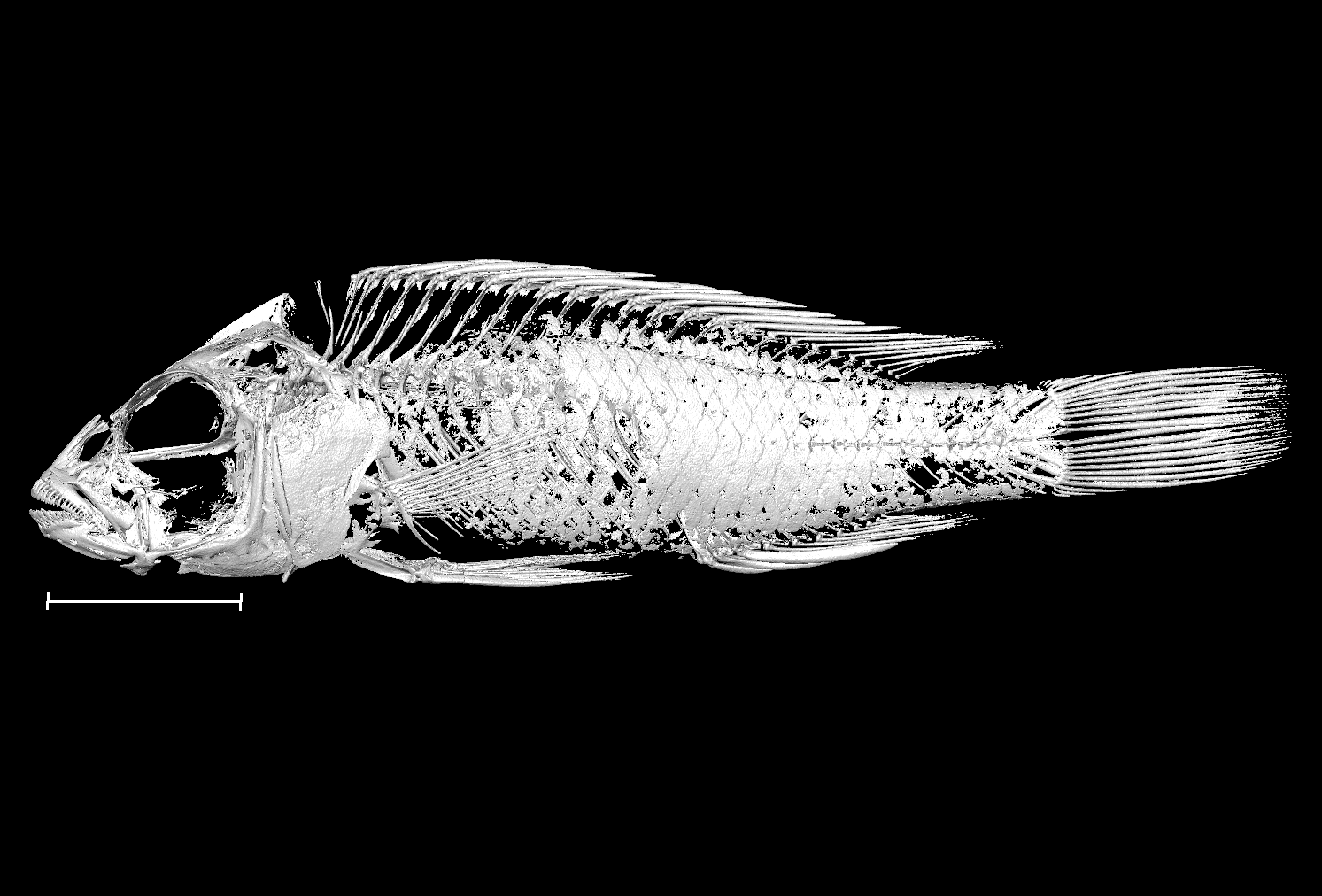

### Astatotilapia_calliptera_UniOxf_AC2_8bit.tif

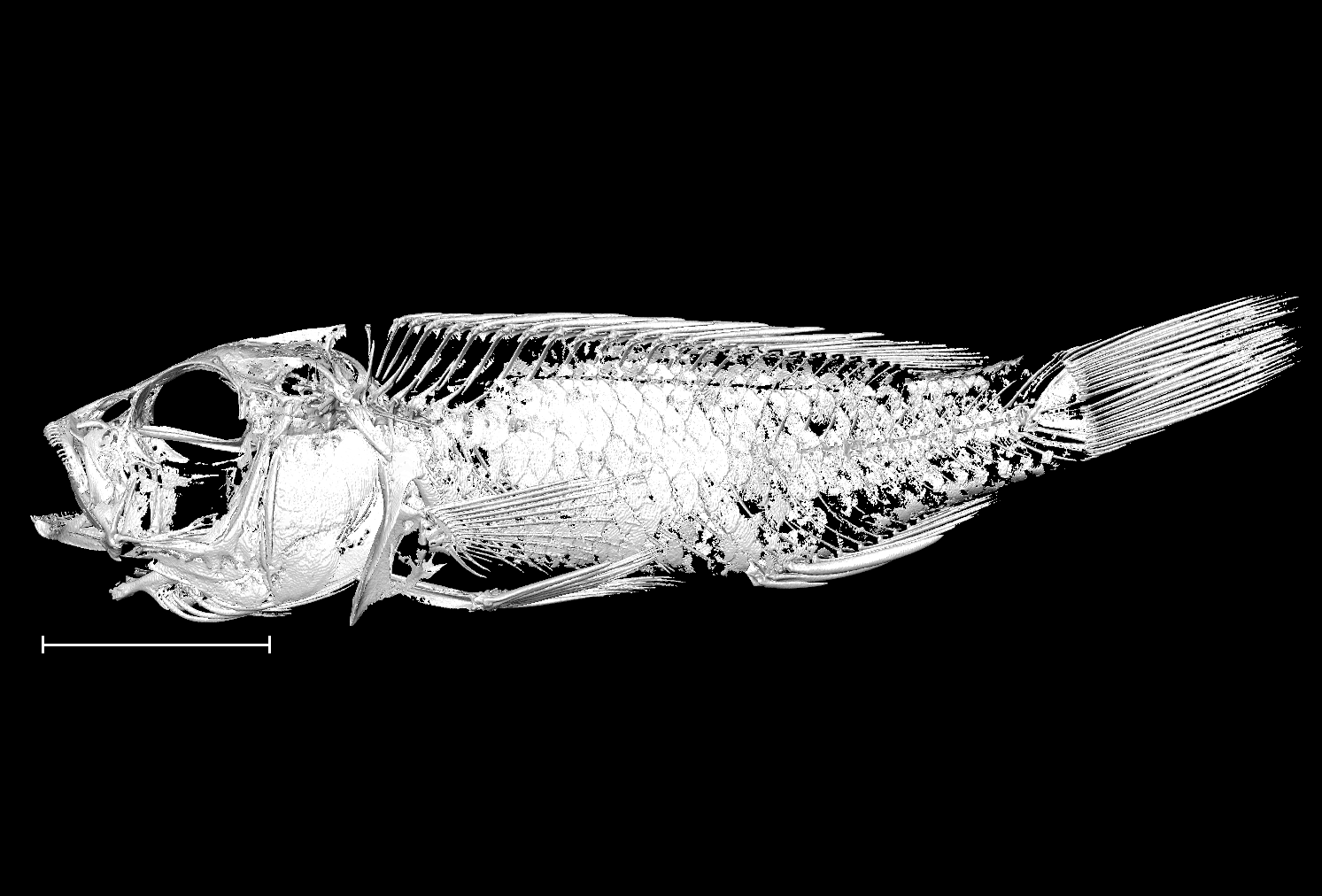

### Astatotilapia_calliptera_UniOxf_AC3_8bit.tif

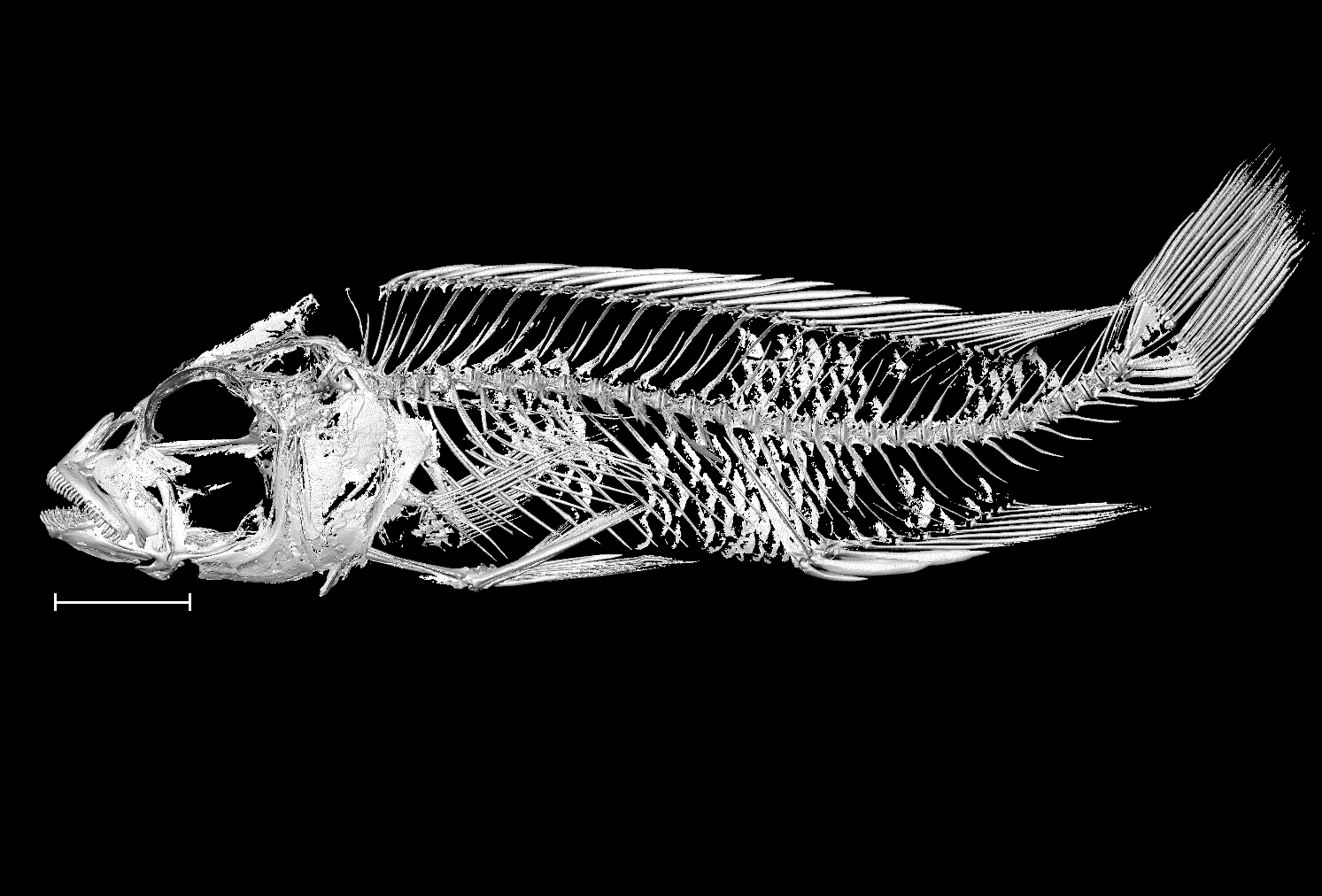

### Astatotilapia_calliptera_UniOxf_AC4_8bit.tif

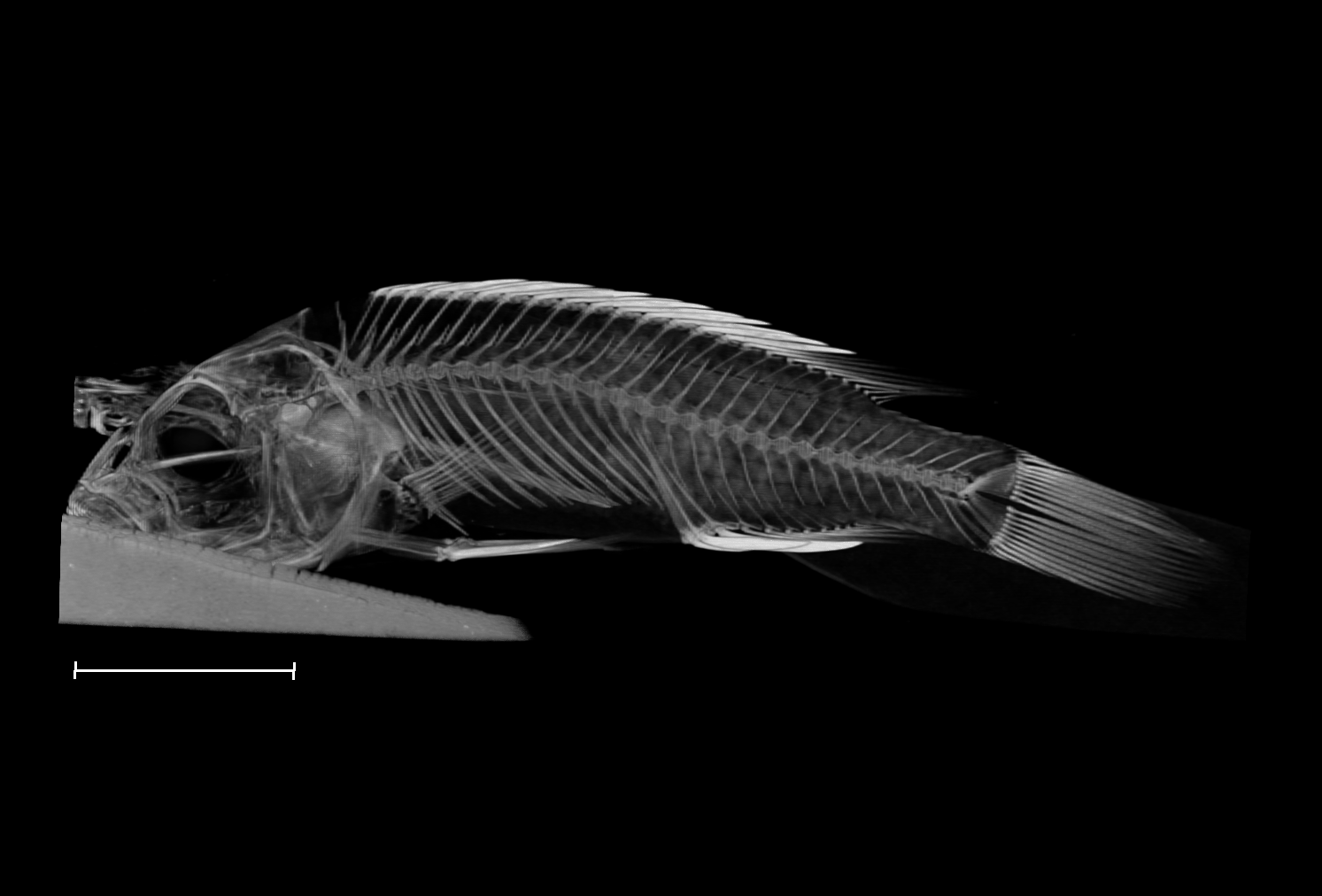

### Astatotilapia_calliptera_UniOxf_AC5_8bit.tif

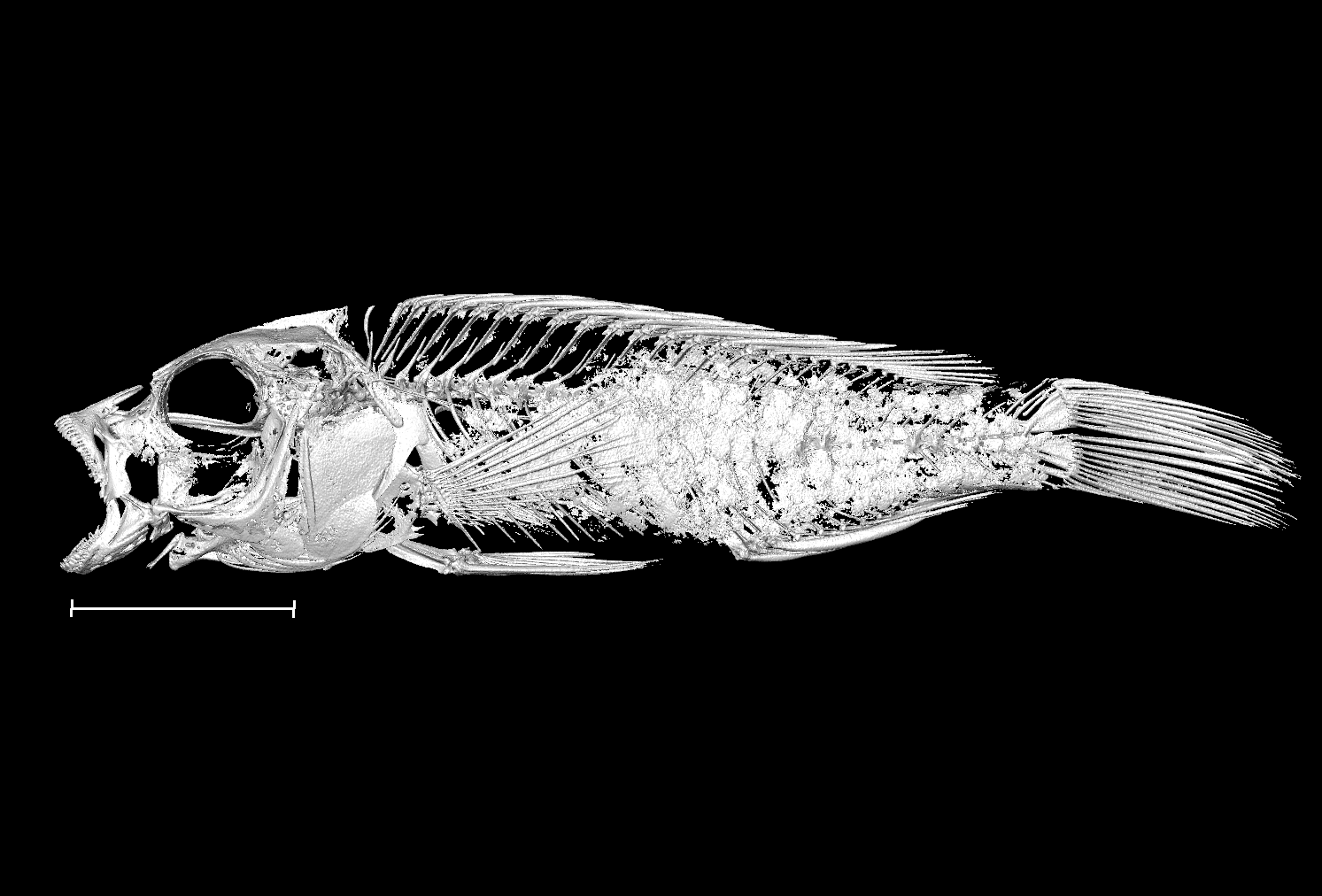

### Astatotilapia_calliptera_UniOxf_AC6_8bit.tif

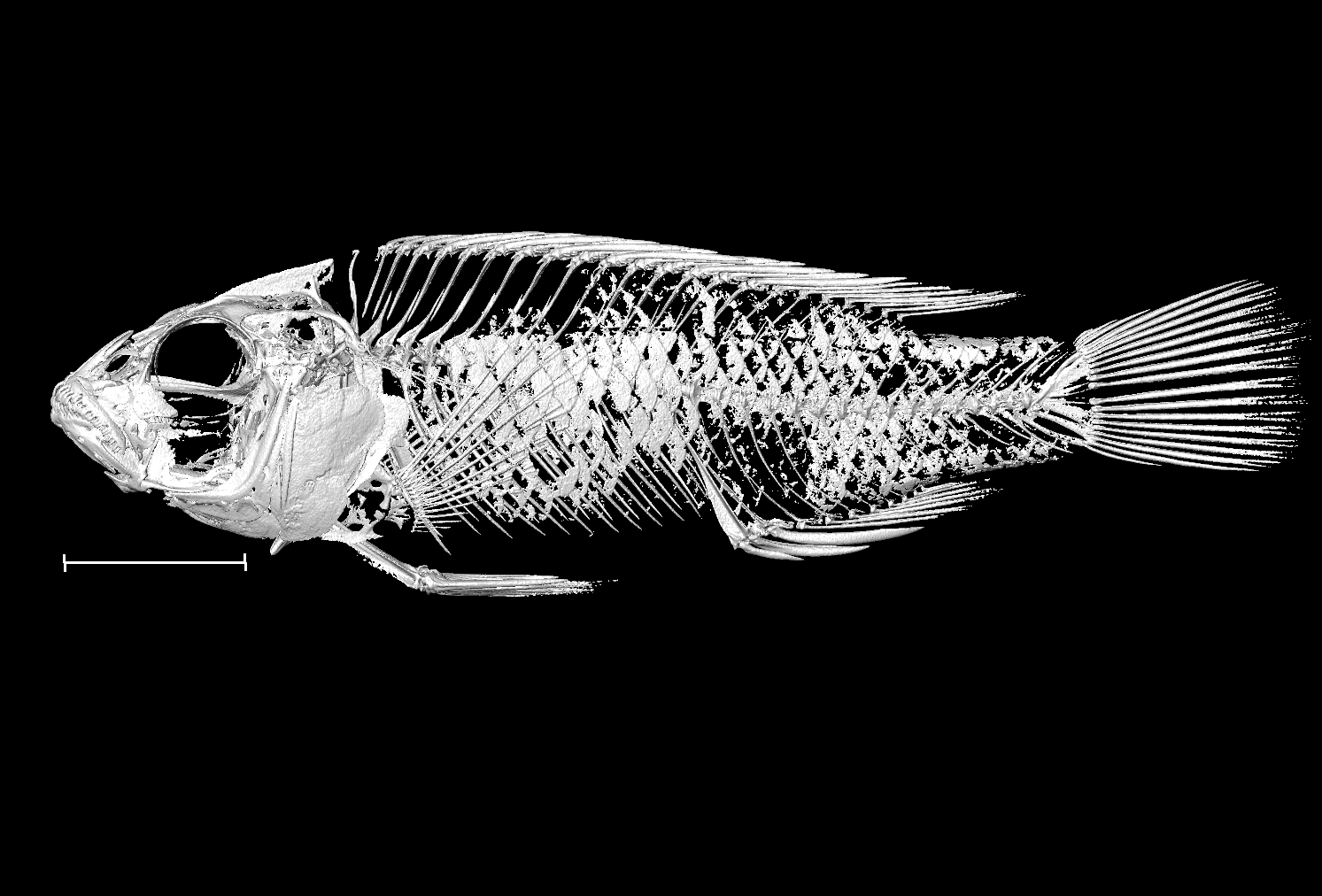

### Astatotilapia_calliptera_UniOxf_AC7_8bit.tif

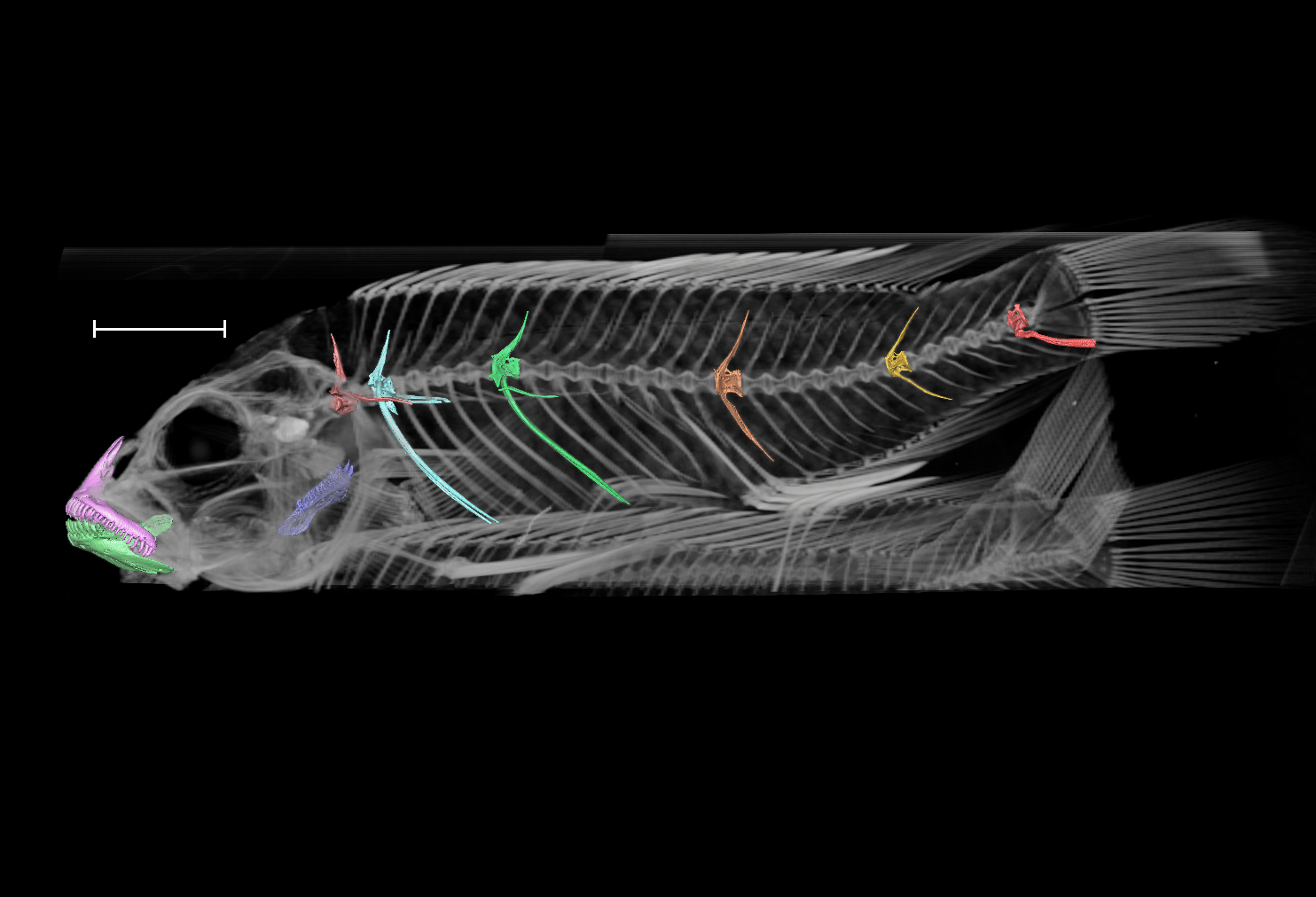

### Astatotilapia_calliptera_UniOxf_AC8_8bit.tif

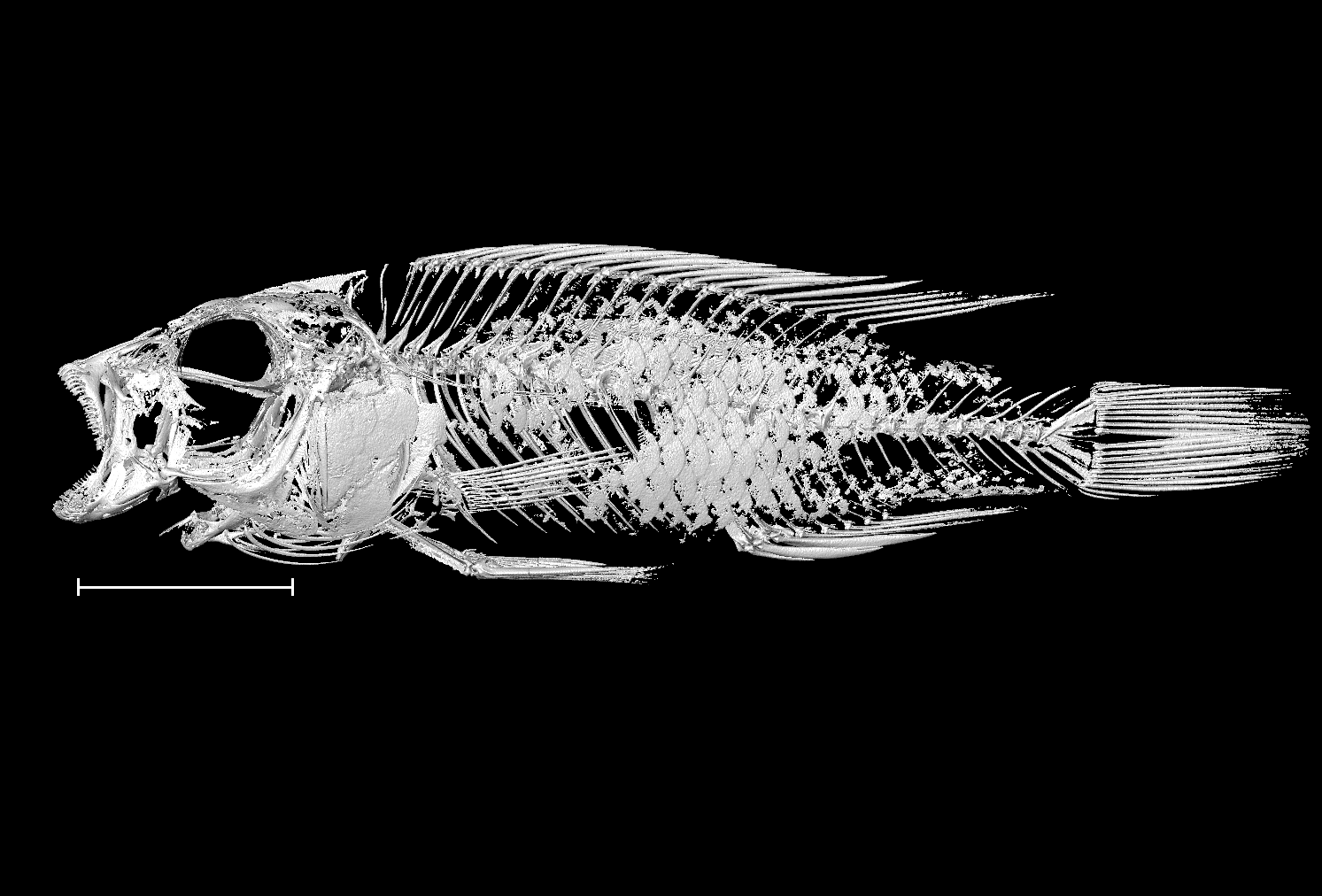

### Astatotilapia_calliptera_UniOxf_AC9_8bit.tif

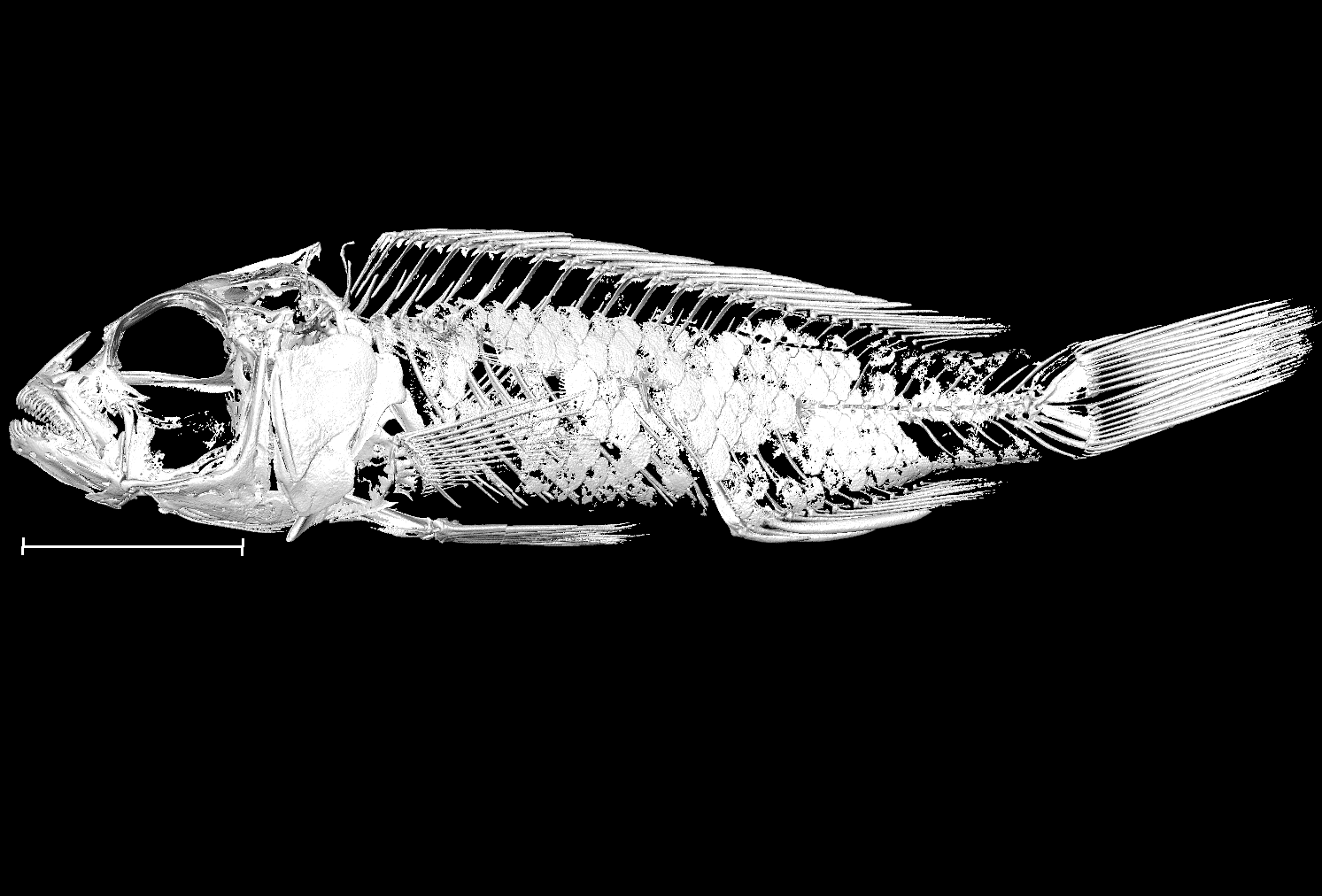

### Astatotilapia_calliptera_UniOxf_H1_8bit.tif

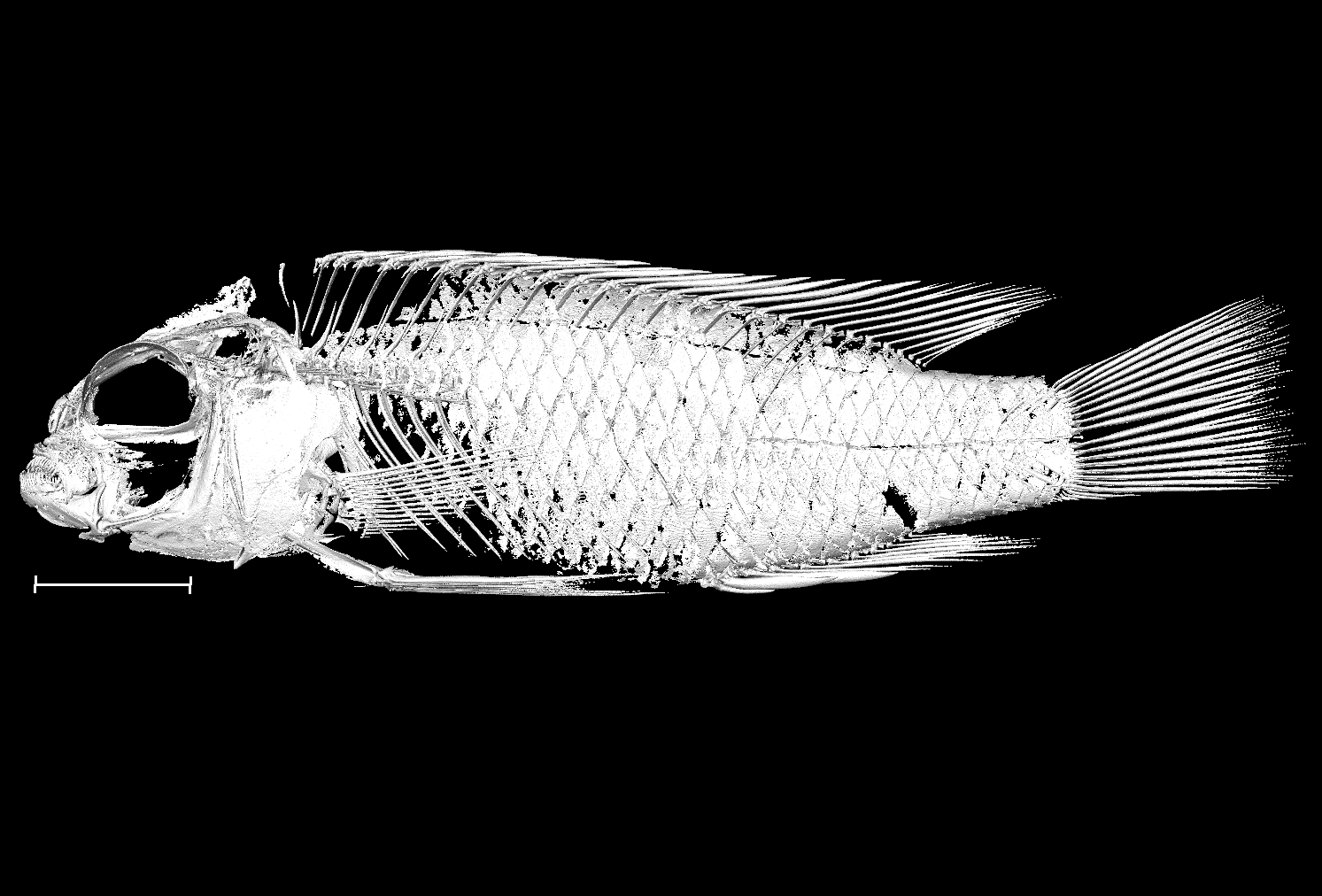

### Astatotilapia_gigliolii_BanUni_G12D10_8bit.tif

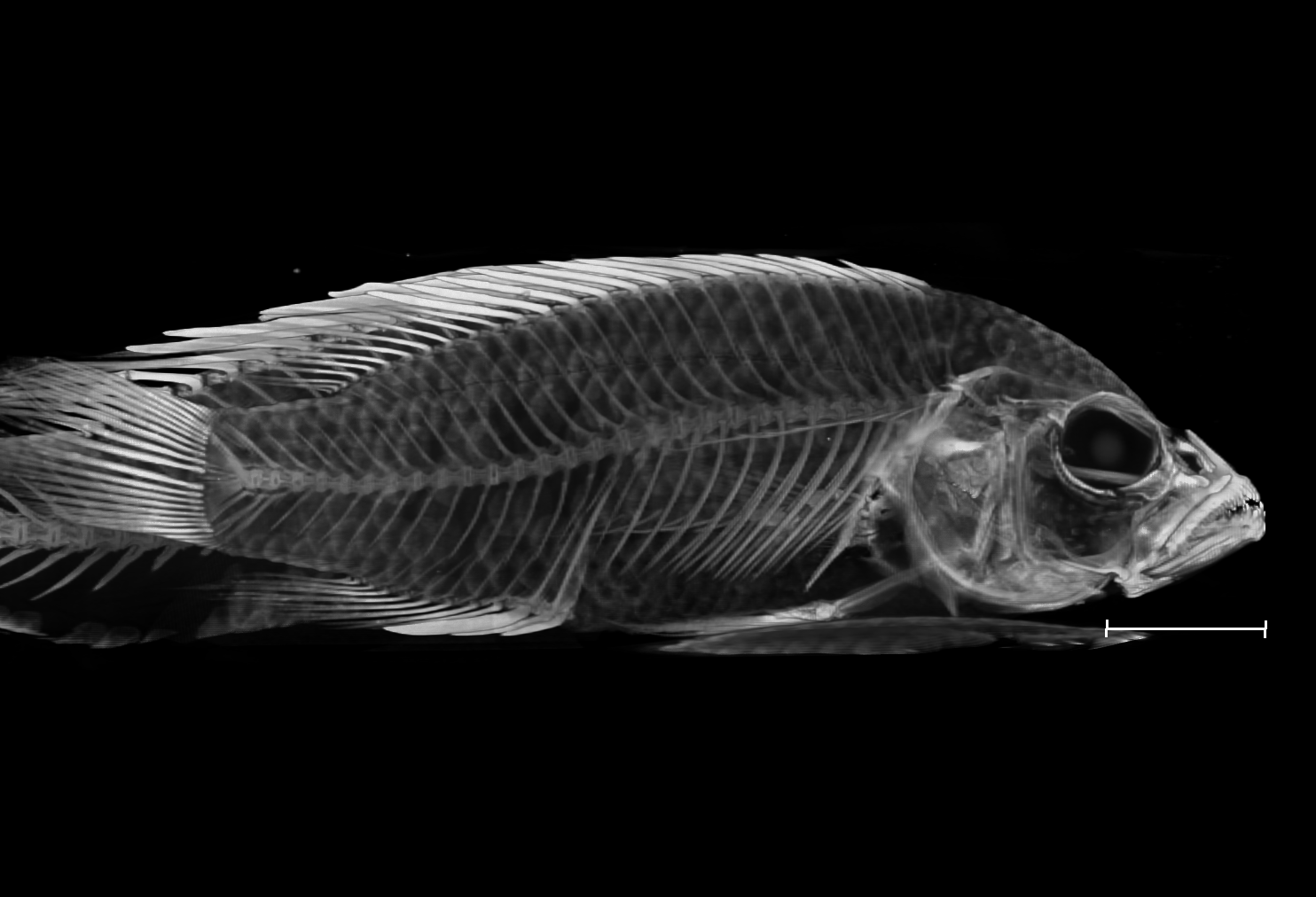

### Astatotilapia_gigliolii_BanUni_G15J08_8bit.tif

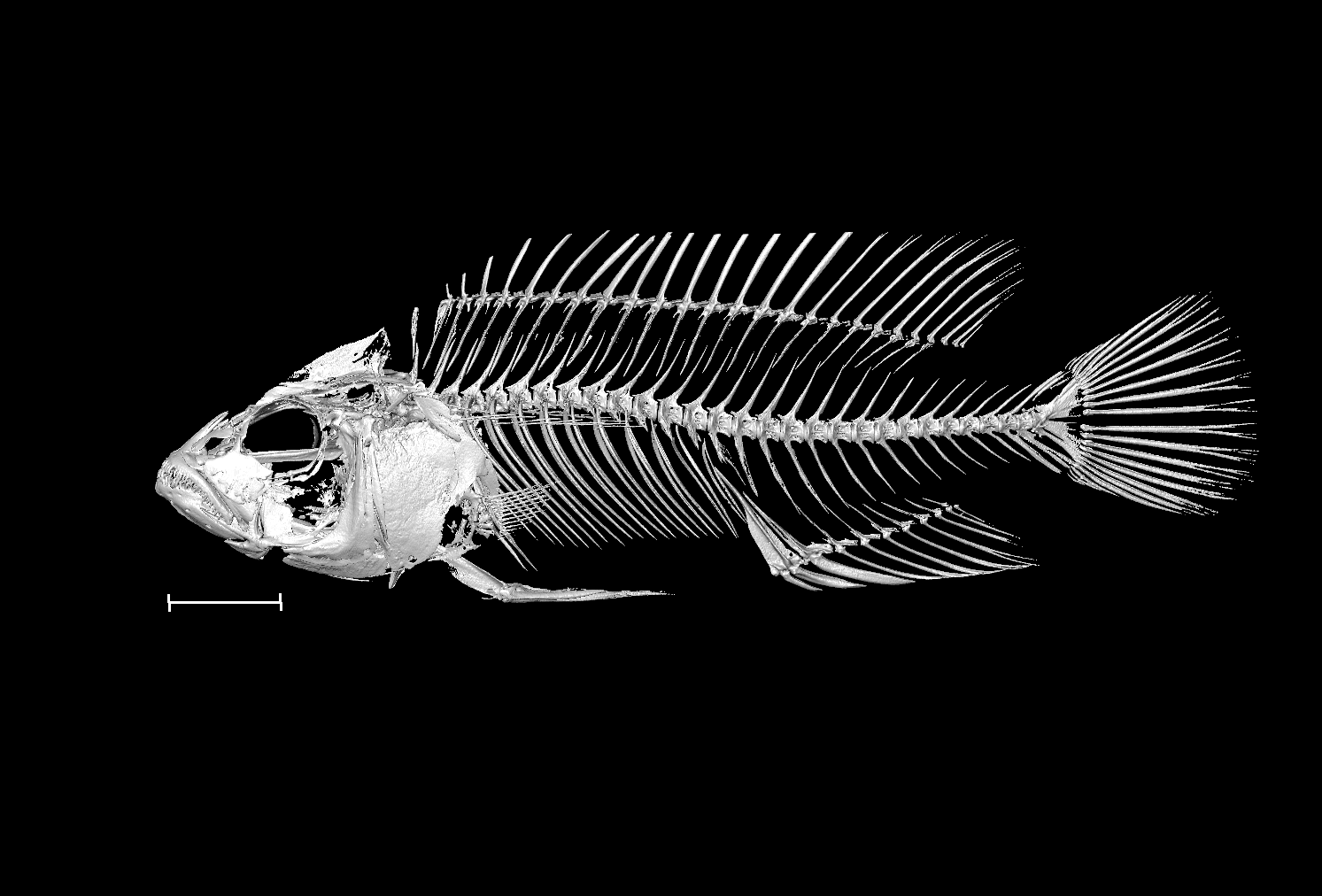

### Astatotilapia_sp_ruaha_blue_BanUni_AU11_8bit.tif

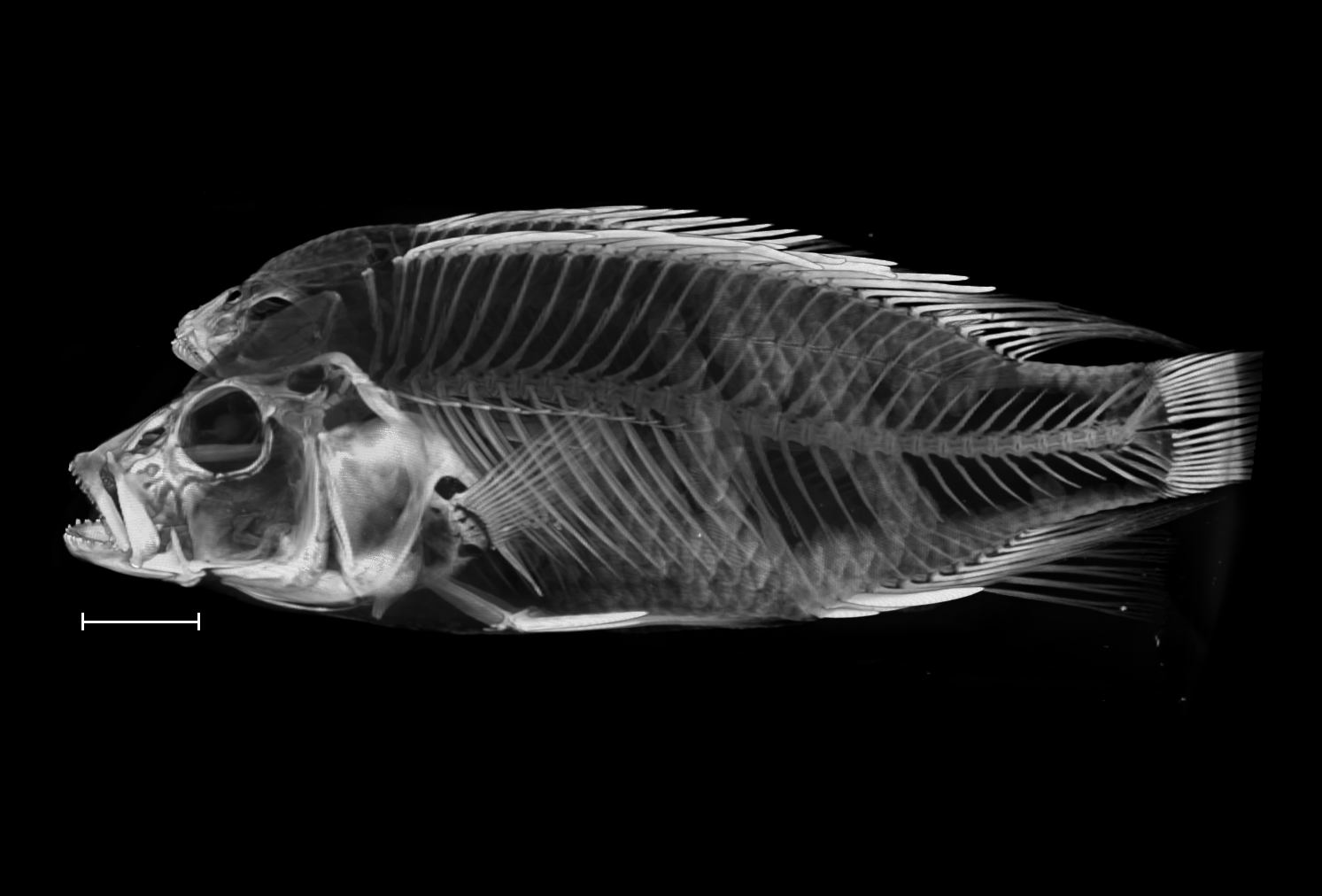

### Astatotilapia_sp_ruaha_blue_BanUni_AU13_8bit.tif

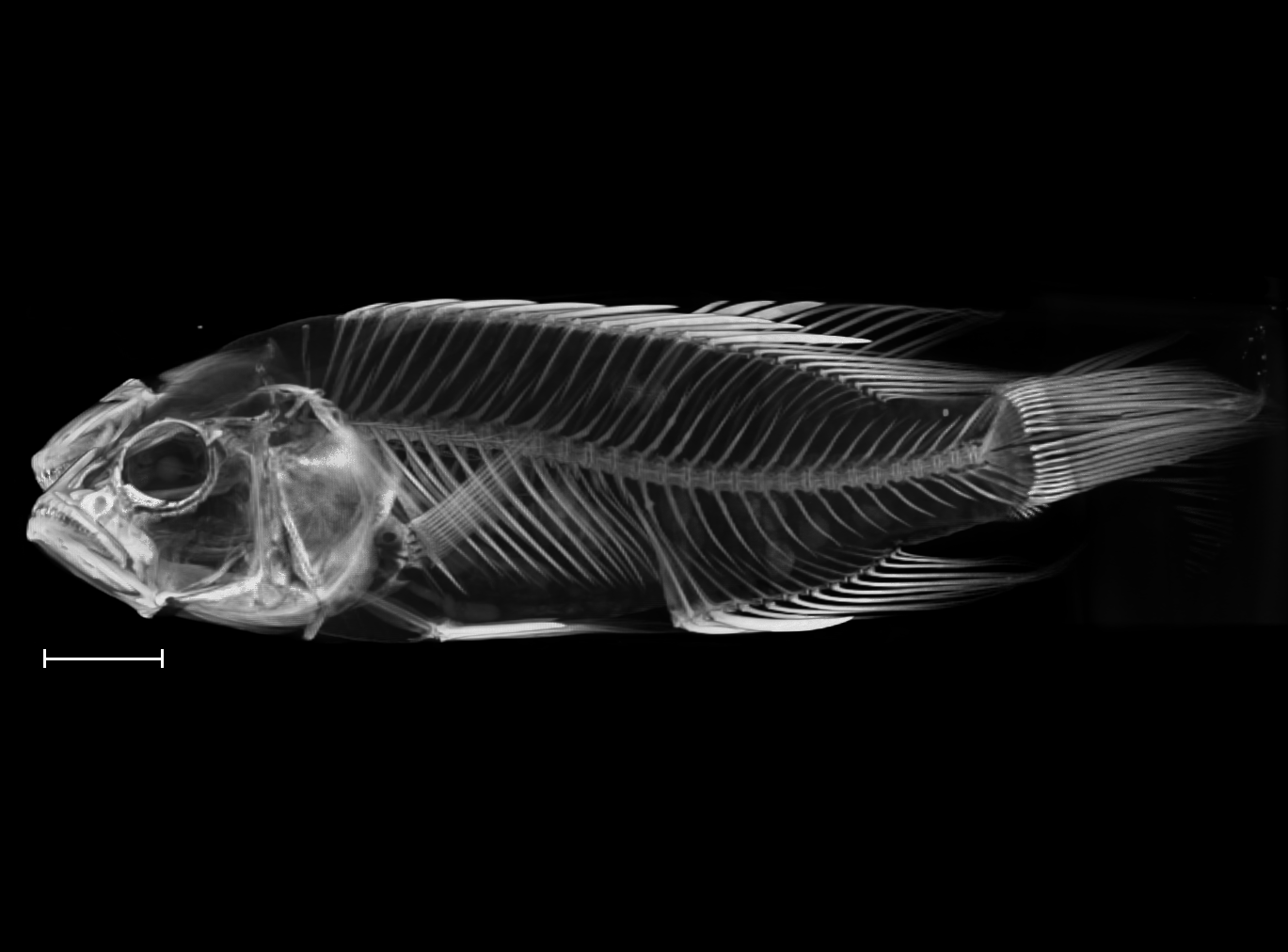

### Copadichromis_likomae_NHMUK_1962_10_18_113_118_8bit_a.tif

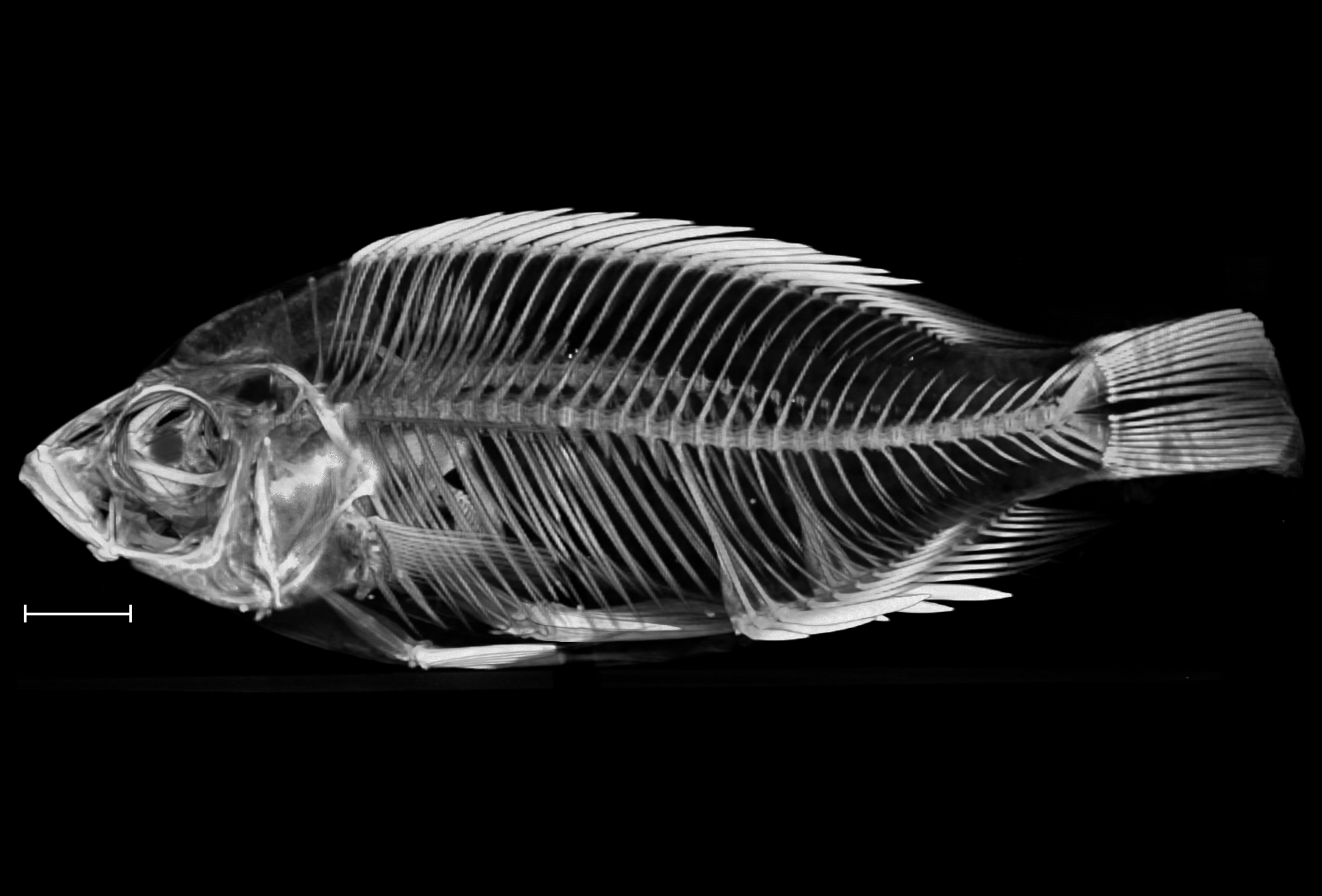

### Copadichromis_likomae_NHMUK_1962_10_18_113_118_8bit_b.tif

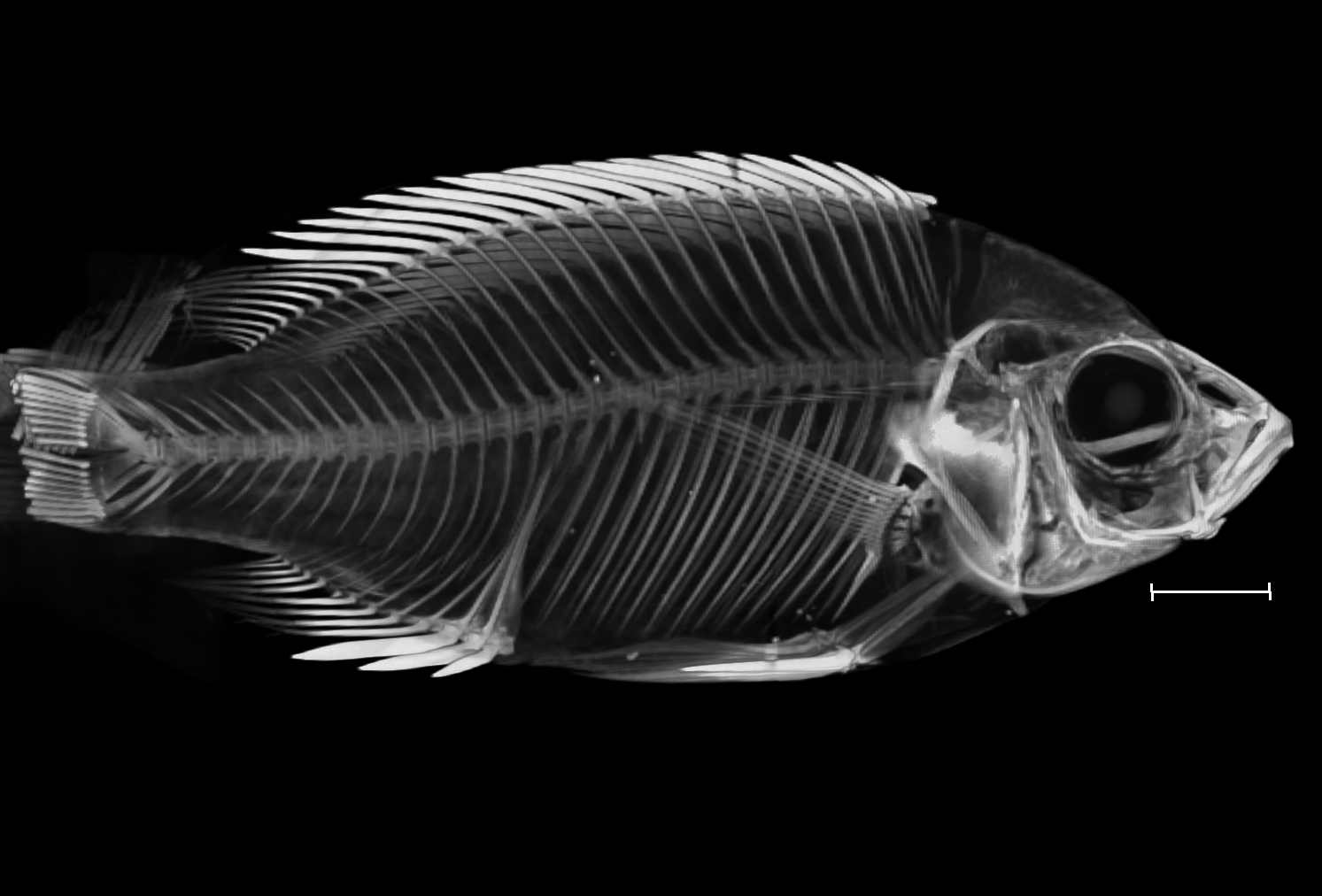

### Copadichromis_quadrimaculatus_NHMUK_1908_10_27_42_48_8bit_a.tif

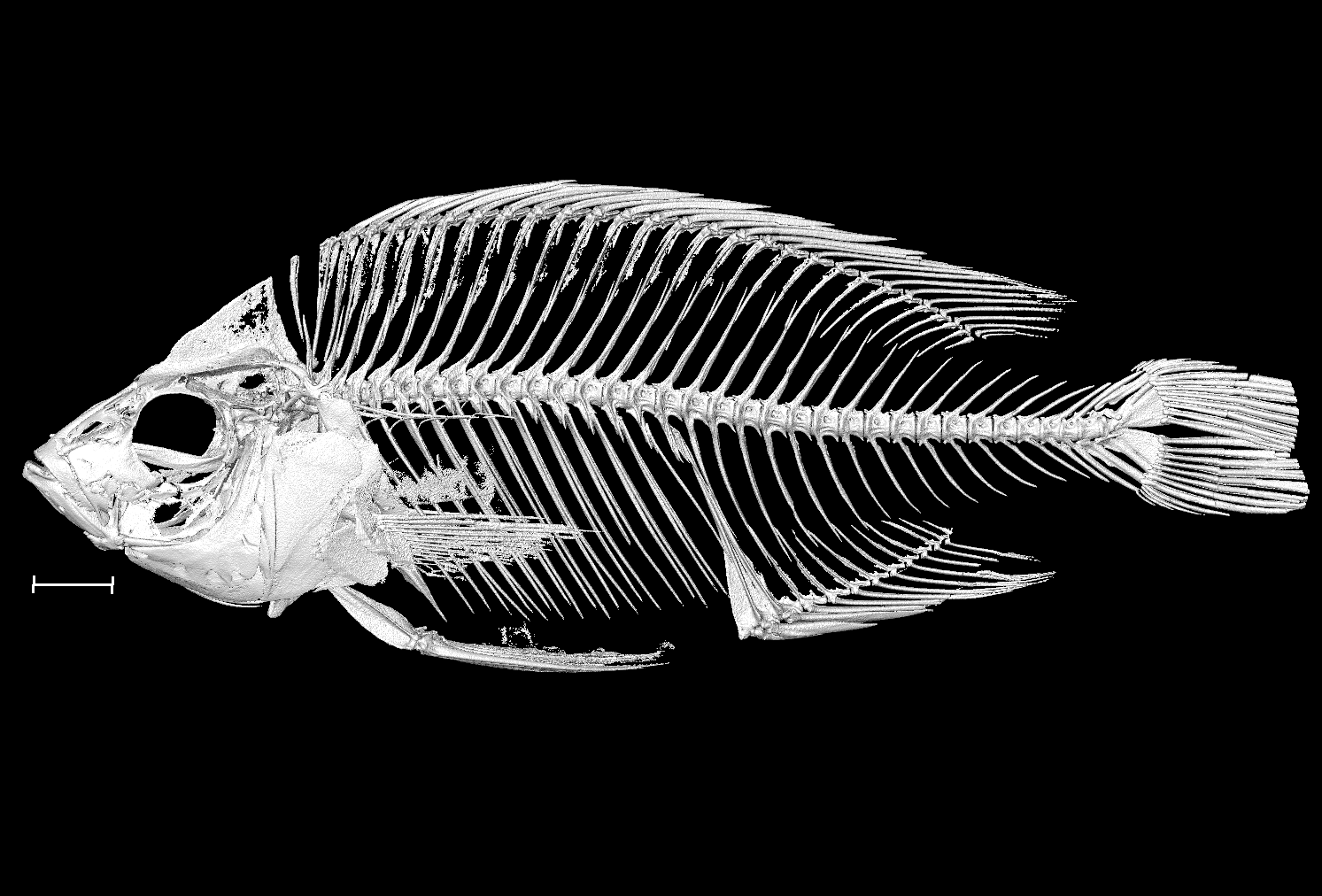

### Copadichromis_quadrimaculatus_NHMUK_1908_10_27_42_48_8bit_b.tif

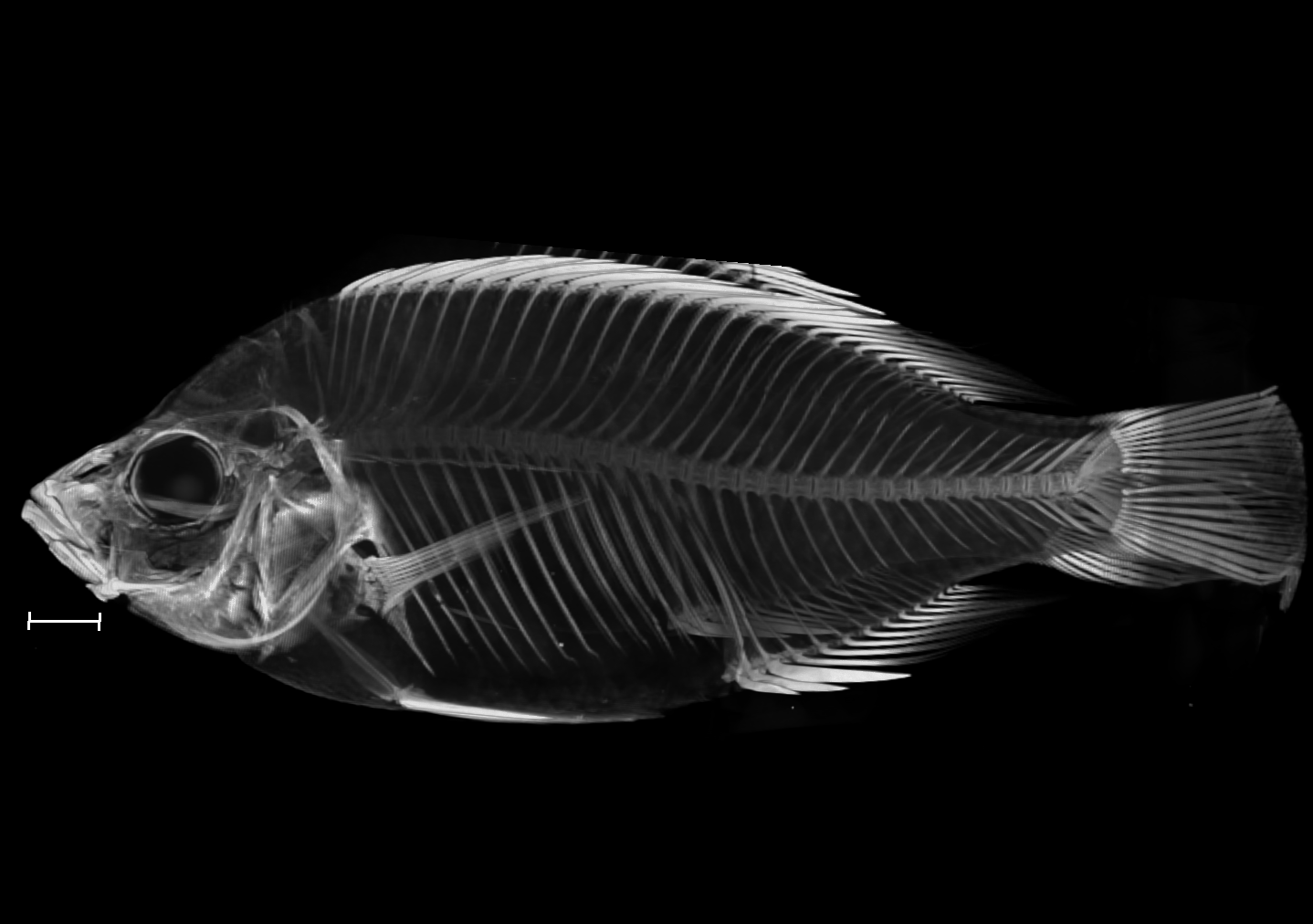

### Copadichromis_trimaculatus_NHMUK_1962_10_18_129_136_8bit_a.tif

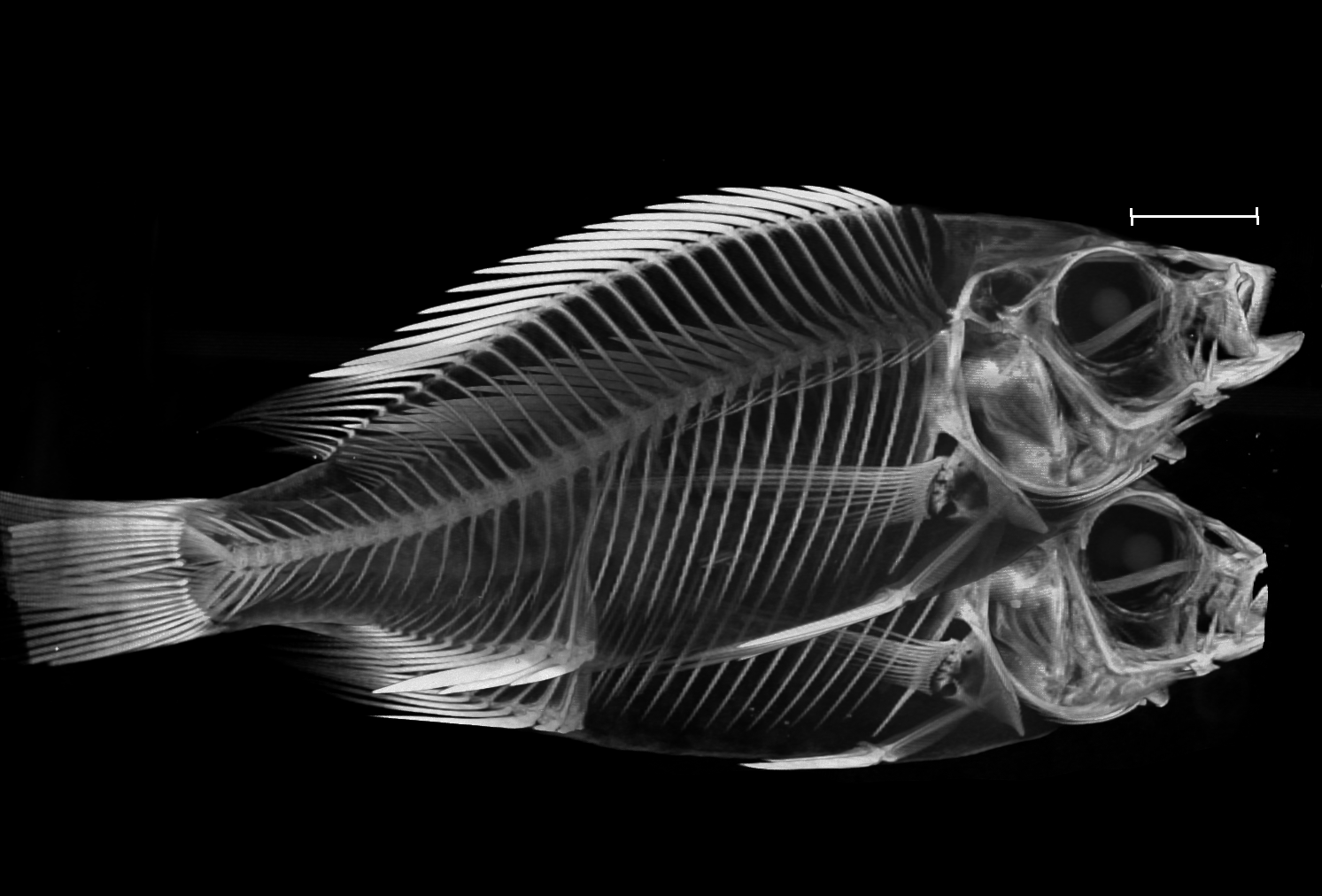

### Copadichromis_trimaculatus_NHMUK_1962_10_18_129_136_8bit_b.tif

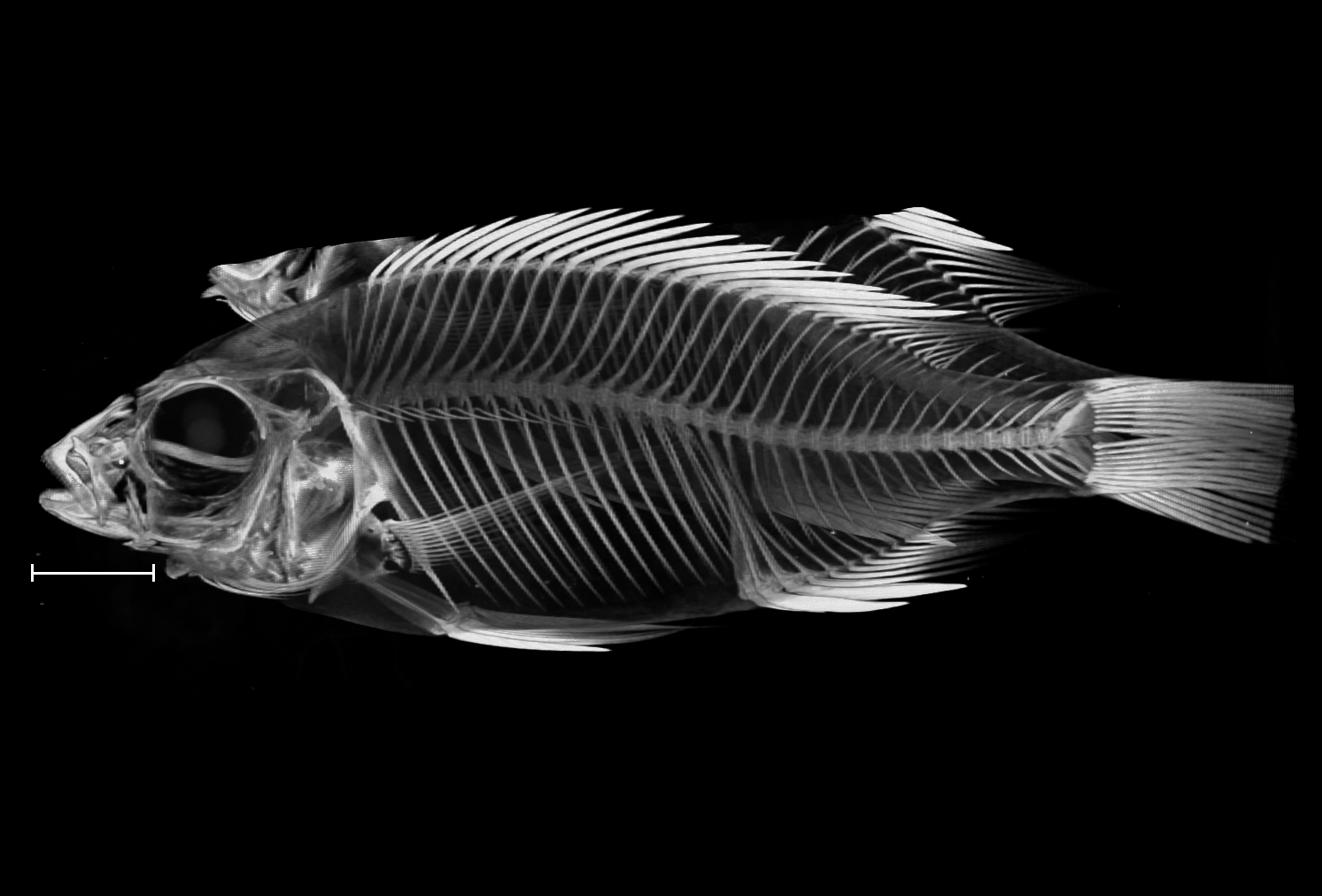

### Copadichromis_virginalis_NHMUK_1962_10_18_32_39_8bit_a.tif

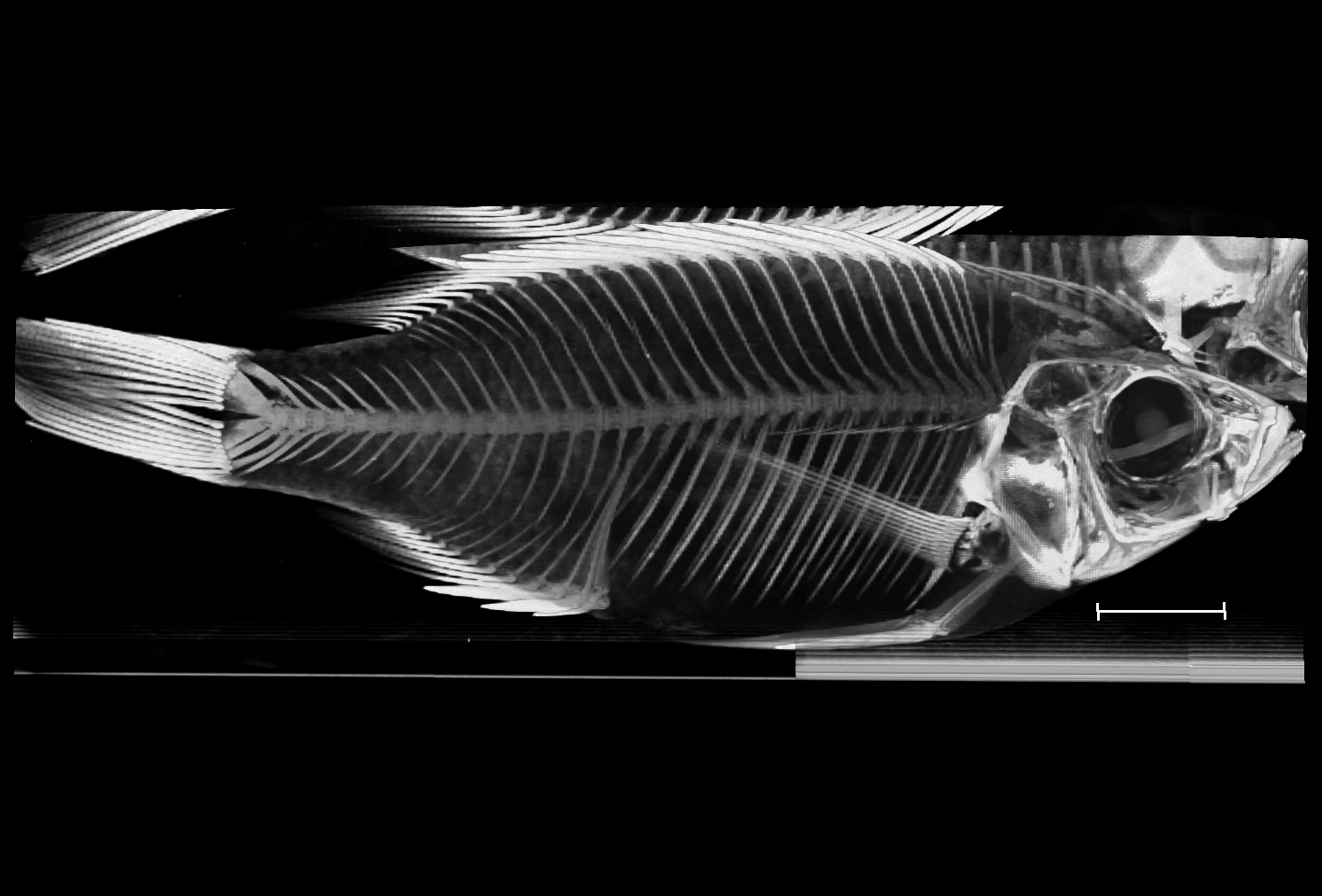

### Copadichromis_virginalis_NHMUK_1962_10_18_32_39_8bit_b.tif

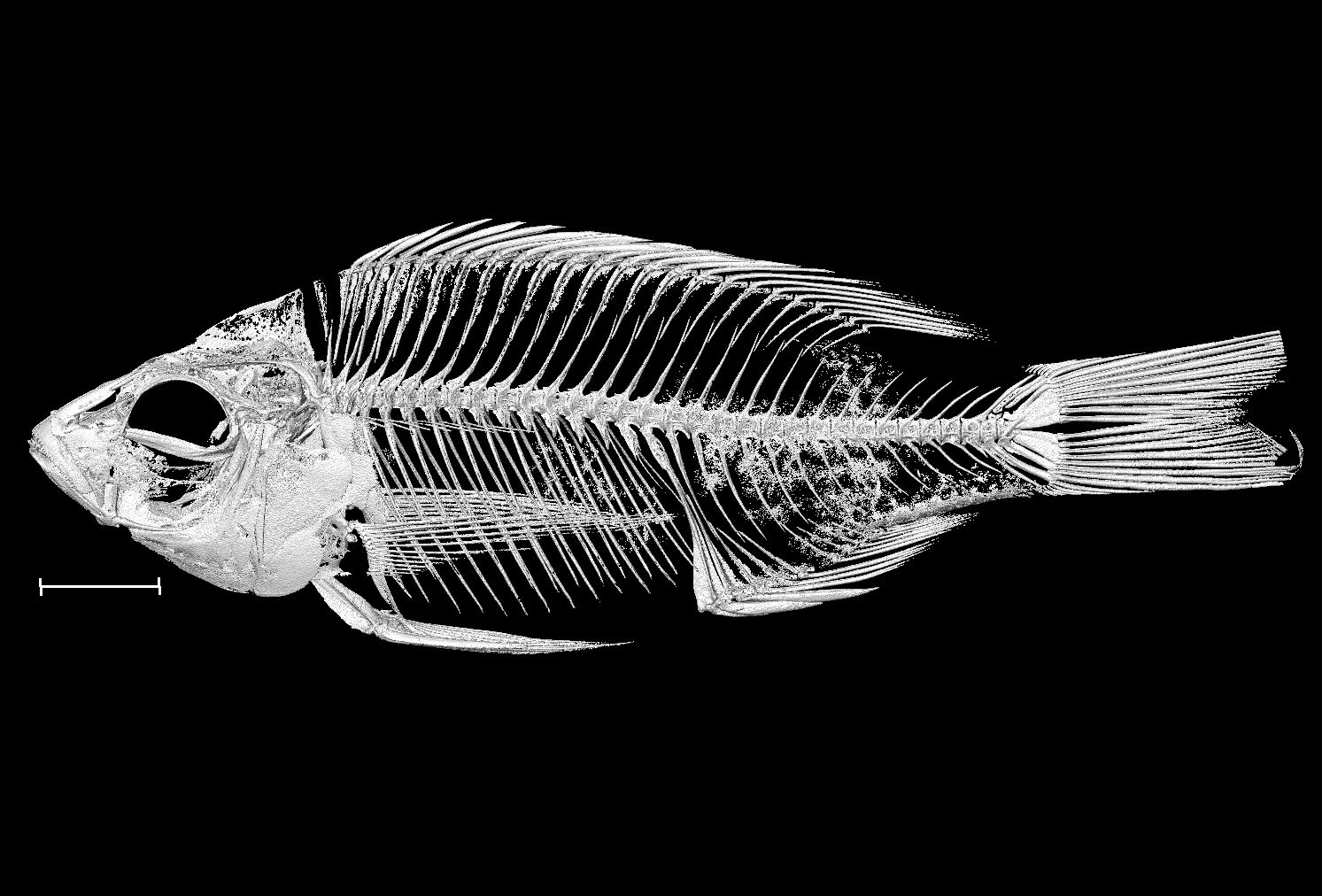

### Cynotilapia_axelrodi_NHMUK_1976_11_19_1_8bit.tif

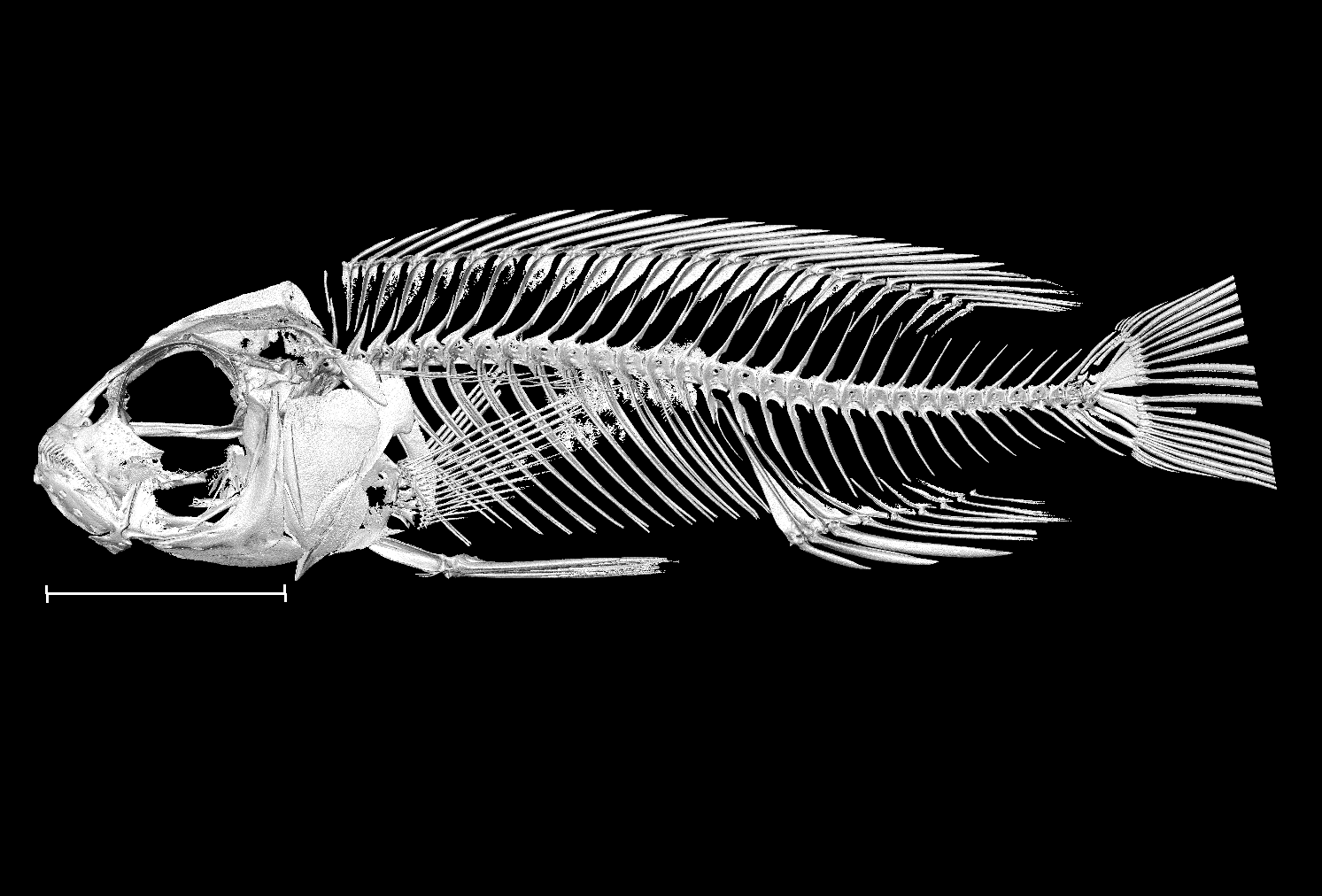

### Dimidiochromis_compressiceps_UniBri_213_8bit.tif

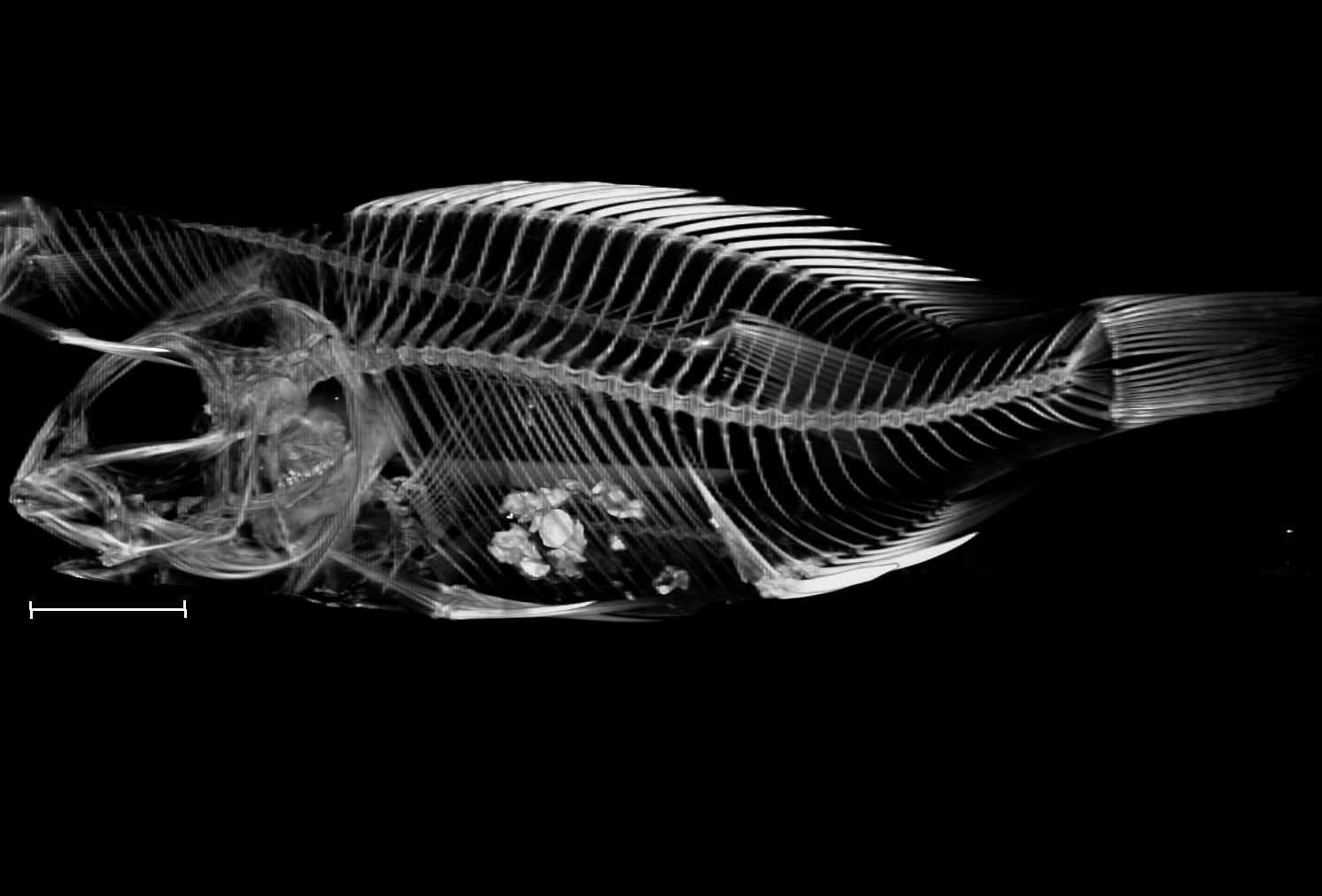

### Dimidiochromis_strigatus_UniBri_39_8bit.tif

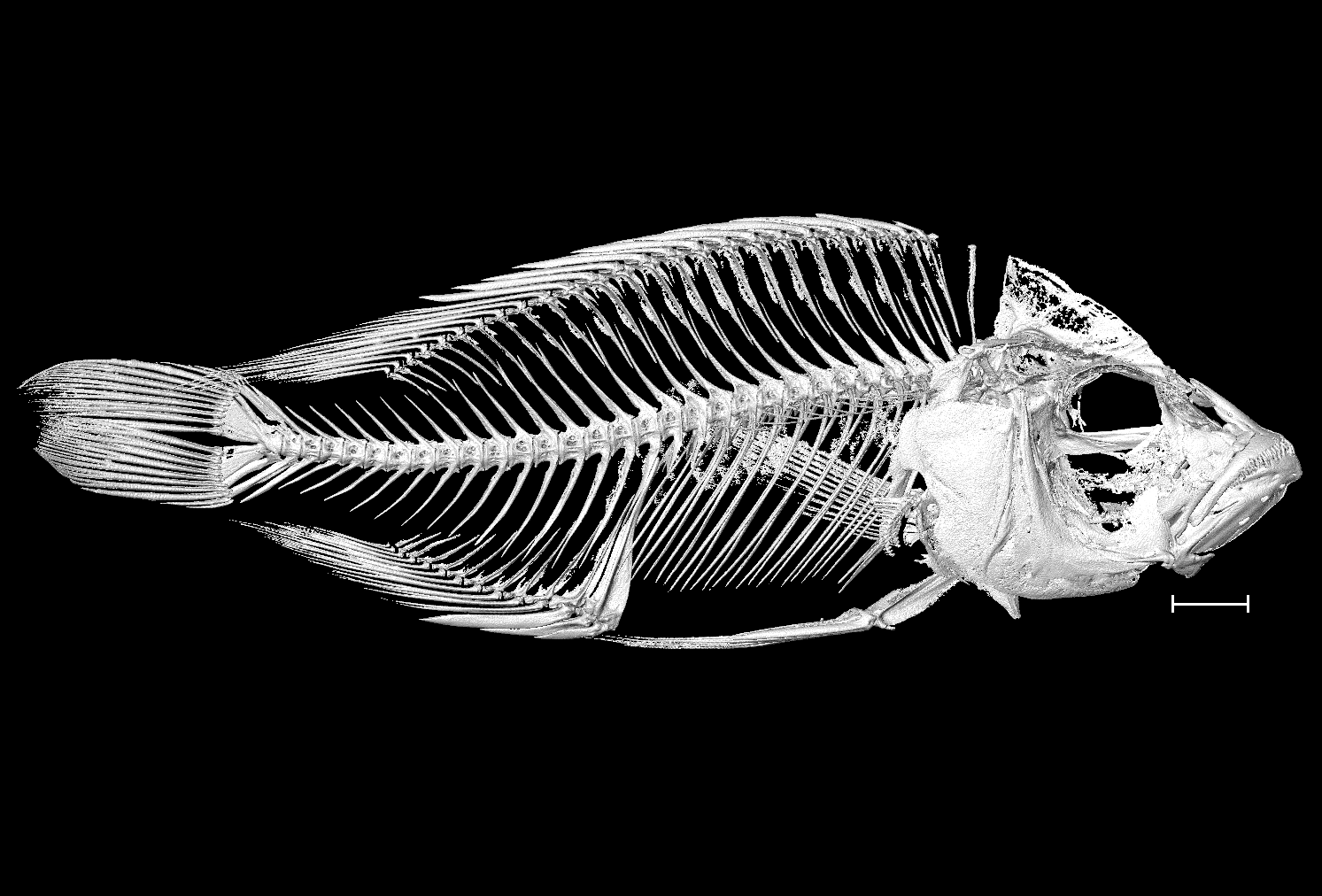

### Diplotaxodon_aeneus_NHMUK_1996_4_30_17_20_8bit_a.tif

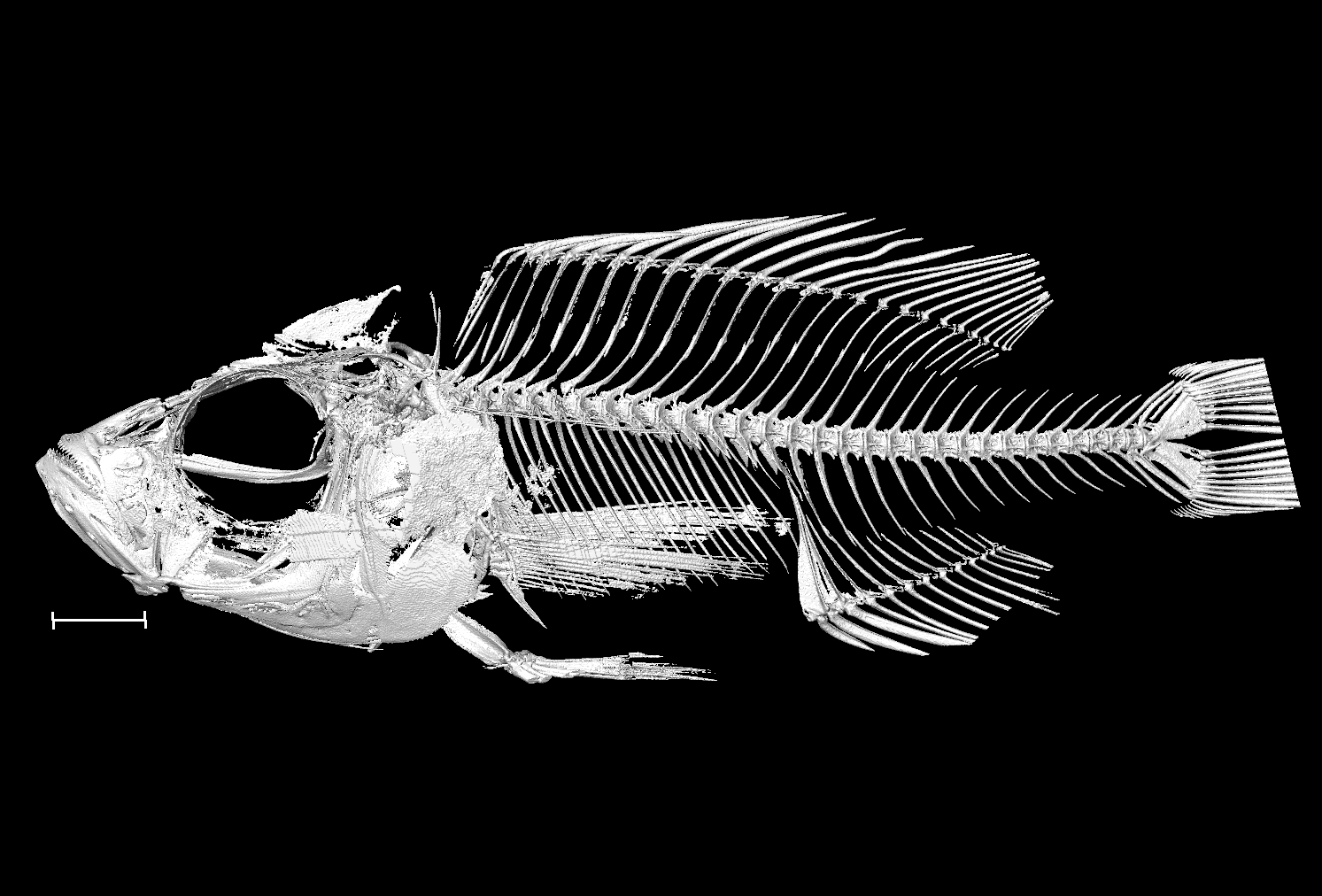

### Diplotaxodon_aeneus_NHMUK_1996_4_30_17_20_8bit_b.tif

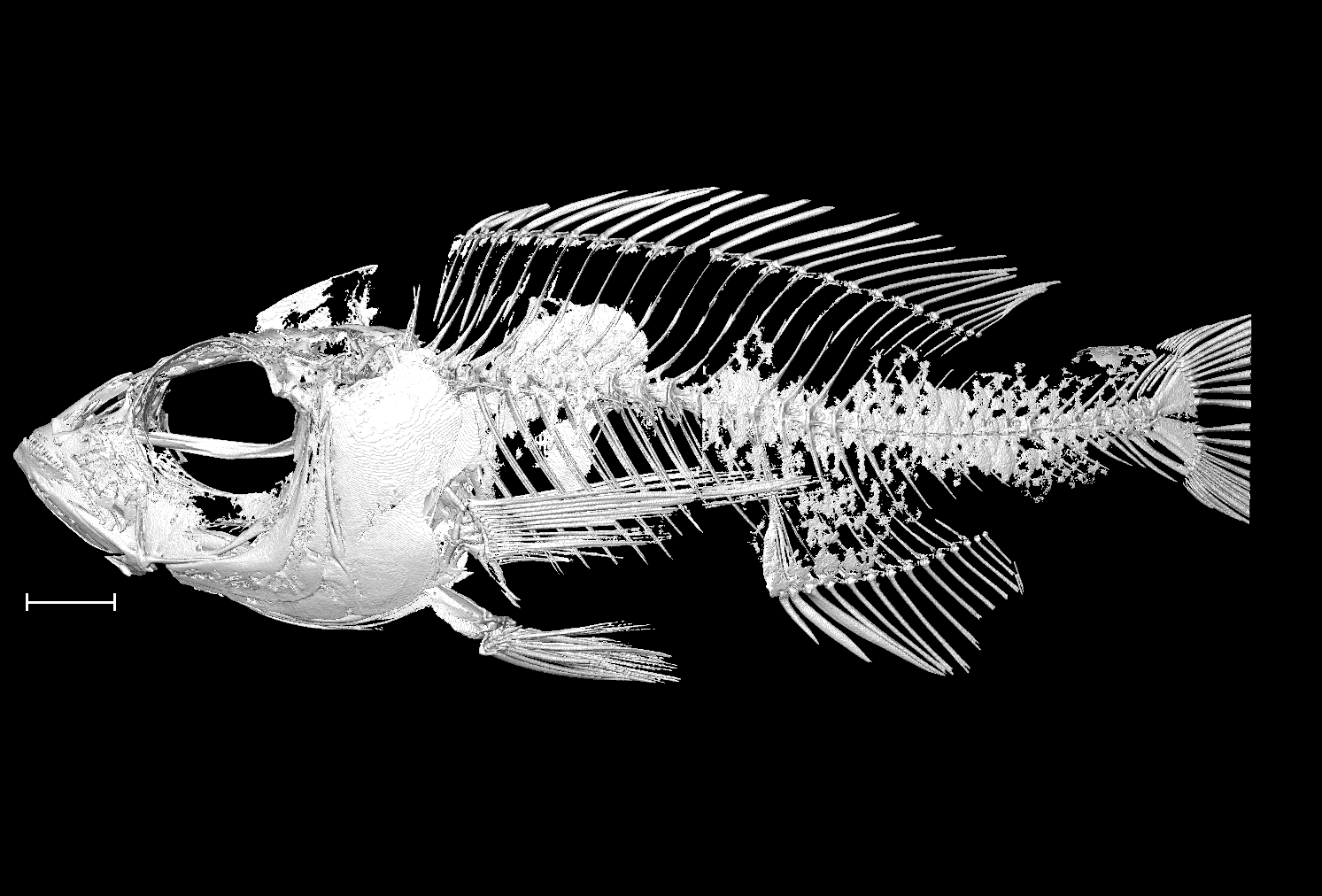

### Diplotaxodon_aeneus_UniBri_DRC051_8bit_a.tif

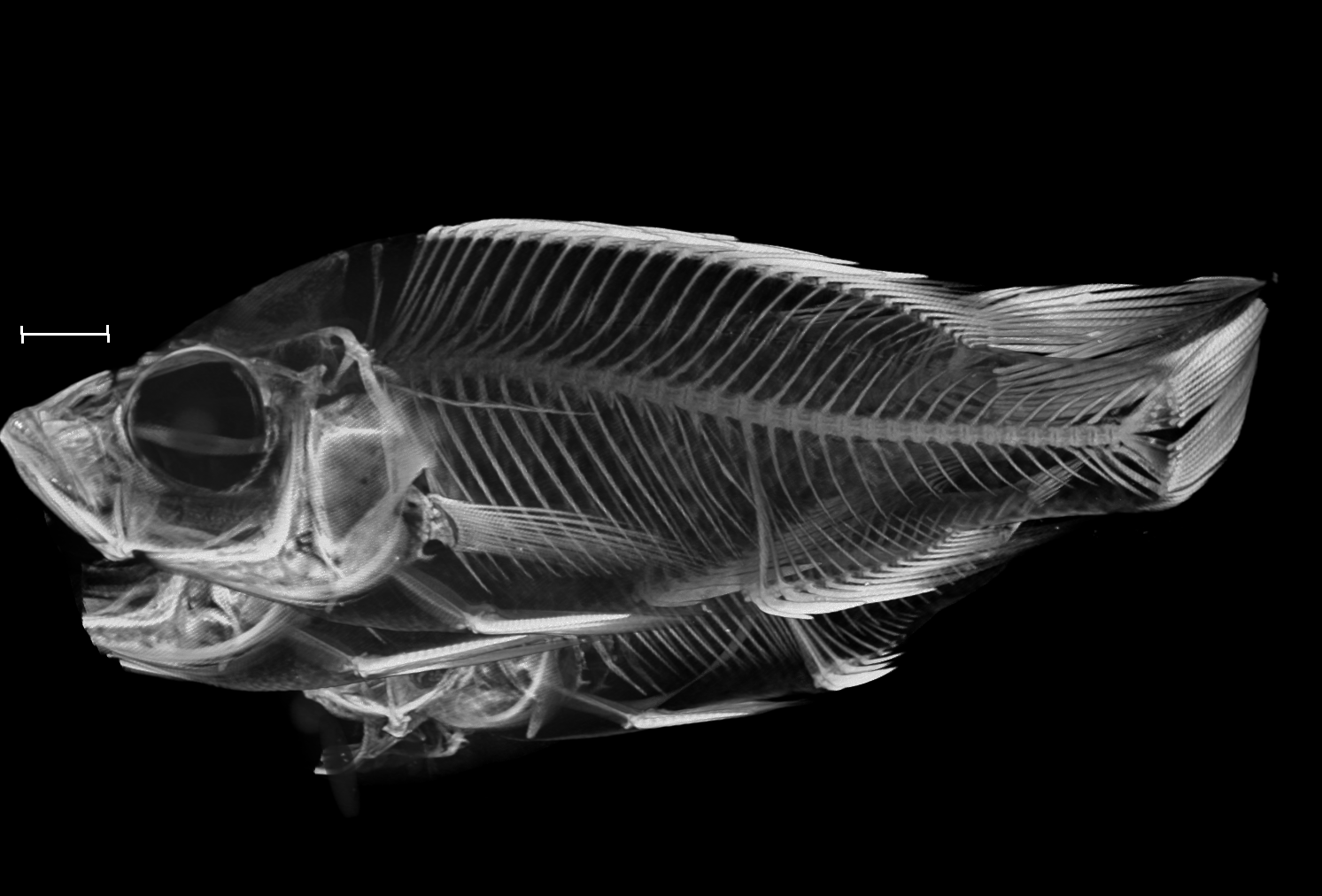

### Diplotaxodon_aeneus_UniBri_DRC051_8bit_b.tif

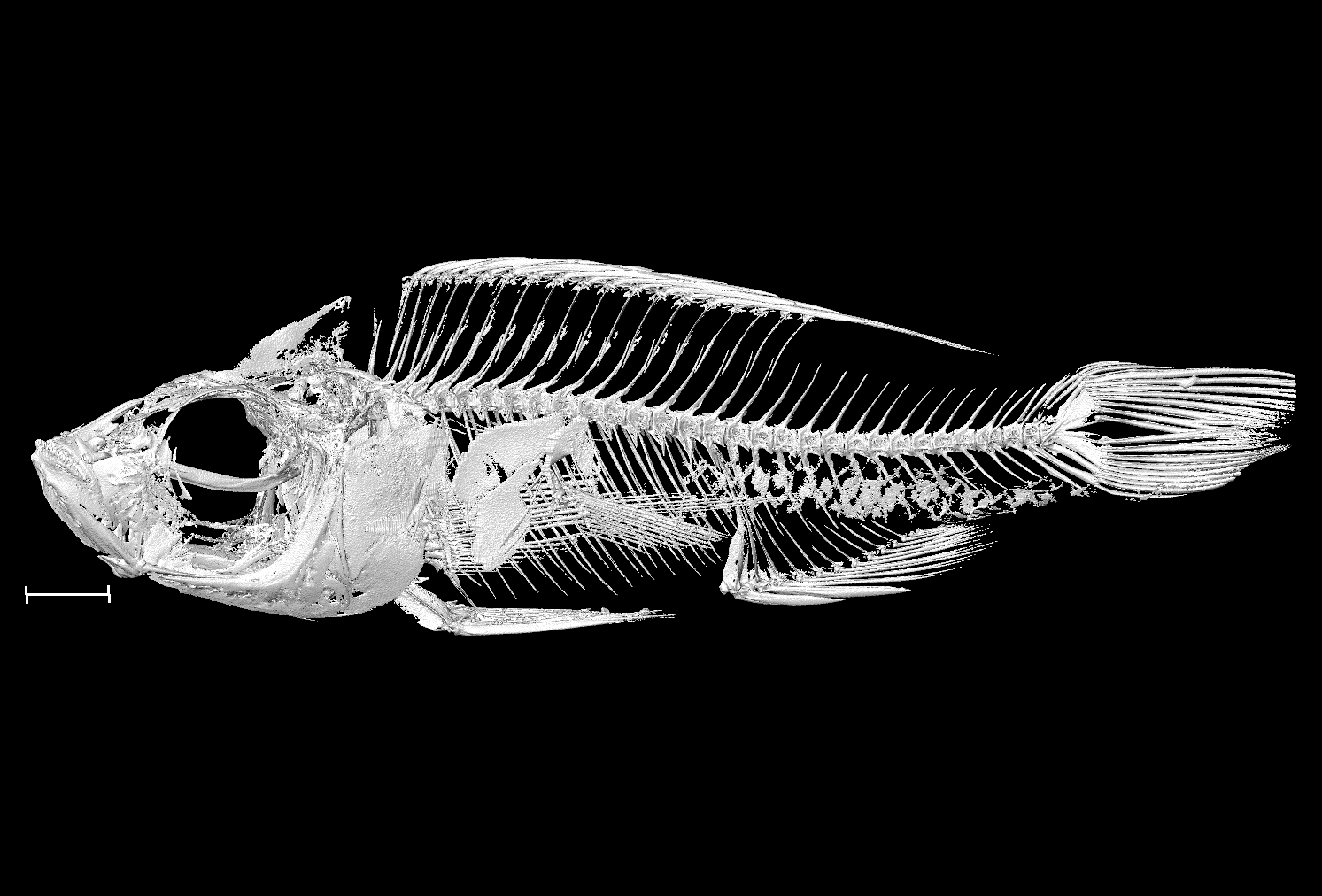

### Diplotaxodon_argenteus_NHMUK_1935_6_14_2281_2282_8bit.tif

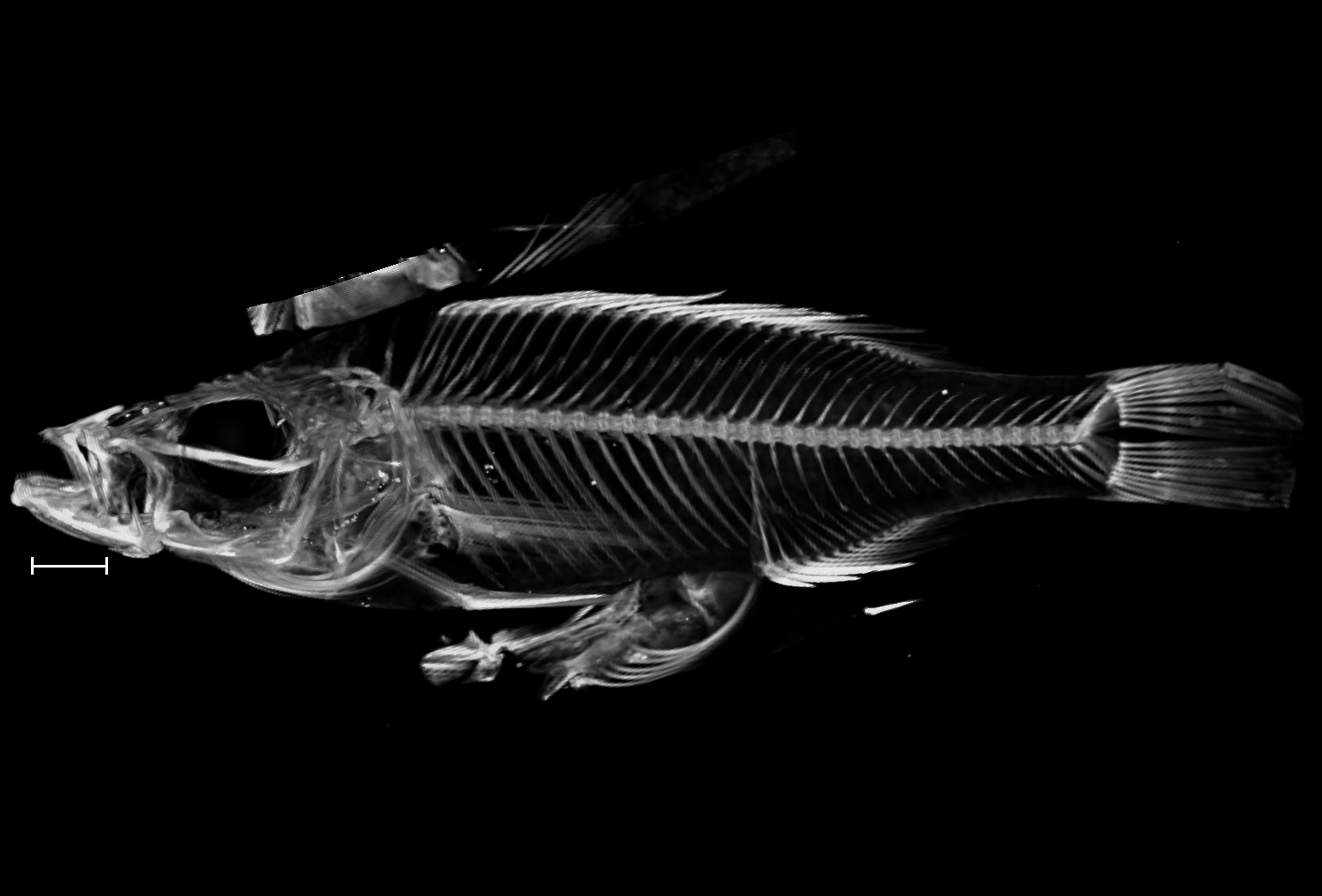

### Diplotaxodon_limnothrissa_NHMUK_1992_3_25_2_17_8bit_a.tif

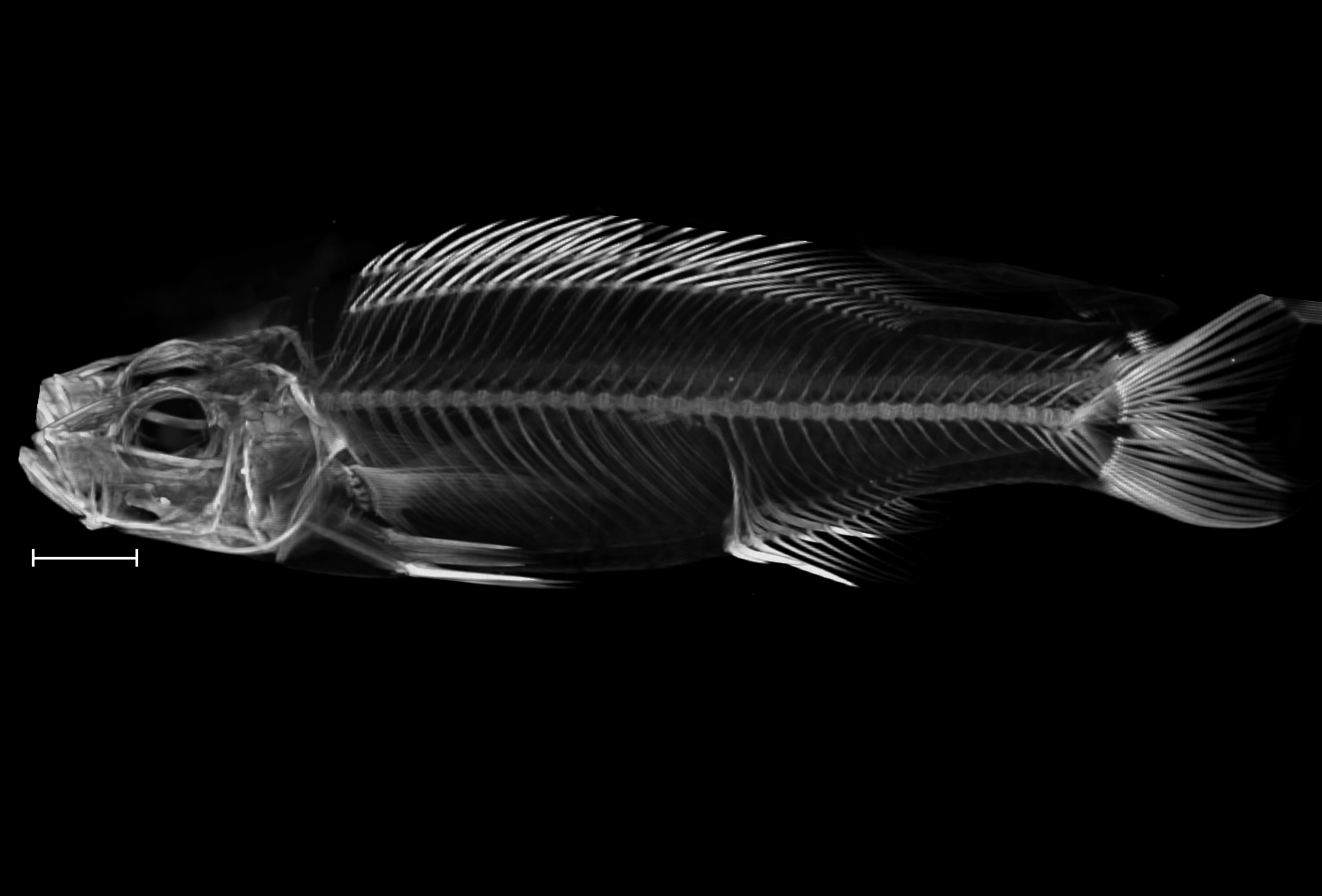

### Diplotaxodon_limnothrissa_NHMUK_1992_3_25_2_17_8bit_b.tif

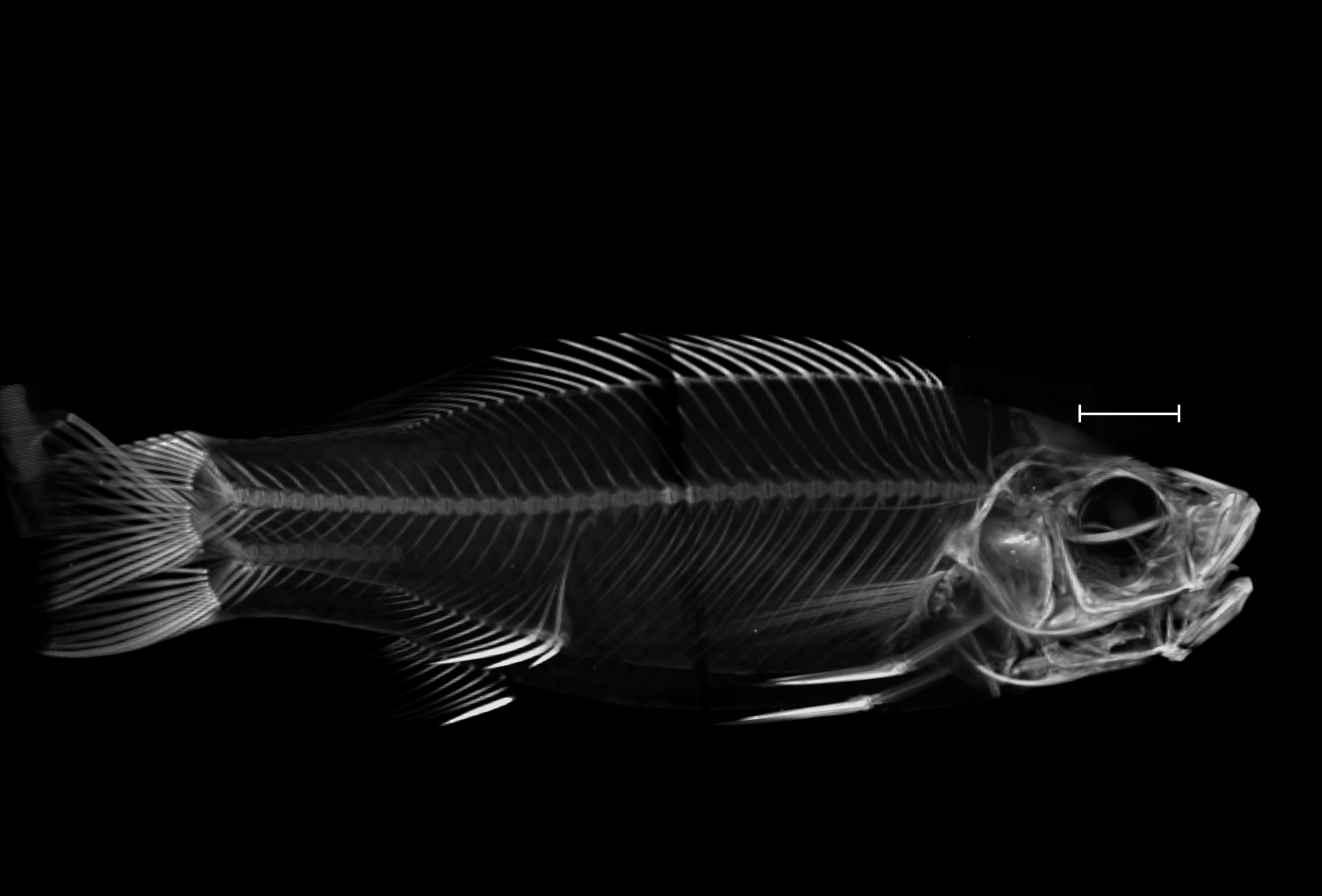

### Diplotaxodon_macrops_NHMUK_1996_4_30_2_10_3_8bit.tif

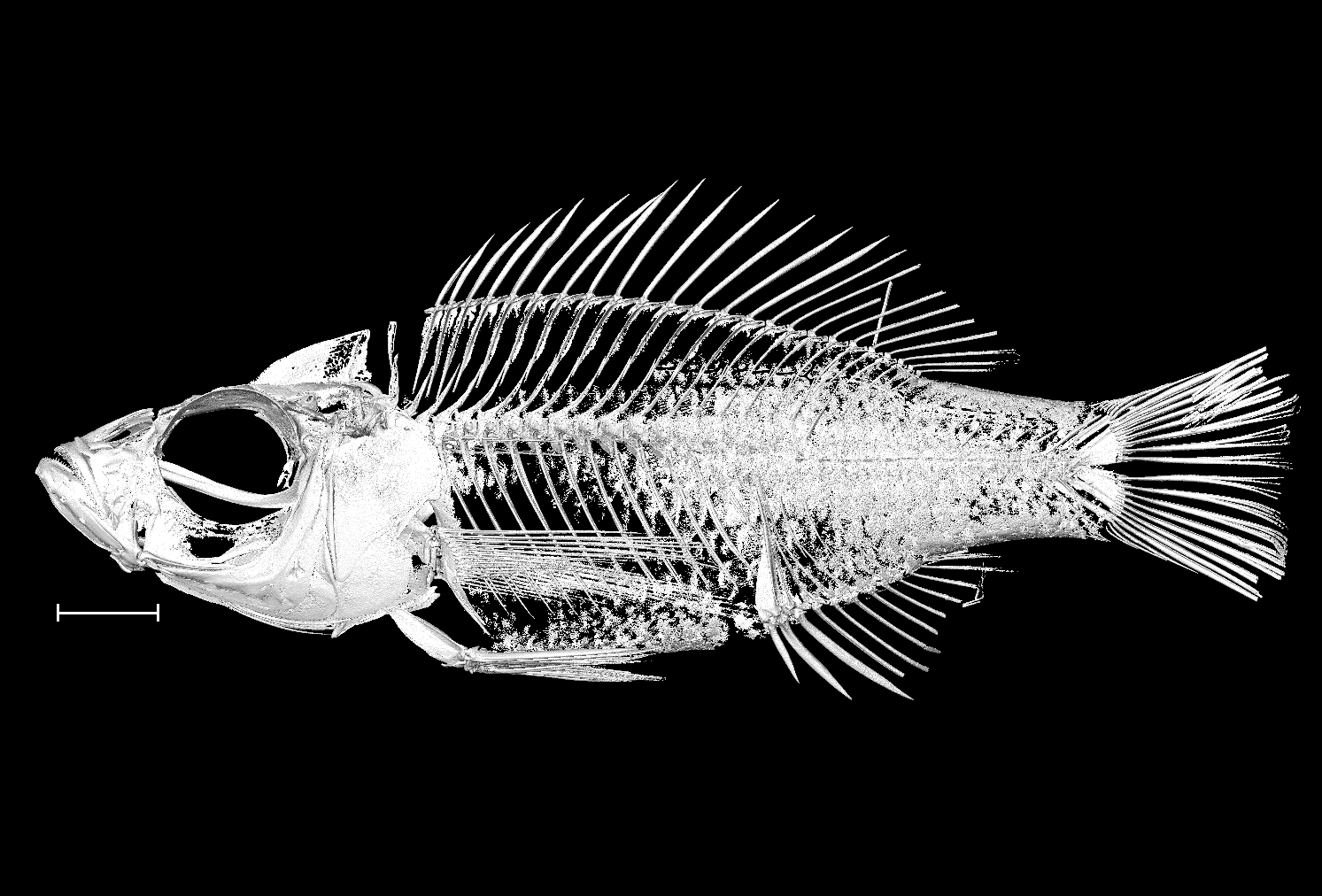

### Diplotaxodon_sp_holochromis_UniBri_DRC001_8bit_a.tif

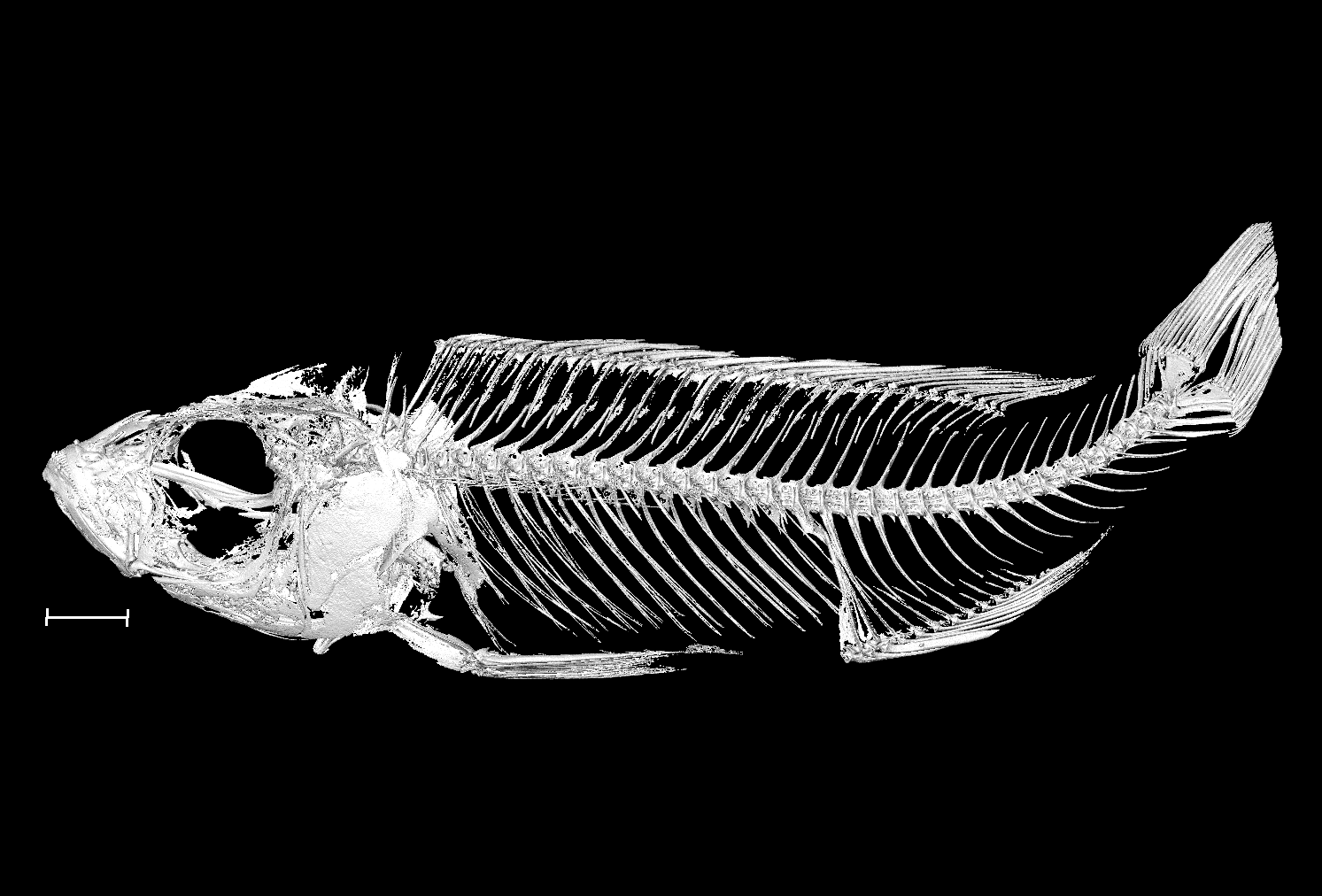

### Diplotaxodon_sp_holochromis_UniBri_DRC001_8bit_b.tif

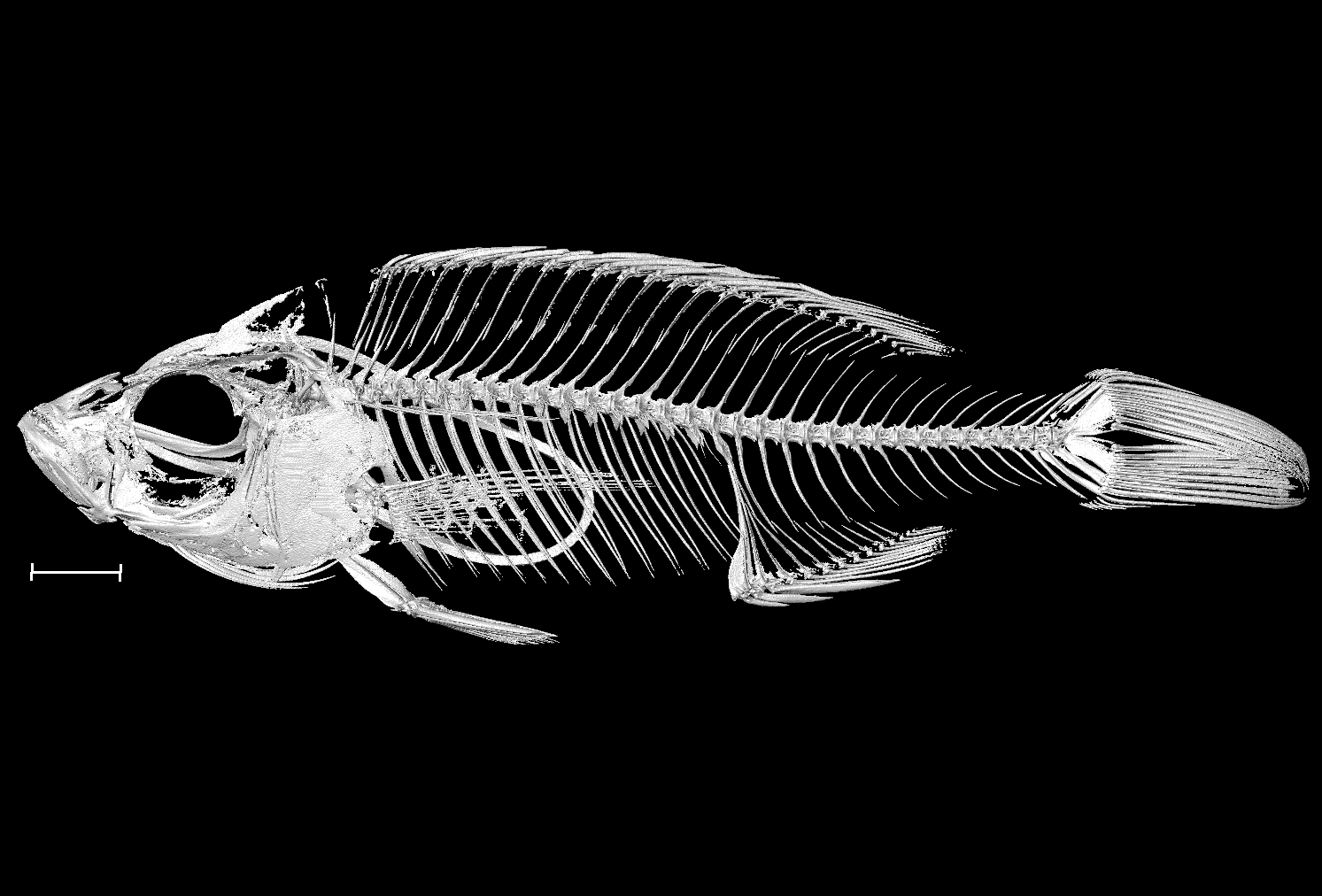

### Diplotaxodon_sp_macrops_black_dorsal_UniBri_DRC090_8bit_a.tif

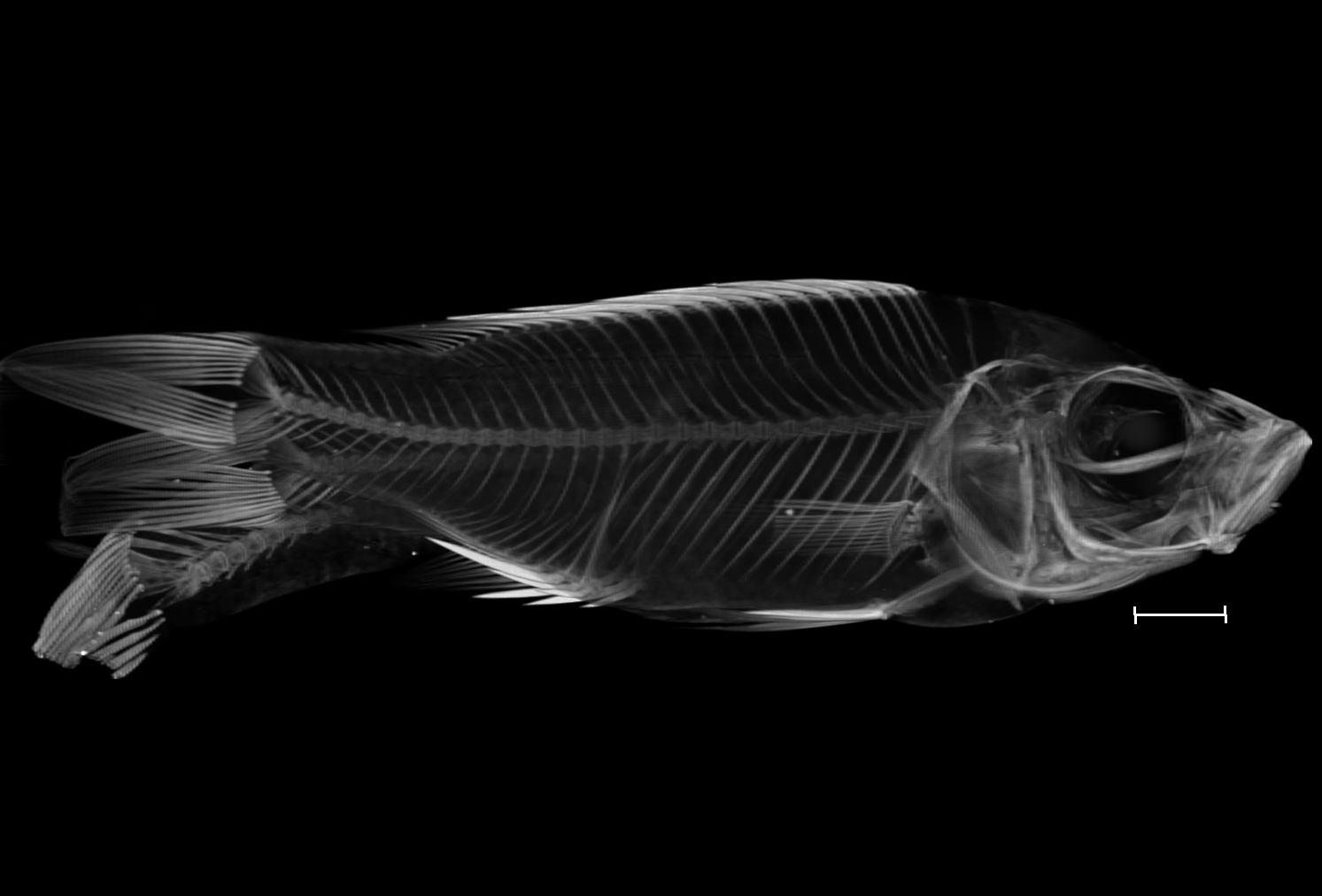

### Diplotaxodon_sp_macrops_black_dorsal_UniBri_DRC090_8bit_b.tif

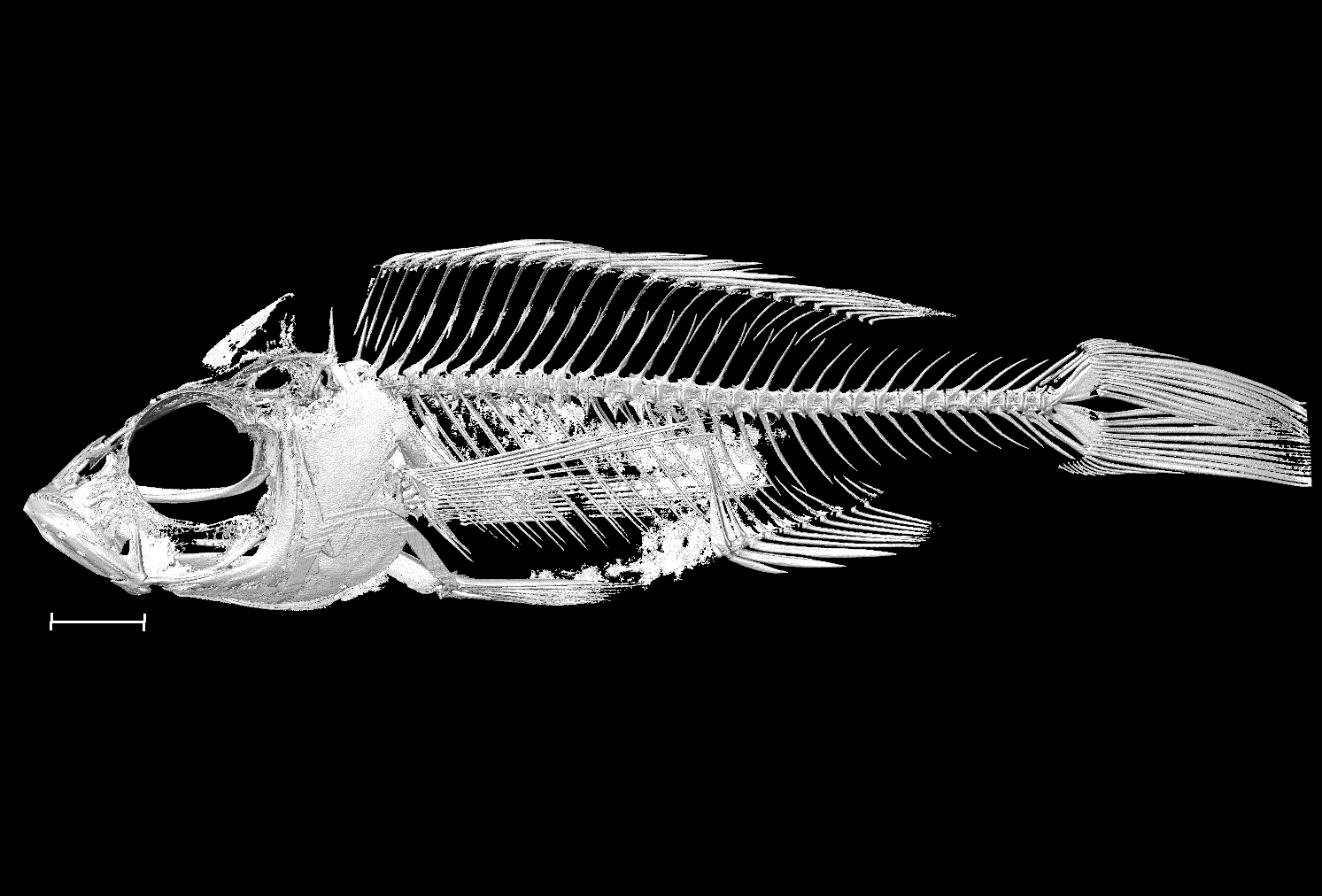

### Diplotaxodon_sp_macrops_ngulube_UniBri_DRC060_8bit_a.tif

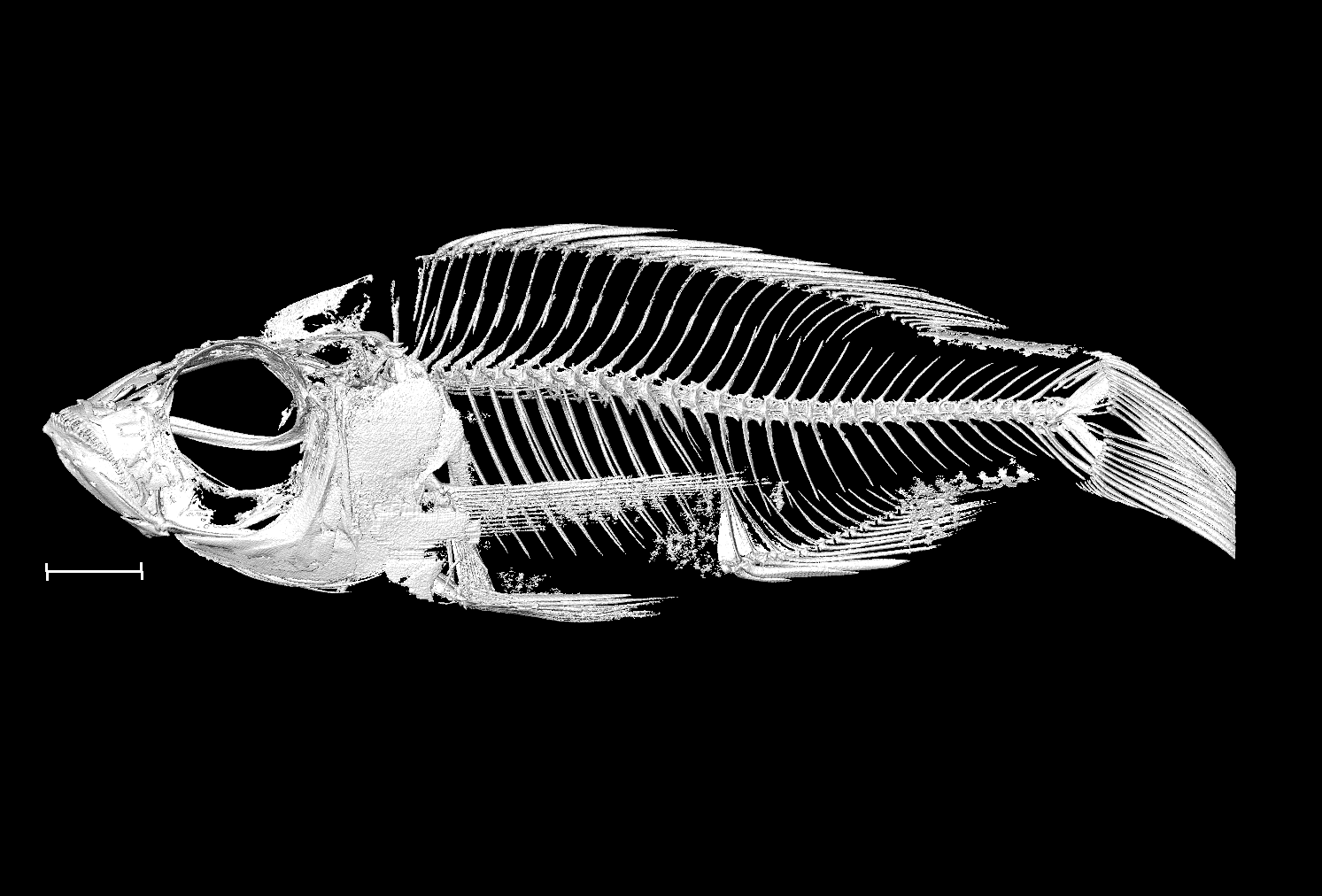

### Diplotaxodon_sp_macrops_ngulube_UniBri_DRC060_8bit_b.tif

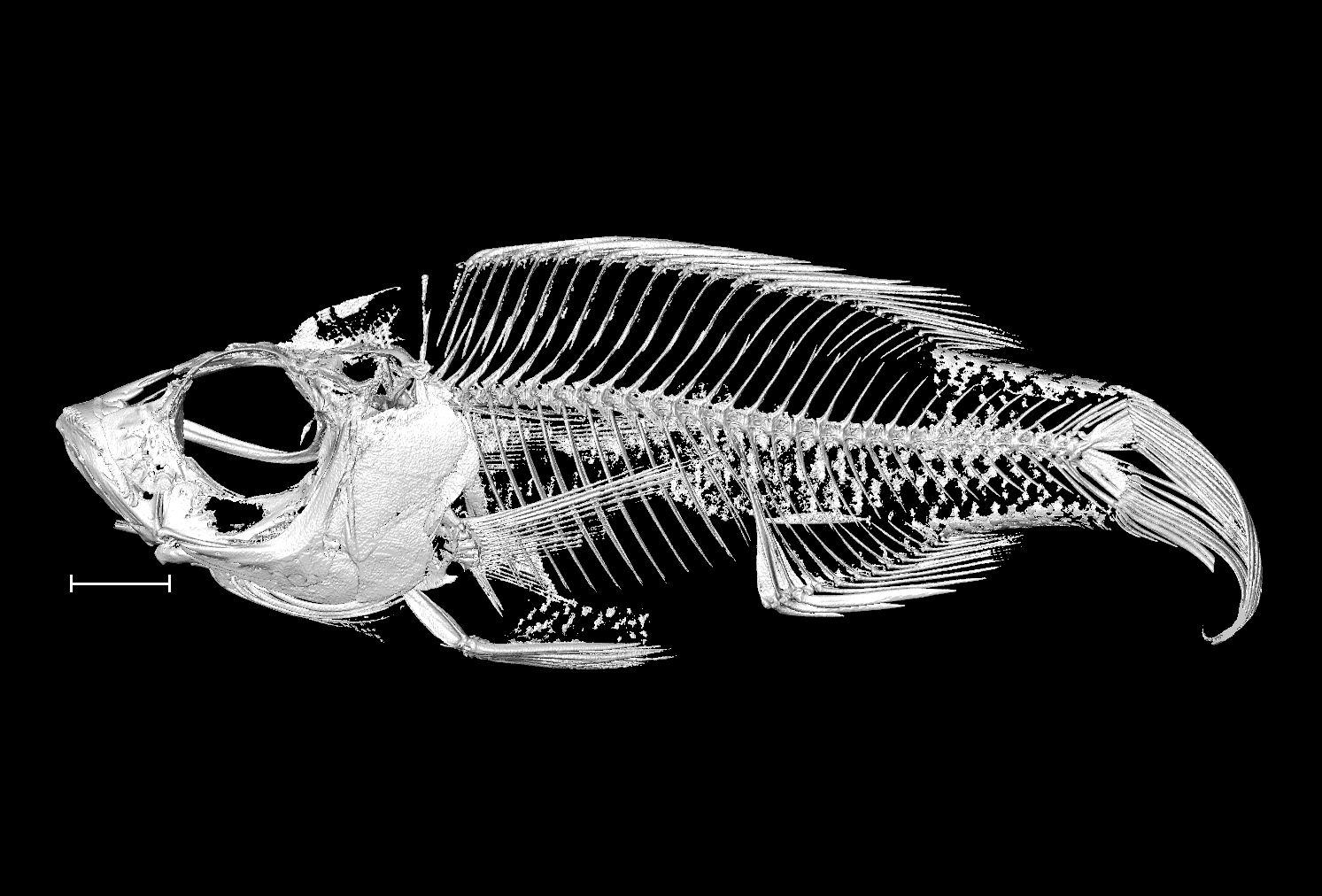

### Diplotaxodon_sp_similis_white_back_north_UniBri_DRC084_8bit.tif

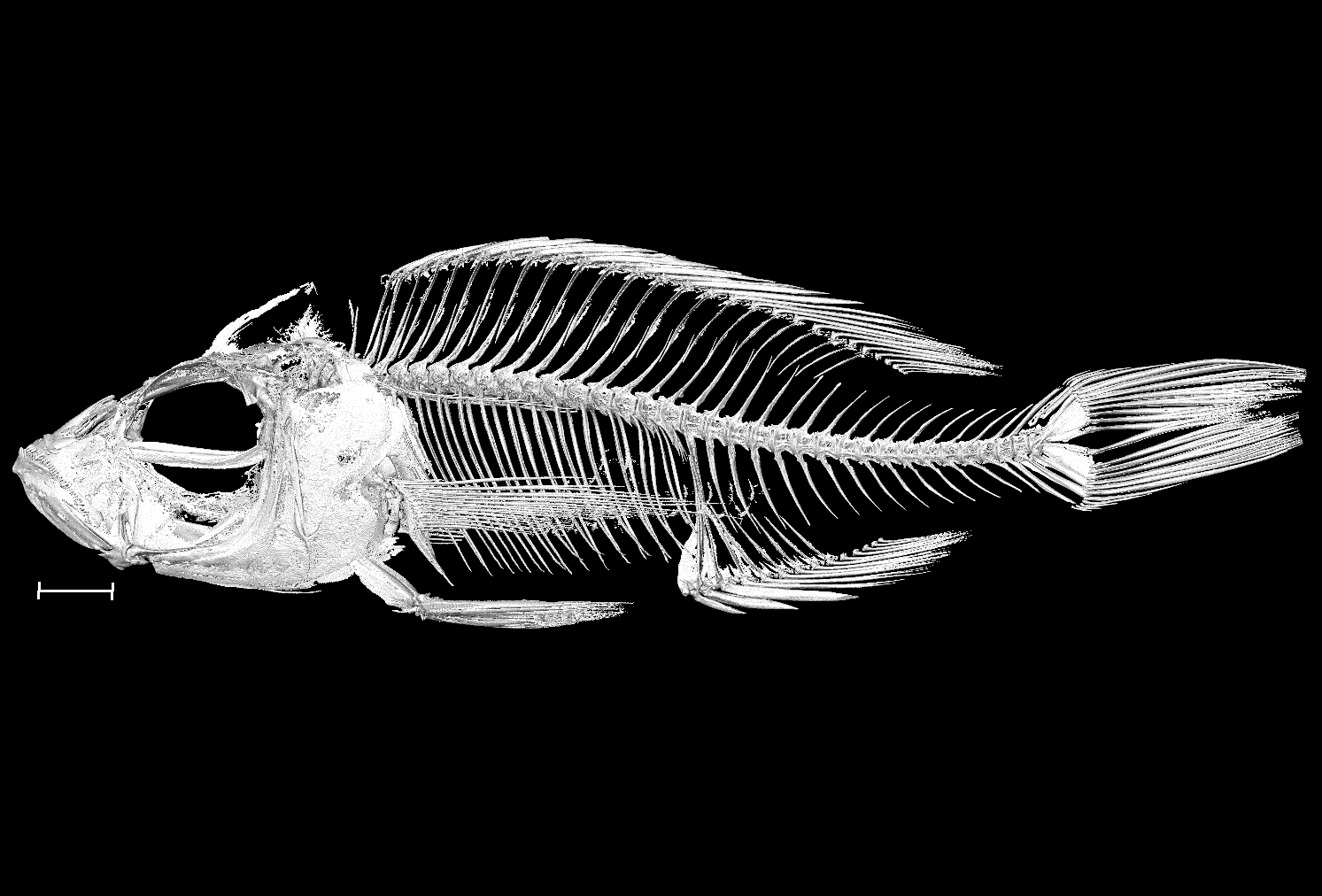

### Diplotaxodon_sp_similis_white_back_north_UniBri_DRC086_8bit.tif

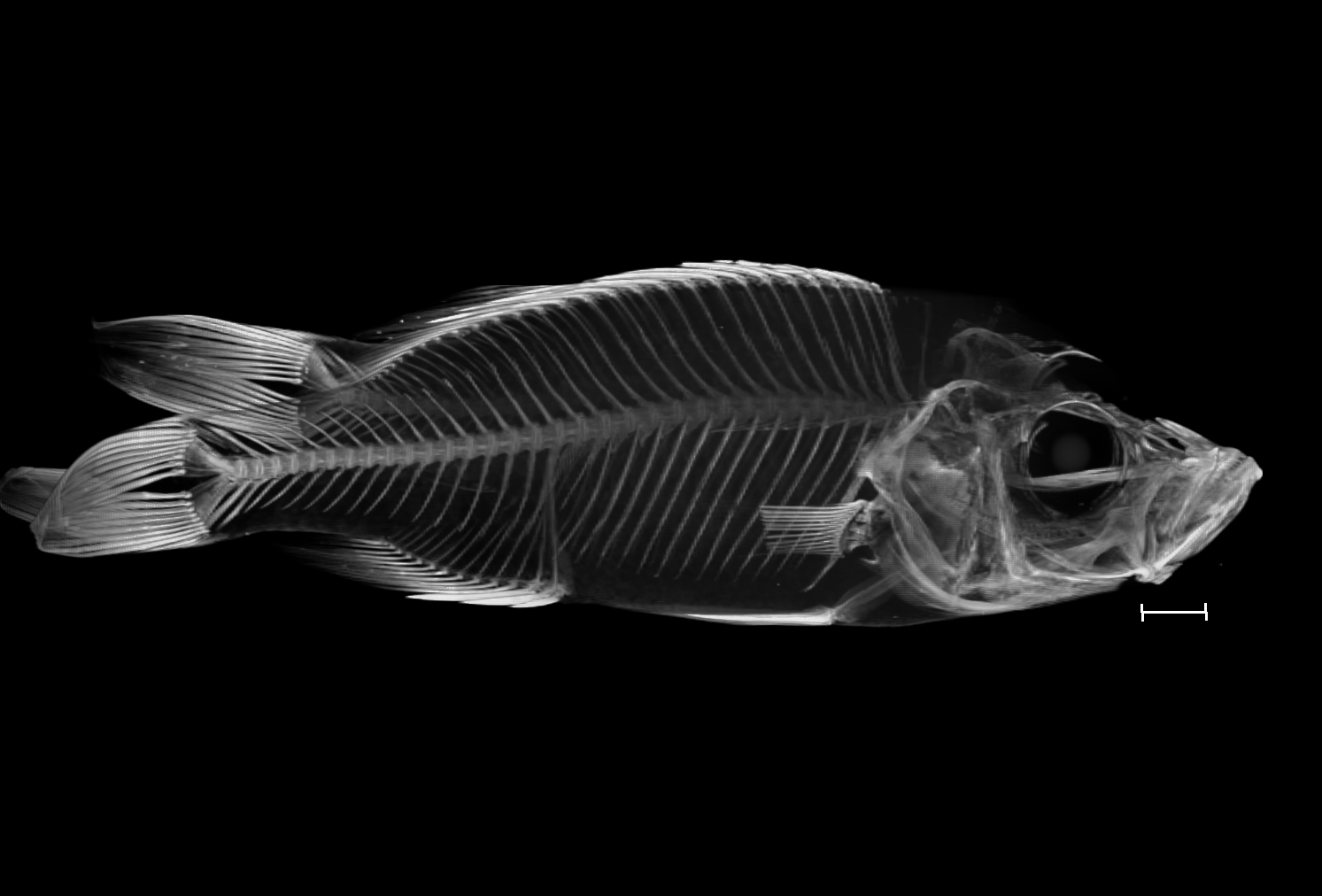

### Genyochromis_mento_NHMUK_1965_10_26_24_29_8bit_a.tif

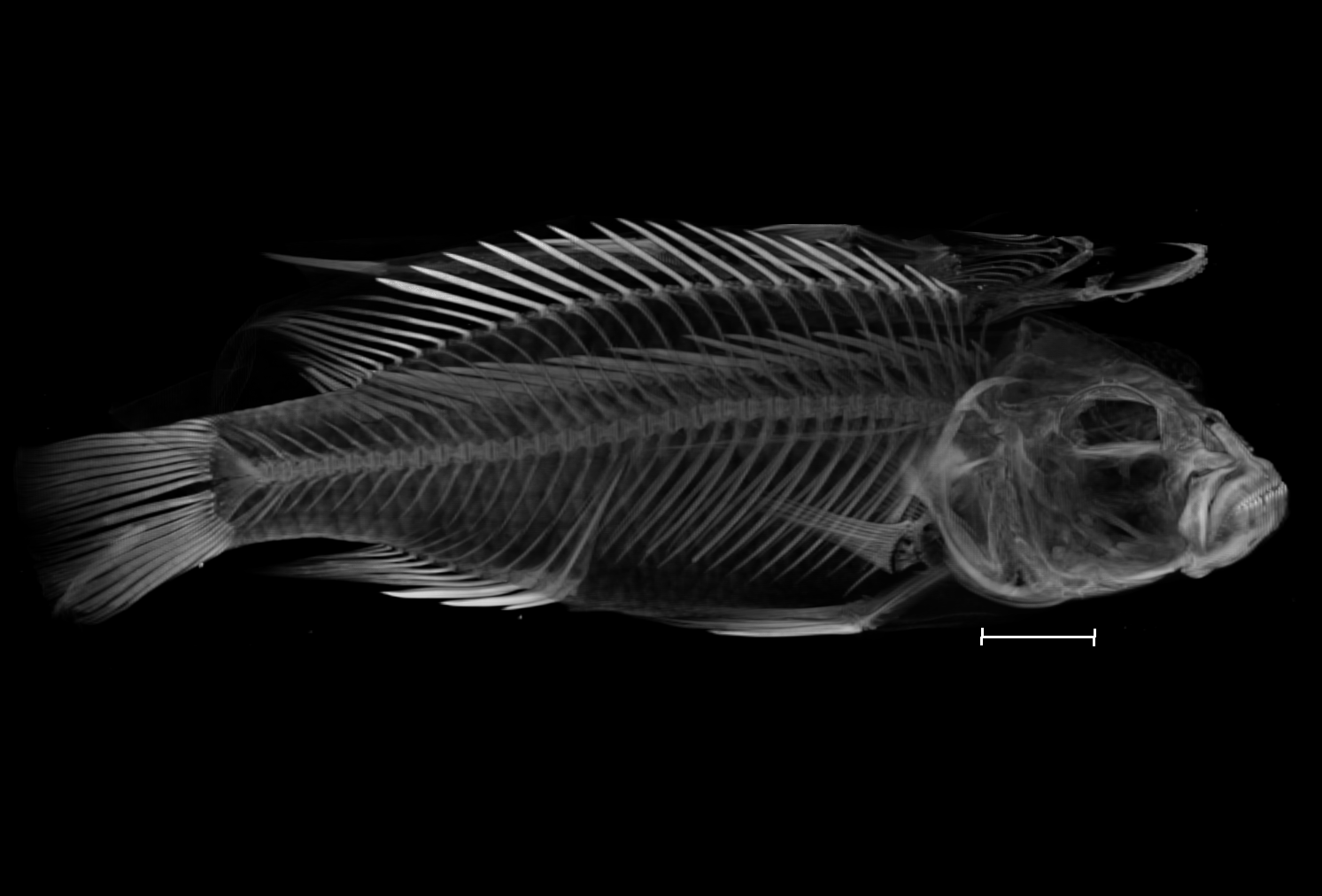

### Genyochromis_mento_NHMUK_1965_10_26_24_29_8bit_b.tif

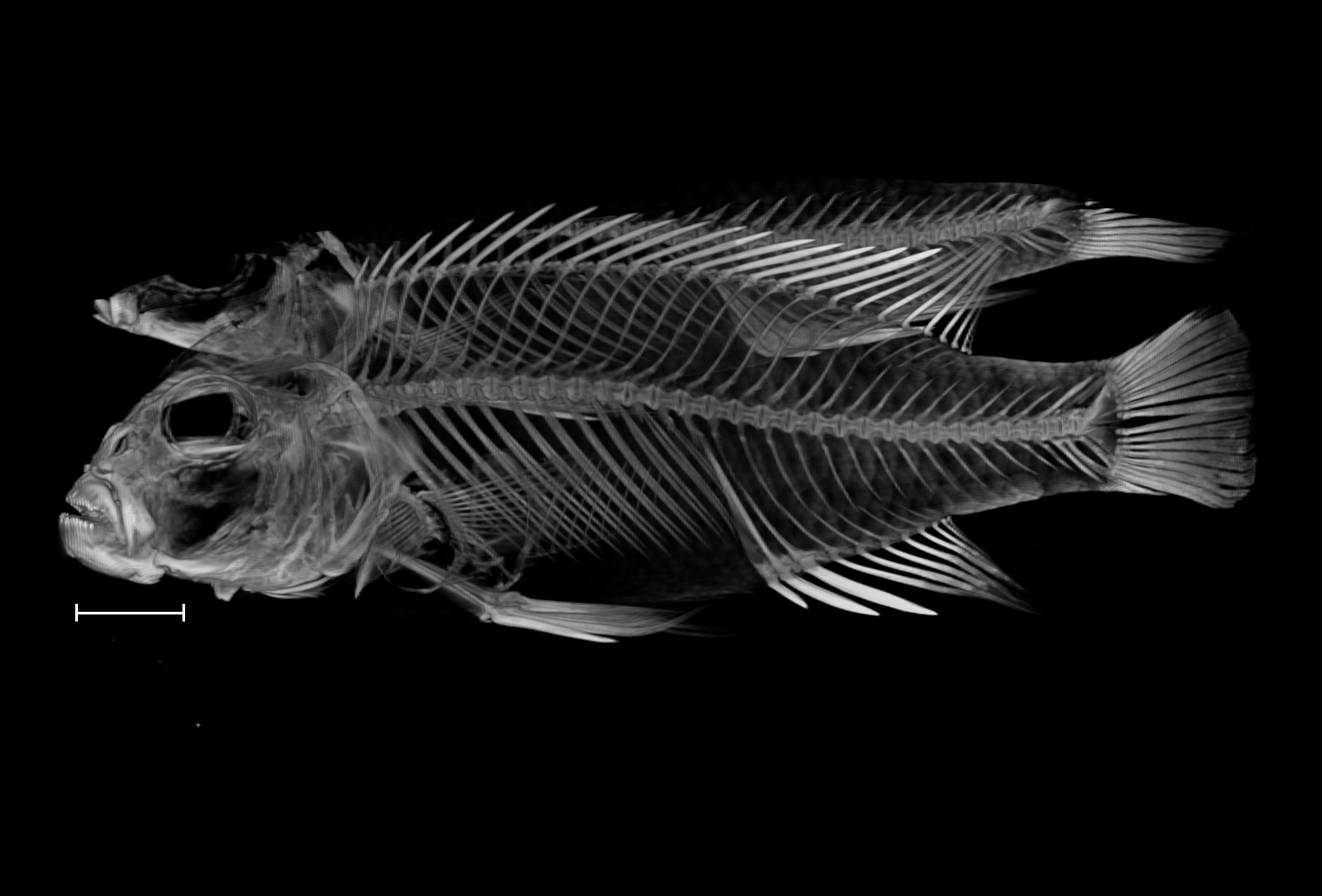

### Hemitilapia_oxyrhynchus_UniBri_40_8bit.tif

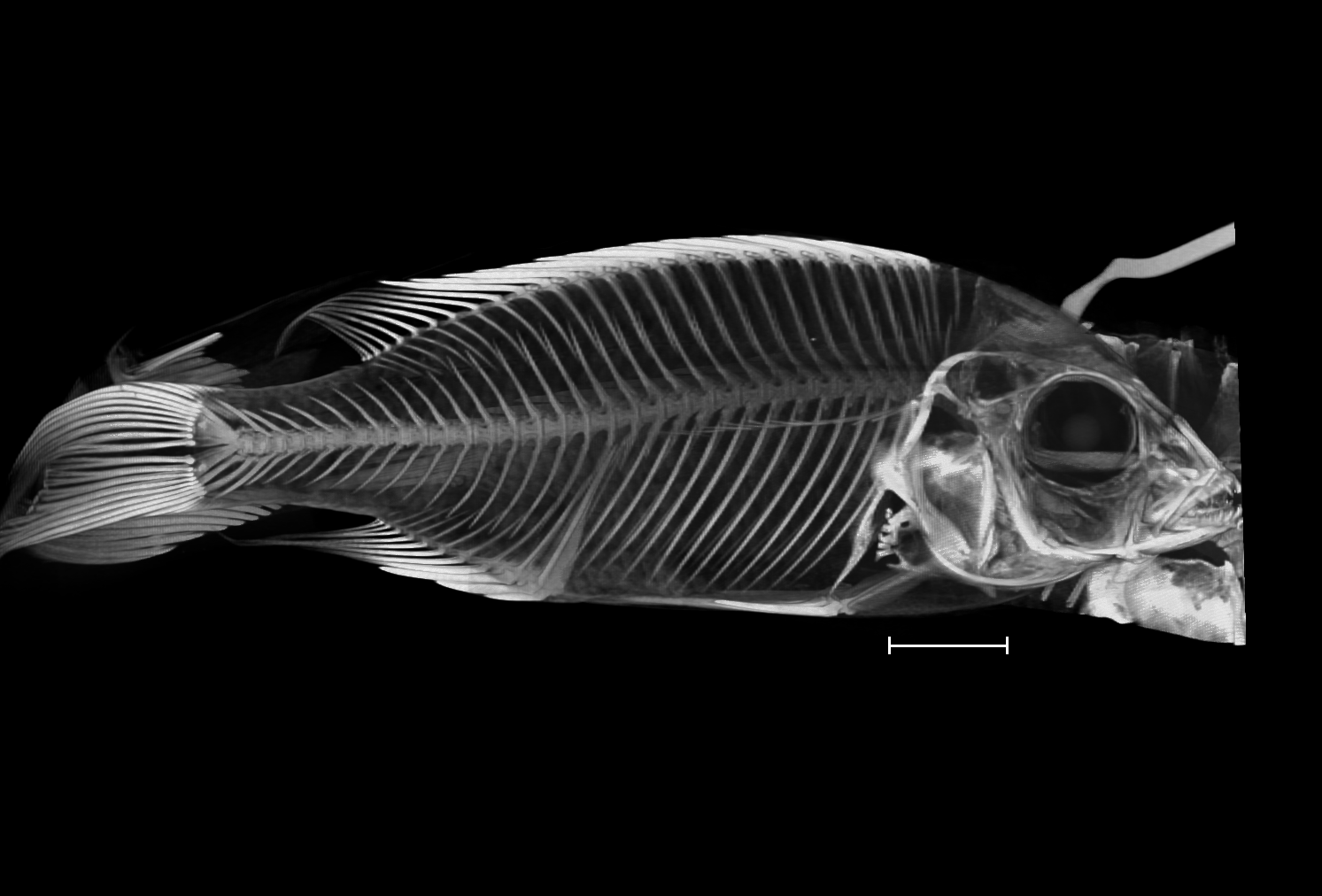

### Iodotropheus_sprengerae_NHMUK_1971_9_8_7_8_8bit_a.tif

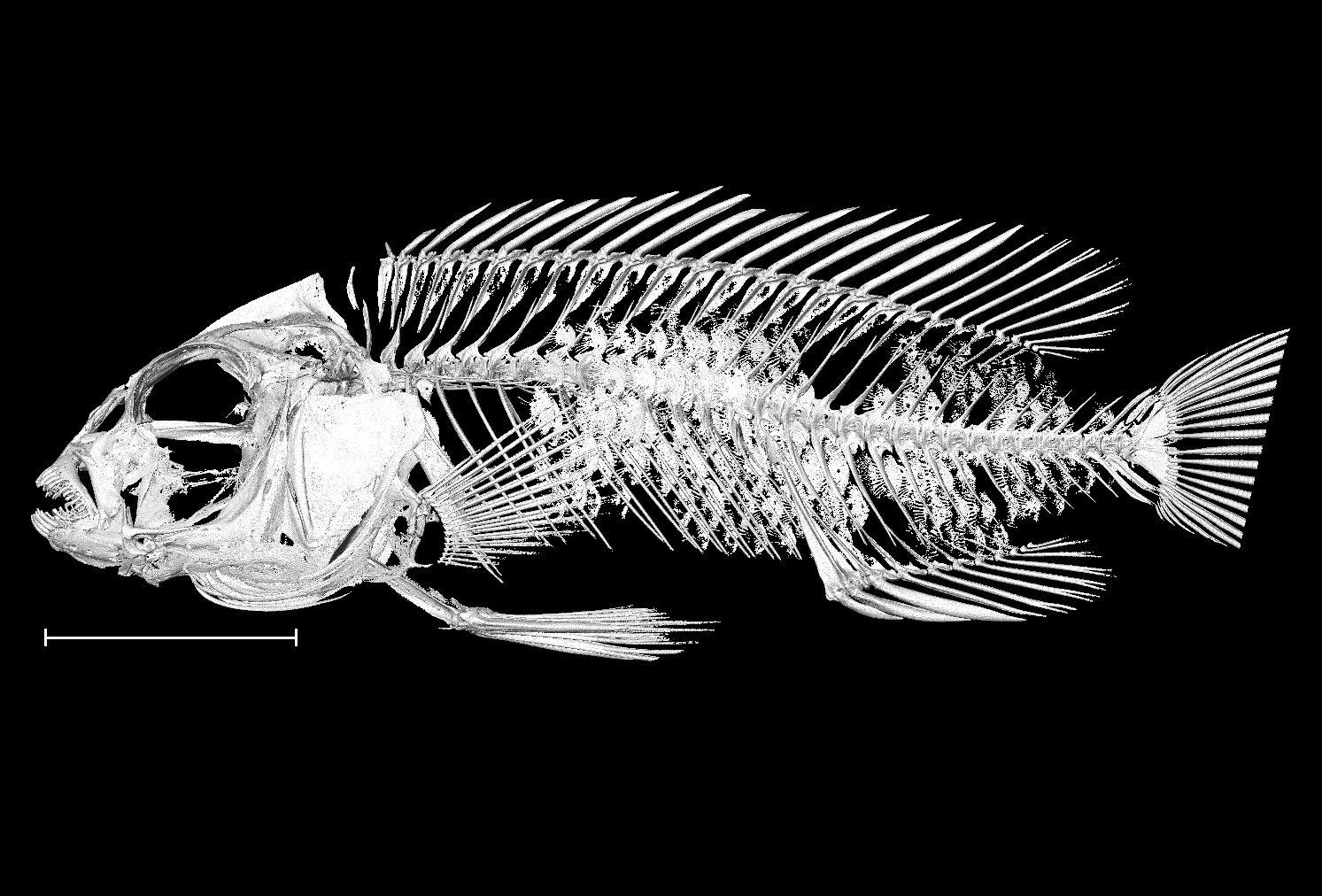

### Iodotropheus_sprengerae_NHMUK_1971_9_8_7_8_8bit_b.tif

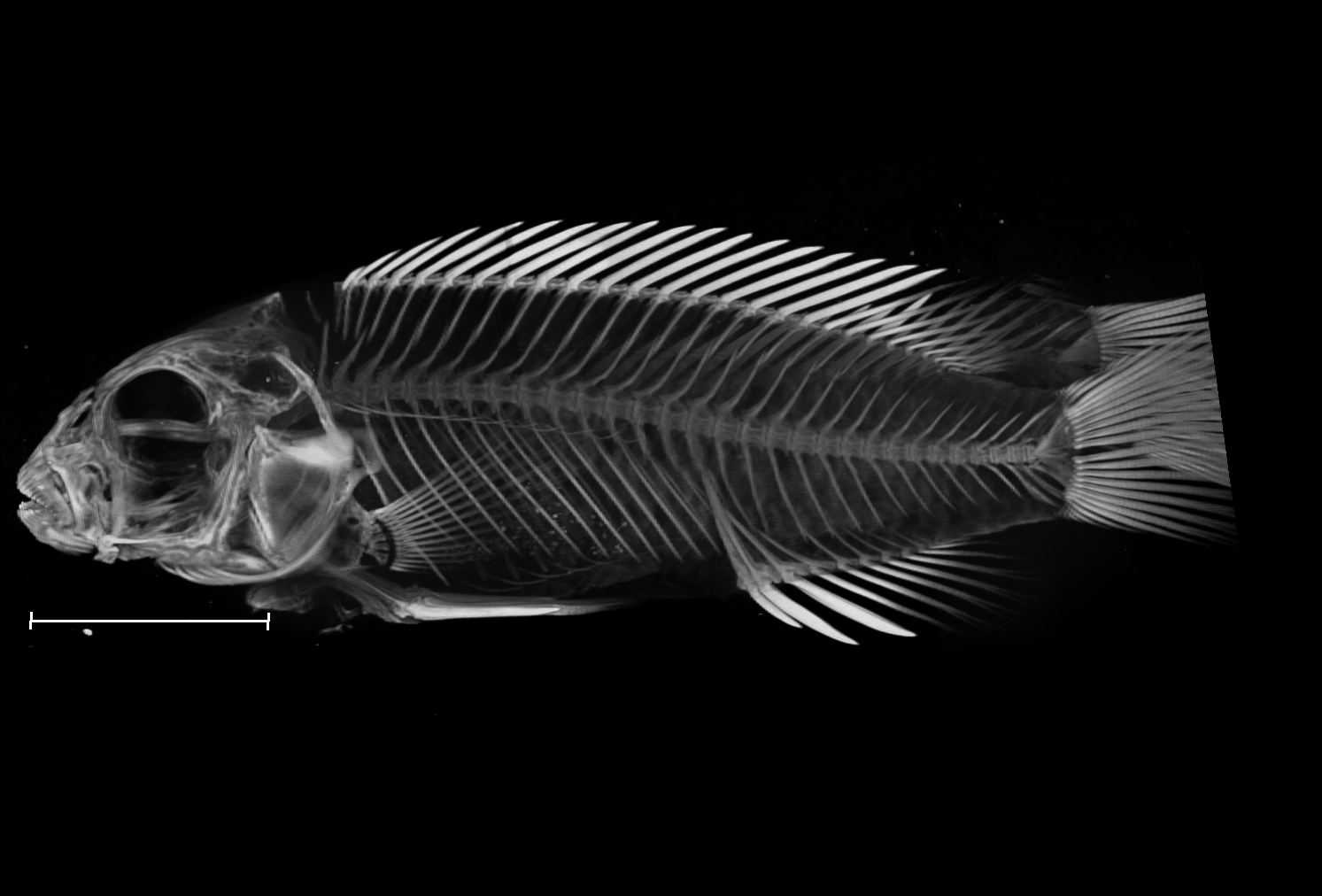

### Labidochromis_strigatus_NHMUK_1981_1_9_28_32_8bit_a.tif

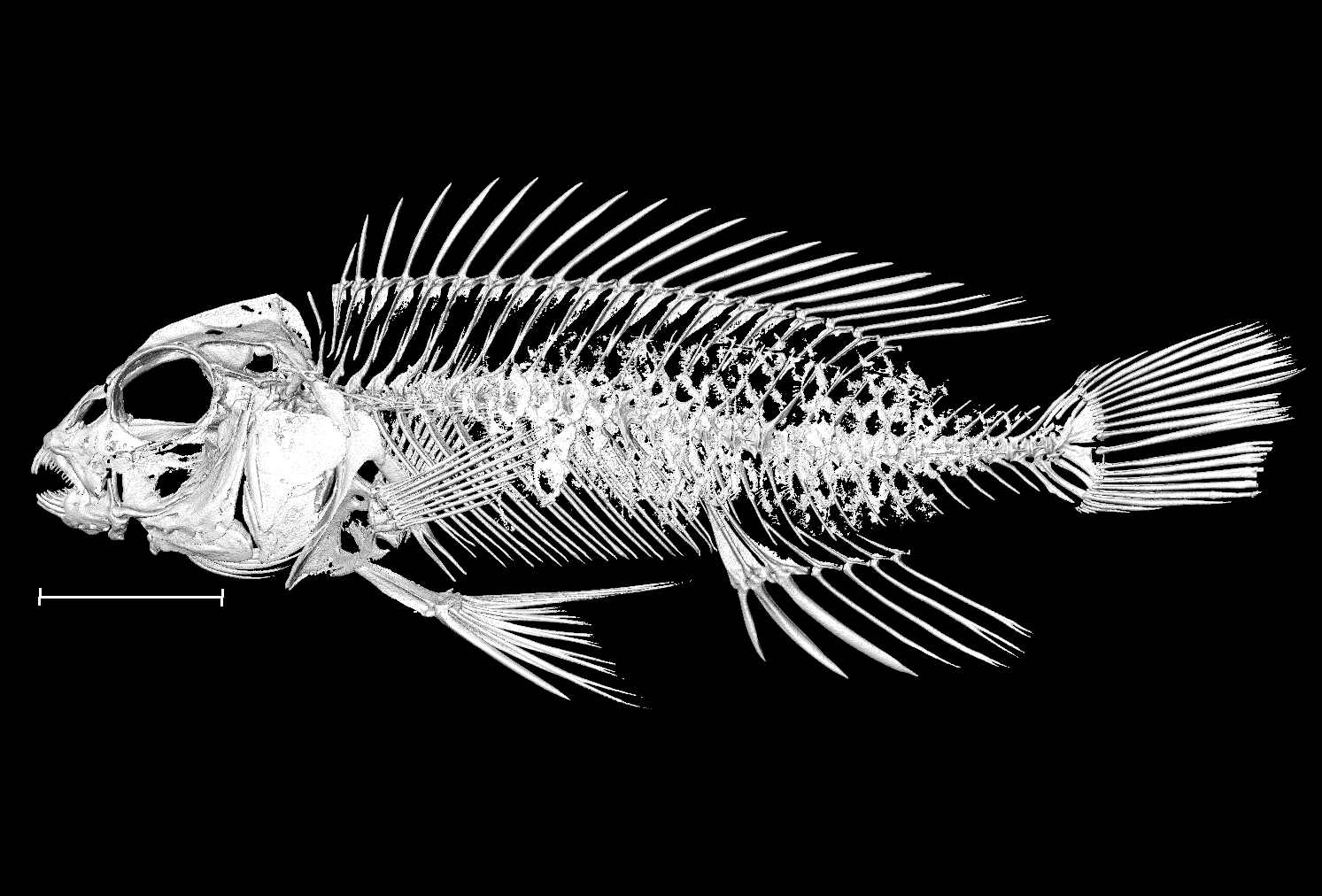

### Labidochromis_strigatus_NHMUK_1981_1_9_28_32_8bit_b.tif

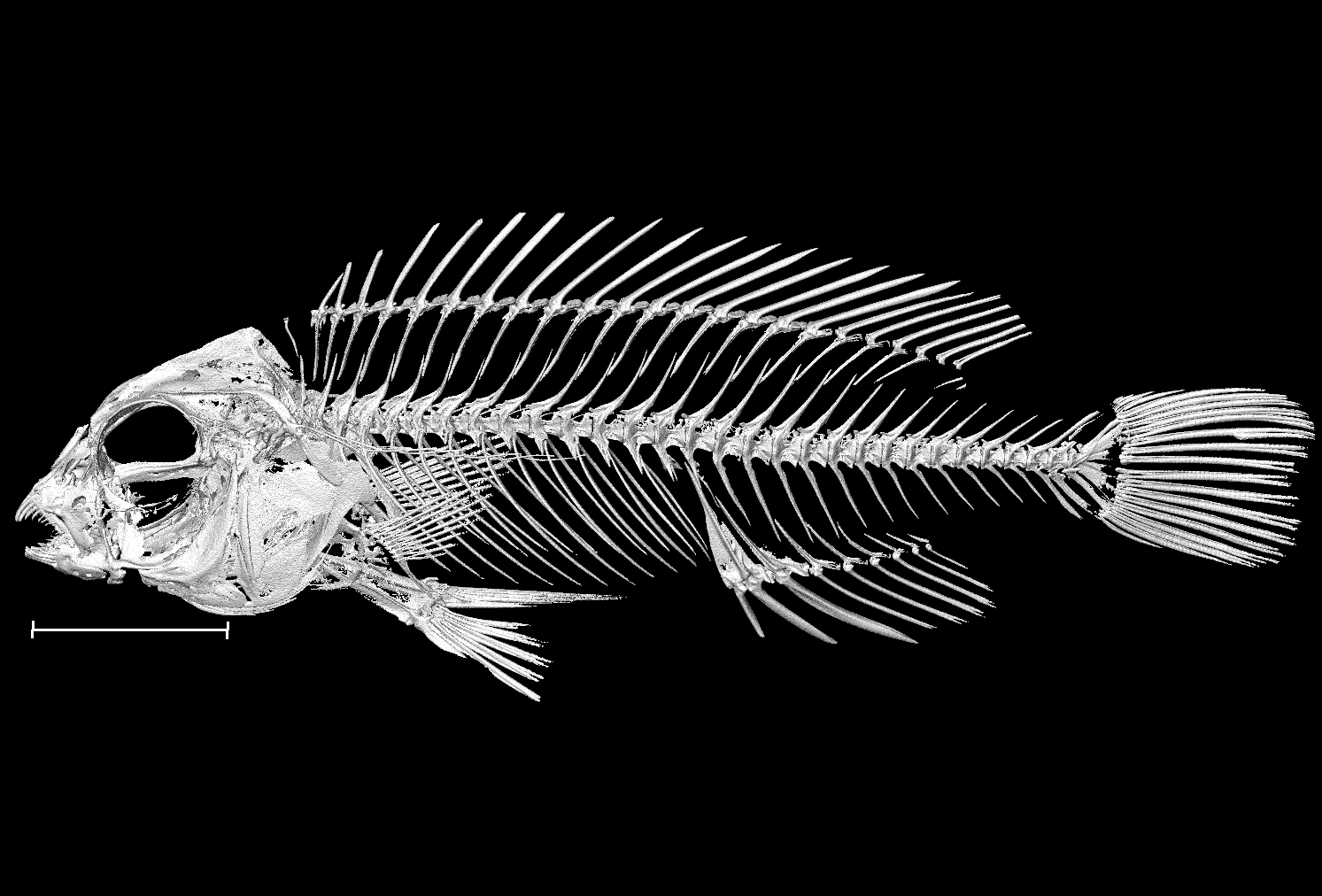

### Lethrinops_albus_NHMUK_1930_1_31_171_180_8bit_a.tif

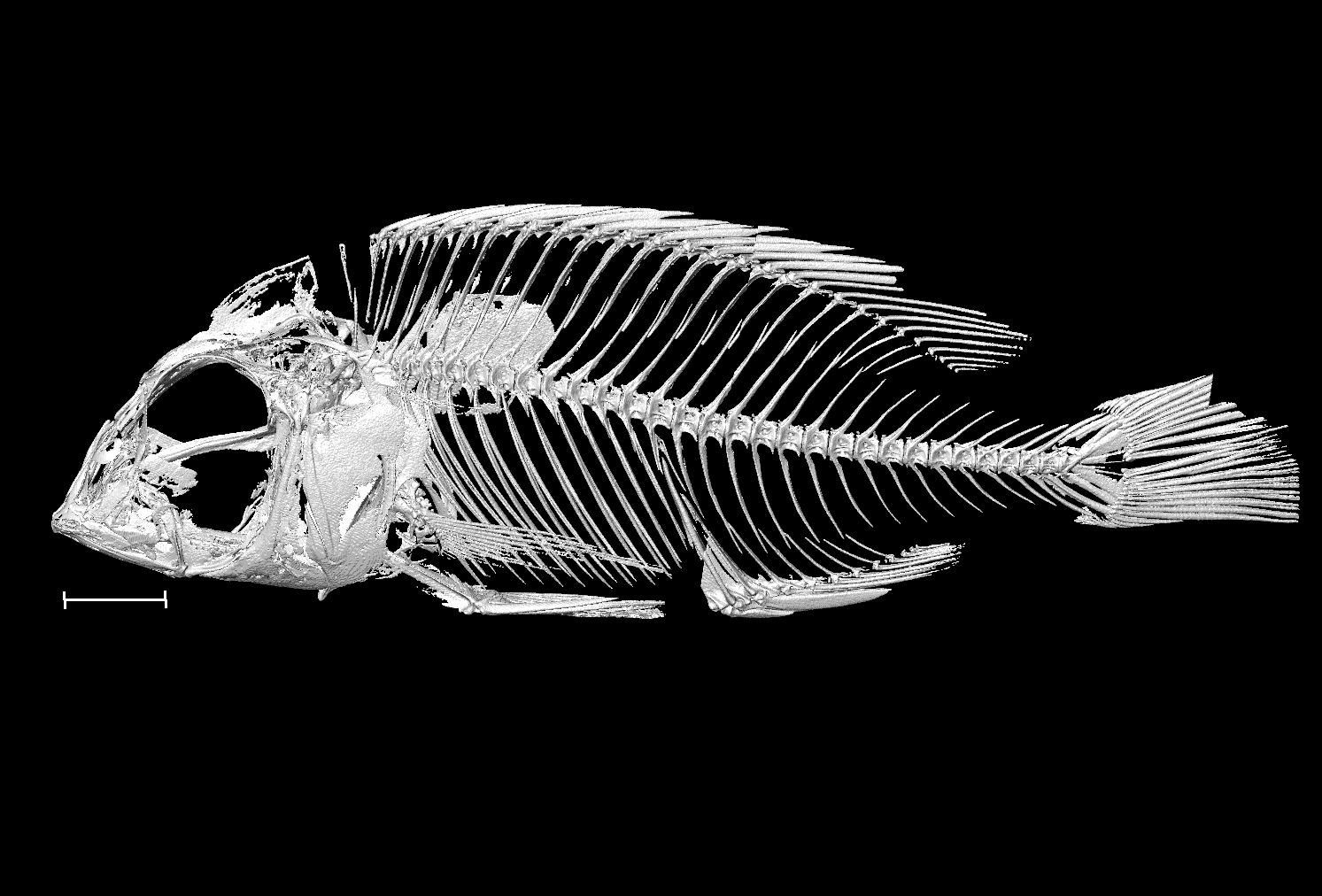

### Lethrinops_albus_NHMUK_1930_1_31_171_180_8bit_b.tif

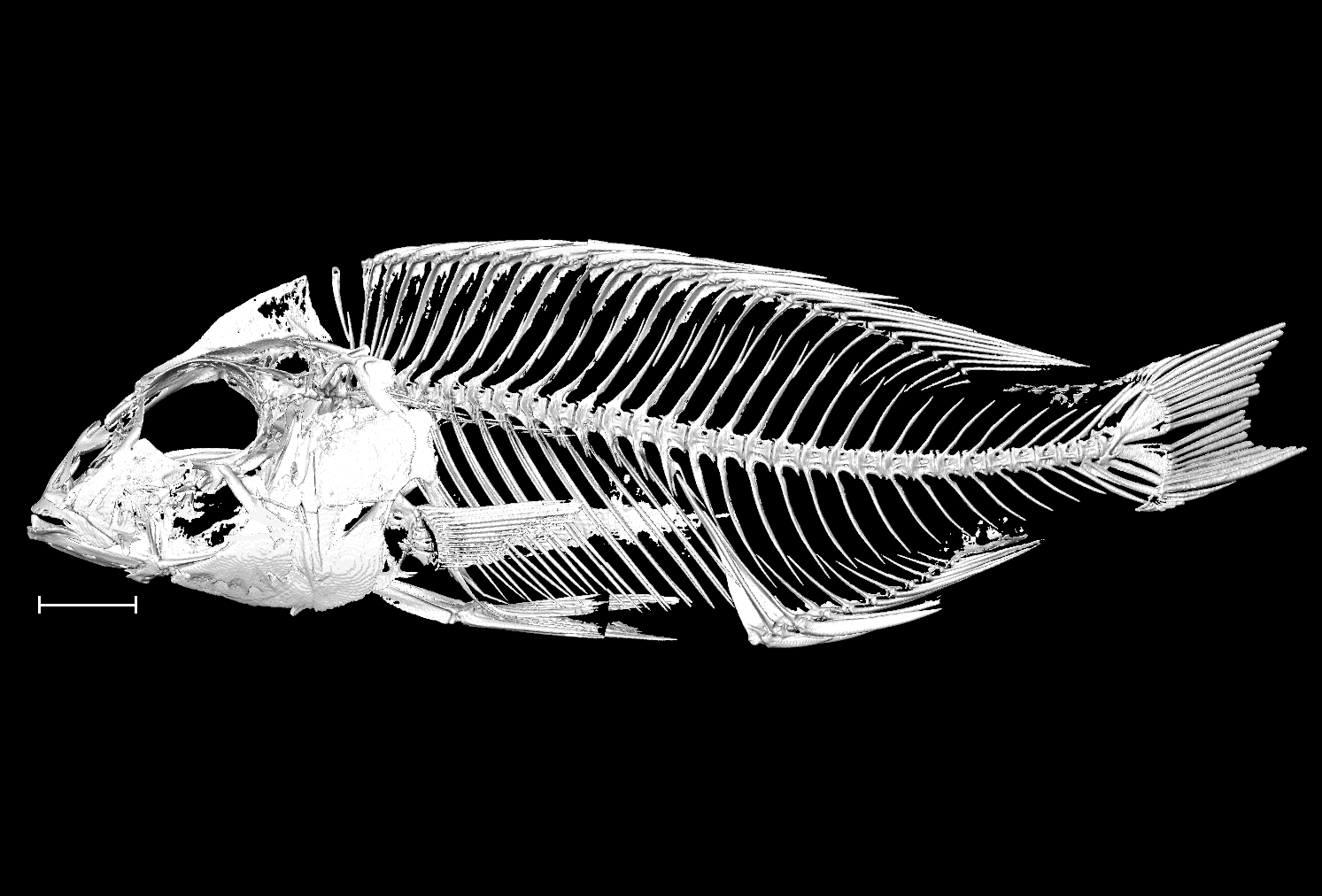

### Lethrinops_auritus_NHMUK_1978_10_31_2_6_8bit_a.tif

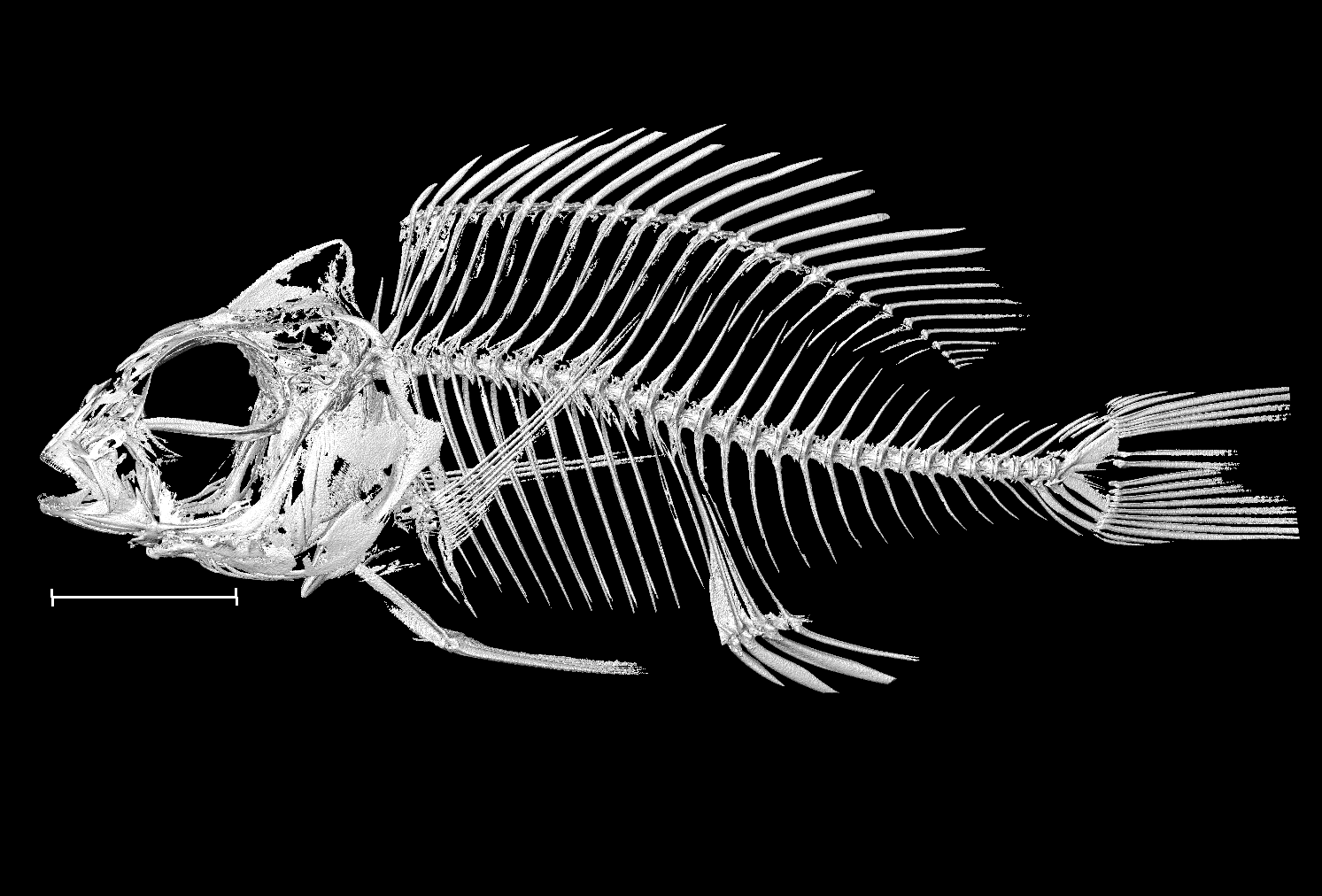

### Lethrinops_auritus_NHMUK_1978_10_31_2_6_8bit_b.tif

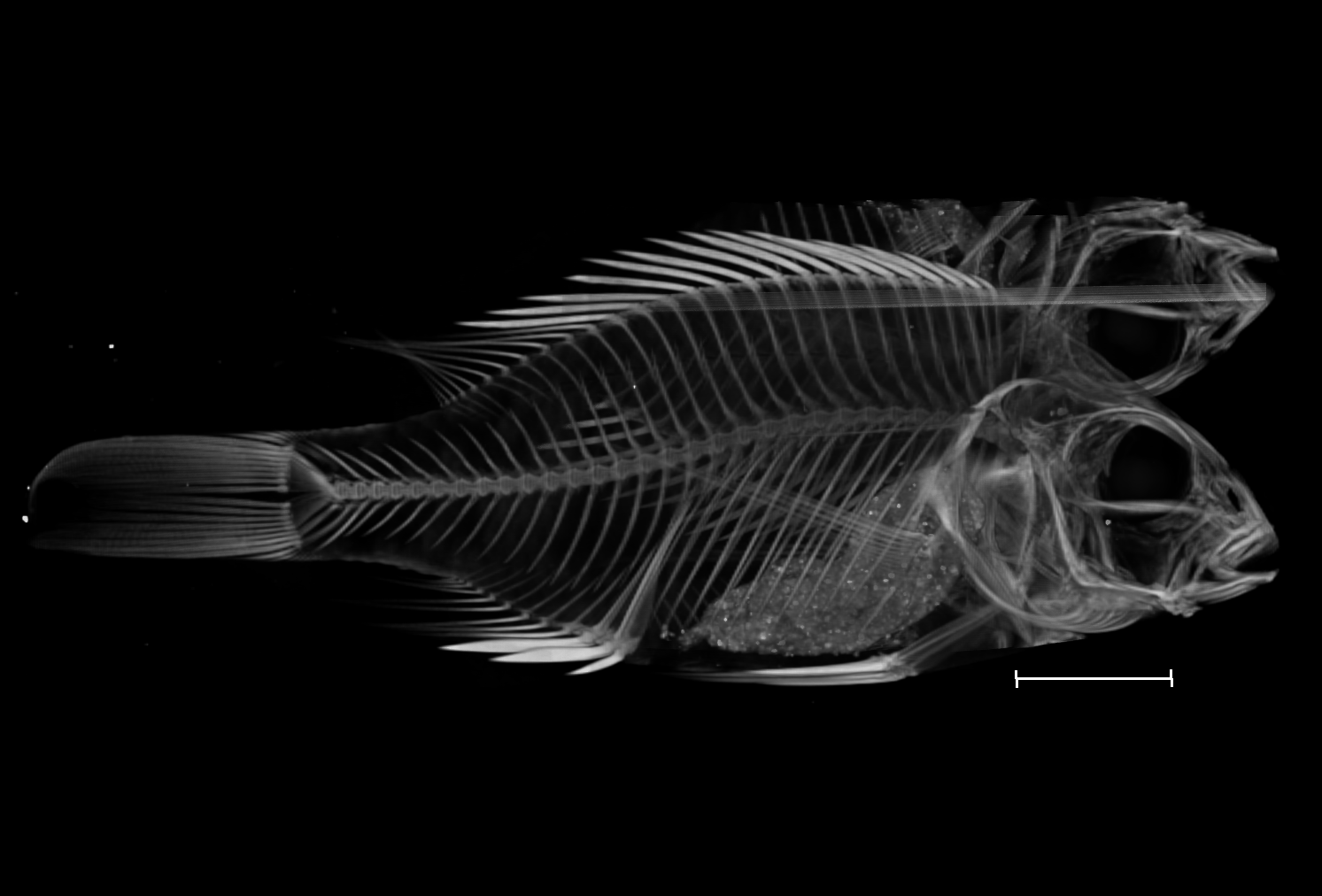

### Lethrinops_gossei_UniBri_refonly_8bit.tif

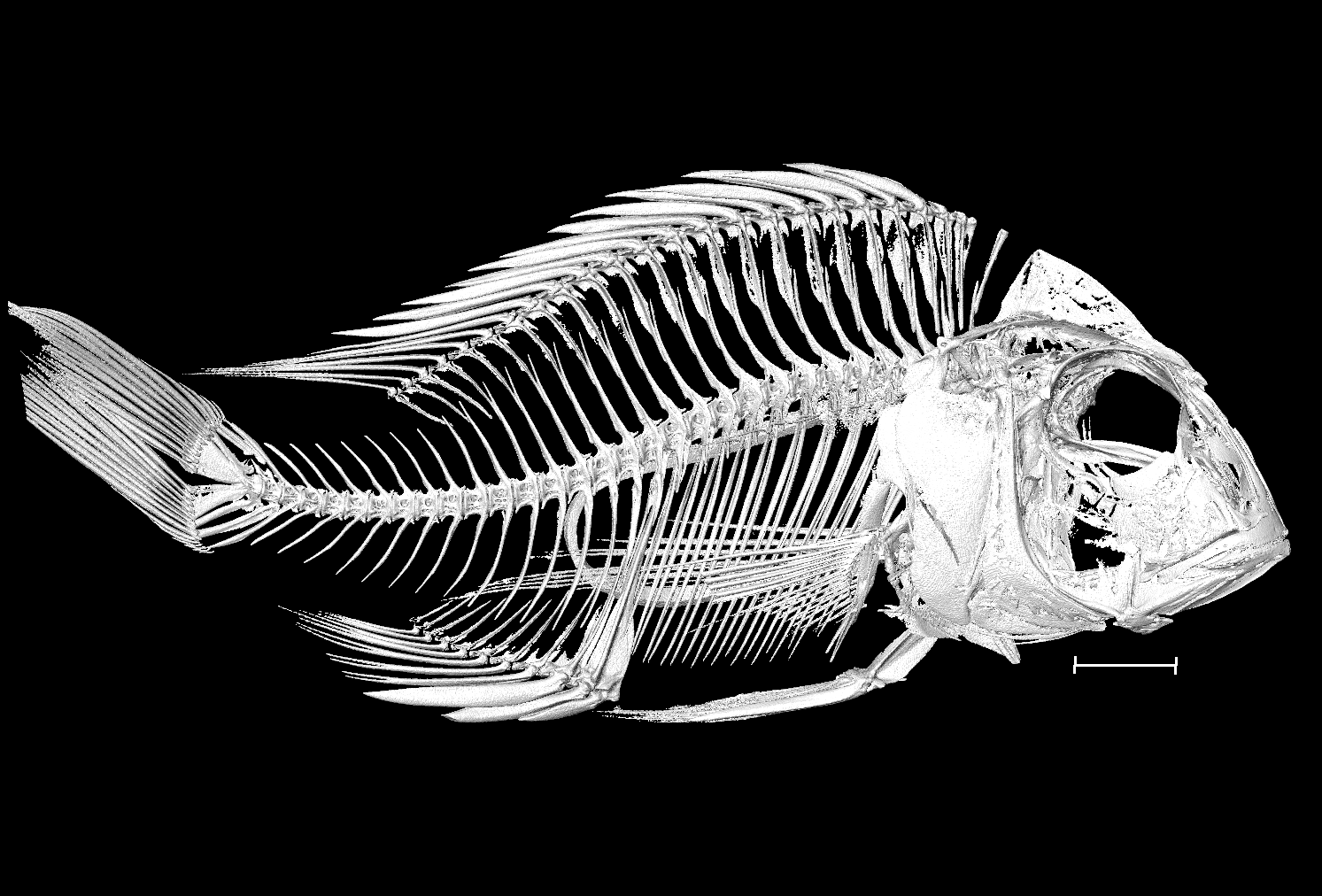

### Maylandia_zebra_UniOxf_M2_8bit.tif

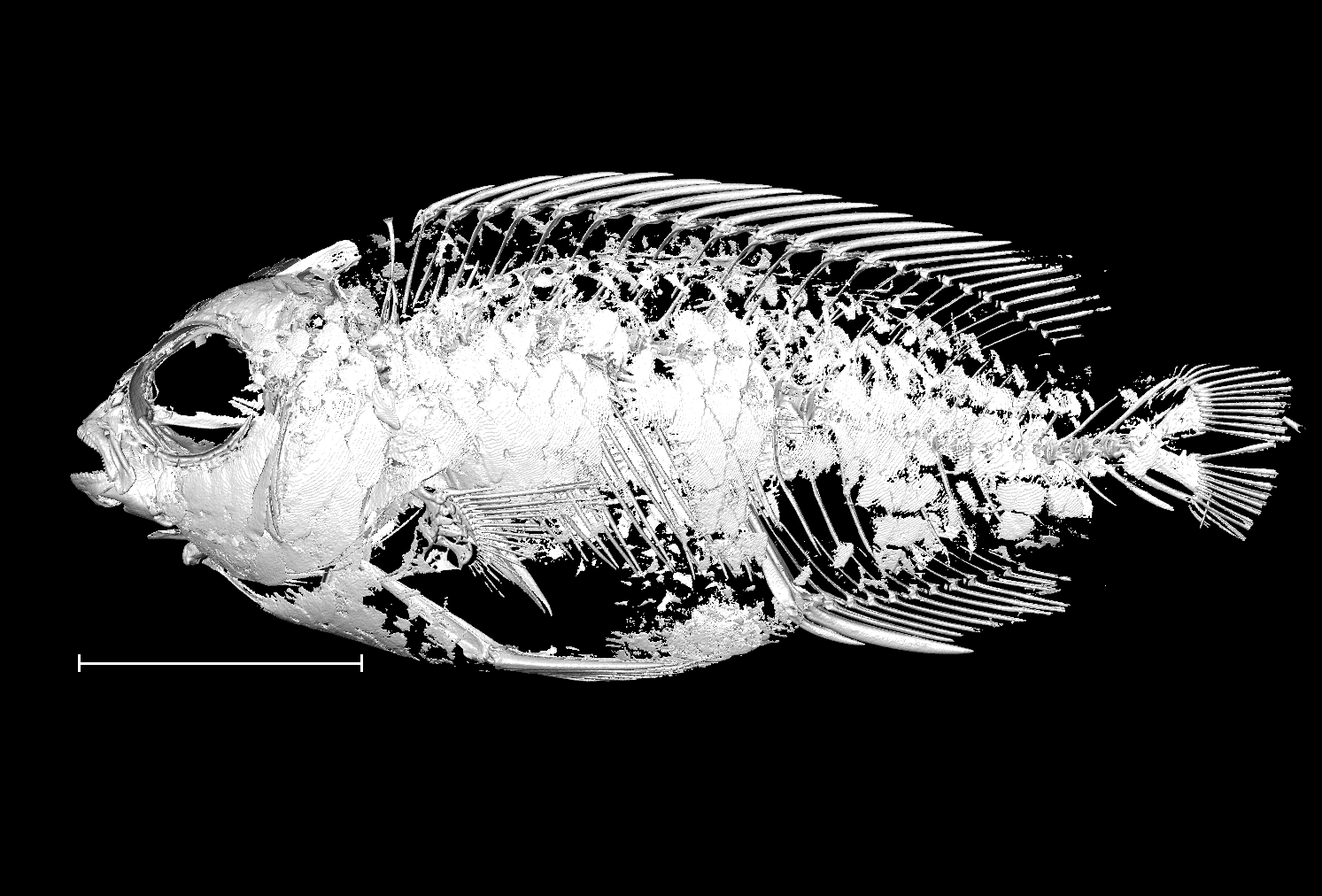

### Maylandia_zebra_UniOxf_M3_8bit.tif

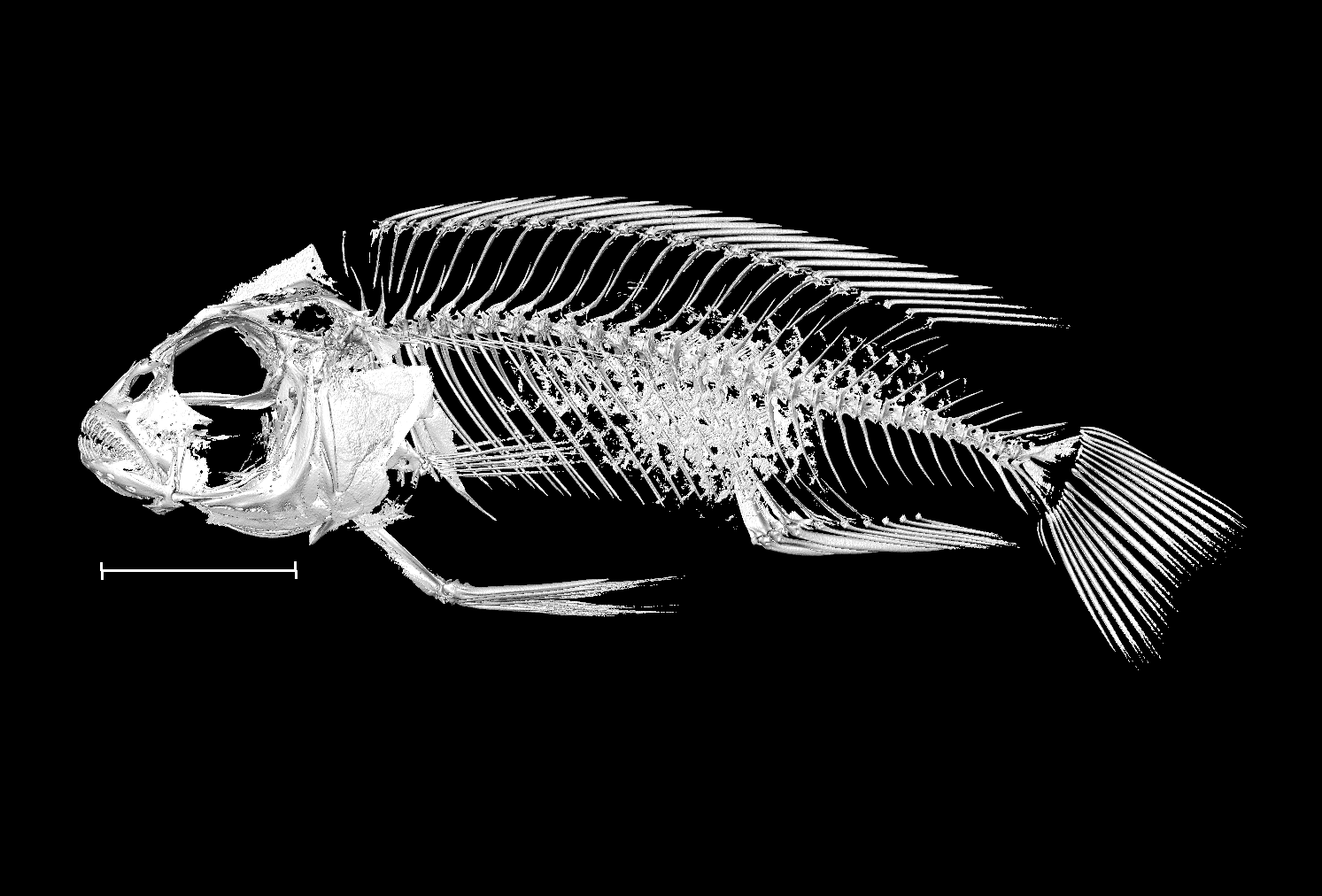

### Maylandia_zebra_UniOxf_M4_8bit.tif

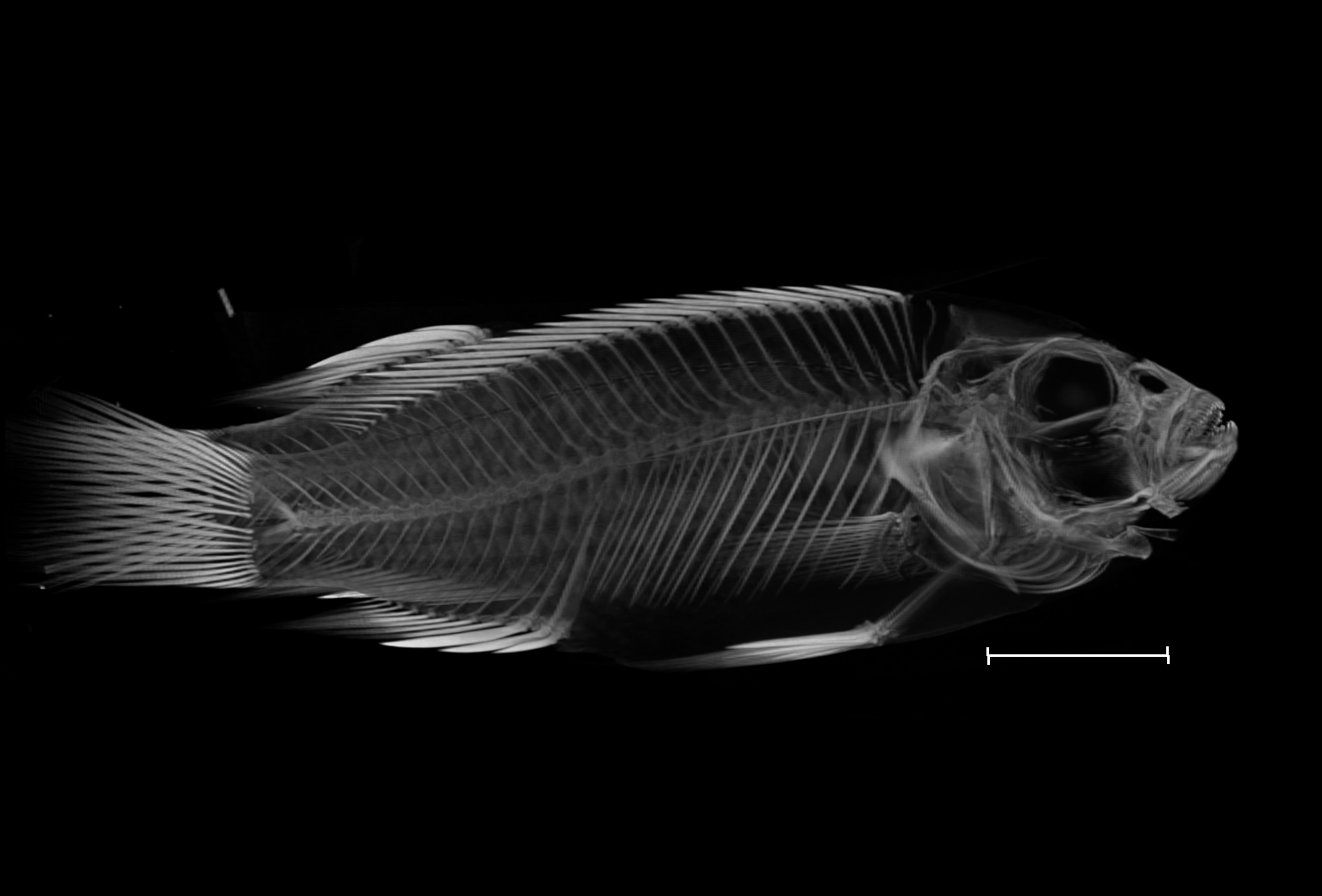

### Maylandia_zebra_UniOxf_M5_8bit.tif

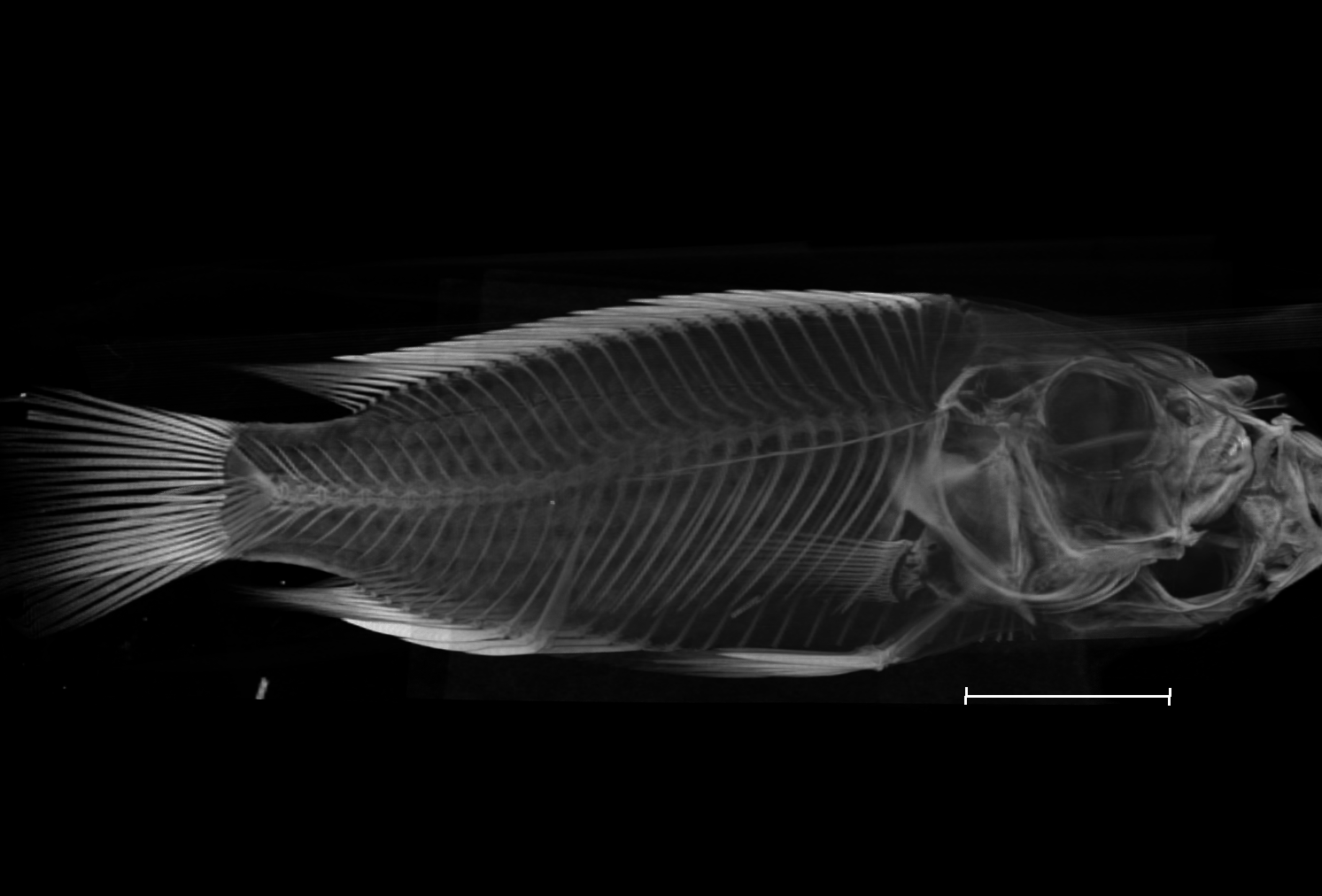

### Maylandia_zebra_UniOxf_M6_8bit.tif

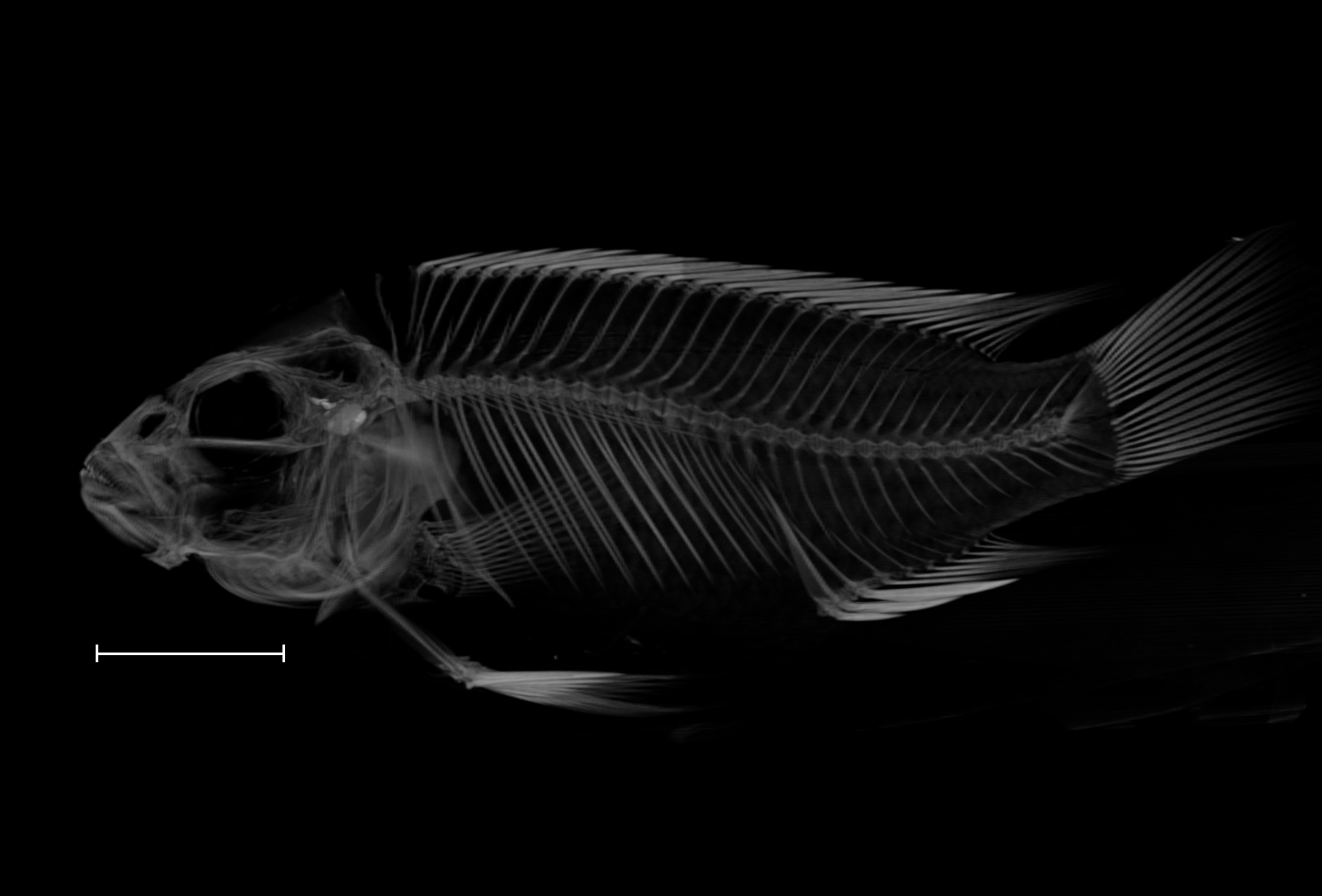

### Mylochromis_anaphyrmus_UniBri_132_8bit.tif

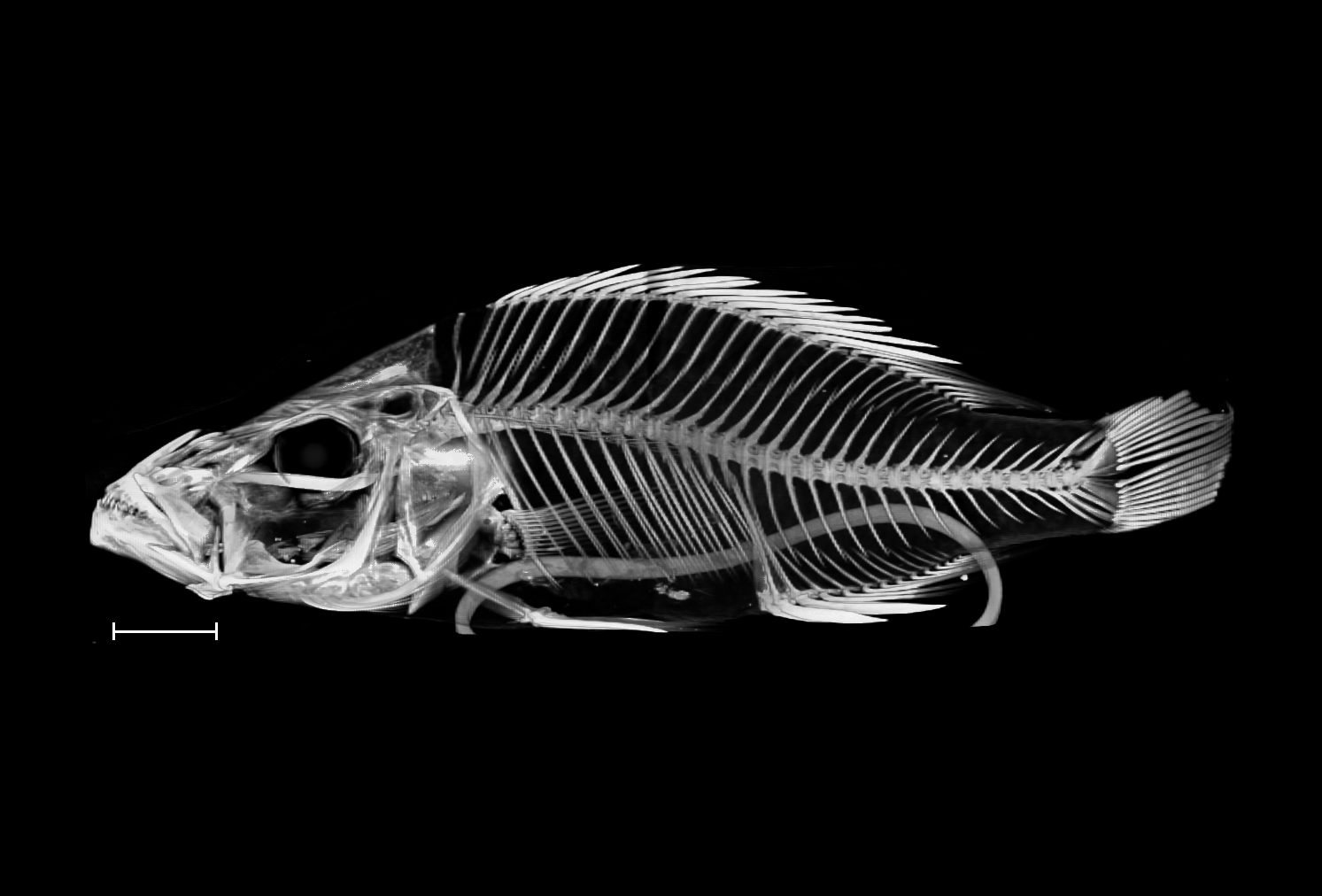

### Nimbochromis_linni_UniBri_440_8bit.tif

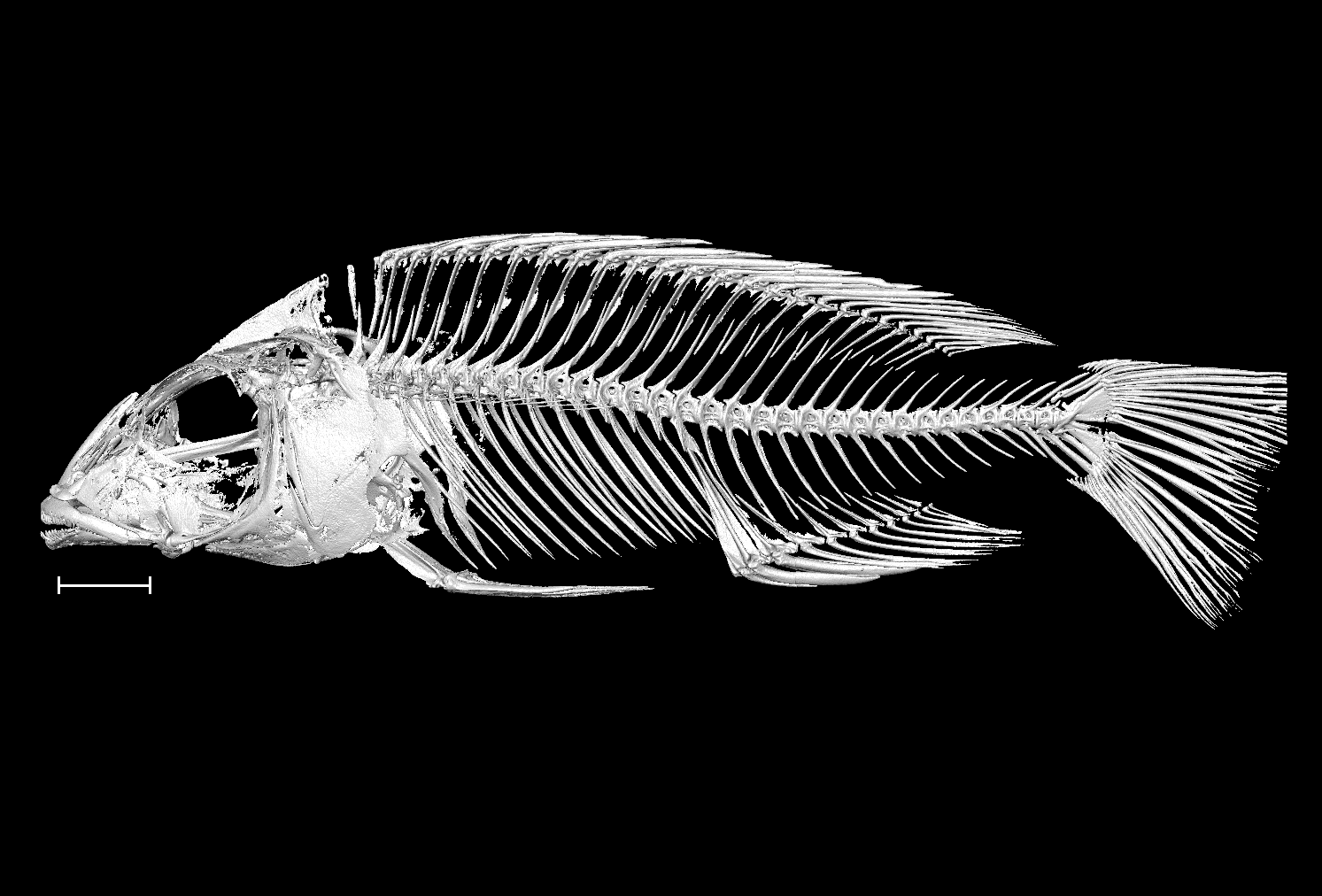

### Nimbochromis_livingstonii_NHMUK_1896_10_5_25_8bit.tif

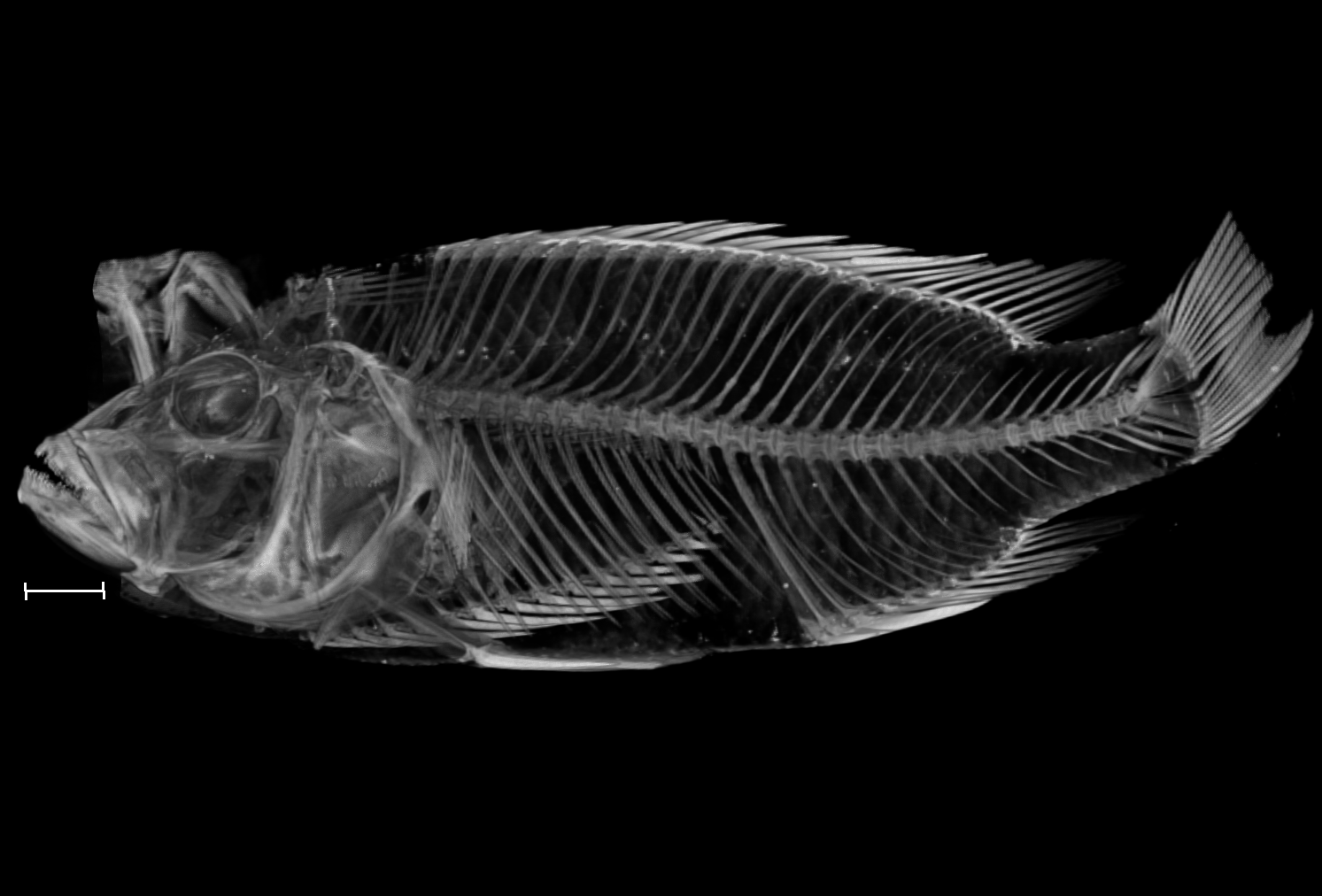

### Nimbochromis_polystigma_NHMUK_1921_9_6_102_107_8bit_a.tif

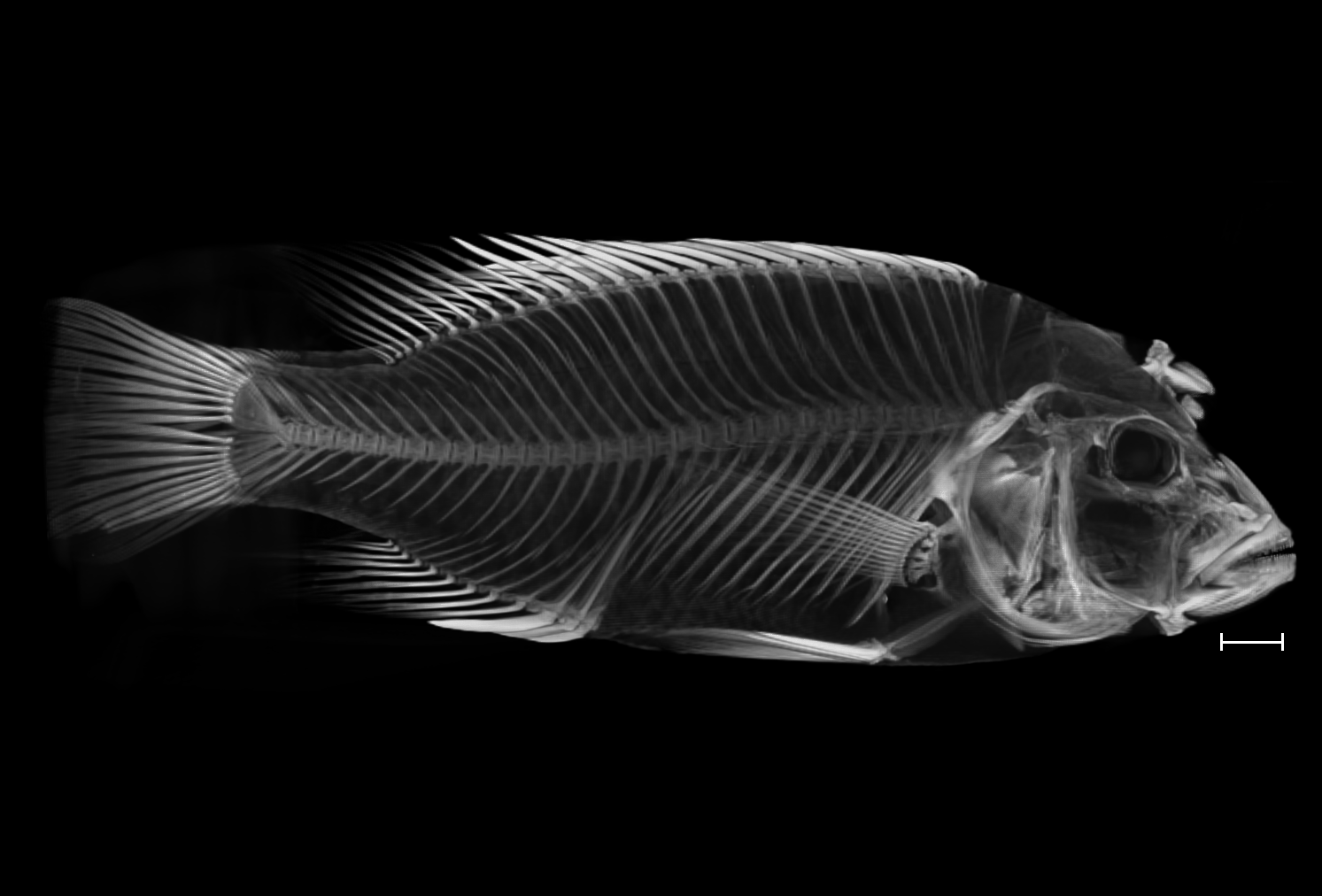

### Nimbochromis_polystigma_NHMUK_1921_9_6_102_107_8bit_b.tif

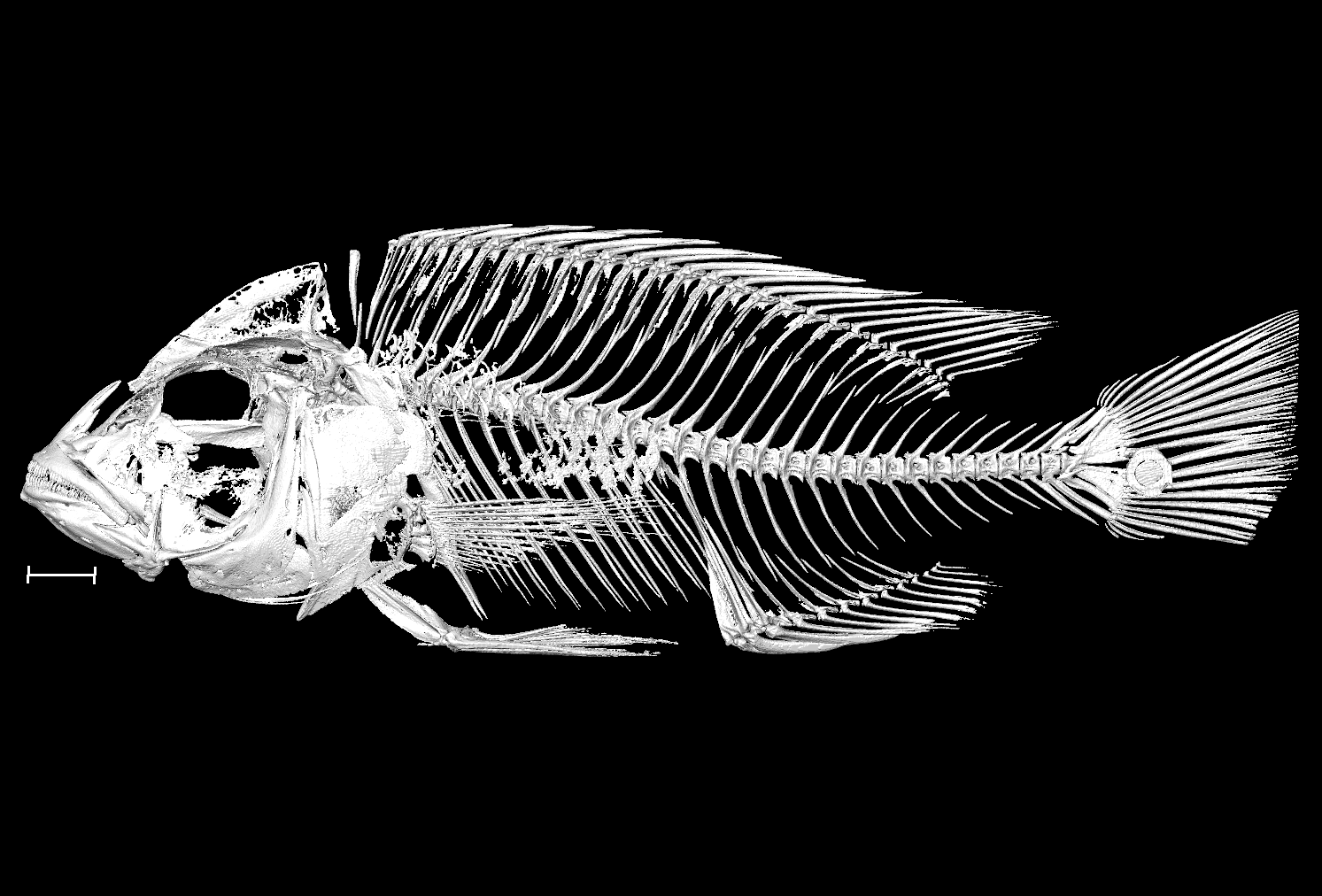

### Otopharynx_lithobates_NHMUK_1974_7_5_1_8bit.tif

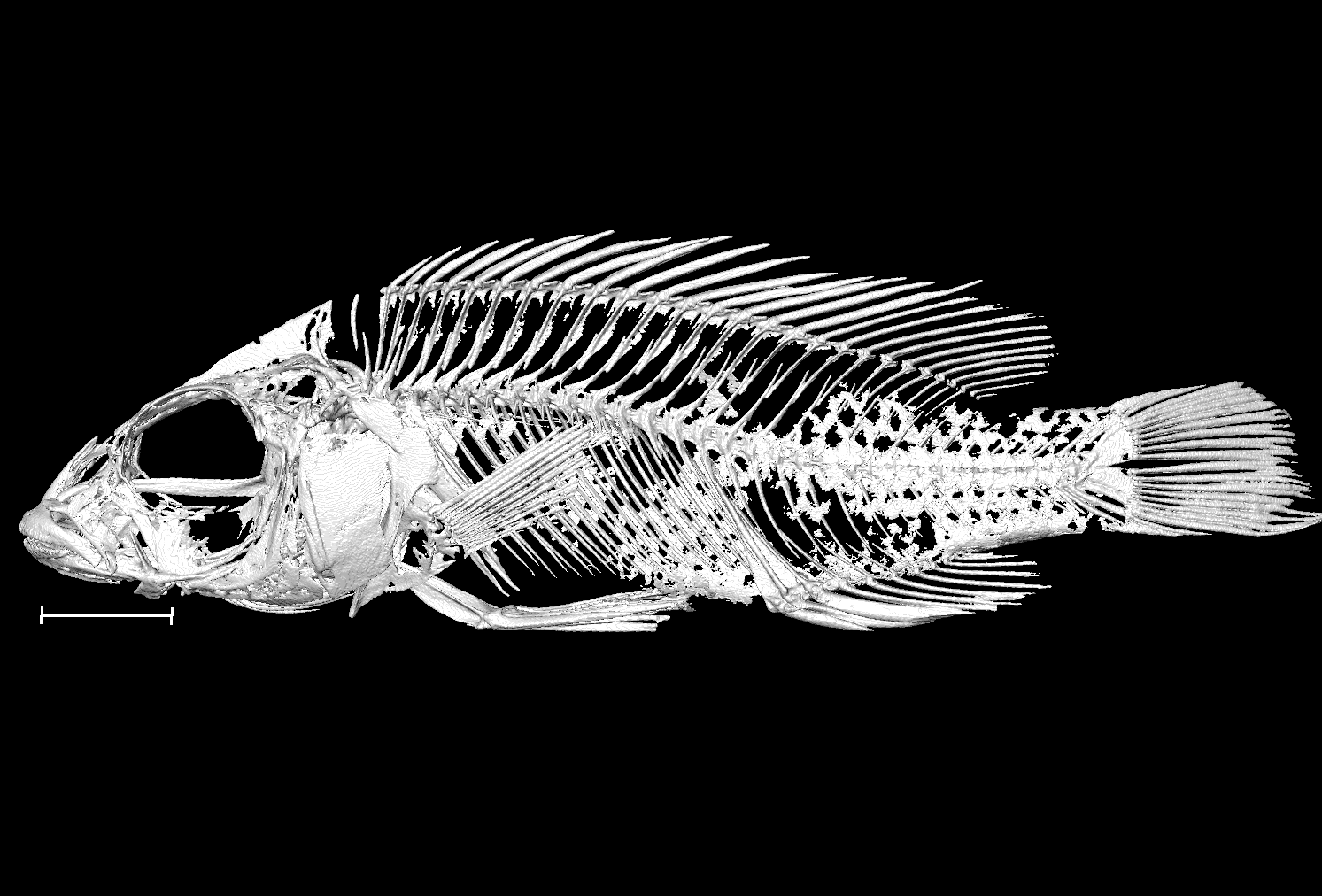

### Otopharynx_lithobates_NHMUK_1974_7_5_2_3_8bit_a.tif

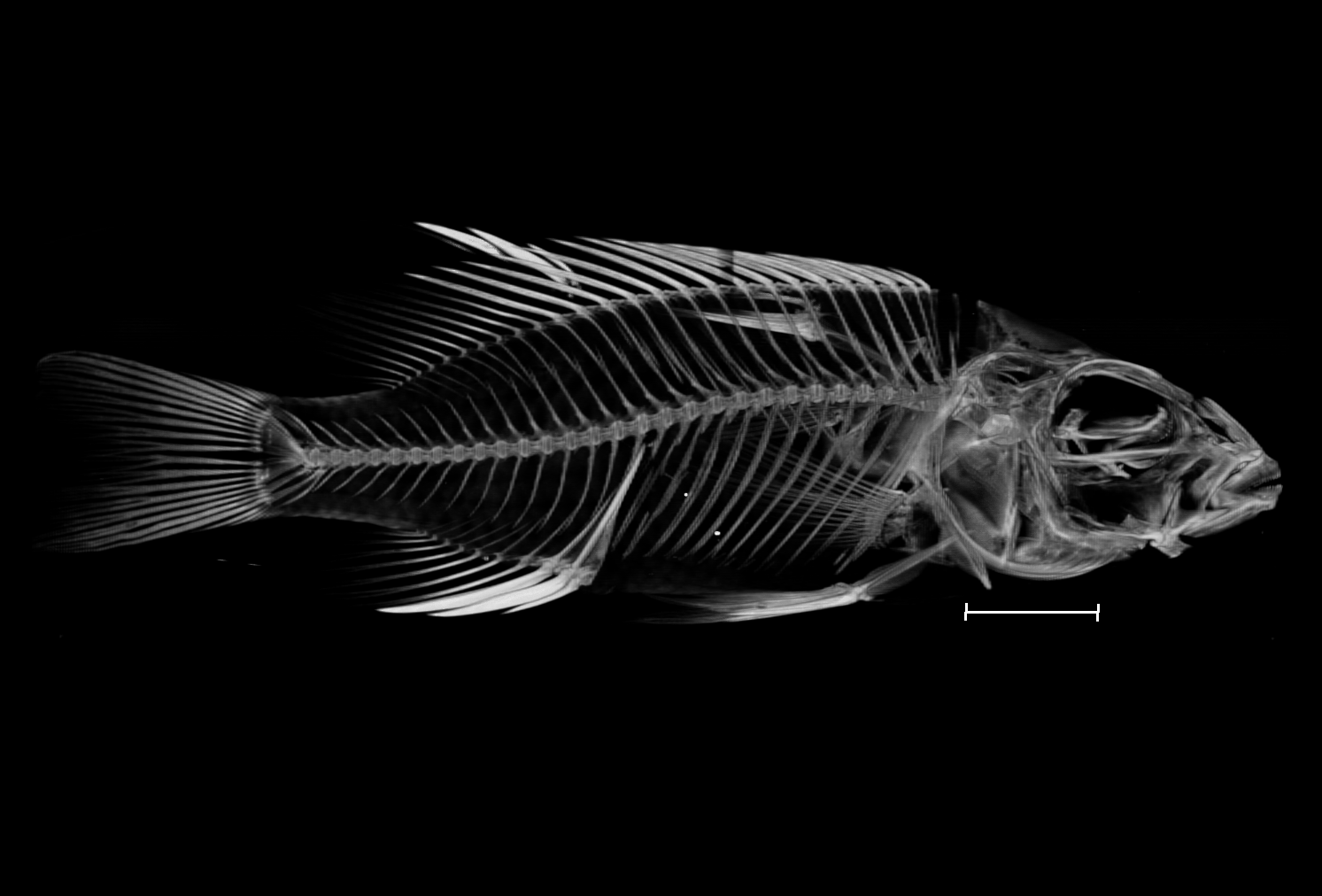

### Otopharynx_lithobates_NHMUK_1974_7_5_2_3_8bit_b.tif

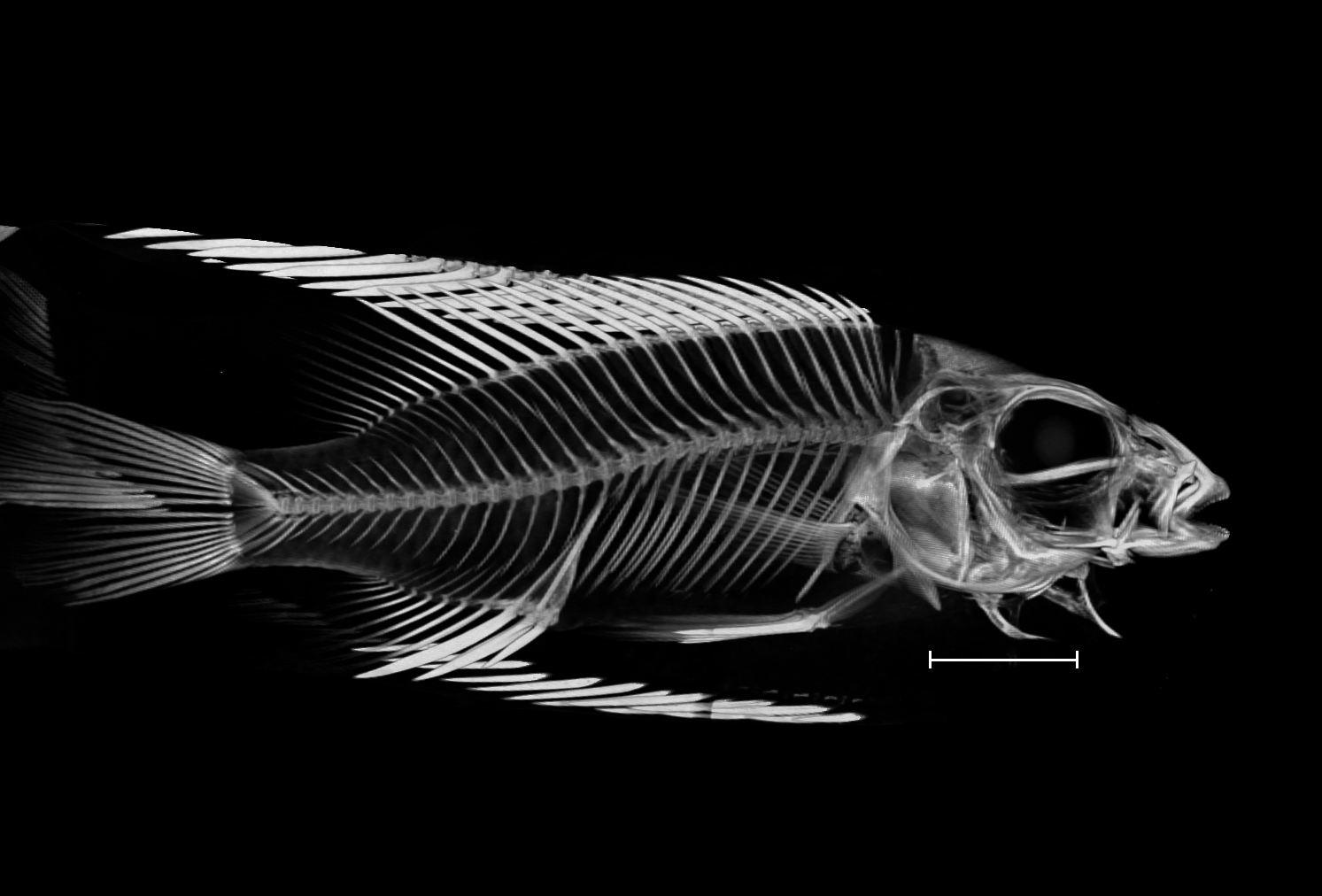

### Otopharynx_sp_brooksi_nkhata_UniBri_115_8bit.tif

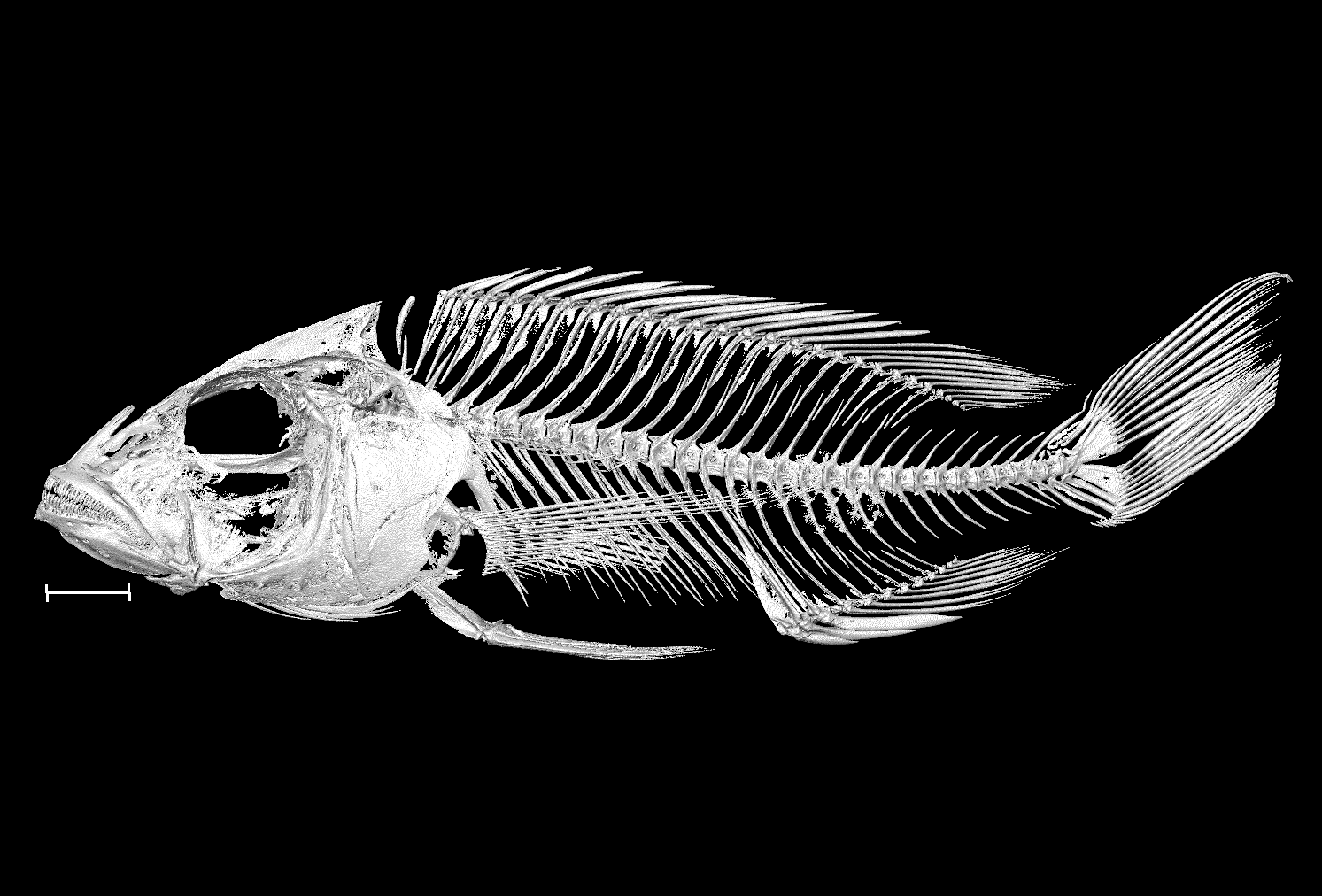

### Otopharynx_speciosus_NHMUK_1935_6_14_1649_8bit.tif

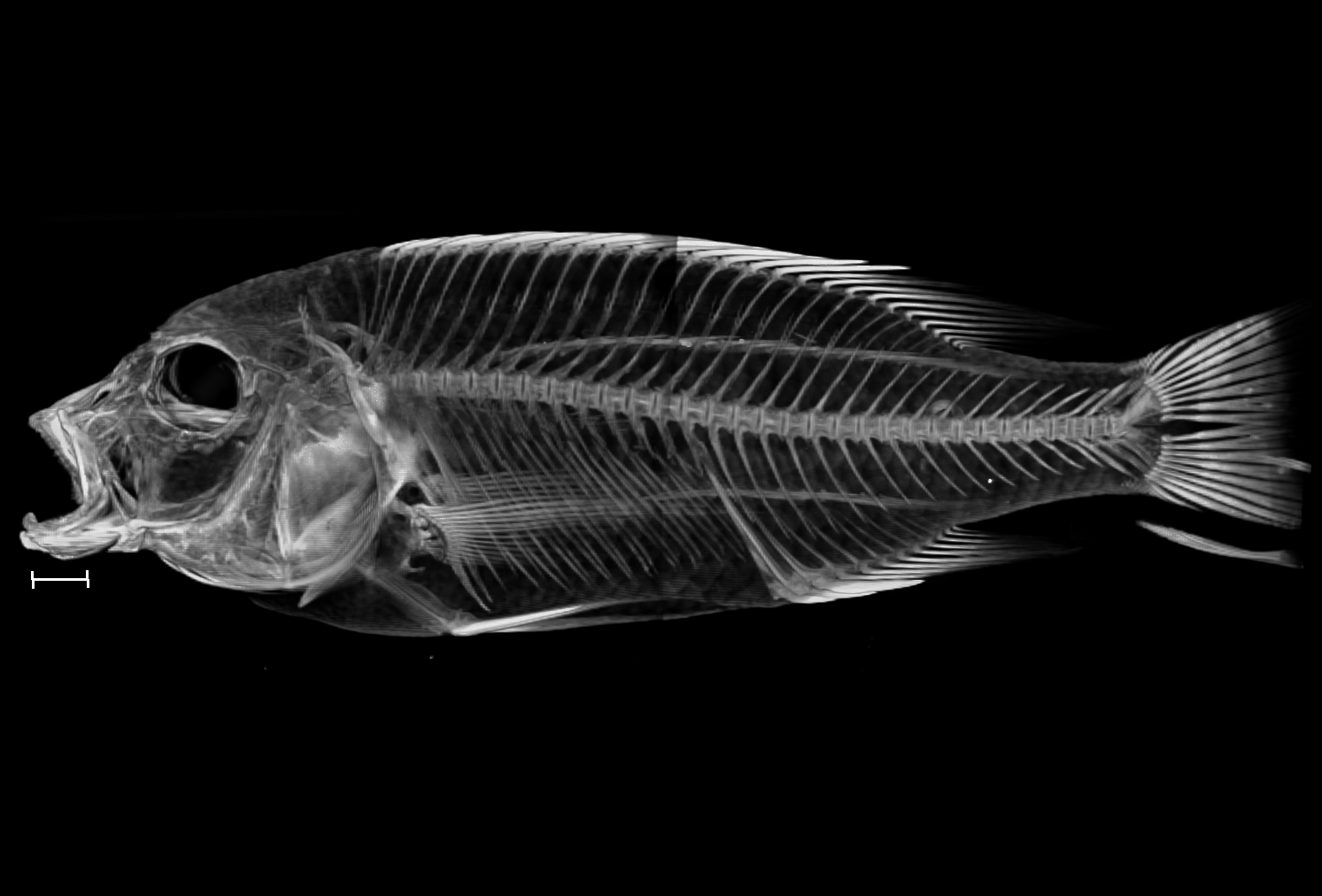

### Otopharynx_speciosus_NHMUK_1935_6_14_1650_8bit.tif

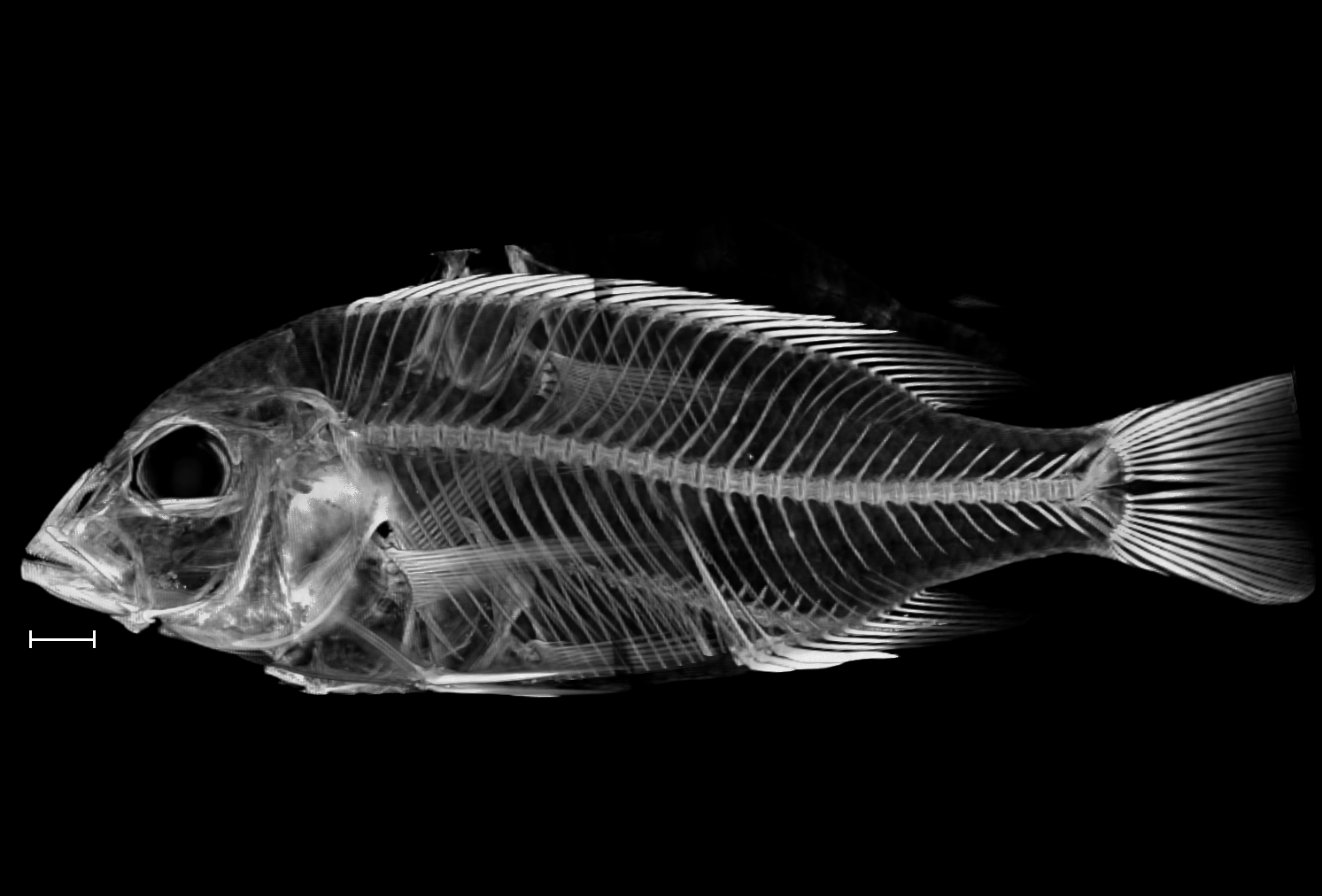

### Otopharynx_tetrastigma_NHMUK_1893_11_15_34_37_8bit_a.tif

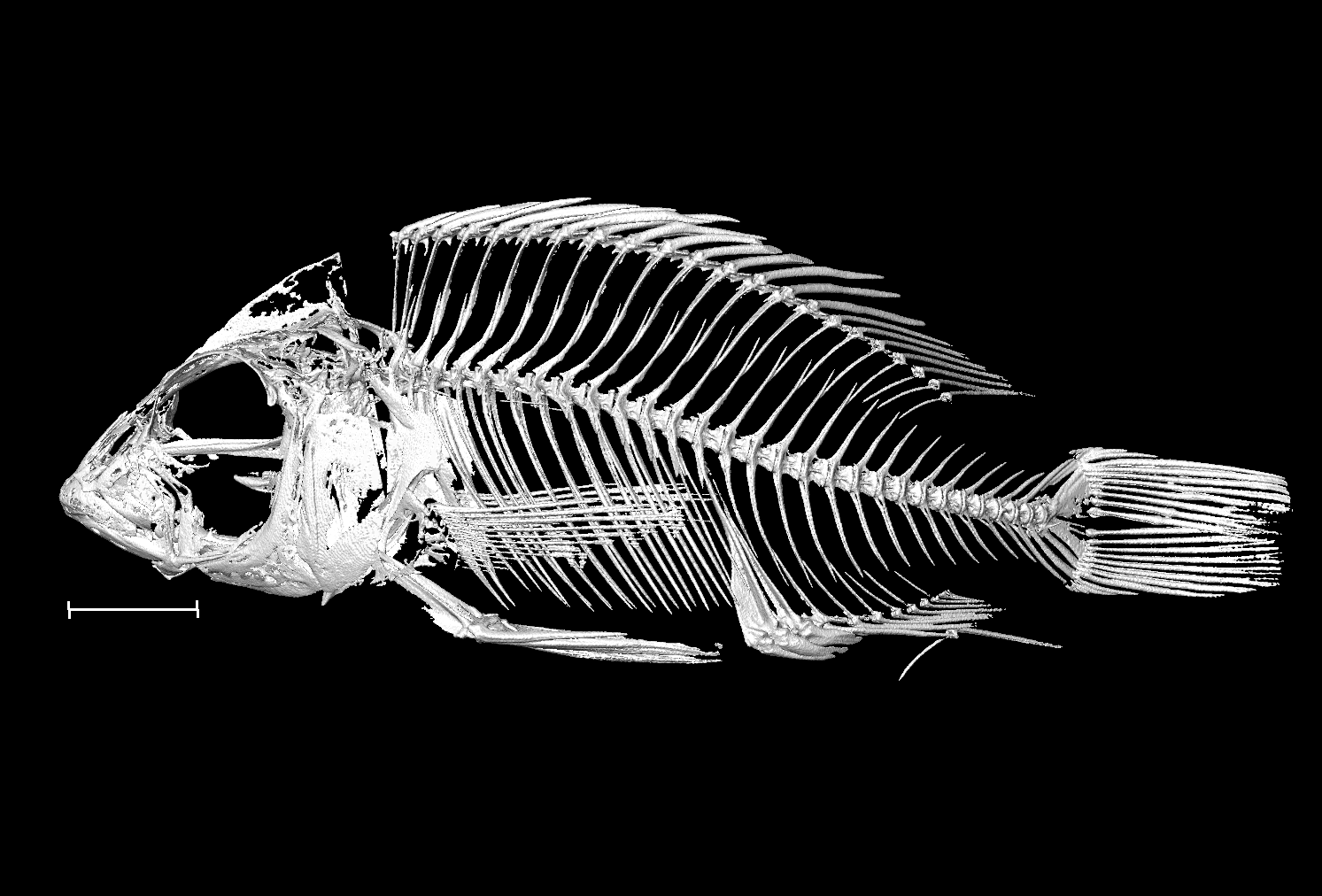

### Otopharynx_tetrastigma_NHMUK_1893_11_15_34_37_8bit_b.tif

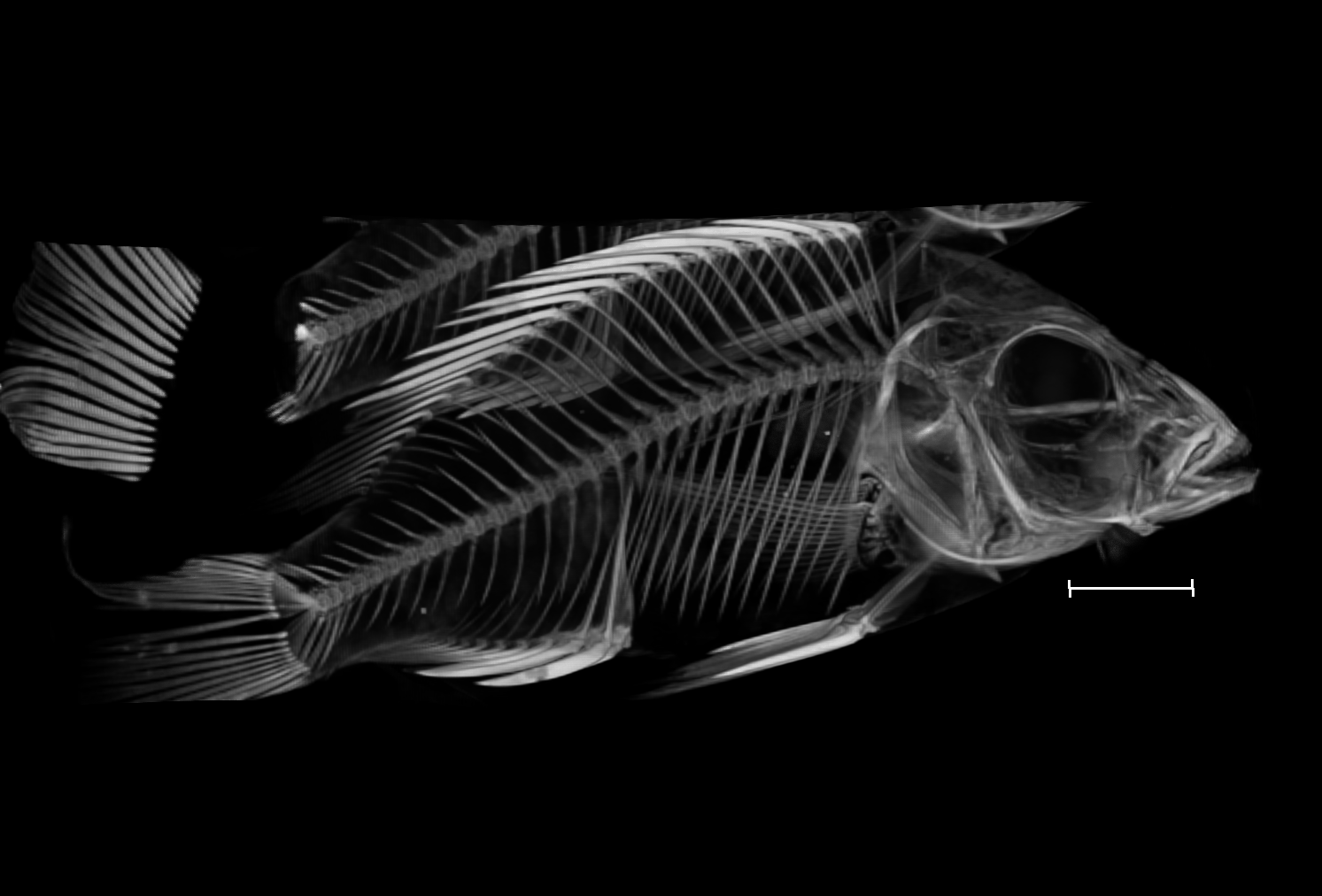

### Pallidochromis_tokolosh_NHMUK_1994_8_11_5_16_8bit_a.tif

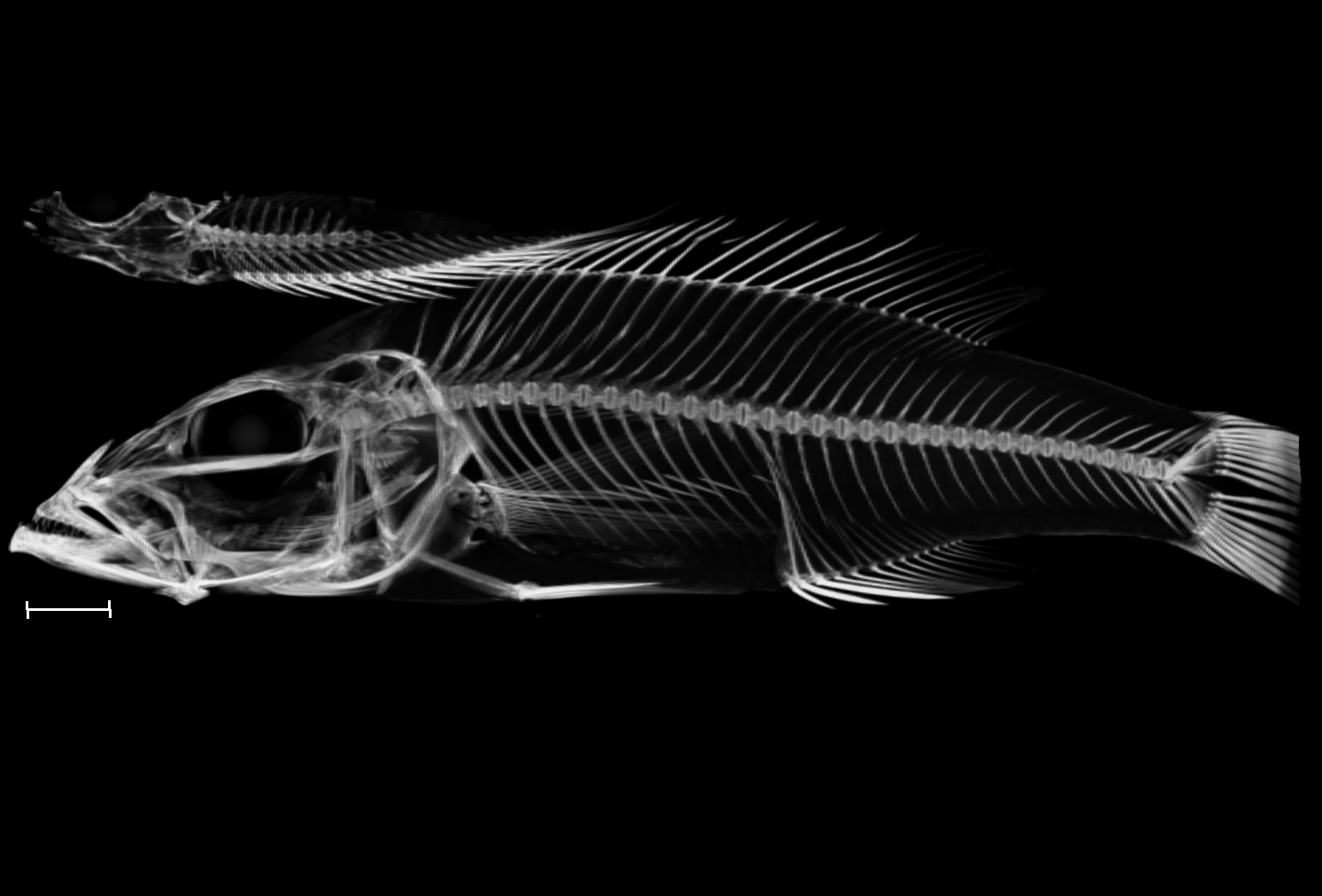

### Pallidochromis_tokolosh_NHMUK_1994_8_11_5_16_8bit_b.tif

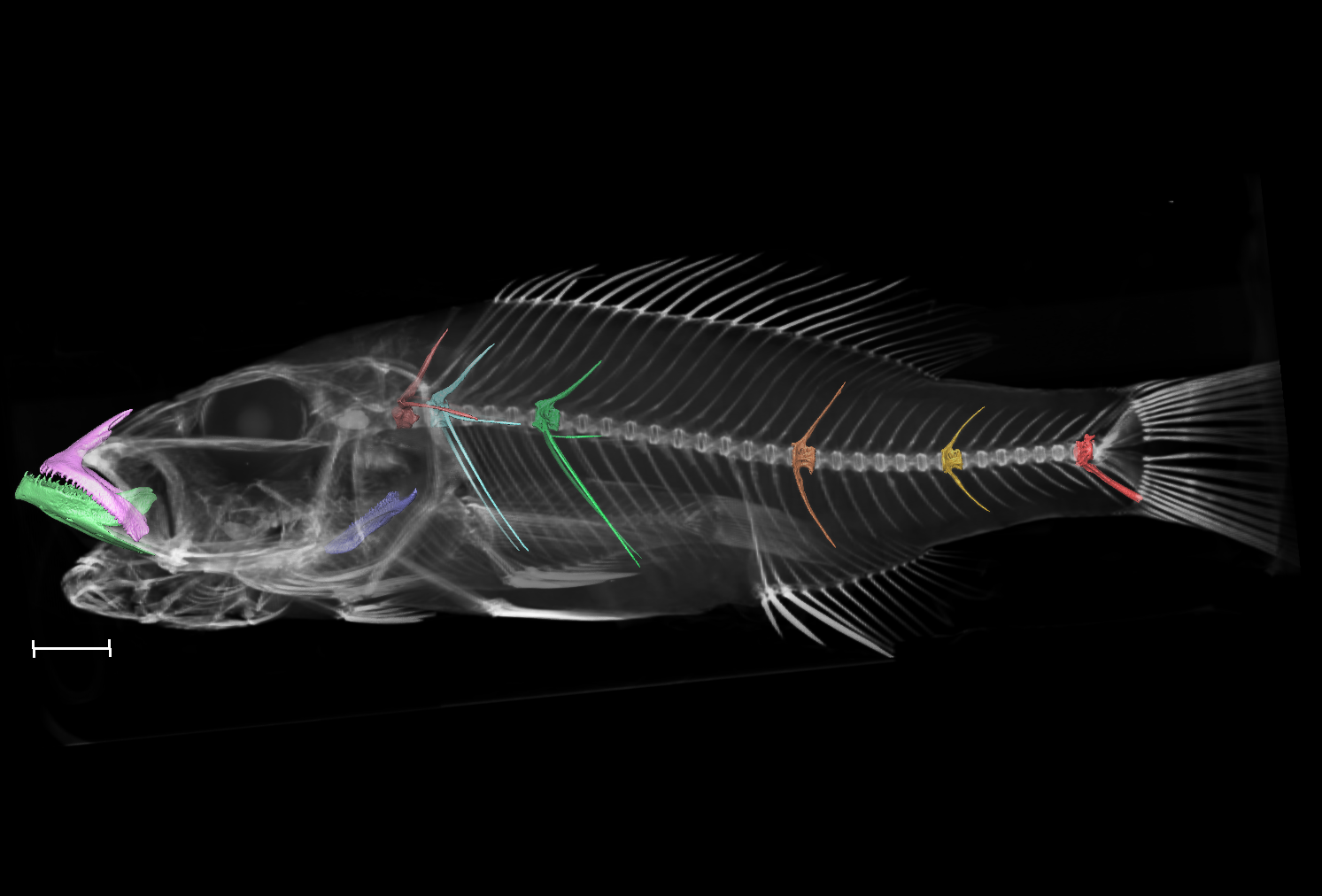

### Petrotilapia_genalutea_NHMUK_1981_2_2_222_226_8bit.tif

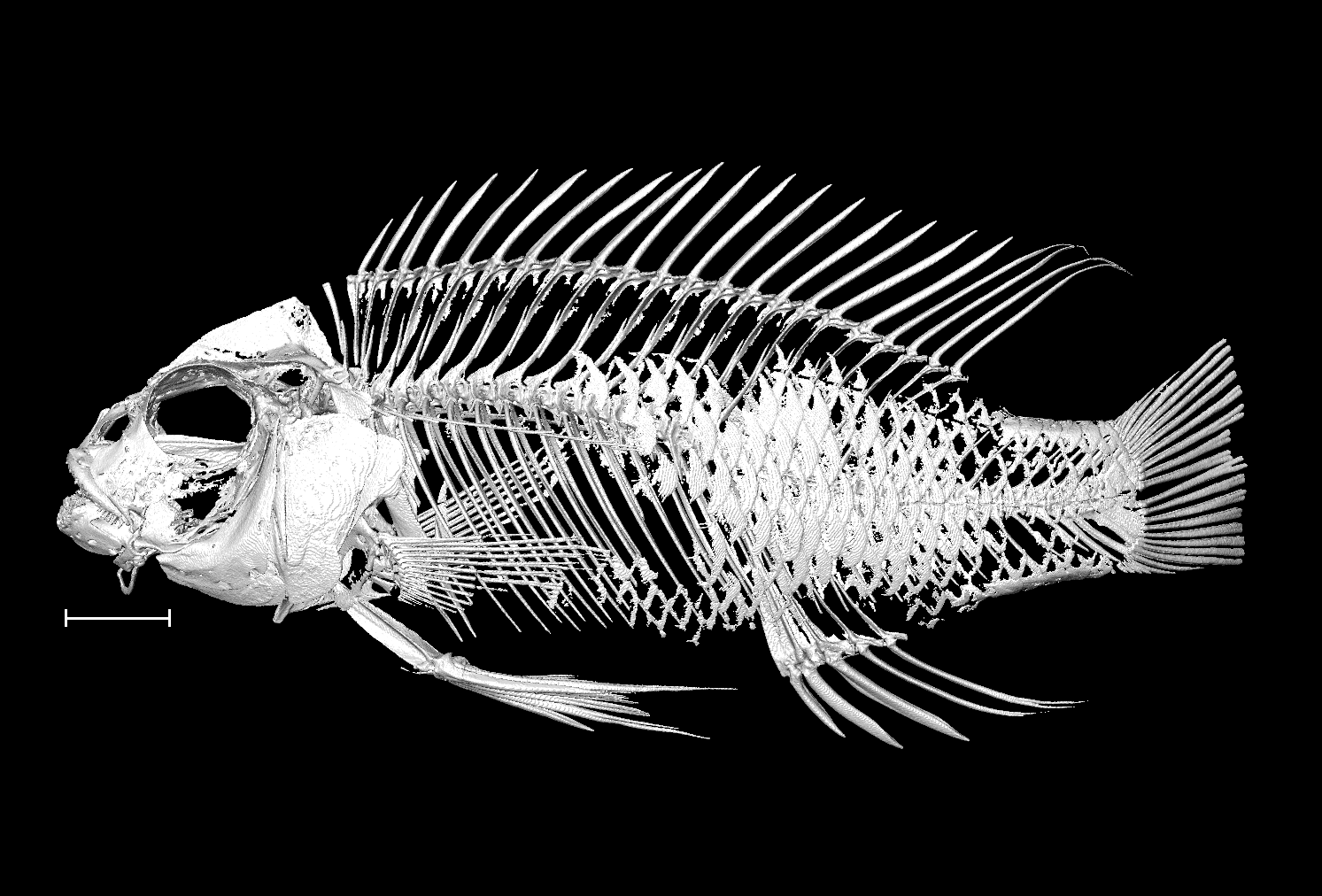

### Placidochromis_electra_UniBri_439_8bit.tif

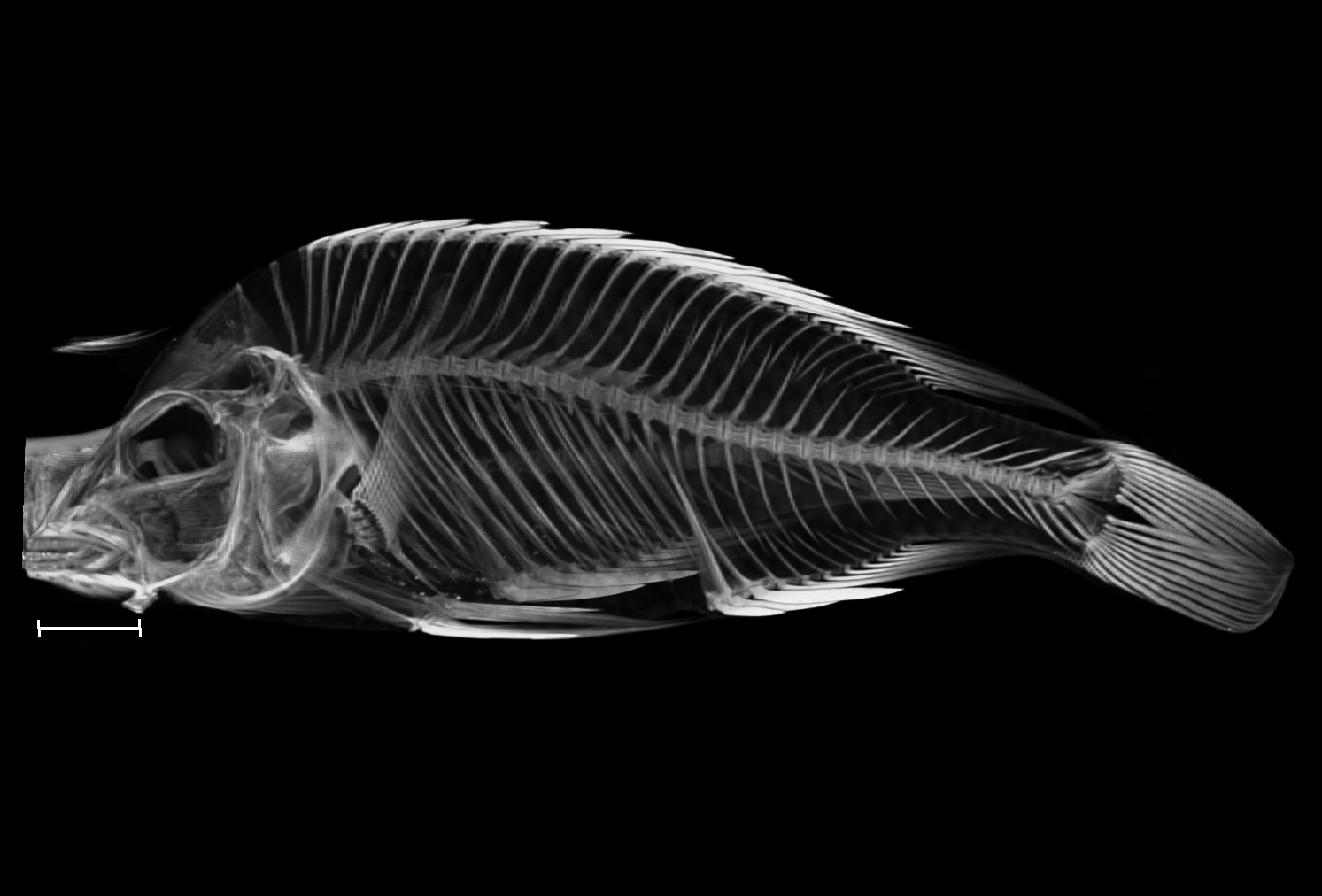

### Placidochromis_johnstoni_UniBri_438_8bit.tif

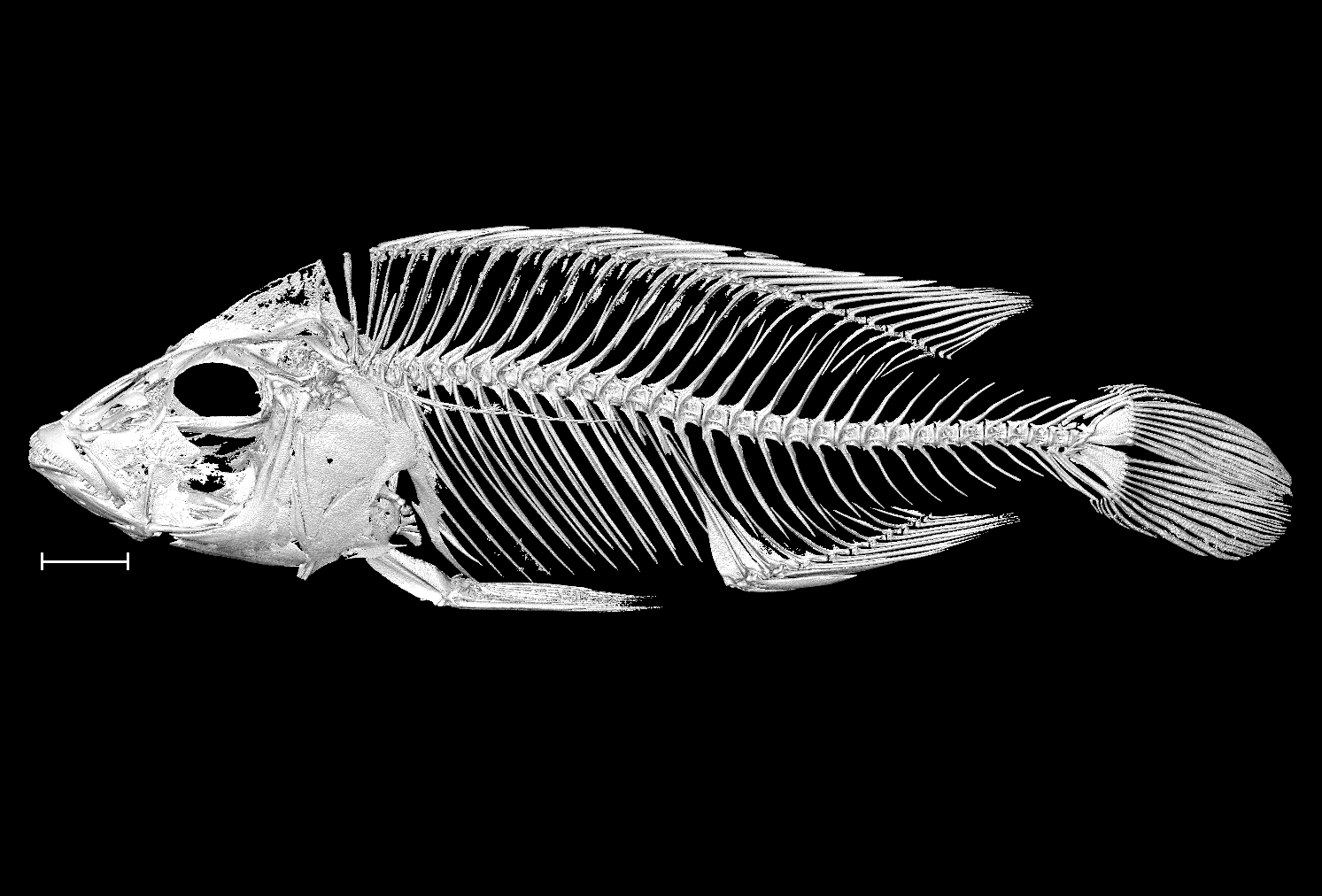

### Placidochromis_milomo_UniBri_206_8bit.tif

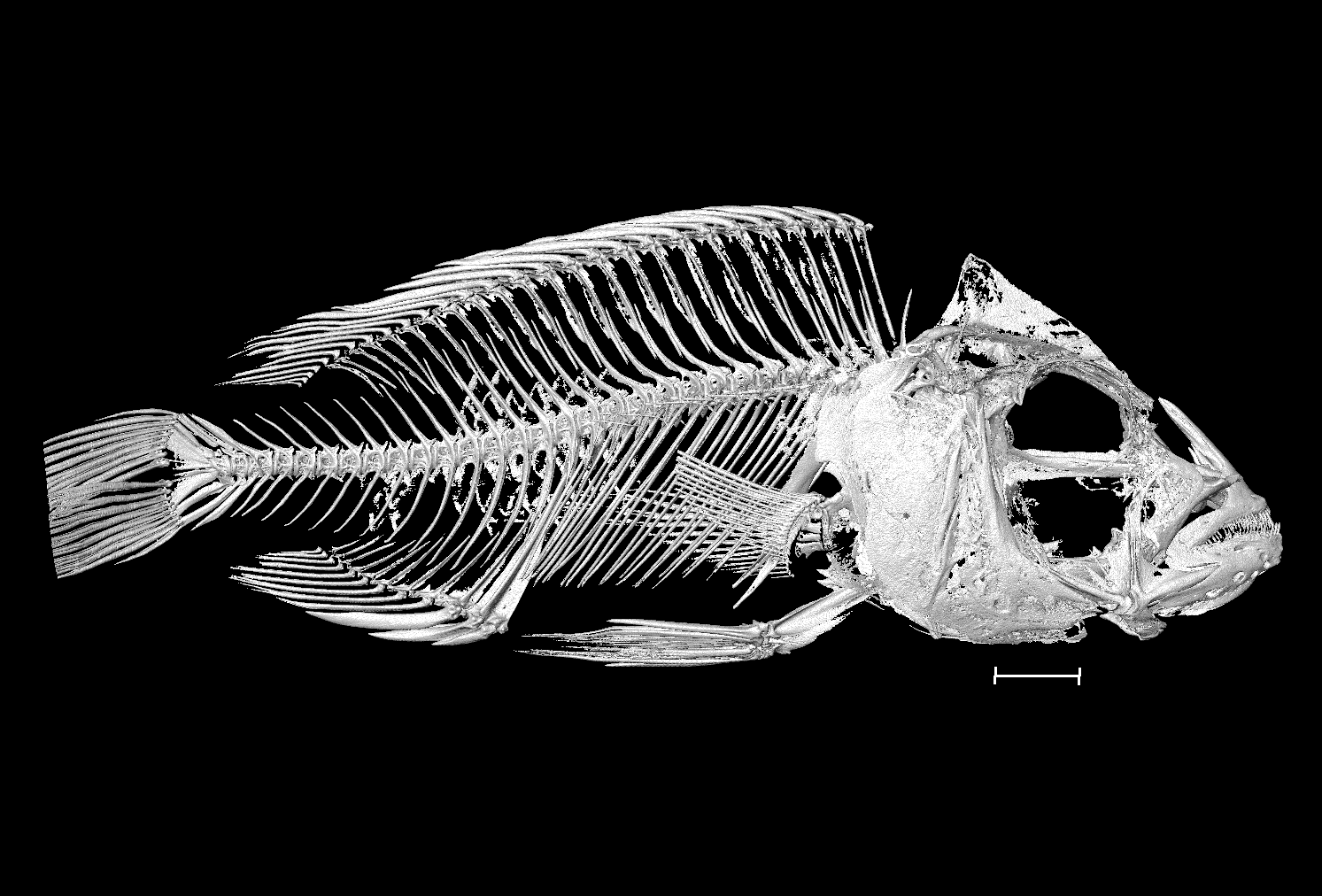

### Placidochromis_milomo_UniBri_437_8bit.tif

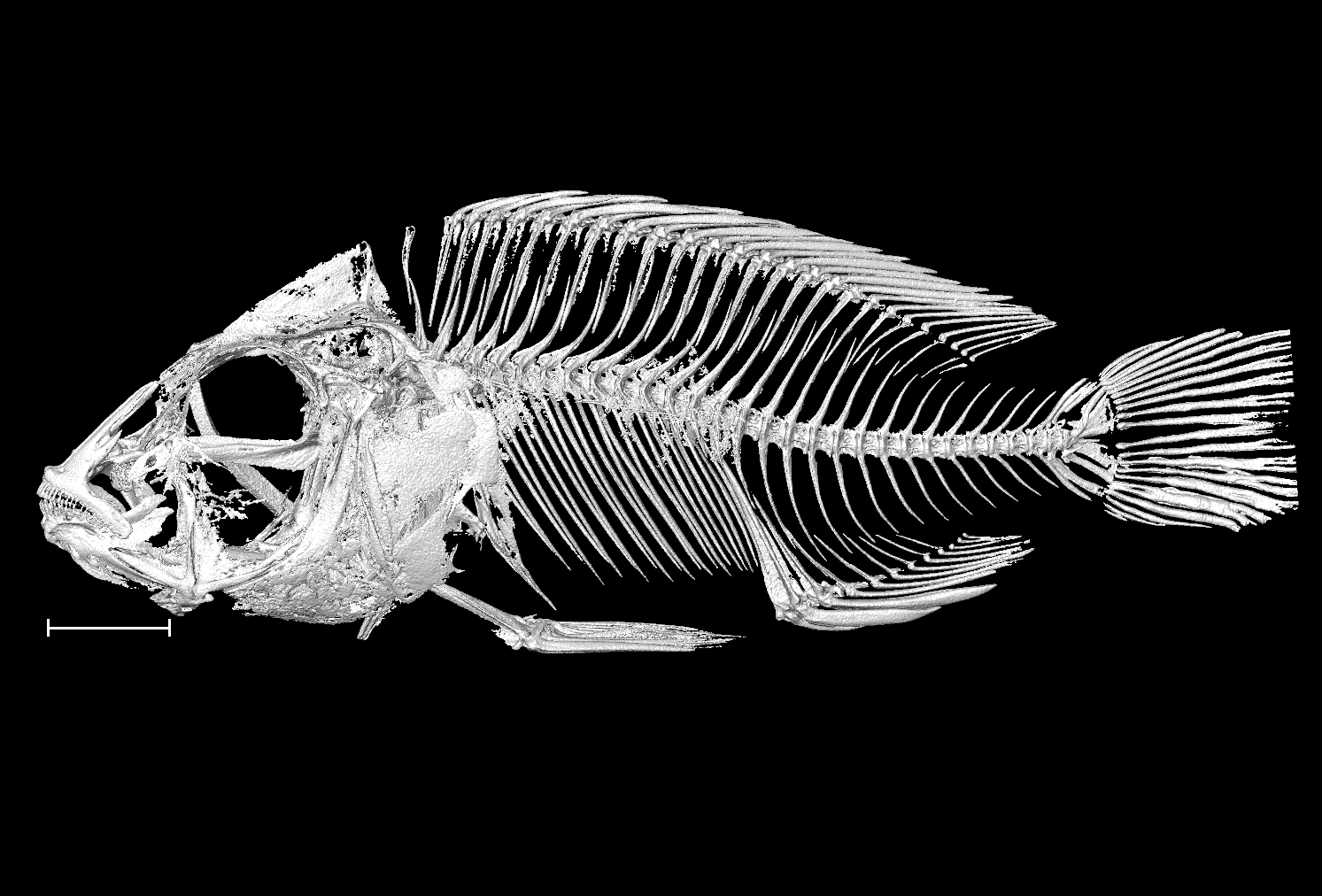

### Protomelas_spilopterus_UniBri_211_8bit.tif

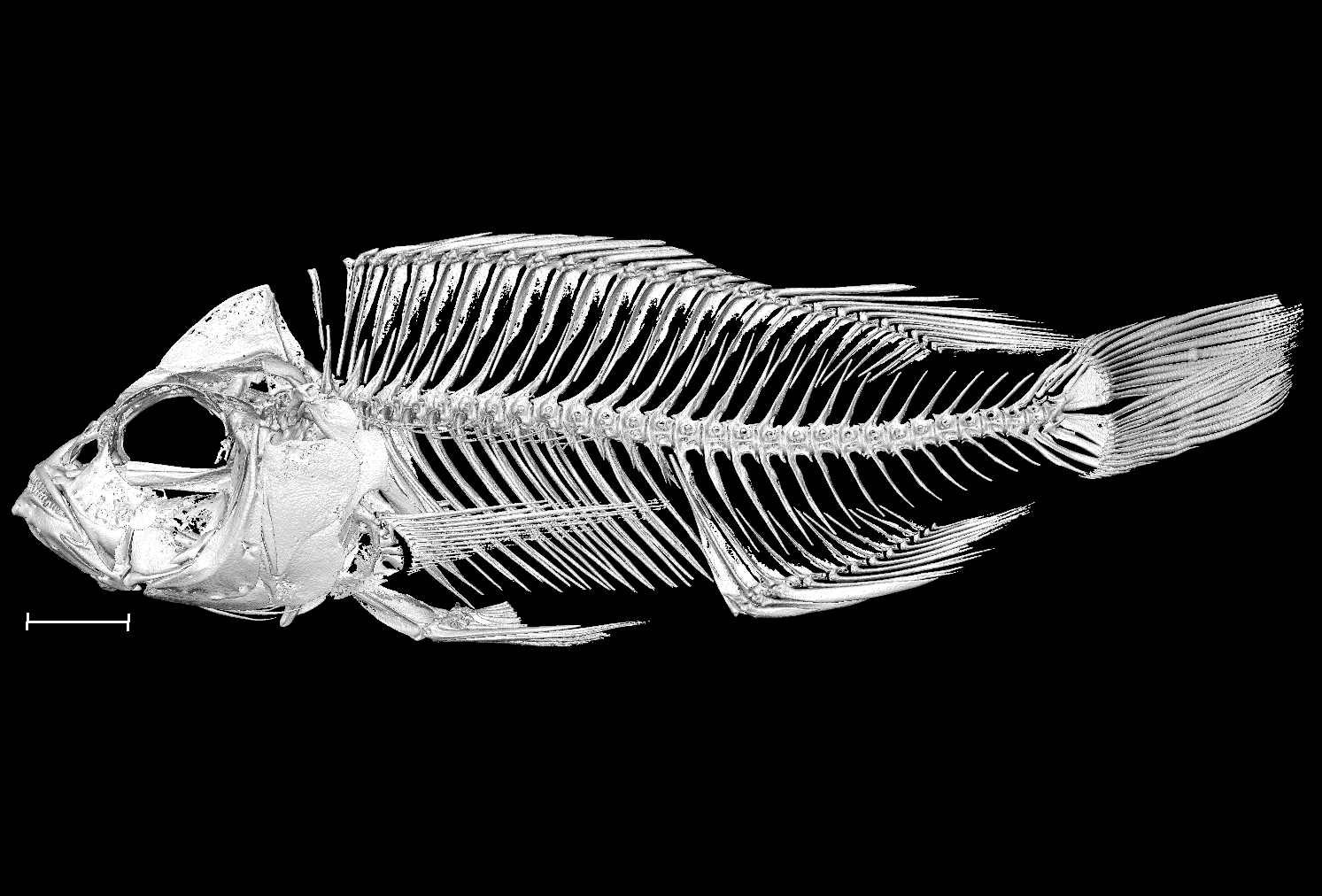

### Rhamphochromis_esox_NHMUK_1986_2_5_49_50_8bit_a.tif

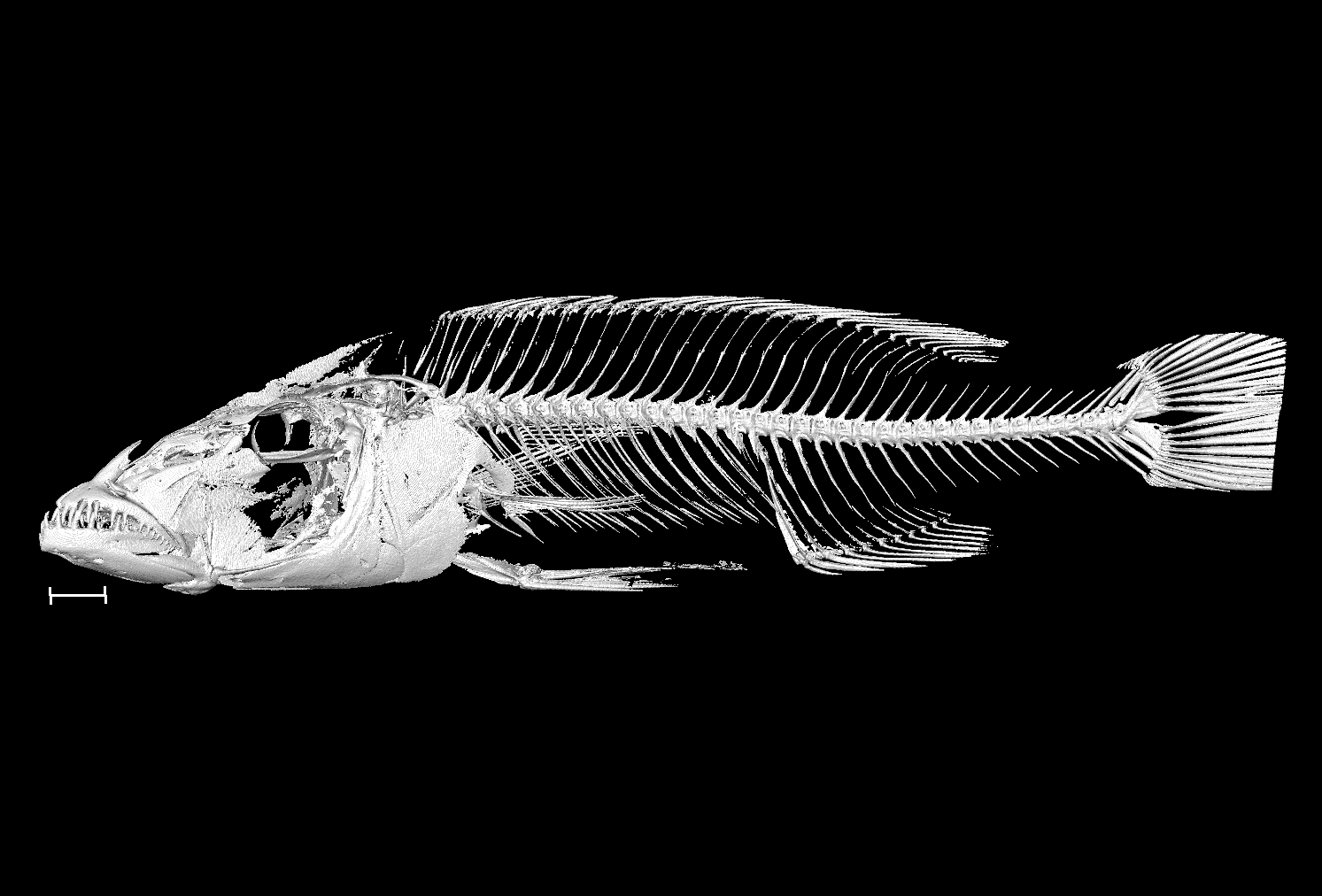

### Rhamphochromis_esox_NHMUK_1986_2_5_49_50_8bit_b.tif

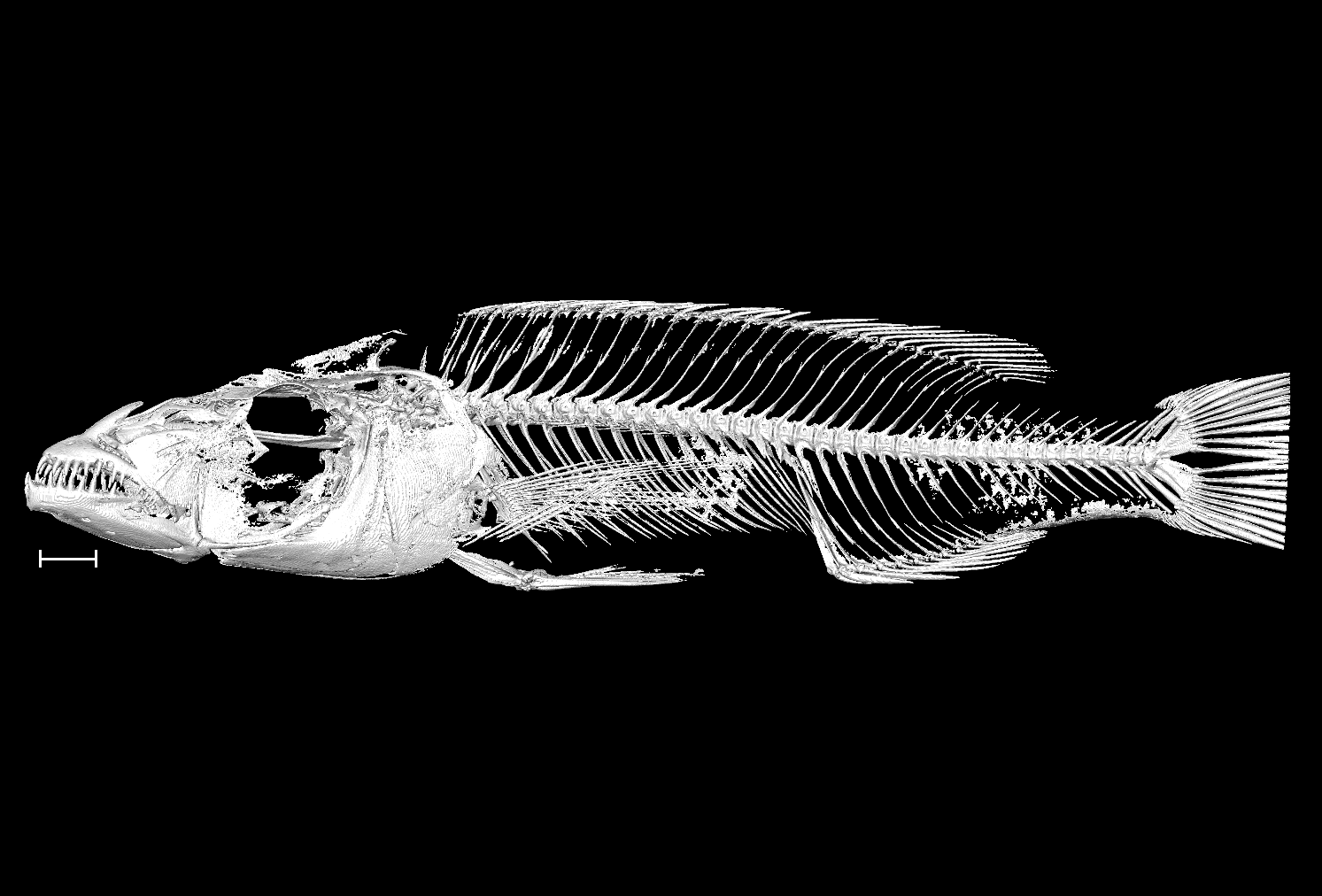

### Rhamphochromis_ferox_NHMUK_1935_6_14_2202_2207_8bit_a.tif

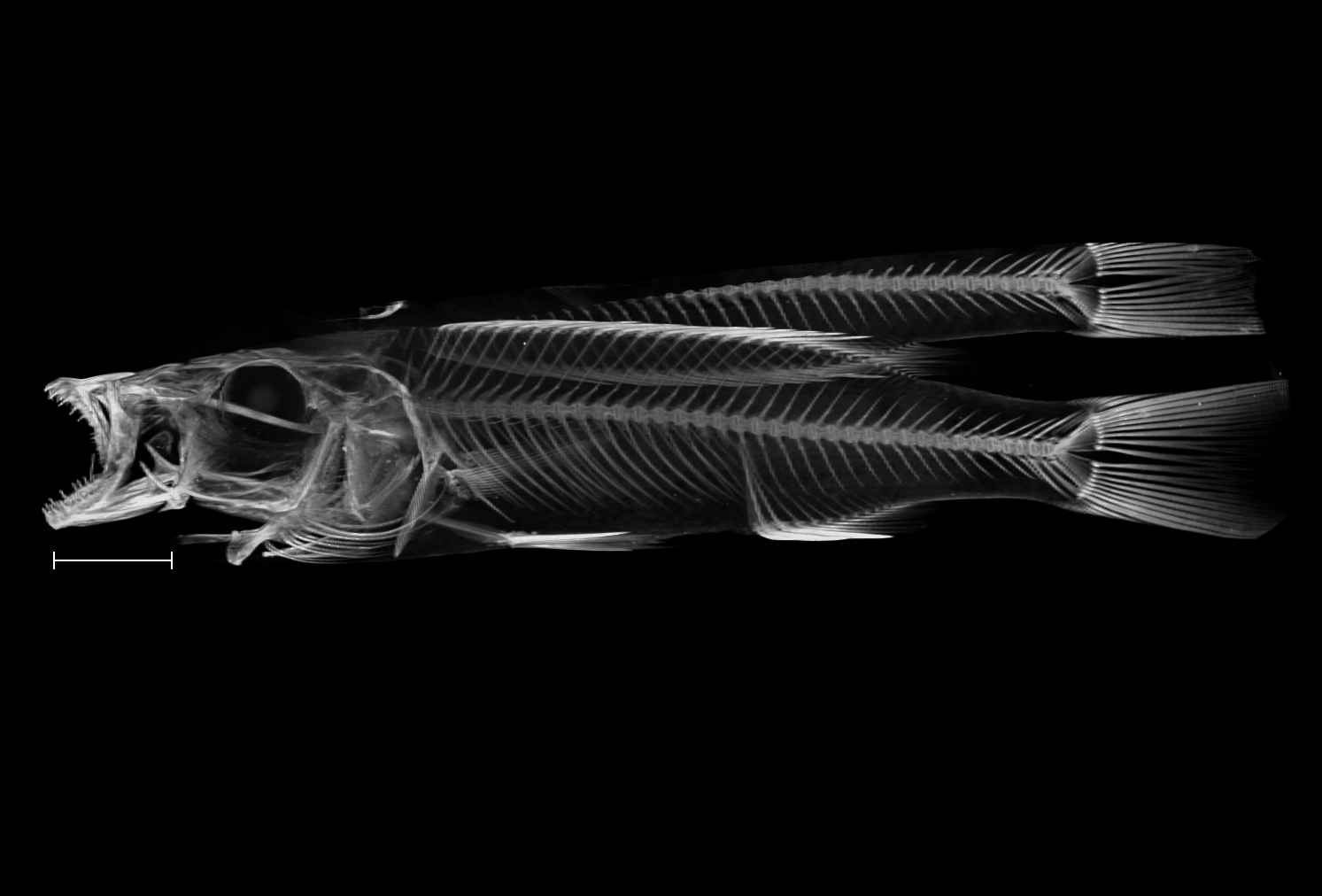

### Rhamphochromis_ferox_NHMUK_1935_6_14_2202_2207_8bit_b.tif

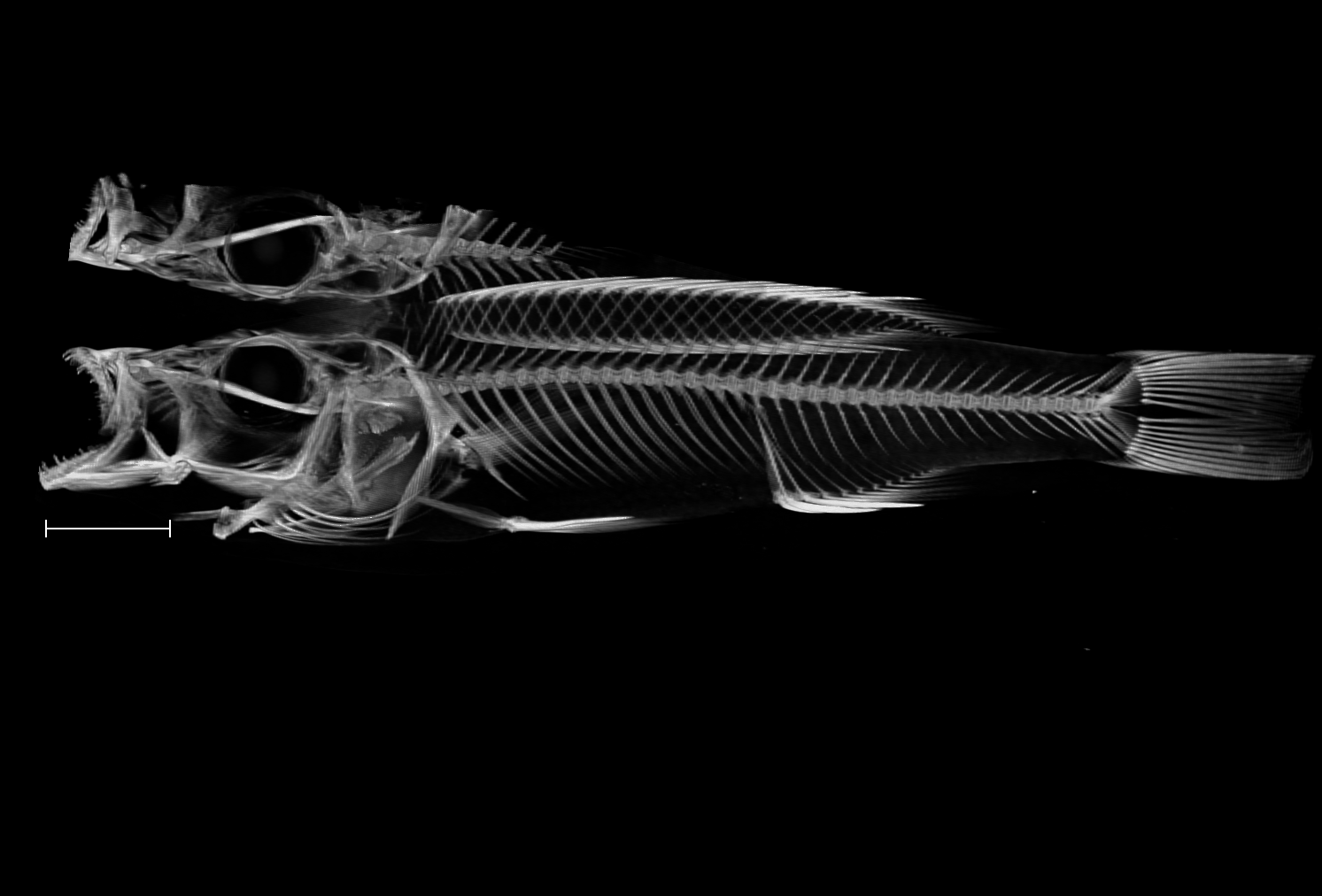

### Rhamphochromis_longiceps_NHMUK_1935_6_14_2175_2176_8bit_a.tif

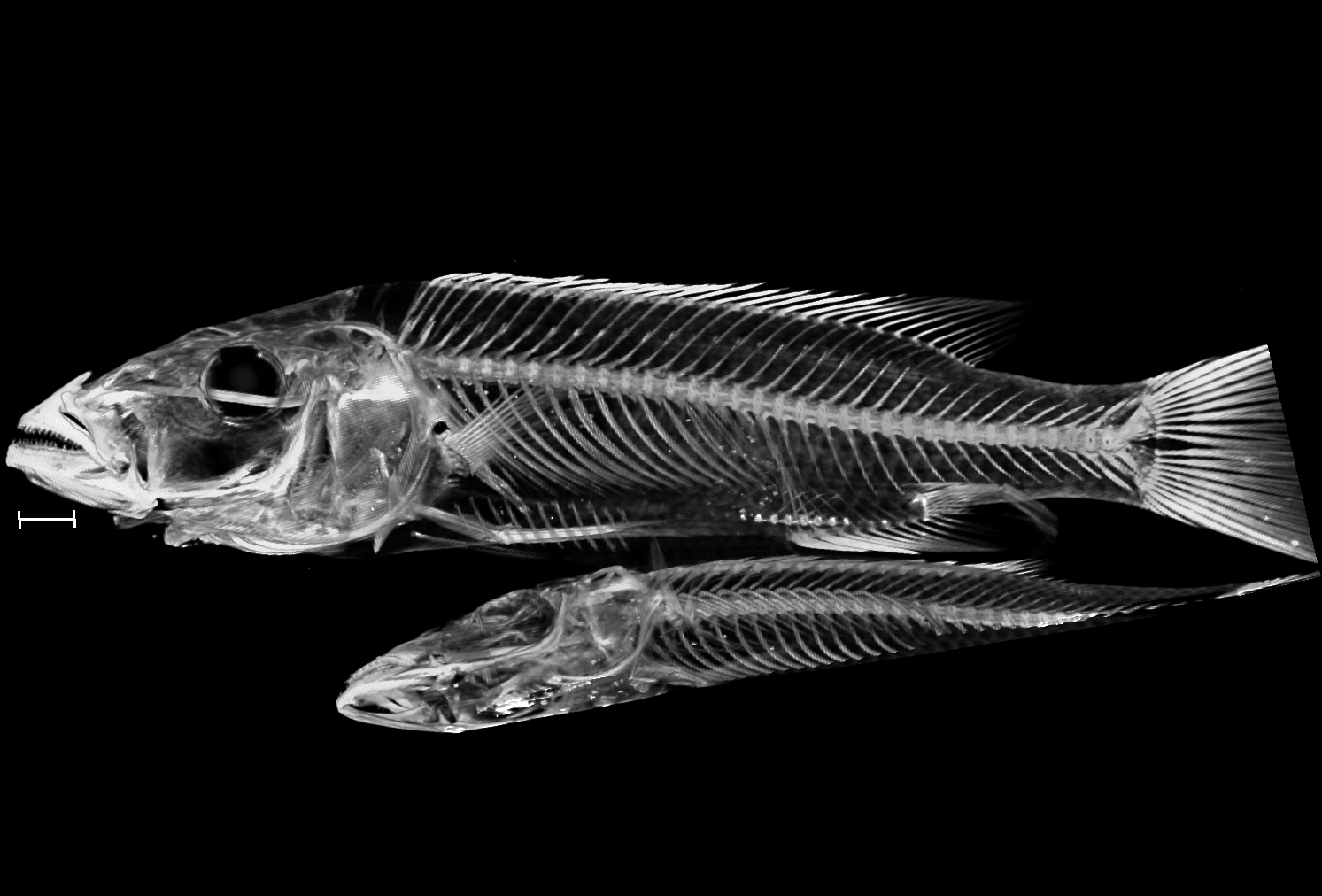

### Rhamphochromis_longiceps_NHMUK_1935_6_14_2175_2176_8bit_b.tif

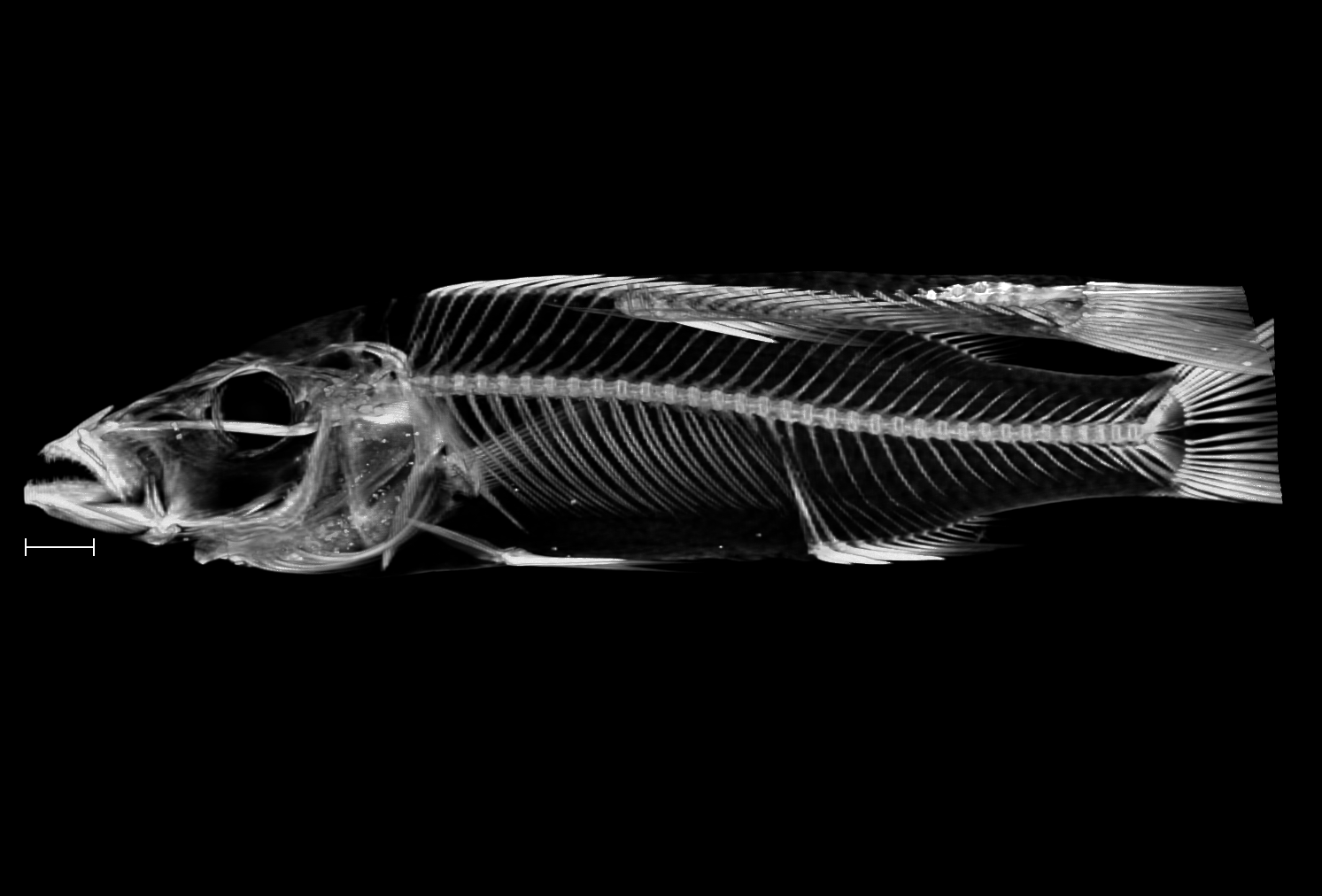

### Rhamphochromis_sp_chilingali_UniBri_RRC013_8bit_a.tif

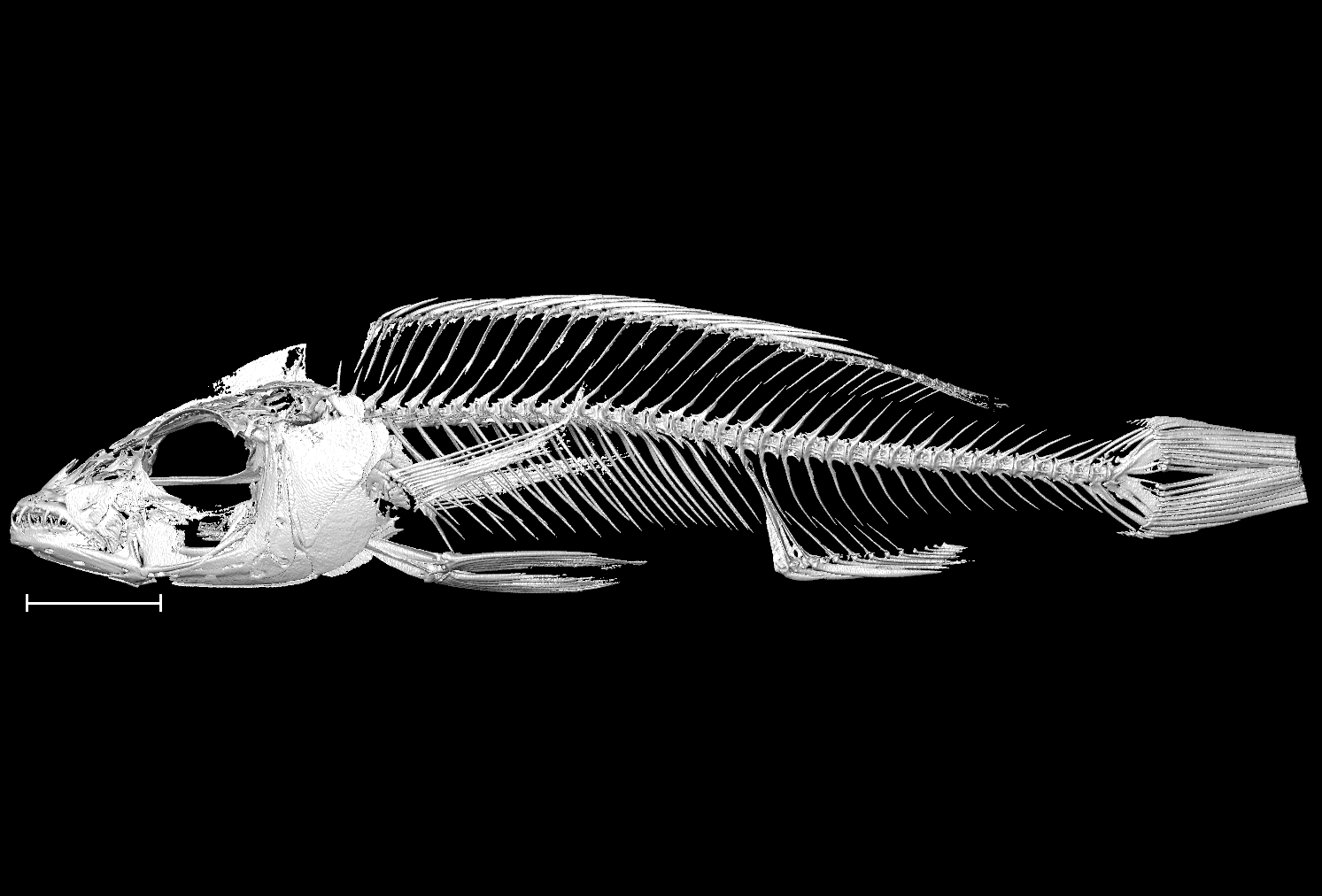

### Rhamphochromis_sp_chilingali_UniBri_RRC013_8bit_b.tif

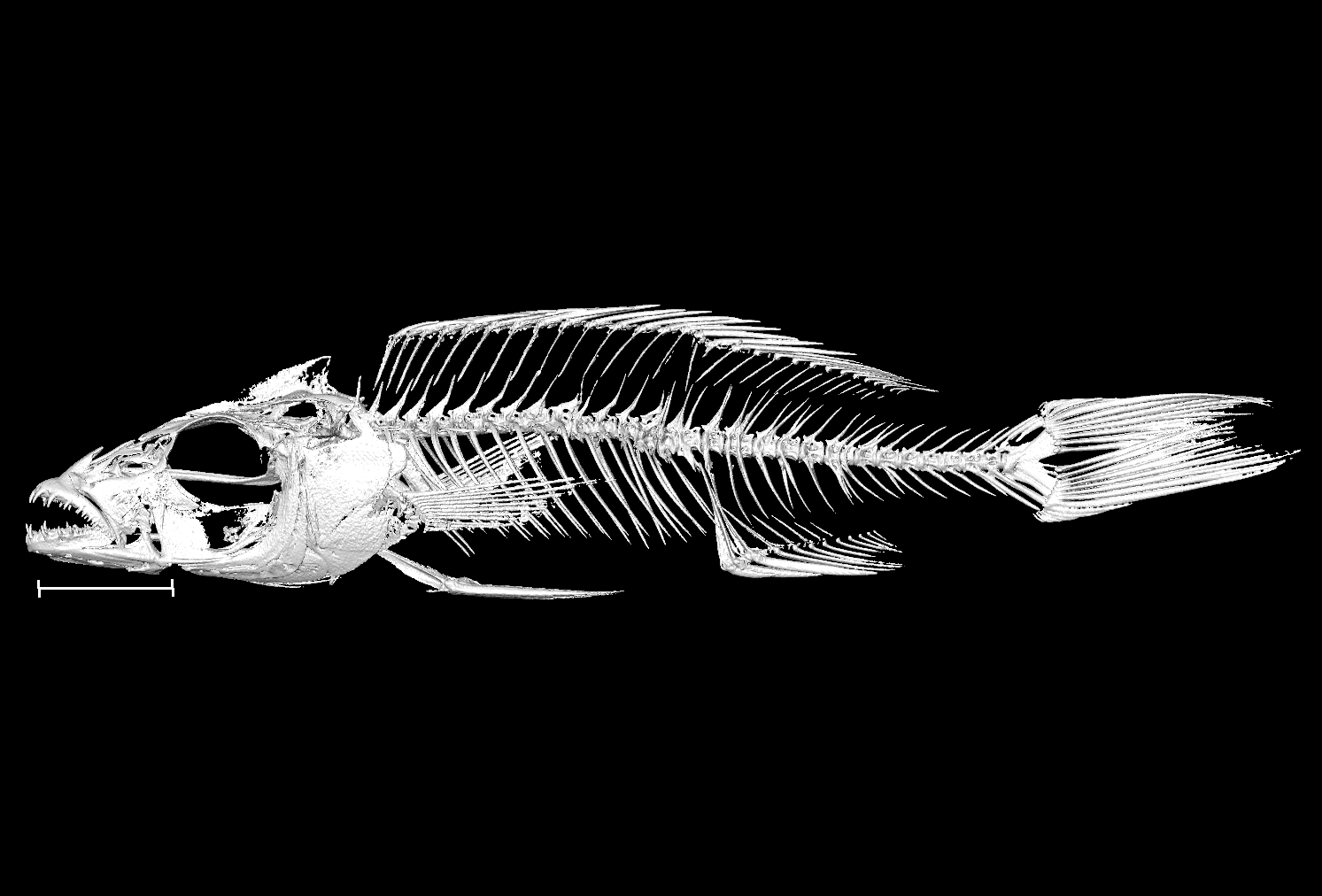

### Rhamphochromis_sp_chilingali_UniOxf_R1_8bit.tif

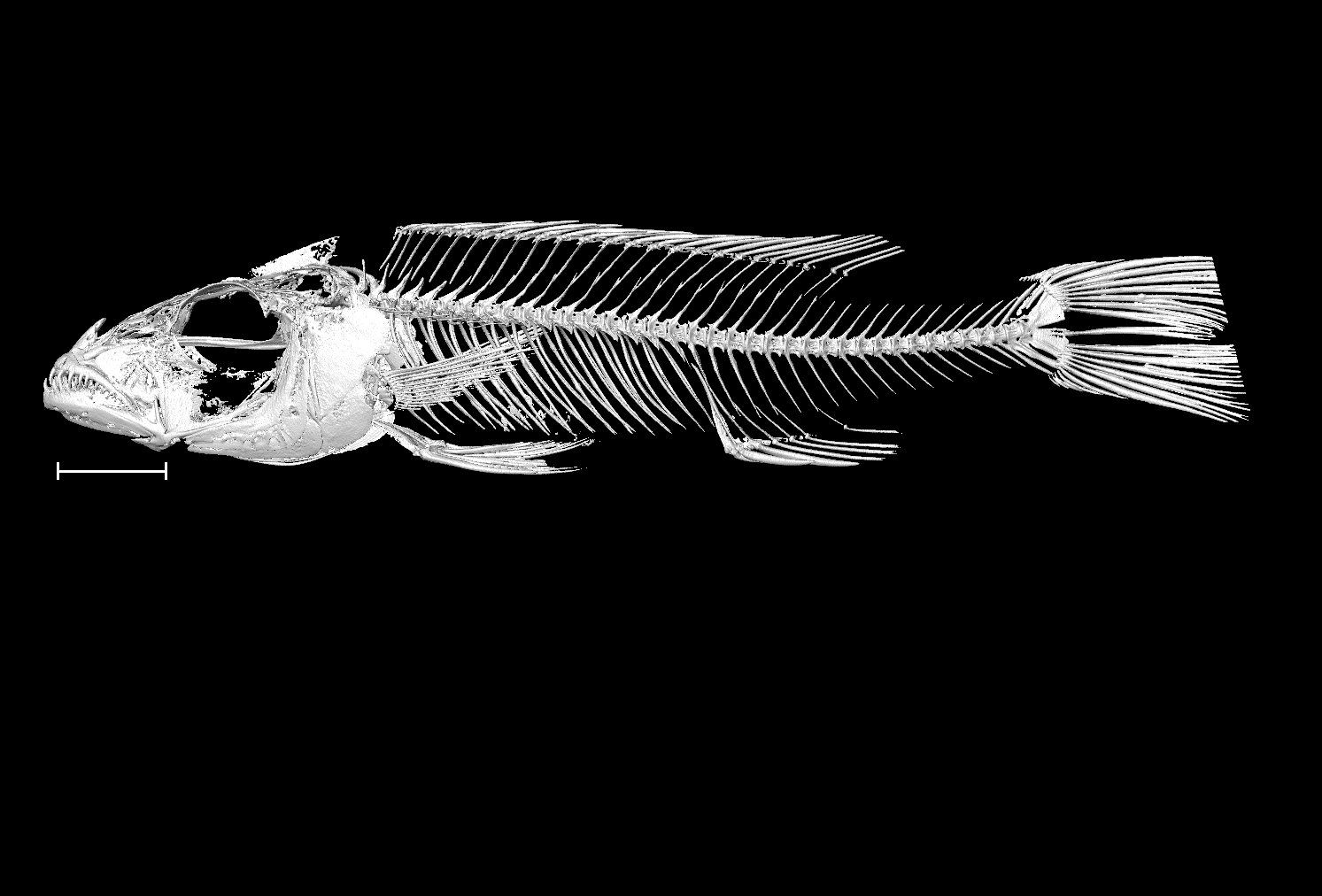

### Rhamphochromis_sp_chilingali_UniOxf_R2_8bit.tif

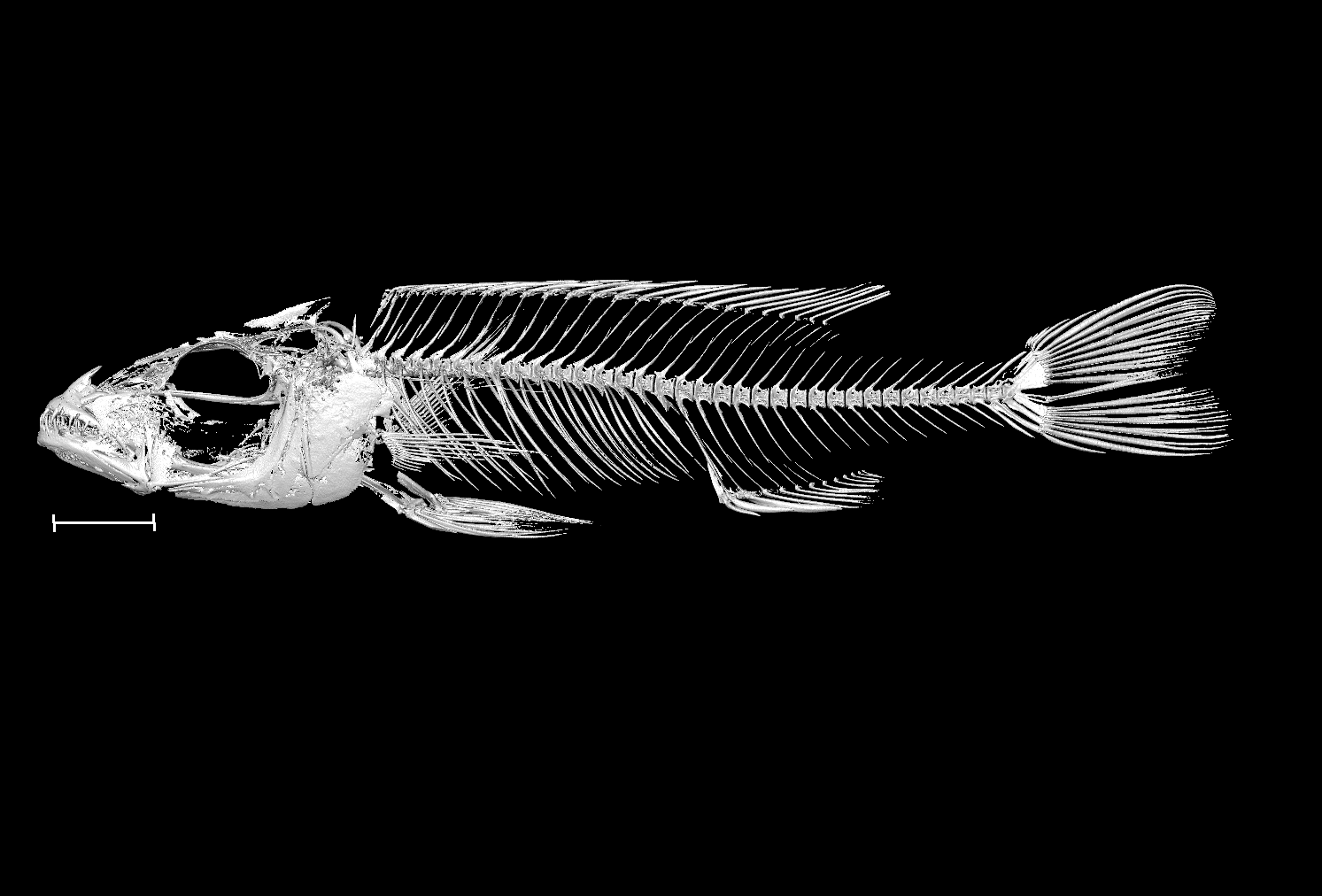

### Rhamphochromis_sp_kingiri_dwarf_UniBri_NA_8bit_a.tif

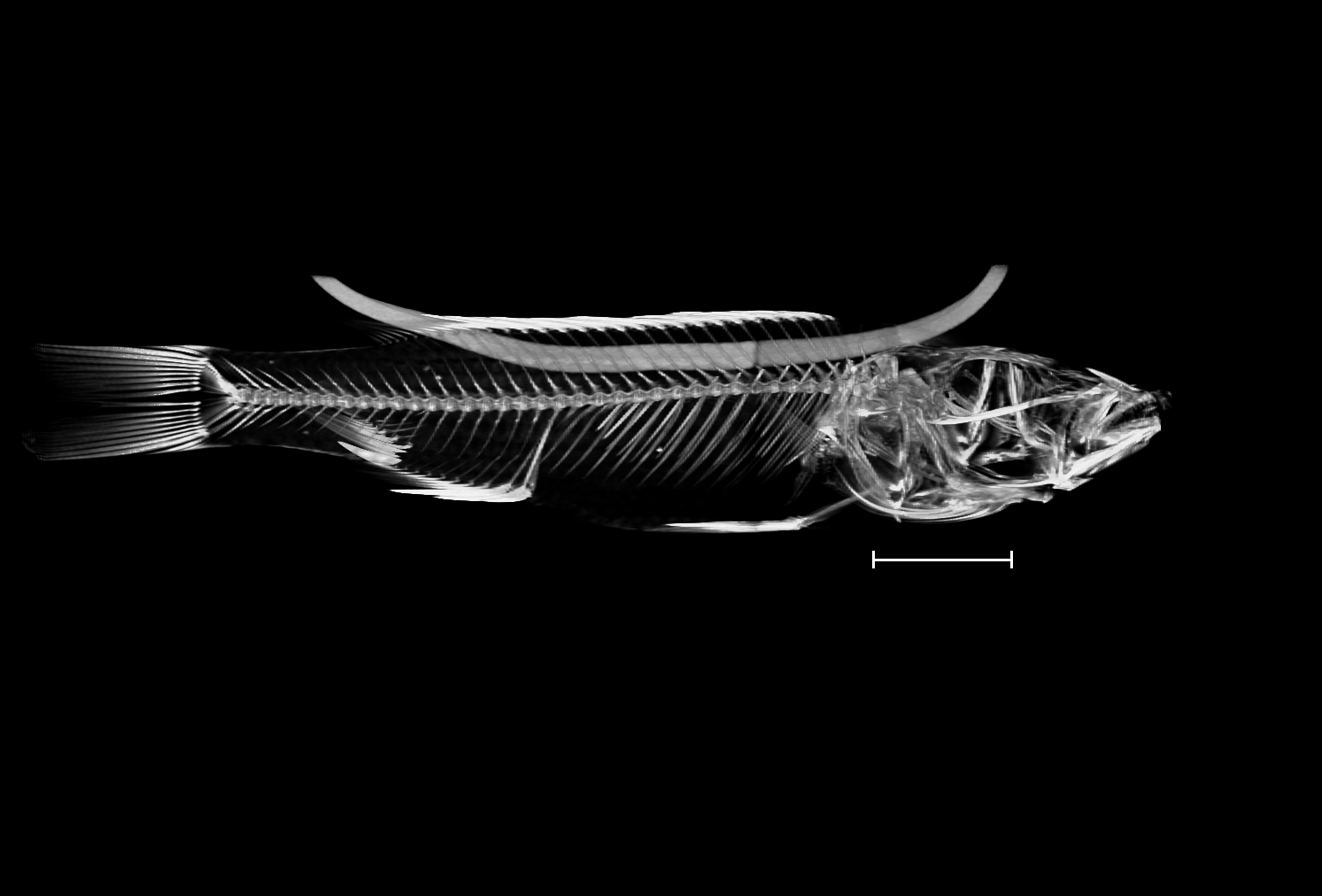

### Rhamphochromis_sp_kingiri_dwarf_UniBri_NA_8bit_b.tif

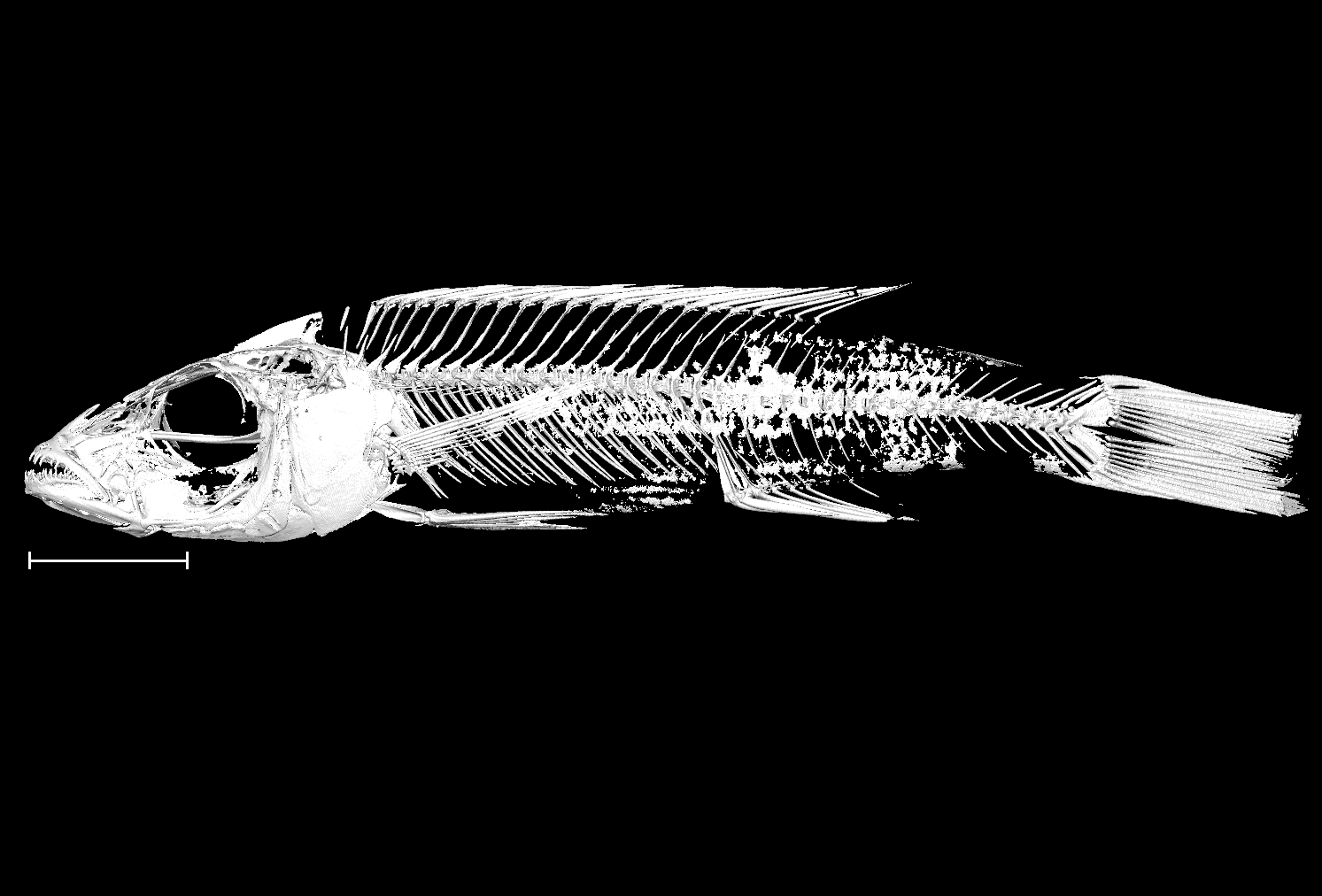
